# Glucocorticoid receptor activation reprograms NK cells to drive AREG-mediated immunosuppression in skin cancer

**DOI:** 10.1101/2023.09.13.557530

**Authors:** Qin Wei, Guirong Liang, Rui zeng, Yuancheng Li, Anlan Hong, Hongsheng Wang, Suying Feng, Yan Wang, Yetao Wang

## Abstract

Natural killer (NK) cells are potent mediators of anti-tumor immunity, yet their functions are frequently subverted by tumor microenvironment-driven immunosuppression. Here, we dissect the molecular mechanisms underlying NK cell dysfunction in cutaneous malignancies and identify a paradoxical cytokine shift in tumor-associated NK cells-reduced production of IFN-γ and TNF-α alongside elevated amphiregulin (AREG), an EGFR ligand linked to tumor progression. Single-cell transcriptomic analysis indicates that this reprogramming correlates with elevated glucocorticoid receptor (GR/NR3C1) pathway activity in tumor-infiltrating NK cells. Functional validation demonstrated that glucocorticoids specifically induce AREG production in NK cells, with tumor-associated prostaglandin E2 (PGE2) augmenting this response. Genetic ablation or pharmacological inhibition of NR3C1 abolished glucocorticoid-driven AREG induction. Moreover, primary GR activation established persistent chromatin accessibility at the AREG locus, sensitizing NK cells to enhanced AREG production upon secondary glucocorticoid exposure. Functionally, AREG counteracts NK cell-mediated tumor apoptosis, while adoptive transfer of AREG-deficient human NK cells significantly suppressed melanoma, cutaneous squamous cell carcinoma (cSCC), and hepatocellular carcinoma growth in NCG mice. These findings establish the GR-AREG axis as a multi-layered therapeutic target for restoring NK cell anti-tumor function.

**Graphical abstract:** 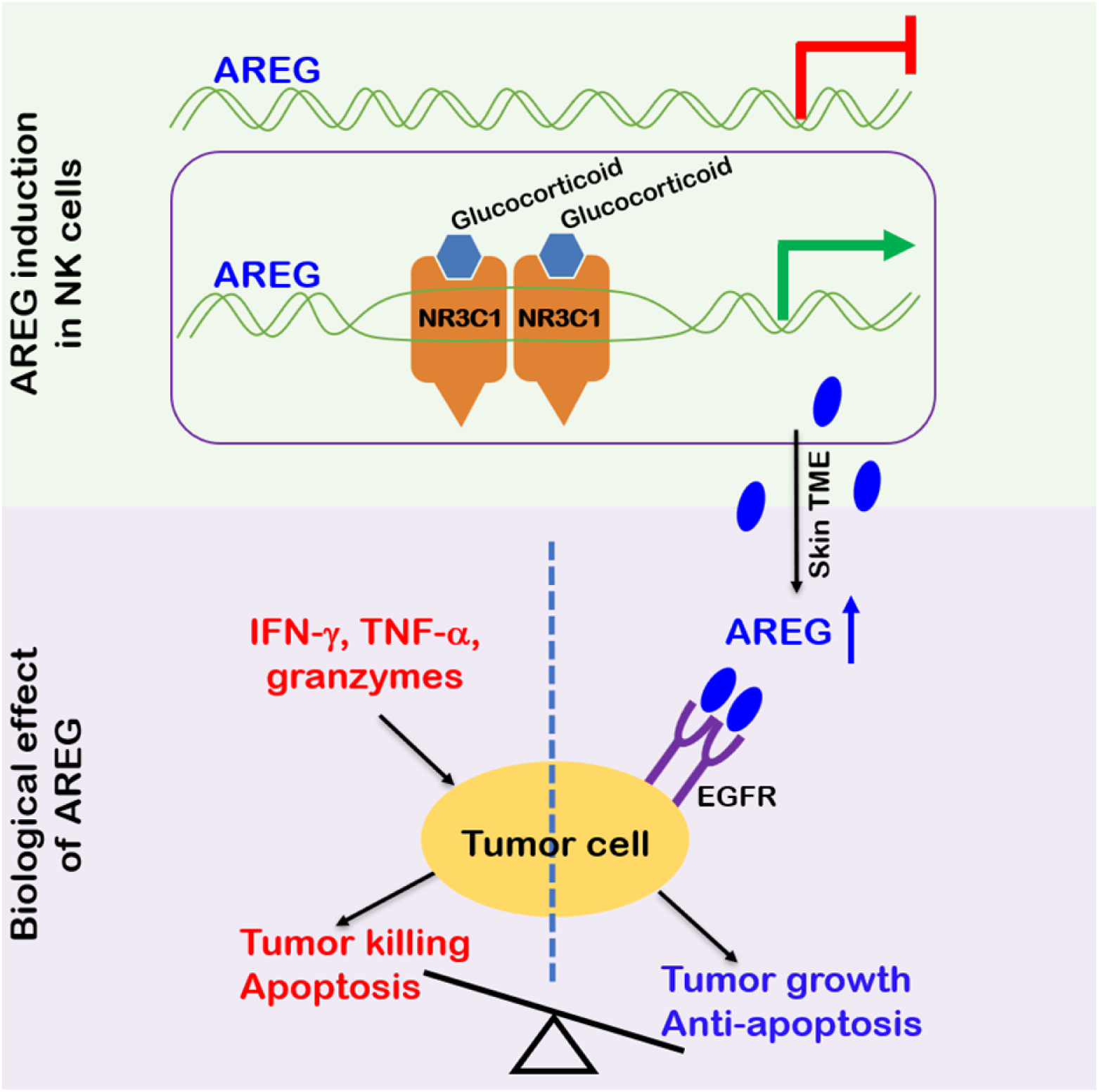

## Introduction

Skin cancers, the most prevalent human malignancies, primarily include melanoma (MM), basal cell carcinoma (BCC), cutaneous squamous cell carcinoma (cSCC), and extramammary Paget disease (EMPD)^1–3^. While surgical resection remains the standard for localized disease, immune checkpoint inhibitors targeting PD-1 and CTLA-4 have demonstrated significant efficacy in advanced or metastatic BCC, cSCC, and MM^2,4–6^. Despite subtype heterogeneity, these malignancies exhibit shared tumor microenvironment (TME) mechanisms driving immune suppression and evasion. Common hallmarks include immunosuppressive cellular networks (e.g., exhausted T cells, regulatory T cells, myeloid-derived suppressor cells)^7–13^, hypoxia-mediated metabolic reprogramming^14–16^, and cancer-associated fibroblast-driven stromal remodeling^17^, which collectively impair cytotoxic immune cell infiltration and function. Nonetheless, TME-driven treatment resistance remains a major barrier to improved clinical outcomes^5,18^, highlighting the need to further define determinants of unresponsive TMEs and their associated biomarkers.

Natural killer (NK) cells serve as sentinel effectors in tumor immune surveillance, eliminating malignant cells by releasing perforin and granzymes, engaging death receptors, and producing anti-tumor cytokines such as IFN-γ and TNF-α^19^. Their therapeutic potential is underscored by two inherent advantages: MHC-unrestricted target recognition circumvents HLA-related limitations, and minimal risk of graft-versus-host disease (GVHD) permits safe allogeneic application^19^. Clinically, NK cells can be sourced from both autologous (peripheral blood-derived) and allogeneic (umbilical cord blood, CD34⁺ progenitor-derived, iPSC-differentiated) origins, with GMP-compliant expansion protocols facilitating standardized production of clinical-grade NK cells^19,20^. This unique combination of multimodal anti-tumor activity and scalable manufacturing establishes NK cells as a versatile platform for cellular therapy, with over 190 active clinical trials investigating their efficacy against hematologic and solid malignancies (ClinicalTrials.gov).

Within tumors, NK cell function is shaped by the TME, exhibiting context-dependent duality. In melanoma, NK cells enhance anti-PD-1 therapy by promoting FLT3LG-dependent intratumoral stimulatory dendritic cells^18^, whereas their dysfunction correlates with cSCC progression^21^. Increasing evidence suggests that NK cell functional suppression arises through conserved mechanisms shared across diverse malignancies^22,23^. For instance, PGE2 disrupts LAMP3⁺DC–NK cell interactions across TMEs, weakening tumor restriction in melanoma and colorectal cancer^23,24^. Tumor-associated NK (TaNK) cells exhibit stress-associated transcriptional features and impaired cytotoxicity, correlating with poor prognosis or immunotherapy resistance in melanoma, breast, lung, and metastatic urothelial carcinoma^22,23^. Further delineating these convergent inhibitory mechanisms is essential for advancing NK cell-based immunotherapies.

Diverging from canonical NK cell effector functions, amphiregulin (AREG) acts as an epidermal growth factor receptor (EGFR) ligand, driving tumor progression by promoting cell survival, proliferation, and immune tolerance^25–28^. Its upregulation in tumors also contributes to therapeutic resistance, positioning AREG as a mediator of both tumor aggression and therapy evasion^27,29–31^. This dual role–directly fueling tumorigenesis while impairing treatment efficacy–establishes AREG as a multifaceted therapeutic target. Our chromatin accessibility analysis revealed a species-specific regulatory divergence: the AREG promoter is constitutively accessible in human NK cells but remains closed in murine counterparts^32^. However, two key questions remain–what upstream signals induce AREG expression in human NK cells, and how NK cell-derived AREG influences their tumor immunity.

Through comparative single-cell transcriptomic profiling of NK cells from matched tumor and peri-tumor tissues, we systematically mapped their functional reprogramming in cutaneous malignancies. This analysis identified glucocorticoids as inducers of AREG production in NK cells, and our experiments demonstrated that NK cell-derived AREG counteracts their tumor-restricting function. These findings highlight the GR-AREG axis as a potential target for restoring NK cell function in cancer immunotherapy.

## Results

### NK cells are stable constituents of skin tumor lymphocytes

To examine whether skin cancers disrupt normal NK cell physiology, we conducted a study using freshly isolated tumor and adjacent normal peri-tumor tissue from patients with BCC, cSCC, EMPD, and acral melanoma (aMM). A total of 12 paired samples (3 donors for each cancer type) were analyzed by single-cell RNA sequencing (scRNA-Seq), and 34 paired samples were analyzed by flow cytometry (Figures 1A, S1, Tables S1 and S2, and Methods).

**Figure 1.**
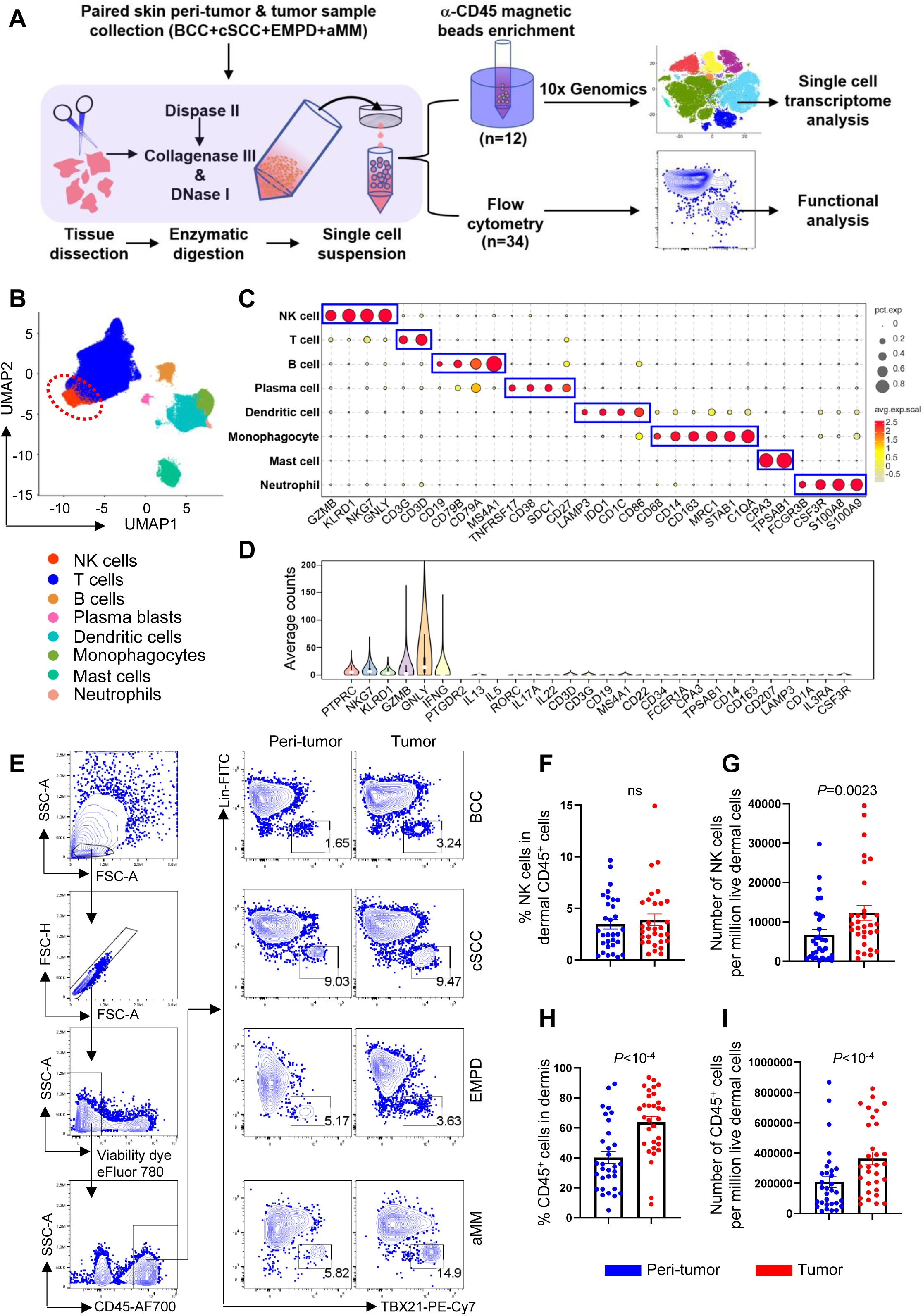
NK cells are stable constituents of skin tumor lymphocytes. **(A)** Experimental design of this study.(B) UMAP of skin tumor and peri-tumor CD45^+^ lymphocytes. (C) The expression of cell type specific markers for (B). (D) The expression of NK cell and other cell type specific markers in the NK cell cluster. (E) Flow cytometry of NK cells in tumor and peri-tumor of BCC, cSCC, EMPD and aMM. (F) Percentage of NK cells in dermal CD45^+^ cells from skin tumor and peri-tumor. (G) Number of NK cells per million live dermal cells from skin tumor and peri-tumor. (H) Percentage of dermal CD45^+^ cells from skin tumor and peri-tumor. (I) Number of CD45^+^ cells per million live dermal cells from skin tumor and peri-tumor. For (F-I) n=31, each dot represents a unique skin tumor donor (BCC, n=11; cSCC, n=14; EMPD, n=4; aMM, n=2), Wilcoxon matched-pairs signed rank test, ns, not significant, data are mean with s.e.m.

To minimize the presence of skin structural cells in the sequenced samples^6,33^, a CD45^+^ cell enrichment step was performed prior to library construction (Figure 1A). After filtering with Seurat and removing non-hematopoietic cells, skin CD45^+^ cells formed eight distinct clusters in uniform manifold approximation and projection (UMAP), representing NK cells, T cells, B cells, plasma blasts, dendritic cells, monocytes, mast cells, and neutrophils (Figures 1B and 1C). NK cells were identified based on their specific expression of NK cell markers (NKG7, GZMB, GNLY, KLRD1, IFNG) and the absence of markers from other cell types (Figures 1C, 1D, and S2A)^32,34,35^.

Flow cytometry was employed to validate the presence of NK cells in human skin. To exclude T cells, B cells, monocytes/macrophages, dendritic cells, and other lineage-positive cells from the CD45^+^ population, a panel of 14 lineage antibodies (CD3, CD4, TCRαβ, TCRγδ, CD19, CD20, CD22, CD34, FcεRIα, CD11c, CD303, CD123, CD1a, and CD14) was used^34^. NK cells were identified as Lin–TBX21^+^ (Figure 1E), encompassing both CD56^+^ and potentially CD56– subsets, as previously described^32,34,36^. Both CD56 and CD16 were detected in Lin–TBX21^+^ population, whereas CD127 was restricted in Lin^-^TBX21^-^ population, confirming that the gated NK cells do not include ILC1s (Figure S2B). Our results showed that NK cells constituted a stable population across all skin tumor and peri-tumor samples (Figure 1E). While the percentage of NK cells within the CD45^+^ population remained unchanged in the tumor, their absolute numbers showed an increase, which correlated with an elevation in CD45^+^ cells (Figures 1F–1I).

### Skin tumor associated NK cell transcriptional features

Reactome analysis of genes upregulated in skin tumors versus peri-tumor tissue identified enriched transcriptional features in tumor-associated NK cells. Notably, both anti-and pro-tumor-associated genes were represented in the enriched pathways (Figure 2A and Table S3). For instance, pathways related to IL-4 and IL-13 signaling (IL4R, FOS, CEBPD, JUNB), which reinvigorate terminally exhausted intratumoral CD8^+^ T cells^37^, and those involved in regulated necrosis and interferon signaling (IFNG, GZMB, CASP4, IRF8, ISG20, IFITM1, IFITM3, GBP5), crucial for tumor restriction, were enriched in tumoral NK cells. Conversely, pathways linked to tumor stress responses were also enriched, including the HSP90 chaperone cycle for steroid hormone receptors (HSPA1A, HSP90AA1, HSPA1B, DANJB1), which enhances glucocorticoid receptor activity and tumor resistance^38–43^, and EGFR signaling in cancer (AREG, UBC, HSP90AA1), which promotes tumor progression^29,44^ (Figure 2A and Table S3). These findings reflect the conserved yet complex immunosuppressive landscape shared across skin cancers^7–15,17^, where skin TMEs, shaped by chronic cytokine exposure and tumor-associated stress, promote a paradoxical coexistence of pro-and anti-tumor transcriptional programs in NK cells.

**Figure 2.**
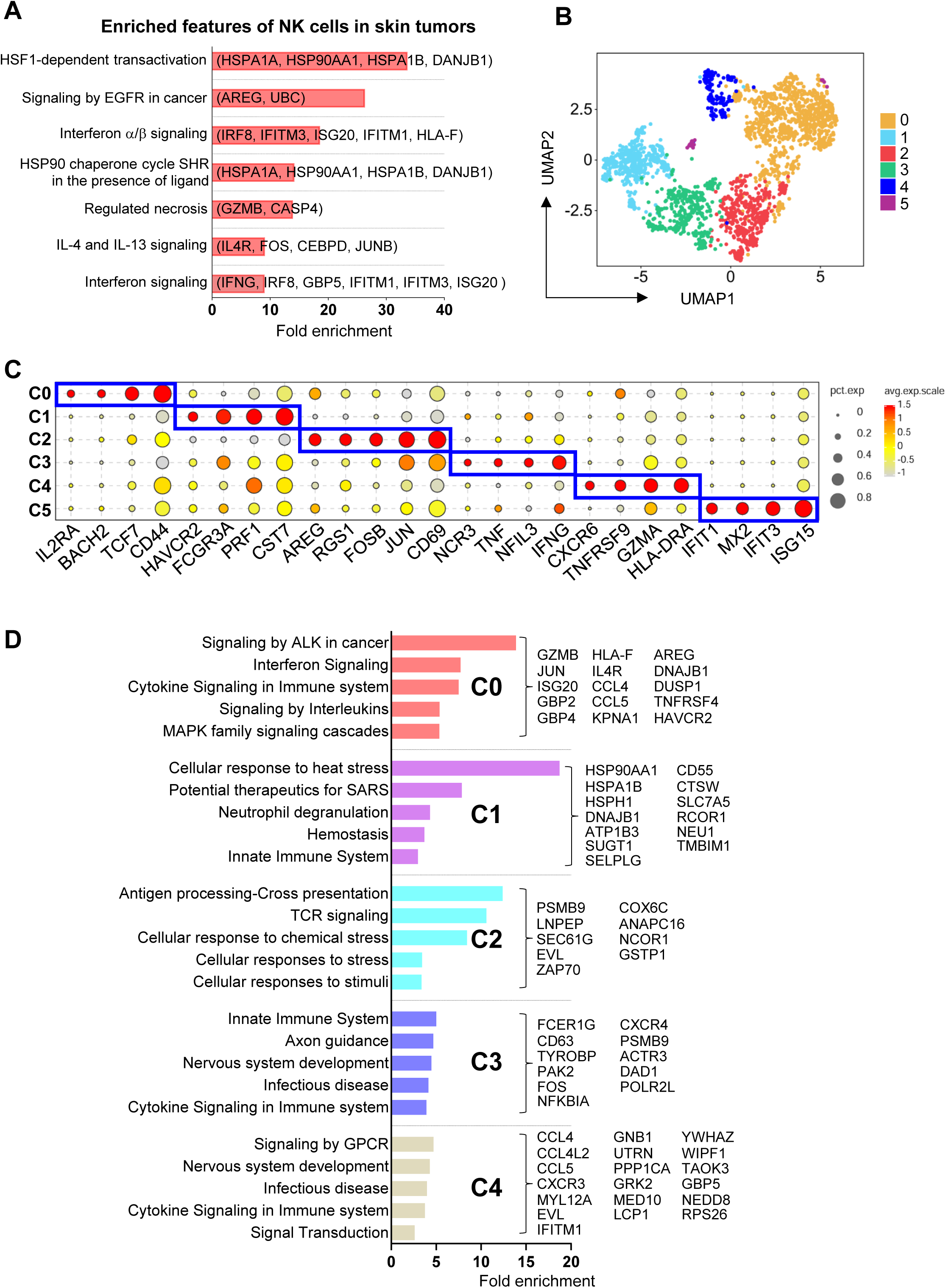
Common transcriptional features of skin tumor and peri-tumor NK cells. (A) Reactome analysis of highly expressed genes in skin tumor NK cells. (B) UMAP of re-clustered skin tumor and peri-tumor NK cells. (C) Dot plot displaying the signature genes in skin NK cell clusters. (D) Reactome analysis of highly expressed genes in each cluster of skin tumor NK cells.

UMAP analysis identified six NK cell clusters, primarily driven by their intrinsic transcriptional characteristics rather than cancer type or tumor microenvironment (tumor vs peri-tumor) (Figures 2B, 2C, S3A and Table S4). Cluster 0 (C0) NK cells exhibited high expression of IL2RA, TCF7, BACH2, and AREG, markers also present in blood CD56^hi^NK cells^32,34,45^. C1 NK cells highly expressed HAVCR2, FCGR3A, PRF1, and CST7, markers commonly found in cytotoxic CD56^dim^NK cells^32,34^. C2 NK cells also showed high AREG expression along with genes associated with tumor-infiltrating NK cells (RGS1, CD69)^23^ and AP-1 family members (JUN, ATF3, FOSB). C3 and C4 NK cells were enriched for genes linked to tissue residency, cytotoxicity, and immune inhibition (IFNG, TNF, CXCR6, GZMA, PRF1, TNFRSF9, TIGIT, LAG3)^23^. C5 NK cells constituted a minor population and exhibited high expression of interferon-stimulated genes (Figure 2C).

Next, the effects of skin tumors on each NK cell cluster were investigated, the enriched pathways by reactome for each cluster are shown in Figure 2D. Consistent with the common features observed in Figure 2A, interferon-or cytokine-signaling associated genes were commonly upregulated in tumor C0, C3, and C4 NK cells (e.g., IL4R, ISG20, GBP2, GBP4, TNFRSF4, JUN). Genes associated with MAPK signaling cascades, a major downstream pathway of EGFR signaling (AREG, JUN, DNAJB1, DUSP1), were upregulated in tumor C0 NK cells. Additionally, genes involved in the cellular response to stress were commonly upregulated in tumor C1 and C2 NK cells (HSP90AA1, HSPA1B, HSPH1, DNAJB1, PSMB9, CSTP1).

### Dysregulated cytokine production in skin tumor NK cells is marked by elevated AREG expression

Transcriptomic analysis revealed that skin tumor NK cells exhibited dual functional features: anti-tumor activity alongside elevated AREG expression, a cytokine that promotes keratinocyte/fibroblast proliferation and therapy resistance via EGFR signaling (Figures 2A and 2D)^27,29–31,46,47^. Flow cytometry confirmed this dichotomy, showing increased AREG production in tumor-infiltrating NK cells compared to their peri-tumor counterparts, while revealing concurrent downregulation of TNF-α and preserved IFN-γ production (Figures 3A-3D).

**Figure 3.**
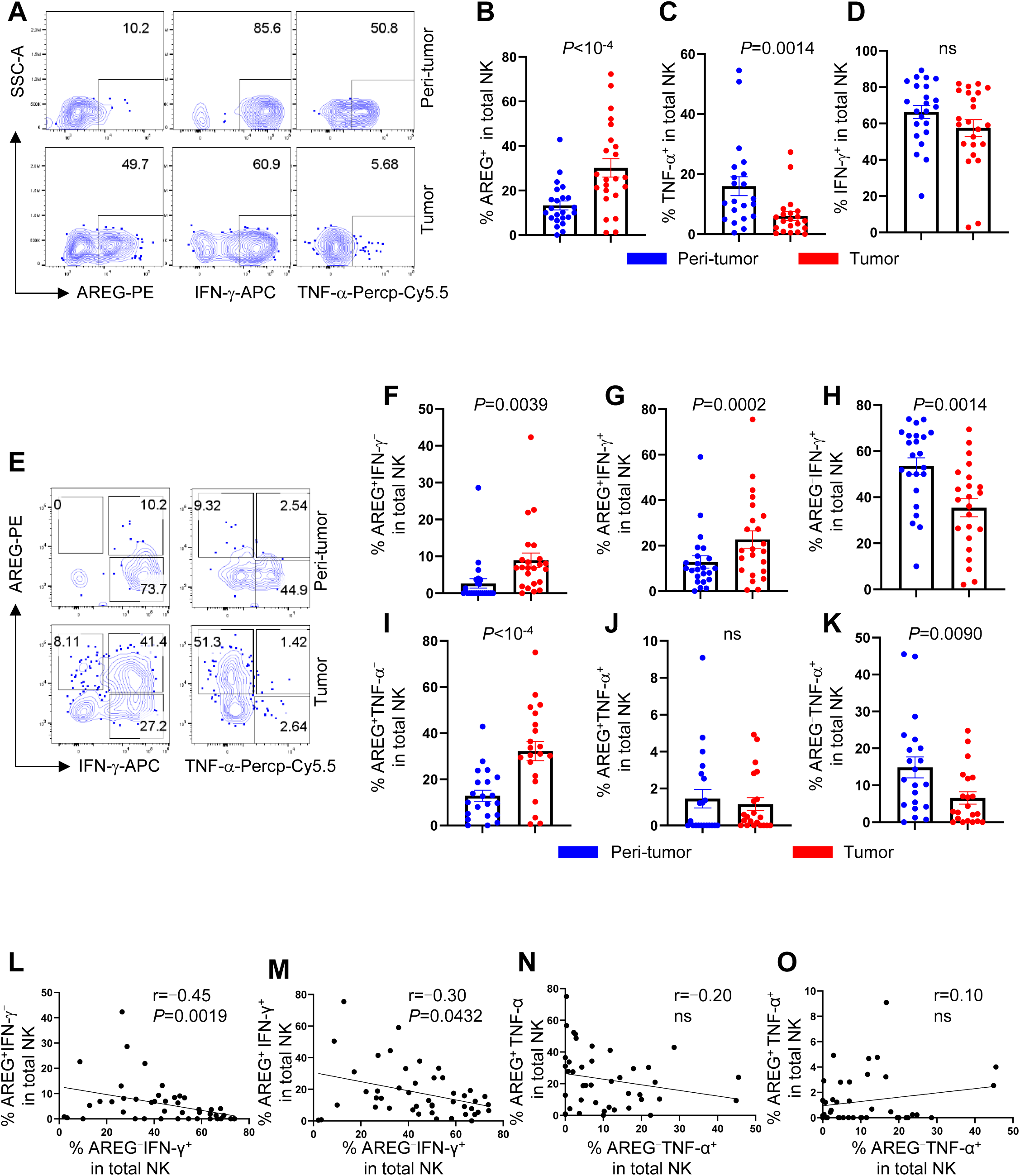
Skin tumor TME upregulates AREG production by NK cells. (A) Skin dermal cells were stimulated with PMA and ionomycin for 2 hrs, the production of AREG, IFN-γ and TNF-α by NK cells (CD45^+^Lin^-^TBX21^+^) were detected by flow cytometry. (B-D) Percentage of AREG^+^ (C, n=23), TNF-α^+^ (D, n=21) and IFN-γ^+^ (E, n=23) NK cells in the tumor and peri-tumor detected in (A). (E) Flow cytometry of NK cells produced AREG vs IFN-γ or TNF-α in the tumor and peri-tumor. (F-K) Percentage of AREG^+^IFN-γ^-^ (F), AREG^+^IFN-γ^+^ (G), AREG^-^IFN-γ^+^ (H) NK cells (n=23) and AREG^+^TNF-α^-^ (I), AREG^+^TNF-α^+^ (J) and AREG^-^TNF-α^+^ (K) NK cells (n=21) detected in (E). (L, M) Correlation of AREG^-^IFN-γ^+^ with AREG^+^IFN-γ^-^ (L) or with AREG^+^IFN-γ^+^ (M) dermal NK cells (n=46). (N, O) Correlation of AREG^-^TNF-α^+^ with AREG^+^TNF-α^-^ (N) or with AREG^+^TNF-α^+^ (O) dermal NK cells (n=42). For (B-D), (F-K), each dot represents a unique skin tumor donor, Wilcoxon matched-pairs signed rank test, data are mean with s.e.m., for (L-O), Spearman correlation, ns, not significant.

Subpopulation analysis revealed increased frequencies of both AREG⁺IFN-γ⁻ and AREG⁺IFN-γ⁺ NK cells within skin tumors (Figures 3E-3G). In contrast, the AREG⁻IFN-γ⁺ subset was markedly reduced (Figure 3H). This polarization pattern extended to TNF-α compartments, with elevated AREG⁺TNF-α⁻ populations and diminished AREG⁻TNF-α⁺ subsets in tumor microenvironments (Figures 3E and 3I-3K). Importantly, the frequency of AREG⁺NK subsets (AREG⁺IFN-γ⁻ and AREG⁺IFN-γ⁺) exhibited a negative correlation with IFN-γ single-positive cells (AREG⁻IFN-γ⁺) (Figures 3L and 3M), whereas no such association emerged between AREG⁺ and TNF-α⁺ compartments (Figures 3N and 3O). Collectively, these data demonstrate a functional reprogramming of NK cells in skin tumors, characterized by coordinated shifts toward AREG-dominant cytokine profiles.

### Glucocorticoids specifically induces NK cell AREG production

To identify potential drivers of AREG production in NK cells within skin tumors, we tested 14 stimulation conditions, including inflammatory cytokines (IL-2, IL-12, IL-15, IL-18, IL-21), activating receptor ligands (4-1BBL, MICA), activating antibodies (anti-CD2, anti-NKp46), and metabolic regulators (IGF-I, IGF-II, insulin, transferrin), both individually and in combination. None of these conditions induced detectable AREG production in NK cells, in contrast to the robust induction observed with PMA and ionomycin (Figure S3B). This lack of response to canonical NK-activating signals suggests that AREG production in tumor-associated NK cells is regulated independently of traditional NK activation pathways.

We next examined whether transcriptional differences between AREG⁺NK cells (C0, C2) and AREG⁻NK cells (C1, C3, C4, C5) could reveal regulatory mechanisms controlling AREG expression (Figures 4A and 4B). Notably, genes associated with glucocorticoid receptor (GR) signaling were consistently upregulated in AREG⁺NK cells (Figure 4C)^48–51^. Moreover, the elevated GR target gene set scores in AREG⁺NK cells persisted across anatomically distinct compartments, with AREG⁺NK cells exhibiting higher scores than AREG⁻NK cells in both tumor and peri-tumor regions (Figures 4D and Table S5). Consistent with their enhanced AREG production (Figure 3B), tumor NK cells displayed greater GR activity than their peri-tumor counterparts (Figure 4E).

**Figure 4.**
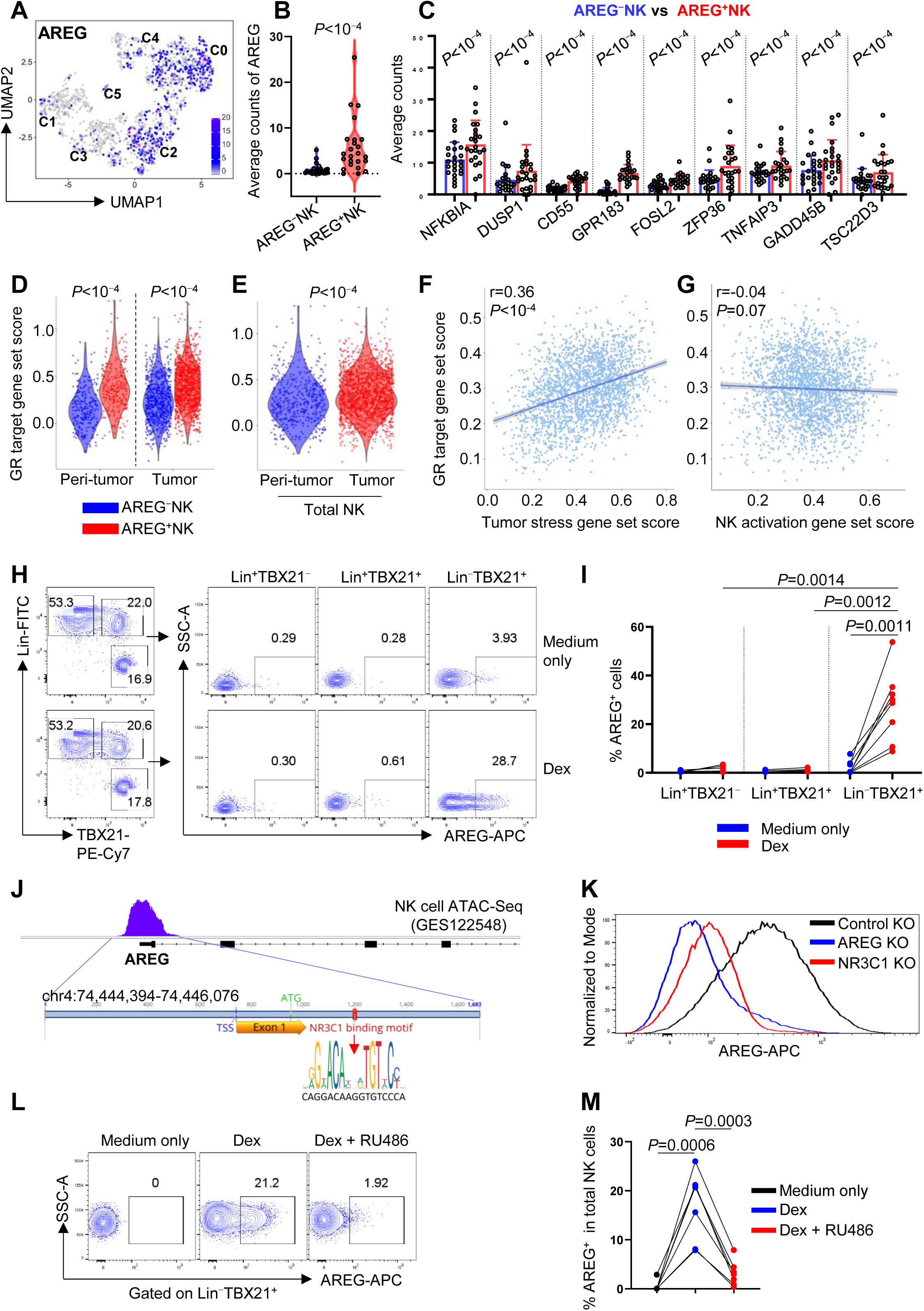
Glucocorticoid receptor activation induces NK cells AREG production. (A) AREG expression across NK cell clusters. (B, C) The average counts of AREG (B) and glucocorticoid target genes (C) between C1, 3, 4, 5 (AREG^-^) and C0, 2 (AREG^+^) NK cells. *P* value was determined by Seurat FindMarkers. (D, E) GR target gene set scores comparing AREG⁻ and AREG⁺ NK cells (D) and total NK cells (E) between tumor and peri-tumor regions. Gene set scores were calculated using AddModuleScore in Seurat, and P values were determined by the Wilcoxon rank-sum test in R. (F, G) Pearson correlation between GR target gene set scores and tumor stress (F) or NK cell activation (G) gene set scores. (H) PMBCs were stimulated with Dex for 16 hrs, the production of AREG in indicated cell populations were detected by flow cytometry. (I) Percentage of AREG^+^ cells in (H), n=8, two-tailed paired t-test. (J) Schematic map of AREG open chromatin region in NK cells (GES122548) and NR3C1 binding motif. (K) Control, AREG or NR3C1 knockout NK cells were cultured in RPMI 1640 for 48 hrs after electroporation, then were stimulated with Dex for 16 hrs, AREG production was detected by flow cytometry. (L) PBMCs were stimulated with Dex or Dex+GR inhibitor (RU486) for 16 hrs, the production of AREG by NK cells was detected by flow cytometry. (M) Percentage of AREG^+^NK cells in (K), n=7, two-tailed paired t-test.

Our analysis showed that GR target gene set scores correlate specifically with tumor stress signatures rather than anti-tumor effector modules in NK cells (Figures 4F and 4G). This aligns with previous studies demonstrating that tumor stress-induced heat shock proteins enhance GR activity by directly binding as coactivators^38–43^, as well as our finding of HSP family gene upregulation in skin tumor NK cells (Figure 2A, Table S3). Collectively, these findings suggest that elevated GR activity may contribute to increased AREG production in tumor associated-NK cells.

To validate this hypothesis, peripheral blood mononuclear cells (PBMCs) were treated with dexamethasone (Dex), a synthetic glucocorticoid widely used in clinical settings. NK cells, but not Lin⁺ cells (including T and B cells), exhibited a marked increase in AREG production upon Dex treatment (Figures 4H, 4I, and S4A). This response was recapitulated with other glucocorticoids (betamethasone and methylprednisolone; Figures S4B and S4C). Despite testing glucocorticoid concentrations spanning nanomolar to micromolar ranges, no dose-dependent increase in the proportion of AREG⁺NK cells was observed (Figures S4A–S4C). This plateau in AREG induction at higher glucocorticoid concentrations was not due to cytotoxicity, as NK cell viability remained unaffected across all tested doses (Figures S4D-S4F). However, exposure of NK cells to other structurally diverse hormones or hormone analogs–including dydrogesterone, estrone, progesterone, salmeterol, testosterone, and triiodothyronine–failed to induce AREG production (Figure S4G). JASPAR motif analysis identified an GR (encoded by NR3C1) binding site within the chromatin-accessible region of the AREG promoter in human NK cells (Figure 4J). Consistent with this, both CRISPR-Cas9-mediated NR3C1 knockout and pharmacological GR antagonism with mifepristone (RU486) abolished Dex-induced AREG production in NK cells (Figures 4K-4M and S4H). These data establish that glucocorticoid-induced GR activation specifically drives AREG expression in NK cells.

### GR activation reprograms the anti-tumor transcriptional profile of NK cells

To comprehensively investigate GR activation in NK cells, we performed bulk RNA-seq on sorted NK cells from untreated or Dex-treated PBMCs under steady-state (Figure S5A). Among 102 Dex-upregulated genes, AREG showed the highest fold change, with enriched pathways including steroid hormone response (FKBP5, KLF9, CFLAR, TSC22D3)^49–51^, and negative regulation of NK/T cell activation, proliferation and apoptosis (CFLAR, PRDM1, TSC22D3, TLE1, TNFAIP8, PIK3IP1, TXNIP, HPGD, PTGER2, ZFP36L2, CD55, FOXO1) (Figures 5A, S5B; Table S6)^50,52–60^. Notably, many of these GR targets are linked to therapy resistance: PRDM1 reduces IL-2 sensitivity and suppresses IFN-γ and TNF-α production in NK cells^61–63^; TSC22D3 impairs anti-tumor responses by inhibiting DC type I interferon signaling and IFN-γ^+^T cell activation^50^; TLE1 attenuates NK effector function and memory responses^34,54,55,64^; and PTGER2 (the PGE2 receptor) suppresses XCL1 and CCL5 secretion by NK cells, hindering cDC1 recruitment and T cell-mediated tumor control (Figure 5A and Table S6)^24,65,66^. In contrast, Dex downregulated genes associated with type I/II interferon responses, positive regulation of TNF production, and programmed cell death, including IFIT1, IFIT2, IFIT3, MX1, DDX58, IRF7, XCL2, STAT1, MYD88, RELB, TRAF1, TNFSF10, FASLG, IL-32, CD226, IL2RB, and IL15RA (Figures 5A, S5B; Table S6)^67,68^.

**Figure 5.**
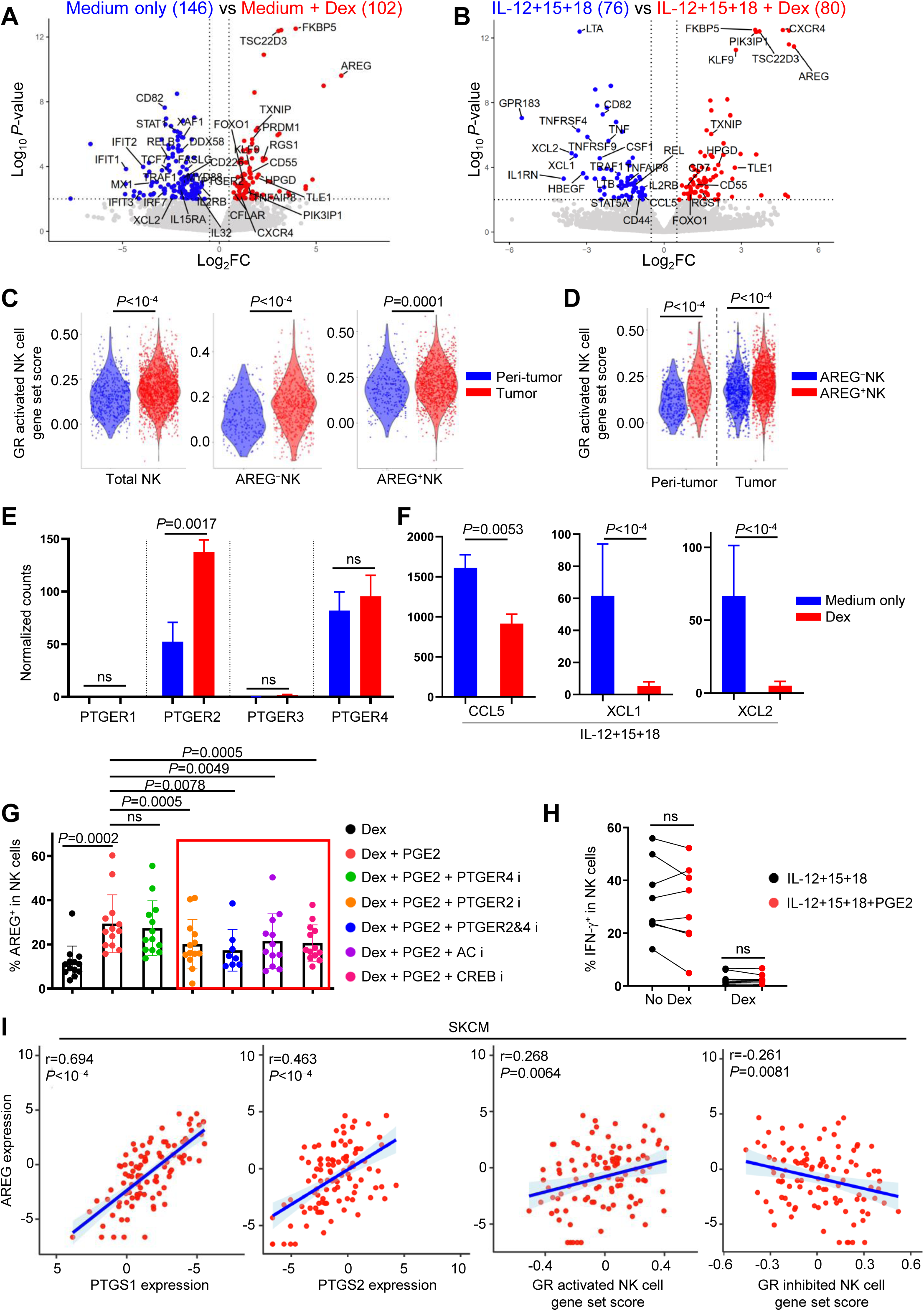
PGE2 signaling potentiates Dex induced AREG production in NK cells. (A) PBMCs were treated with or without Dex in RPMI 1640 for 16 hrs, CD56^dim^NK cells were stored and were subjected to bulk RNA-Seq, the DEgenes were shown in volcano plot (Log_2_FC> 0.5, *P*<0.01, determined by DESeq2, n=3). (B) As performed in (A), DEgenes of Dex treated or untreated NK cells in the condition of IL-12+IL-15+IL-18 stimulation were shown (n=3). (C, D) Using Dex-upregulated genes from (B) as a reference, GR-activated gene set scores were compared between skin tumor and peri-tumor regions for total NK cells, AREG⁻NK cells, and AREG⁺NK cells (C), and between AREG⁻ and AREG⁺NK cells within each region (D). (E, F) Normalized counts of indicated genes from (A) and (B). (G) PBMCs were untreated or incubated with PF-04418948 (PTGER2 inhibitor), L-161982 (PTGER4 inhibitor), SQ22536 (adenylate cyclase inhibitor) or 666-15 (CREB inhibitor) as indicated for 6 hrs, then were treated with Dex alone or together with PGE2 for 16 hrs, AREG production by NK cells were detected by flow cytometry. (H) PMBCs incubated with or without Dex were treated with IL-12+IL-15+IL-18 in the presence or absence of PGE2 for 16 hrs, IFN-γ production by NK cells were detected by flow cytometry. (I) Pearson correlation of AREG with PTGS1 or PTGS1 expression in SKCM from TCGA, and Pearson correlation of AREG expression in SKCM from TCGA with GR activated or inhibited NK cell signature derived from our RNA-Seq data. For (G), Wilcoxon matched-pairs signed rank test, for (H), two-tailed paired t-test. Data are mean with s.e.m., each dot represents one donor, ns, not significant.

Many of these genes regulate NK cell anti-tumor activity: for example, CD82 suppresses cancer cell invasiveness^69^; interferon-induced XAF1 promotes tumor cell apoptosis^70,71^; and the activating receptor CD226 enhances anti-tumor function by counteracting Dex-induced FOXO1-mediated suppression (Figure 5A and Table S6)^72^.

NK cells were additionally sorted from PBMCs treated with IL-12+IL-15+IL-18 alone or in combination with Dex and subjected to bulk RNA-seq. Consistent with steady-state NK cells, AREG exhibited the highest fold change among the 80 Dex-upregulated genes in cytokine-activated NK cells (Figure 5B, Table S6), with significant overlap in Dex-upregulated genes between activated and steady-state conditions (Figures 5A, 5B, S5C and Table S6). These findings demonstrate that GR activation suppresses transcriptional programs linked to NK cell anti-tumor activity, with AREG emerging as the most upregulated gene in both states. Using this approach, we performed GR target gene set score analysis in skin tumor NK cells with an NK cell-specific reference gene list derived from Dex-upregulated genes (Table S6), thereby circumventing the non-specific effects associated with published GR targets from other cell types used in Figures 4D and 4E^48–51^. Skin tumor NK cells exhibited higher GR activation scores than peri-tumor NK cells (Figure 5C), and AREG⁺ NK cells showed elevated scores compared to AREG⁻ NK cells (Figure 5D), further supporting the interplay between elevated GR activity in skin tumor NK cells and increased AREG production.

### PGE2 signaling enhances Dex-induced AREG production by NK cells

Tumor cell-derived PGE2 drives immune evasion by disrupting communication and inducing dysfunction in conventional dendritic cells (cDCs), NK cells, and cytotoxic T cells, while its metabolite 15-keto-PGE2 enhances the immunosuppressive activity of regulatory T cells^24,65,66,73^. Bulk RNA-seq revealed that human NK cells predominantly express the PGE2 receptors PTGER2 (EP2) and PTGER4 (EP4), with Dex selectively upregulating PTGER2 expression (Figure 5E). Interestingly, Dex mimicked PGE2-mediated suppression of NK cell-derived CCL5, XCL1, and XCL2—chemokines essential for cDC1 recruitment and anti-tumor immunity (Figure 5F)^24^. We therefore investigated whether PGE2 regulates AREG production in NK cells.

Our experiments demonstrated that although PGE2 alone failed to stimulate AREG production in NK cells (Figure S5D), it enhanced AREG production induced by Dex (Figure 5G). This synergistic effect was mediated specifically through PTGER2, as pharmacological inhibition of PTGER2, but not PTGER4, attenuated PGE2’s enhancement of AREG (Figure 5G). Combined inhibition of both PTGER2 and PTGER4 did not yield further suppression, confirming the dominant role of PTGER2 (Figure 5G). Furthermore, blocking downstream components of the PTGER2 pathway, adenylate cyclase (AC) and CREB, abolished PGE2-enhanced AREG production in NK cells treated with Dex (Figure 5G), implicating the PGE2–PTGER2–cAMP–CREB axis in amplifying GR-driven AREG expression. In contrast to its selective modulation of AREG, PGE2 had no effect on IFN-γ levels in NK cells, with or without Dex, despite Dex alone potently suppressing IL-12+IL-15+IL-18-induced IFN-γ production (Figure 5H).

RNA-seq data from skin cutaneous melanoma (SKCM) in The Cancer Genome Atlas (TCGA) was analyzed to assess the clinical relevance of AREG and its association with PGE2 synthesis. In SKCM, AREG expression positively correlated with levels of PTGS1 and PTGS2–genes encoding prostaglandin-endoperoxide synthase 1 and 2 (COX-1 and COX-2), which catalyze PGE2 biosynthesis (Figure 5G). Furthermore, AREG expression in SKCM exhibited a positive correlation with GR activation signatures and a negative correlation with GR inhibition signatures derived from our NK cell RNA-seq data (Figure 5G; Table S6). Thus, analysis of this independent dataset from TCGA supports a potential role for the PGE2 and GR pathway in regulating AREG expression in NK cells.

### GR activation primes AREG promoter accessibility in NK cells for amplified production upon restimulation

We next checked whether the GR activation in NK cells creates a sustained effect, PBMCs were either initially activated with Dex or remained unstimulated. Both sets of NK cells were then allowed to rest for 5 days, with a low dose of IL-15 to sustain NK cell viability. RNA-Seq and ATAC-Seq were performed on sorted NK cells after 16 hours of initial stimulation or after 5 days of rest to capture transcriptome and chromatin accessibility alterations associated with GR activation. Following secondary stimulation, AREG production was compared between the two groups (Figure 6A).

**Figure 6.**
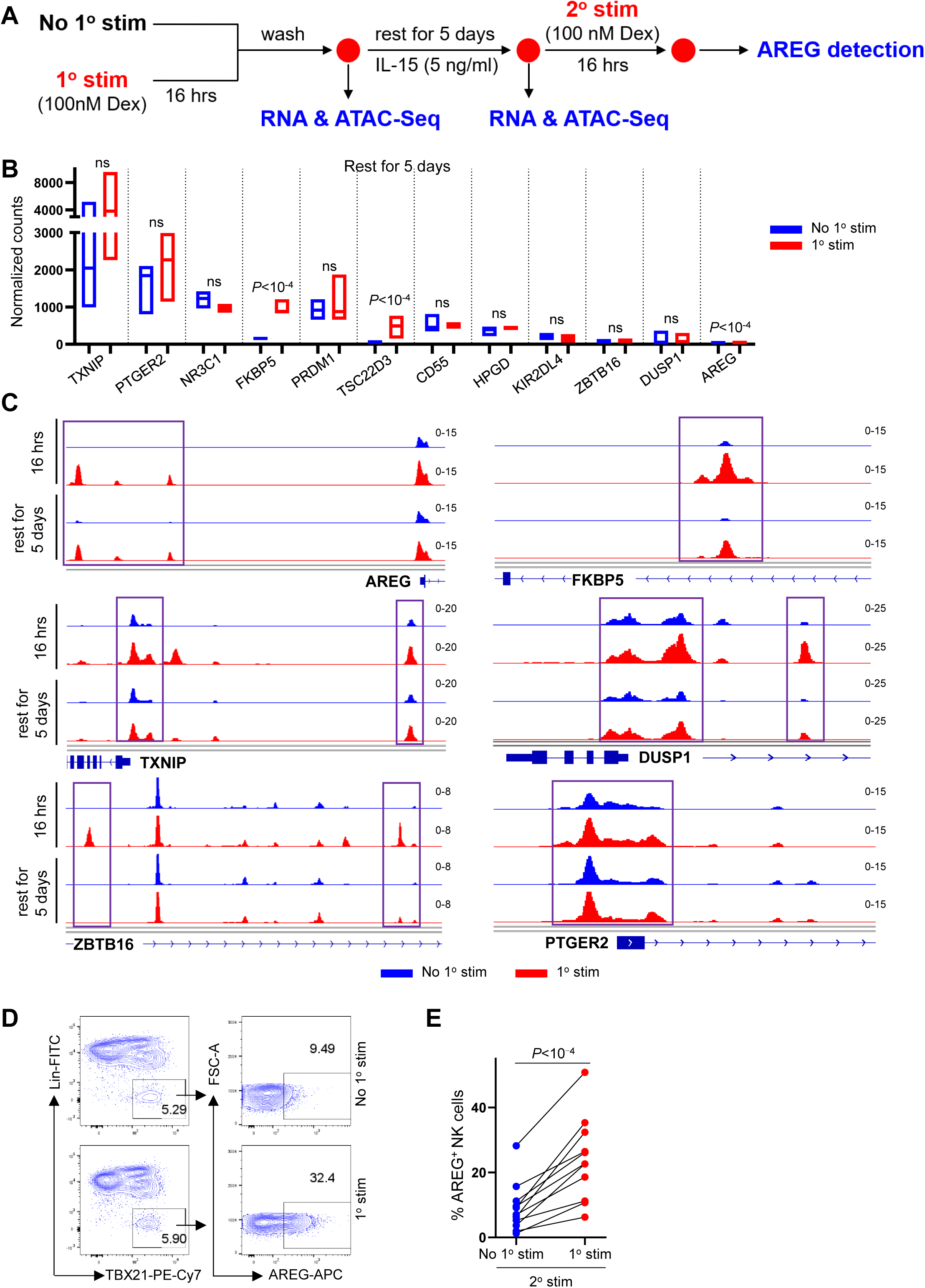
GR activation sustains chromatin accessibility to enhance AREG production upon restimulation. (A) Experimental schematic for detecting the effect of primary Dex treatment on the transcriptome, chromatin accessibility and AREG production of NK cells. (B) Expression of indicated genes in NK cells with or without primary Dex stimulation following a 5-day rest. *P* value was determined by DESeq2, n=4, ns, not significant. (C) ATAC-Seq analysis of sorted blood NK cells with or without primary Dex stimulation for 16 hrs, and after rest for 5 days. (D) PBMCs with and without primary Dex stimulation were stimulated with Dex after resting for 5 days, AREG production of NK cells were detected by flow cytometry. (E) The percentage of AREG^+^NK cells detected in (D) (n=11), two-tailed paired t-test, each dot represents one donor, ns, not significant.

Our results revealed that most genes induced by Dex after 16 hours were no longer differentially expressed or had returned to baseline following a 5-day rest (Figure 6B). Although AREG mRNA remained elevated after the 5-day rest, this residual expression likely reflected baseline differences rather than sustained induction (Figure 6B), as AREG protein remained undetectable in Dex-primed NK cells after rest, mirroring levels in unprimed cells (Figure S5E).

In contrast, Dex treatment rapidly increased chromatin accessibility at AREG promoter loci within 16 hours, and this open state persisted for at least 5 days after Dex removal (Figure 6C). This sustained accessibility pattern was shared by FKBP5 and TXNIP, whereas DUSP1 and ZBTB16 loci reverted to baseline (Figure 6C). Notably, chromatin accessibility at NR3C1 and PTGER2, critical regulators of AREG expression, remained unchanged by GR activation (Figure 6C and S5F). Together, these findings demonstrate that GR activation transiently boosts AREG production in NK cells while establishing persistent chromatin accessibility at AREG loci, even when protein levels are undetectable (Figure S5E). This primed chromatin state ultimately enabled Dex-primed NK cells to exhibit enhanced AREG production compared to unprimed counterparts upon restimulation (Figures 6D and 6E).

### NK cell-derived AREG mitigates NK cell-induced apoptosis in target cells

To assess the role of NK cell-secreted AREG in modulating cytotoxicity against target cells, we performed killing assays comparing wild-type (WT) and AREG knockout (KO) NK cells. The magnetic bead-enriched NK cells were expanded in NK MACS medium for 10 days to become competent for Cas9/RNP-mediated knockout. During this culture period, the medium intrinsically induced NK cell AREG production^32^. AREG knockout consistently achieved around 70% reduction in AREG production without altering IL-12+IL-15+IL-18 induced IFN-γ production or cell viability (Figures 7A-7D). The killing assay showed that WT and AREG KO NK cells induced comparable specific lysis of melanoma (A375, A2058) and liver cancer (Huh7) cell lines (Figure 7E). AREG KO NK showed a mild increase in lysis of the liver cancer cell line HepG2 and the colorectal cancer cell line HCT116 (Figure 7E). Degranulation (surface CD107a), as well as levels of GZMA, GZMB, NKG2A, NKG2D, and CD158, were similar between WT and AREG KO NK cells (Figure 7F), indicating that AREG production by NK cells does not affect their direct killing capacity.

**Figure 7.**
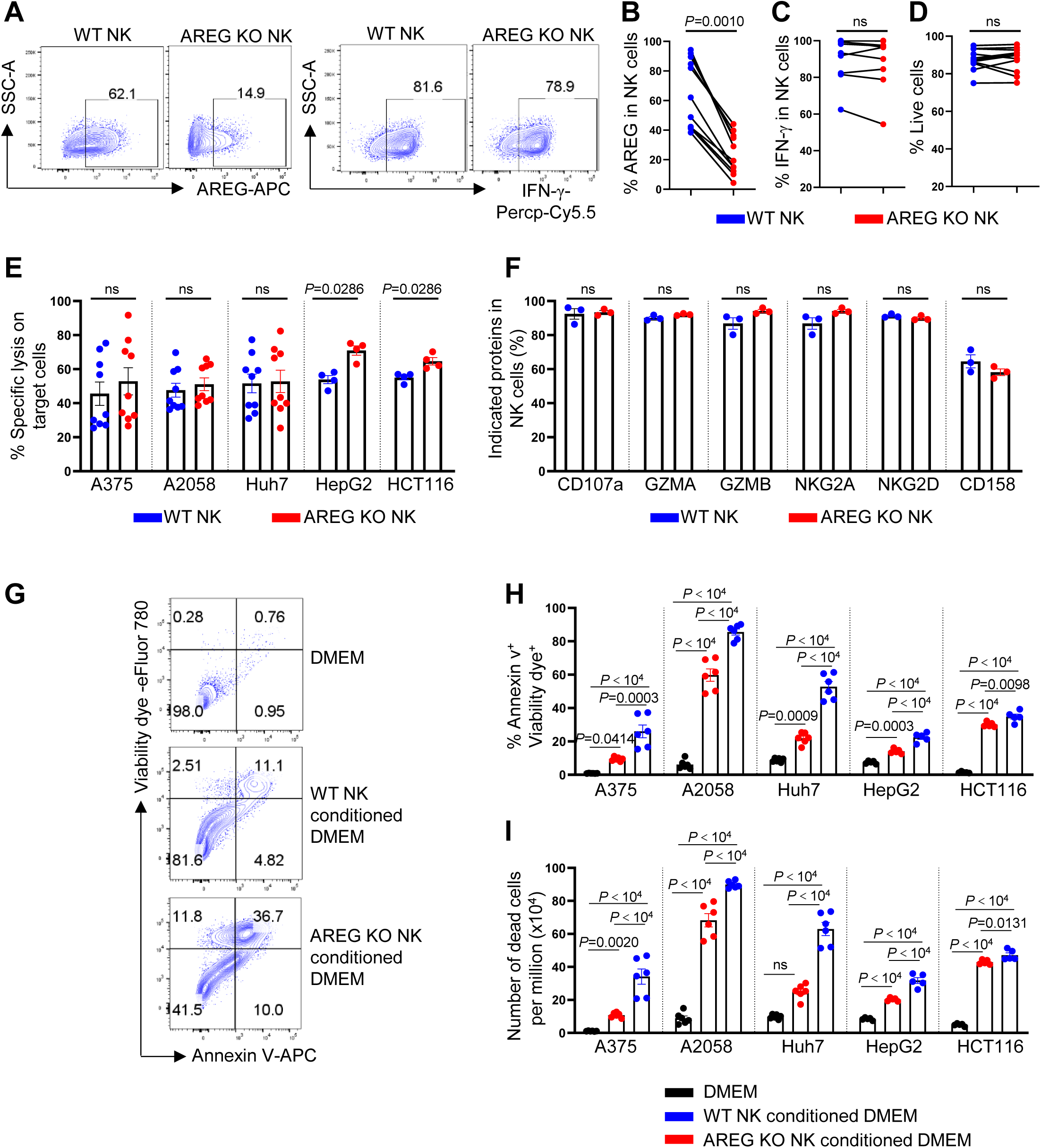
NK cell-produced AREG attenuates NK cell-induced apoptosis in target cells. (A) WT and AREG KO NK cells were cultured in NK MACS medium for 48 hrs after electroporation and stimulated with IL-12, IL-15, and IL-18 for 16 hrs. AREG and IFN-γ production were assessed by flow cytometry. (B–D) Percent of AREG⁺ (B), IFN-γ⁺ (C), and live NK cells (D) from (A). (E) Percent of specific lysis of A375 (n=9), A2058 (n=9), Huh7 (n=9), HepG2 (n=4), and HCT116 (n=4) by WT and AREG KO NK cells. (F) Percent of indicated proteins in WT and AREG KO NK cells detected by flow cytometry (n=3). (G) WT and AREG KO NK cells were stimulated with IL-12, IL-15, and IL-18 for 16 hrs in NK MACS medium, followed by washing and replacement of the culture medium with DMEM for an additional 48 hrs to generate NK cell-conditioned DMEM. A375 cells were then incubated with control DMEM, WT NK cell-conditioned DMEM, or AREG KO NK cell-conditioned DMEM for 48 hrs. Cells were subsequently subjected to Annexin V and viability dye staining, and analyzed by flow cytometry. (H, I) Percent of Annexin V⁺ viability dye⁺ cells (H) and number of dead cells (I) in A375 (n=6), A2058 (n=6), Huh7 (n=6), HepG2 (n=5), and HCT116 (n=5) after treatment as (G). For (B, C, F), Wilcoxon matched-pairs signed rank test; for (D), paired t-test; for (E), Mann-Whitney test; for (H, I), one-way ANOVA. Data are mean ± s.e.m. ns, not significant.

We then tested the effect of conditioned medium from WT or AREG KO NK cells on target cell survival. Target cells incubated with AREG KO NK-conditioned medium exhibited increased apoptosis and a higher number of dead cells compared to those treated with WT NK-conditioned medium across all tested cell lines (Figures 7G-7I). These results suggest that NK cell-derived AREG counteracts apoptosis in target cells triggered by NK-derived soluble factors (e.g., IFN-γ), thereby reducing overall cytotoxicity.

### NK cell-produced AREG compromises anti-tumor efficacy

Our previous study revealed a critical species-specific difference: Human NK cells produce AREG, whereas mouse NK cells lack this ability due to epigenetically inaccessible Areg chromatin loci^32^. This divergence limits the utility of genetic mouse models for studying NK cell-derived AREG. To address this, we investigated AREG’s biological role in human NK cells by engrafting NCG mice (NOD/ShiLtJGpt-*Prkdc^em26Cd52^*Il2rg*^em26Cd^*^22^*/Gpt*) with human melanoma or cSCC cell lines and evaluating the tumor-restricting effects of adoptively transferred WT or AREG KO human NK cells, expanded in NK MACS medium and stimulated with IL-12+IL-15+IL-18 (Figure 8A).

**Figure 8.**
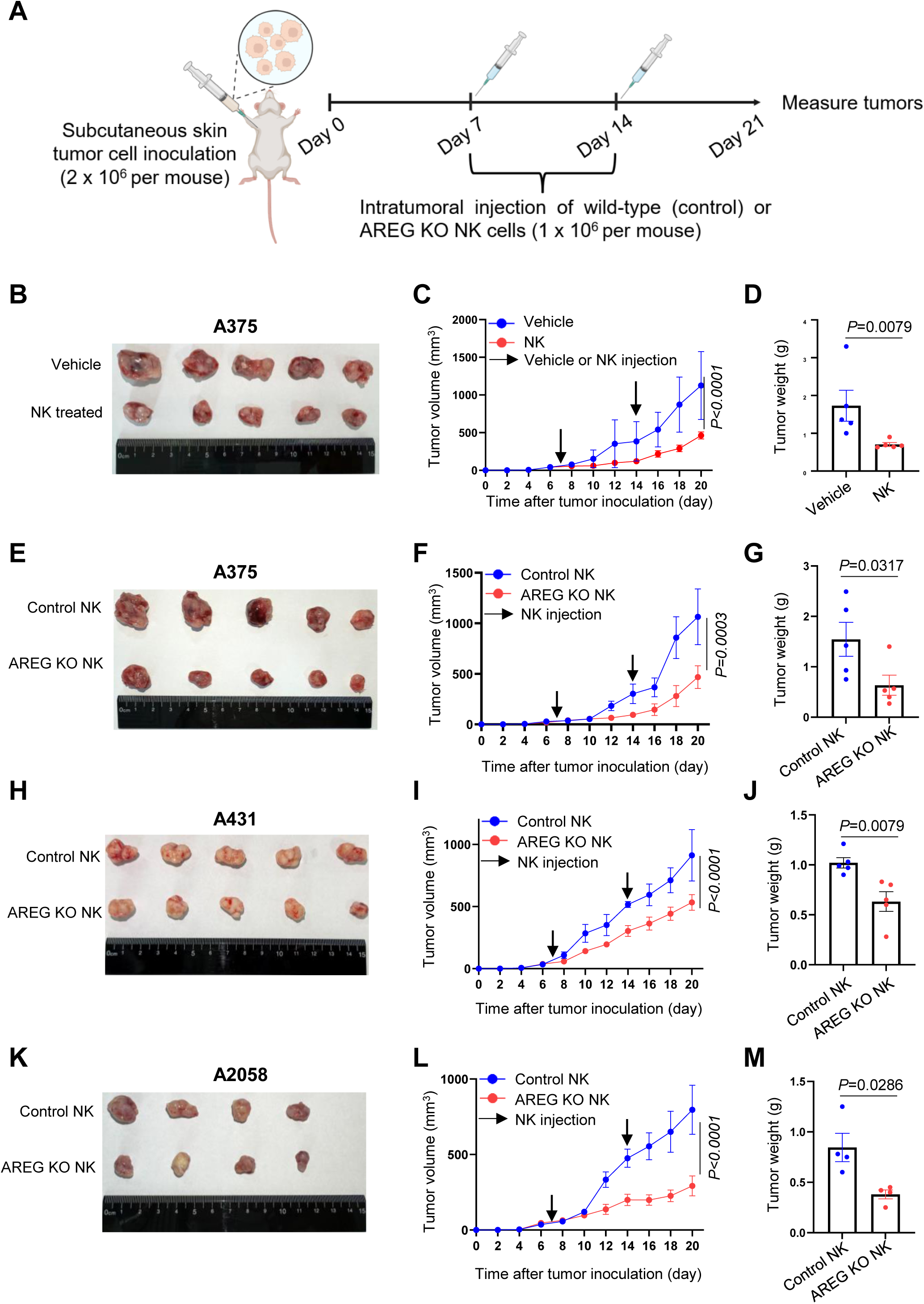
AREG knockout in NK cells enhances their skin tumor restriction. Skin cancer cell lines were inoculated subcutaneously in the axillary region of NCG mice. Wild type or AREG knockout NK cells were stimulated with IL-12, IL-15, and IL-18 for 16 hrs before intratumoral injection on day 7 and 14 post-tumor inoculation. (B) A375 inoculated NCG mice were injected intratumorally with 0.9% NaCl (vehicle) or wild-type NK cells as described in (A), tumors from control and NK cell-treated mice were dissected, and tumor volume (C) and weight (D) were measured (n=5). (E-G), The experiment was performed as in (B), except NCG mice were treated with wild type NK cells (control) and AREG knockout NK cells (n=5). (H-M), The experiment was performed as in (E-G), except NCG mice were inoculated with A431 cells (n=5) (H-J) and A2085 cells (n=4) (K-M). For (C, F, I, L), Two-way ANOVA, for (D, G, J, M), Mann-Whitney test.

Intratumoral injection of WT NK cells significantly suppressed tumor progression in NCG mice engrafted with the melanoma cell line A375, as evidenced by reduced tumor volume and weight compared to untreated controls (Figures 8B-8D). Moreover, AREG KO NK cells exhibited enhanced tumor-restricting activity, further inhibiting A375 tumor growth compared to WT NK cells (Figures 8E-8G). To assess the broader relevance of this finding, we tested AREG KO NK cells against cSCC (A431) and an additional melanoma line (A2058). Consistent with the A375 results, AREG KO NK cells markedly improved tumor control in both models, underscoring AREG function as a conserved negative regulator of NK cell anti-tumor activity (Figures 8H–8M). Conversely, Dex treated NK cells abrogated their ability to restrict A375-derived tumor growth (Figures 9A-9C).

**Figure 9.**
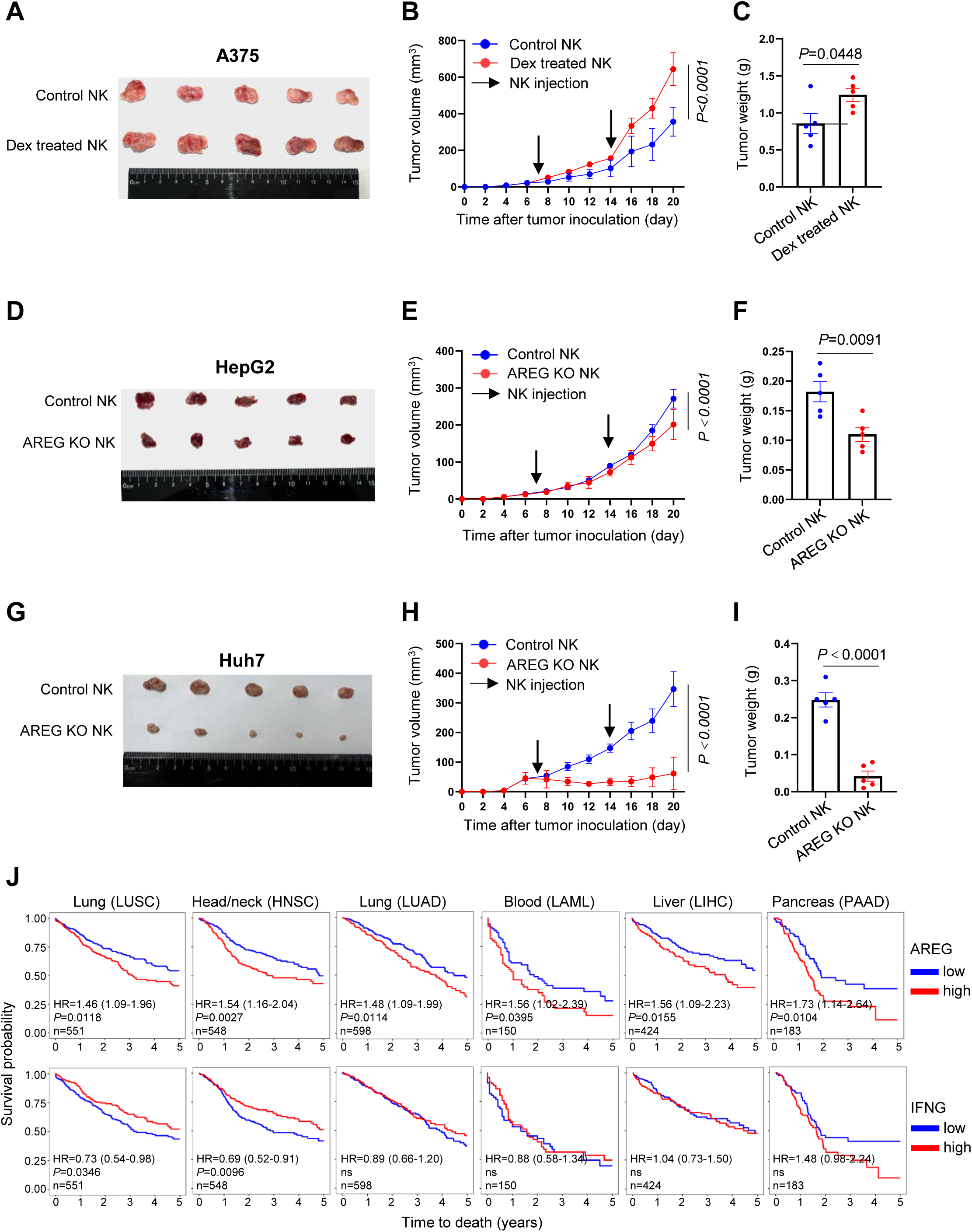
AREG knockout in NK cells enhances their liver tumor restriction. (A-C) A375 inoculated NCG mice were injected intratumorally with Dex untreated (control) or treated wild-type NK cells, tumor volume and weight were measured (n=5). (D-F) HepG2 inoculated NCG mice were injected intratumorally with wild type (control) or AREG knockout NK cells, tumor volume and weight were measured as in (A-C) (n=5). (G-I) The experiment was performed as in (D-F), except NCG mice were inoculated with Huh7 (n=5). (J) The association of AREG or IFNG expression with the survival probability in different cancer types based on the TCGA database. HR (hazard ratio), Lung Squamous Cell Carcinoma (LUSC), Lung Adenocarcinoma (LUAD), Head-Neck Squamous Cell Carcinoma (HNSC), Acute myeloid leukemia (LAML), Liver Hepatocellular Carcinoma (LIHC), Pancreatic adenocarcinoma (PAAD). For (B, E, H), Two-way ANOVA, for (C, F, I), Mann-Whitney test.

To assess whether NK cell-derived AREG impairs anti-tumor activity beyond skin cancers, we tested AREG KO NK cells in NCG mice engrafted with HepG2 and Huh7 liver cancer cell lines. AREG KO NK cells showed enhanced tumor suppression against HepG2-derived liver tumors (Figures 9D–9F) and completely blocked the growth of Huh7-derived tumors compared to WT NK cells (Figures 9G–9I). TCGA data analysis revealed that high AREG expression correlates with poor prognosis in lung squamous cell carcinoma (LUSC), head and neck squamous carcinoma (HNSC), lung adenocarcinoma (LUAD), acute myeloid leukemia (LAML), liver hepatocellular carcinoma (LIHC), and pancreatic adenocarcinoma (PAAD) (Figure 9J). In contrast, high IFNG expression showed no consistent association with improved survival in these cancers (Figure 9J). Collectively, these findings indicate that AREG production by NK cells impairs their anti-tumor efficacy across diverse malignancies.

## Discussion

The therapeutic potential of NK cells in cancer immunotherapy remains constrained, despite advances such as immune checkpoint blockade and ex vivo activation protocols^19^. Here, we identify a transcriptional and functional reprogramming of NK cells within skin TMEs, characterized by elevated AREG production. Tumor-associated NK cells exhibit heightened GR target gene signatures, correlating with AREG upregulation. Glucocorticoid-driven GR activation selectively induces AREG production in NK cells. NK-derived AREG suppresses NK cell cytotoxicity in vitro by inhibiting target cell apoptosis and compromises their tumor-restricting function in vivo in NCG mice. These findings establish a mechanistic link between TME-driven NK cell dysfunction and GR-associated transcriptional rewiring, offering insight into the compromised anti-tumor efficacy of NK cells in therapeutic settings.

The presence of NK cells co-expressing both anti-and pro-tumor transcriptional signatures highlights their functional plasticity in skin cancers (Figures 2A and 2D). This duality aligns with emerging evidence of immune cell “exhaustion” or “dysfunction” in the TME but extends this paradigm by implicating active transcriptional reprogramming through elevated GR activity in tumor-associated NK cells. The identification of GR signaling as the central regulator of AREG production provides a mechanistic explanation for this functional dichotomy (Figure 4). Notably, glucocorticoids–widely used to manage inflammation in cancer patients–may inadvertently exacerbate immune suppression by skewing NK cells toward AREG-dominant states. These findings highlight the need for caution in clinical glucocorticoid use and suggest GR antagonists (e.g., mifepristone) as potential adjuvants to preserve NK cell function.

The amplification of GR-driven AREG by PGE2 via PTGER2-cAMP-CREB signaling reveals a previously unrecognized crosstalk between prostaglandin and glucocorticoid pathways in immune evasion (Figures 5E-5H). This synergy, aligns with the established immunosuppressive role of PGE2 in impairing dendritic cell and T cell function^24,65,66,73^, extends its impact to NK cells as additional targets. The clinical correlation between AREG and COX-1/COX-2 expression in melanoma further supports targeting prostaglandin synthesis (e.g., COX inhibitors) as a strategy to disrupt this axis (Figure 5I).

GR activation establishes a persistent chromatin state in NK cells, priming them for enhanced AREG production upon subsequent stimulation. Bulk RNA-seq and ATAC-seq revealed that Dex treatment rapidly increases chromatin accessibility at the AREG promoter, an effect that persists for at least five days after Dex withdrawal, even as AREG mRNA and protein return to baseline (Figures 6C and S5E). This sustained chromatin remodeling creates a “primed” epigenetic state, enabling heightened AREG induction upon restimulation. These findings suggest that even transient glucocorticoid exposure (e.g., for chemotherapy side effect management) may have lasting effects on NK cell function. Modulating chromatin accessibility, such as with histone acetyltransferase (HAT) inhibitors, could reverse this priming and restore NK cell anti-tumor activity.

The finding that AREG impairs NK cell cytotoxicity by protecting tumor cells from apoptosis reconciles the paradox of retained IFN-γ production yet diminished anti-tumor efficacy in TMEs (Figure 7). AREG-deficient NK cells exhibit enhanced tumor control across multiple cancer models (Figures 8 and 9), while elevated AREG expression correlates with poor prognosis in diverse malignancies (Figure 9J), establishing NK cell-derived AREG as a therapeutic target. Strategies such as pharmacological GR antagonists to suppress AREG induction, COX/PGE2 pathway inhibitors to disrupt synergistic signaling, AREG-neutralizing antibodies to block its biological activity, or adoptive transfer of AREG-KO or GR-KO NK cells could enhance existing immunotherapies and improve clinical outcomes.

### Limitations of this study

While this study identified the immunosuppressive role of NK cell-derived AREG in skin cancer, we acknowledge that the heterogeneity of clinical skin cancer subtypes was not independently analyzed due to the limited sample size for each subtype in our cohort. Although AREG-KO NK cells demonstrated enhanced anti-tumor efficacy in both melanoma-and cSCC-derived tumors in NCG mice, further studies using single-cell multi-omics or spatial profiling are needed to dissect NK cell interactions with distinct TMEs (e.g., immune cell composition, metabolic landscapes). These insights could expand the therapeutic potential of AREG blockade beyond its universal role in NK cell dysfunction, informing subtype-adapted combinatorial strategies.

## Acknowledgments

This study was supported by National Natural Science Foundation of China (82471876), CAMS Innovation Fund for Medical Sciences (CIFMS) (2024-I2M-3-005) and Hospital for Skin Diseases, Institute of Dermatology, Chinese Academy of Medical Sciences and Peking Union Medical College grant (3301030103119) to Yetao W., National key research and development program (2022YFC2504700, 2022YFC2504701, 2022YFC2504705) to Yan W., The open project of Jiangsu provincial science and technology resources (Clinical Resources) coordination service platform (TC2022B016), The Jiangsu provincial natural science funds for young scholars (SBK2023041928) and The young scientists fund of the national natural science foundation of China (82304236) to Y.L.. We thank the study participants who provided skin and blood samples. Wei Cheng, Hao Chen and Zhiming Chen assisted in acquiring skin cancer and histology pictures. All staff in the biobank of Institute of Dermatology, Chinese Academy of Medical Sciences, Jiangsu Biobank of Clinical Resources assisted with clinical sample collection. Guangzhou Genedenovo Biotechnology Co., Ltd assisted with sequencing data analysis.

## Author contributions

Yetao Wang: Conceptualization; experiment design; data analysis; resources; supervision; validation; investigation; visualization; methodology; data curation; software; writing – original draft, review and editing; project administration; funding acquisition. Yan Wang: resources. Qin Wei: experiment design, performing experiments; data analysis; validation; visualization, data curation. Guirong Liang: performing experiments. Ruizeng: performing experiments; data analysis; validation. Yuancheng Li: data analysis; validation; visualization, software. Anlan Hong: resources. Hongsheng Wang: resources. Suying Feng: resources.

## Methods

### Cell culture and stimulation conditions

All cells were cultured at 37°C containing 5% CO_2_.

**Figures 3A-3K:** Dermal cells were stimulated with PMA (81 nM) and ionomycin (1.34 uM) (1:500, eBioscience, 00-4970-03) in RPMI 1640 for 2 hrs.

**Figures 4H and 4I:** PBMCs were stimulated with Dex (100 nM) in RPMI 1640 for 16 hrs.

**Figure 4K**: After electroporation for 48 hrs, NK cells were stimulated with Dex (100 nM) in RPMI 1640 for 16 hrs.

**Figures 4L and 4M:** PBMCs were stimulated with Dex (100 nM) in the presence or absence of RU486 (100 nM) in RPMI 1640 for 16 hrs.

**Figure 5G**: PBMCs were pre-treated with PF-04418948 (5 µM), L-161982 (5 µM), SQ22536 (10 µM) or 666-15 (1 µM), or left untreated for 6 hrs, then were treated with Dex (100 nM) alone or with PGE2 (1 µM) for 16 hrs in RPMI 1640.

**Figure 5H**: PMBCs incubated with or without Dex (100 nM) were treated with IL-12 (10 ng/ml) + IL-15 (50 ng/ml) + IL-18 (50 ng/ml) in the presence or absence of PGE2 (1 uM) for 16 hrs in RPMI 1640.

**Figures 6D and 6E:** PBMCs with and without primary Dex (100 nM) stimulation were stimulated with Dex (100 nM) after resting for 5 days in RPMI 1640 containing 5 ng/ml IL-15.

**Figures 7A-7D and 7F:** After electroporation for 48 hrs, NK cells were stimulated with IL-12 (10 ng/ml) + IL-15 (50 ng/ml) + IL-18 (50 ng/ml) in NK MACS medium for 16 hrs.

**Figure 7E**: The stimulated NK cells in Figure 7A were co-incubated with target cells (A375, A2058, Huh7, HepG2, or HCT116) in RPMI 1640 medium for 6 hrs.

**Figures 7G-7I:** A375, A2058, Huh7, HepG2, or HCT116 cells were incubated with DMEM, WT NK cell-conditioned medium, or AREG KO NK cell-conditioned medium for 48 hrs.

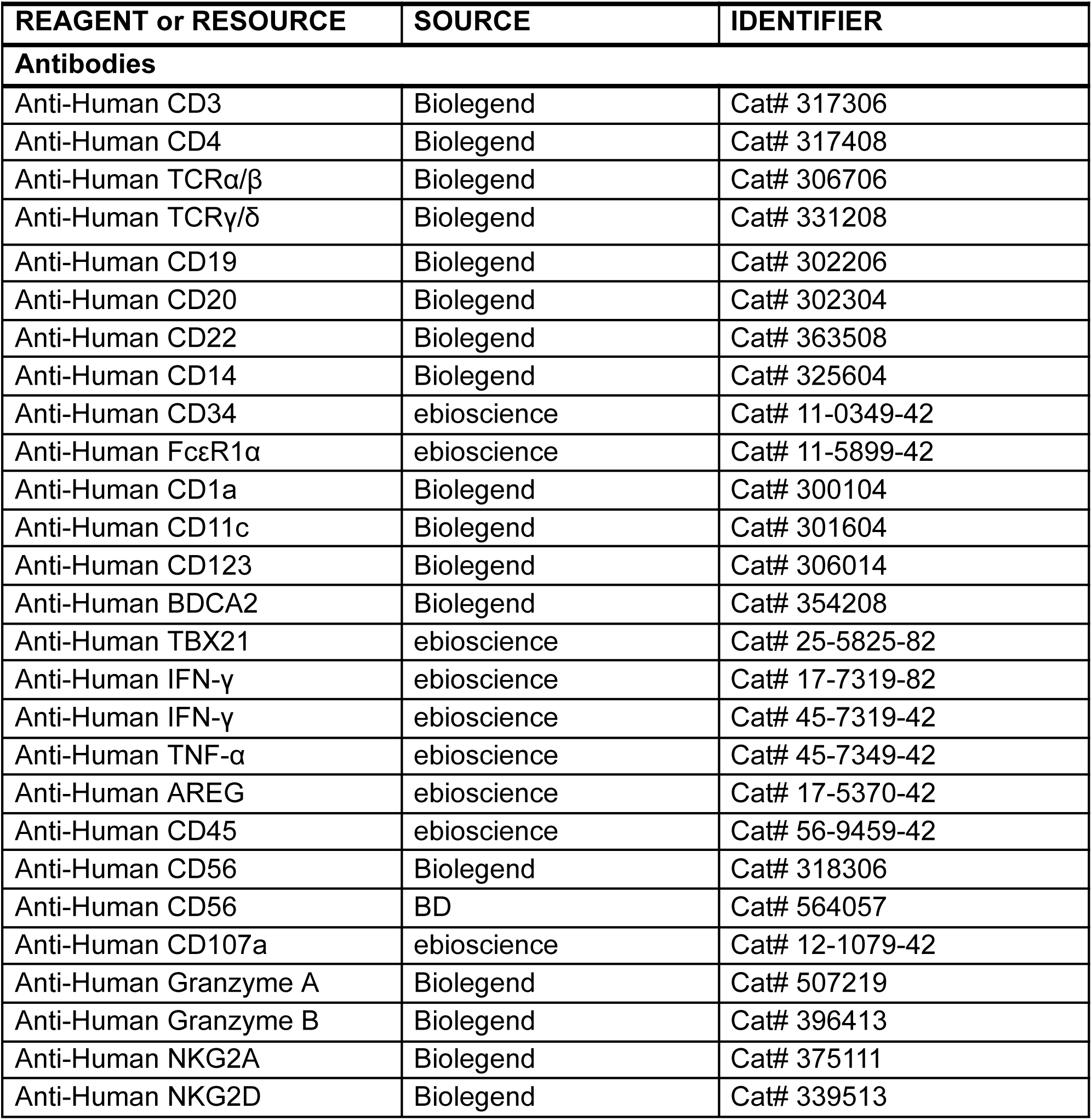
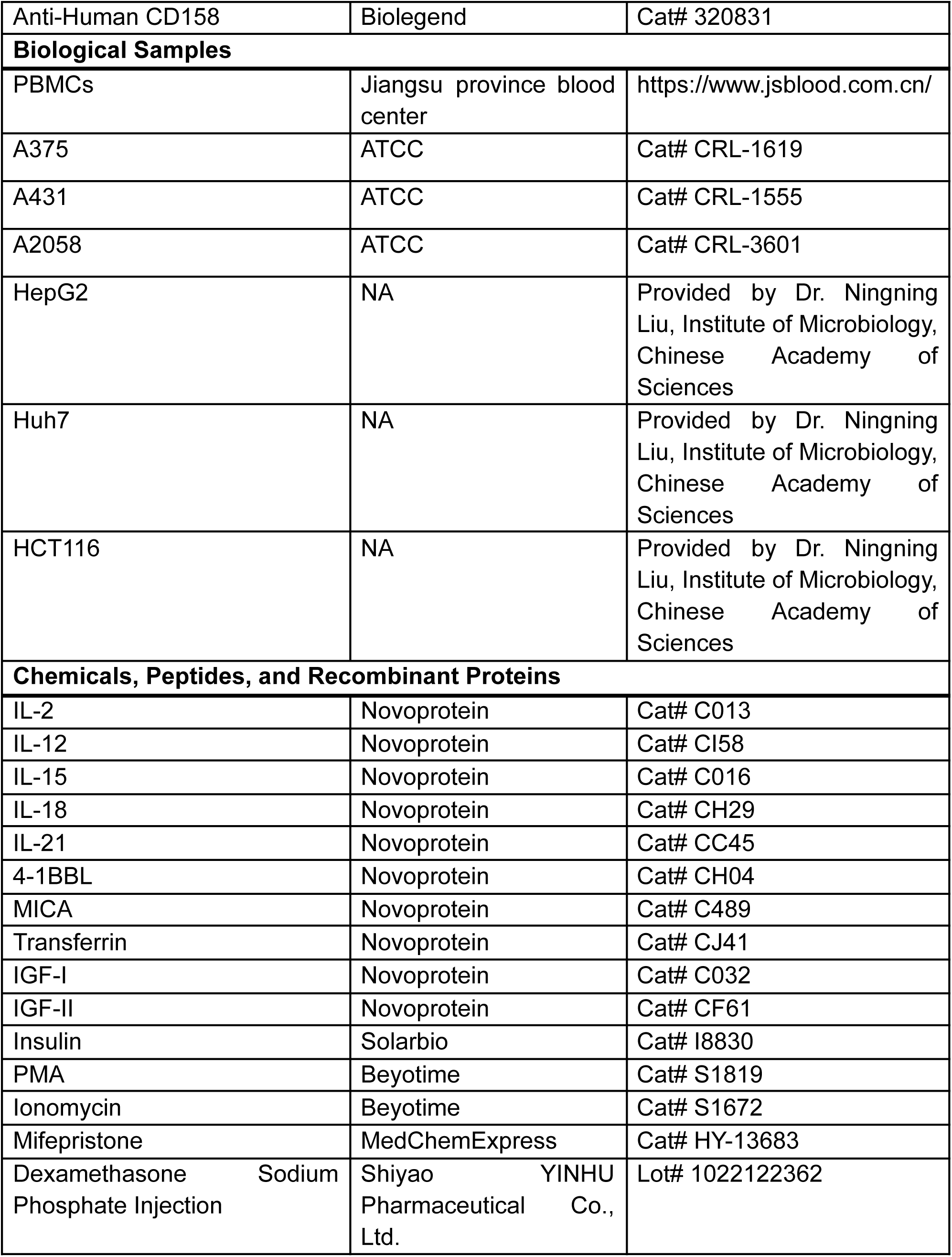
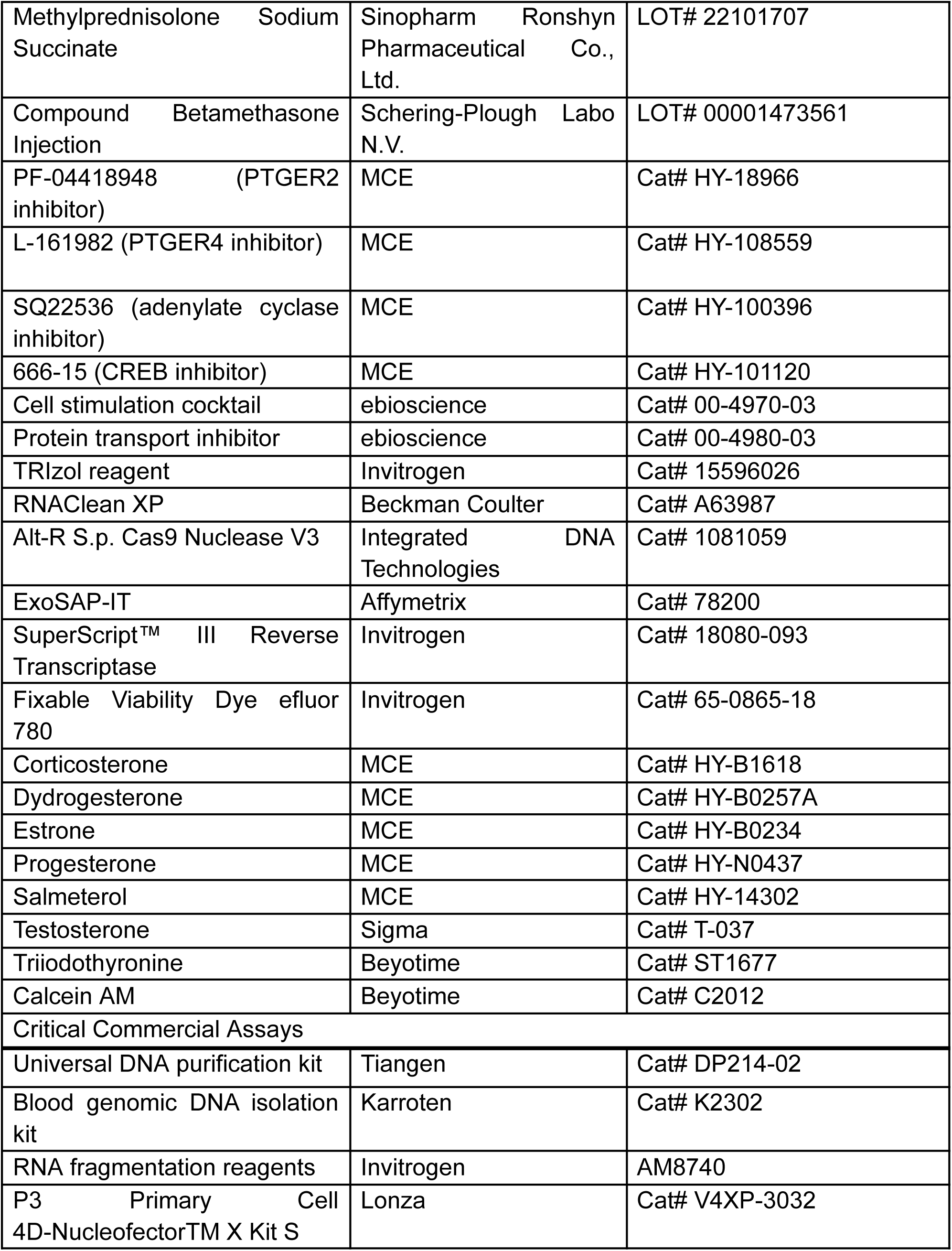
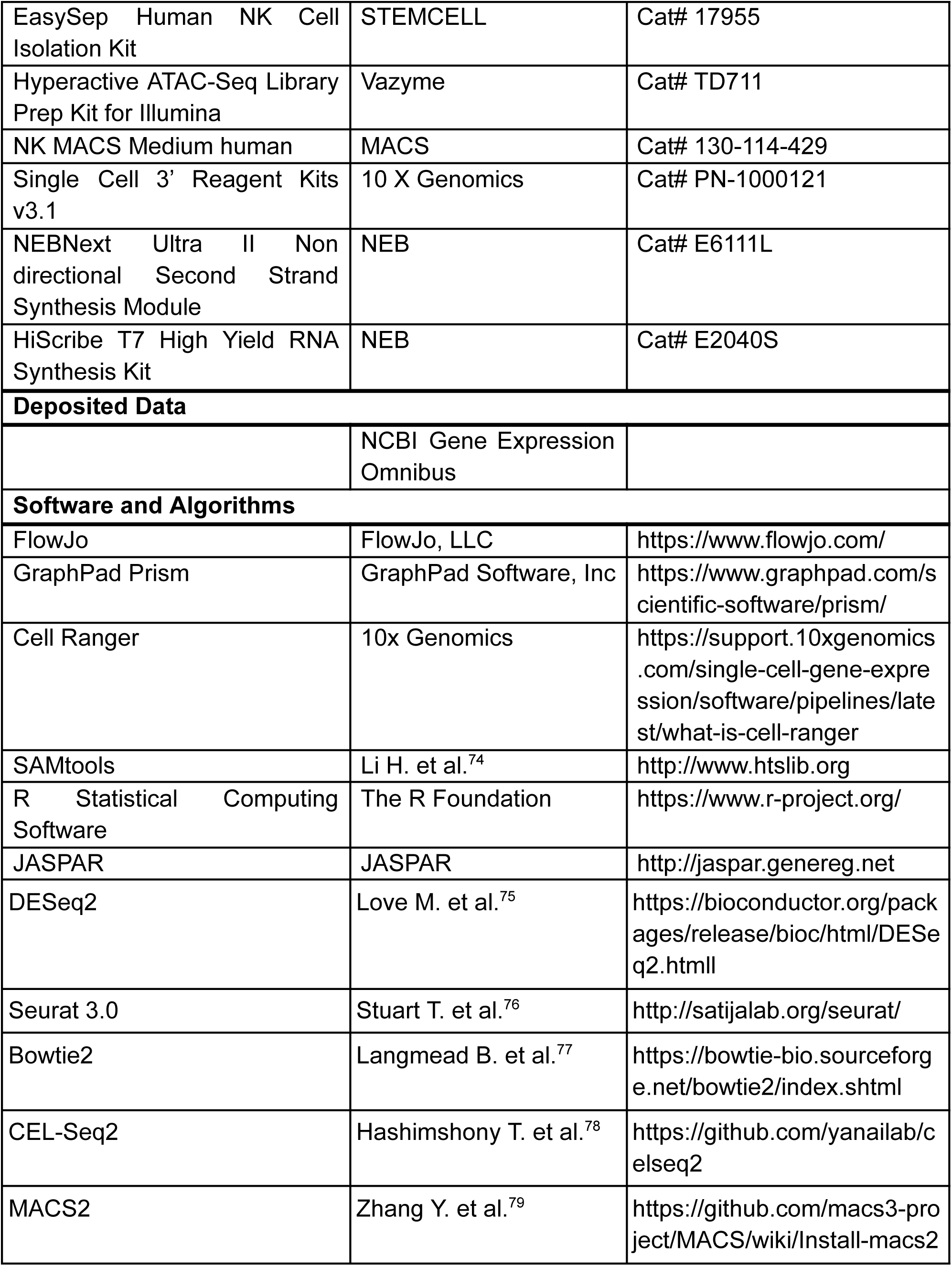
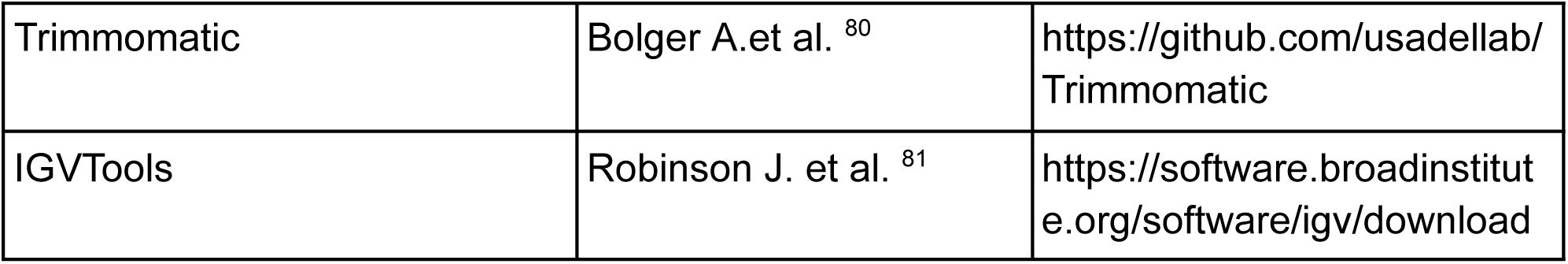

## Resource availability

Further information and requests for resources and reagents should be directed to and will be fulfilled by the lead contact, Yetao Wang.

## Materials availability

This study did not generate new unique reagents.

## Data availability

The datasets produced in this study, including scRNA-Seq data of skin tumor and peri-tumor, and bulk RNA-Seq data of Dex treated and untreated NK cells for 16 hrs, are available in NCBI Gene Expression Omnibus (GEO) with accession number: GSE242941.

Bulk RNA-Seq data of Dex-primarily treated and untreated NK cells after 5 days of rest, and ATAC-Seq data of Dex-primarily treated and untreated NK cells after 16 hrs and after 5 days of rest, are available in the NCBI Sequence Read Archive (SRA), BioProject: PRJNA1119074.

### Clinical samples

Skin tumor and peri-tumor samples were obtained from the biobank of Institute of Dermatology, Chinese Academy of Medical Sciences, Jiangsu Biobank of Clinical Resources (BM2015004). Blood samples were obtained from Jangsu province blood center. All participants provided written informed consent for protocols that were included in the study of cellular immunity in skin cancers, in accordance with procedures approved by the ethics committee of Institute of Dermatology, Chinese Academy of Medical Sciences and Peking Union Medical College. The clinical characteristics including sex, age and skin cancer types for each donor of skin cancer were provided in Figure S1 and Table S1. No blinding of investigators or subjects during the conduct or analysis of the study. No power analysis was used to calculate the group size in this study.

### Human skin cell preparation

Isolation of dermal single cells from peri-tumor and tumor tissues for *in vitro* culture and flow cytometry: 1. After scraping off the subcutaneous fat tissue, skin biopsies were transferred to 1 ml dispase (5 U/mL in PBS containing 1% penicillin/streptomycin) and were incubated in 37°C for 2 hrs to separate epidermis from dermis; 2. Dermis was washed briefly in PBS and was transfer to dermis digestion solution (Collagenase III 3mg/ml + DNase (5 ug/ml) in 10% FBS/RPMI 1640) at 37°C for 2 hrs, shaking vigorously every 30 min; 3. The digested dermis was filtered through a 70 um strainer, the flow through was collected in a 15 ml tube, after centrifuge at 700 x g for 3min, the cells were resuspended in 500 ul MACS buffer (0.5% BSA and 2 mM EDTA in PBS), ready for use.

Isolation of skin CD45^+^ cells for scRNA-Seq: Skin biopsies were digested according to the instruction of the whole skin dissociation kit (130-101-540, MACS). Briefly, after scraping off fat tissue, skin biopsies were cutted into small pieces (< 4 mm^2^), each piece was digested with a buffer containing 435 ul buffer L, 12.5 ul enzyme P, 50 ul enzyme D and 2.5 ul enzyme A on 37°C shaker for 3 hrs. The digested skin tissue was diluted with ice cold DMEM containing 2% BSA, then was filtered through a 70 um strainer, the flow through was centrifuged at 300 x g for 10 min at 4°C. The isolated skin cells were resuspended in MACS buffer and were subjected to CD45^+^ cell enrichment using magnetic beads (130-045-801, MACS).

### Peripheral blood mononuclear cells (PBMCs) isolation

Each human peripheral blood leukopak was washed with 50 ml PBS, and was overlaid on lymphoprep (STEMSELL, 07851), then was centrifuged at 500 x g for 30 min in room temperature. After washing with MACS buffer for 3 times, the isolated PBMCs were either used immediately or frozen in FBS containing 10% DMSO.

### Flow cytometry

Skin cells or PBMCs were first stained with fixable viability dye efluor 780 (Invitrogen, 65-0865-18). For surface staining, cells were incubated in MACS buffer with antibodies at 4°C for 30 min in the dark. For intracellular staining, cells were fixed and permeabilized using Foxp3 staining kit (eBioscience, 00-5523-00), then cytokines or TBX21 were stained with antibodies in the permeabilization buffer at 4°C for 30 min in the dark. After washing with MACS buffer, the cells were ready for flow cytometry detection. For samples that do not contain enough cells to analyze were dropped out.

### NK cell sorting

PBMCs were stained with fixable viability dye efluor 780, lineage antibodies (against CD3, CD4, TCRαβ, TCRγδ, CD19, CD20, CD22, CD34, FcεRIα, CD11c, CD303, CD123, CD1a, and CD14) and antibody against CD56. CD56^dim^NK cells were sorted as indicated in Figure S5A using BD FACSAria IIu. The sorted NK cells were subjected to bulk RNA-Seq and ATAC-Seq library preparation.

### scRNA-Seq library preparation

The scRNA-Seq libraries were prepared by Single Cell 3’ Reagent Kits v3.1 (10 x Genomics, PN-1000121). The enriched skin CD45^+^ cells were washed and resuspended in 1 x PBS containing 0.05% BSA. Cell number and viability were measured by trypan blue staining under microscope, cell concentration was adjusted to 1,000-1,500 cells/ul (viability > 90%). Single cell suspension was loaded onto Chromium Controller (10 x Genomics) to participate 8,000–10,000 single cells into gel beads in emulsions (GEMs). The quality of amplified cDNA and final sequencing library were measured by Agilent 2100 Expert (Agilent Technologies).The sequencing depth was controlled around 50,000 mean reads/cell. The libraries were sequenced by Illumina NovaSeq 6000.

### Bulk RNA-Seq library preparation

The bulk RNA-Seq libraries of sorted Dex treated and untreated CD56^dim^NK cells were prepared using CEL-Seq2^78^. Total RNA of sorted cells was extracted using TRIzol reagent (ThermoFisher, 15596026). 100 ng RNA for each library was used for first strand cDNA synthesis using barcoded primers as follows (barcode underlined): Dex untreated repeat 1: 5’-GCCGGTAATACGACTCACTATAGGGAGTTCTACAGTCCGACGATCNNNNNN<u>AGA CTC</u>TTTTTTTTTTTTTTTTTTTTTTTTV-3’; Dex untreated repeat 2: 5’-GCCGGTAATACGACTCACTATAGGGAGTTCTACAGTCCGACGATCNNNNNN<u>CATG AG</u>TTTTTTTTTTTTTTTTTTTTTTTTV-3’; Dex untreated repeat 3: 5’-GCCGGTAATACGACTCACTATAGGGAGTTCTACAGTCCGACGATCNNNNNN<u>CAG</u> <u>ATC</u>TTTTTTTTTTTTTTTTTTTTTTTTV-3’; Dex treated repeat 1: 5’-GCCGGTAATACGACTCACTATAGGGAGTTCTACAGTCCGACGATCNNNNNN<u>AGCT AG</u>TTTTTTTTTTTTTTTTTTTTTTTTV-3’; Dex treated repeat 2: 5’-GCCGGTAATACGACTCACTATAGGGAGTTCTACAGTCCGACGATCNNNNNN<u>CATG CA</u>TTTTTTTTTTTTTTTTTTTTTTTTV-3’; Dex treated repeat 3: 5’-GCCGGTAATACGACTCACTATAGGGAGTTCTACAGTCCGACGATCNNNNNN<u>TCAC AG</u>TTTTTTTTTTTTTTTTTTTTTTTTV-3’; Cytokine stimulated, Dex untreated repeat 1: 5’-GCCGGTAATACGACTCACTATAGGGAGTTCTACAGTCCGACGATCNNNNNN<u>AGCT CA</u>TTTTTTTTTTTTTTTTTTTTTTTTV-3’; Cytokine stimulated, Dex untreated repeat 2: 5’-GCCGGTAATACGACTCACTATAGGGAGTTCTACAGTCCGACGATCNNNNNN<u>CATG TC</u>TTTTTTTTTTTTTTTTTTTTTTTTV-3’; Cytokine stimulated, Dex untreated repeat 3: 5’-GCCGGTAATACGACTCACTATAGGGAGTTCTACAGTCCGACGATCNNNNNN<u>AGG ATC</u>TTTTTTTTTTTTTTTTTTTTTTTTV-3’; Cytokine stimulated, Dex treated repeat 1: 5’-GCCGGTAATACGACTCACTATAGGGAGTTCTACAGTCCGACGATCNNNNNN<u>AGCT TC</u>TTTTTTTTTTTTTTTTTTTTTTTTV-3’; Cytokine stimulated, Dex treated repeat 2: 5’-GCCGGTAATACGACTCACTATAGGGAGTTCTACAGTCCGACGATCNNNNNN<u>CACT AG</u>TTTTTTTTTTTTTTTTTTTTTTTTV-3’; Cytokine stimulated, Dex treated repeat 3: 5’-GCCGGTAATACGACTCACTATAGGGAGTTCTACAGTCCGACGATCNNNNNN<u>AGT GCA</u>TTTTTTTTTTTTTTTTTTTTTTTTV-3’. The second strand was synthesized using NEBNext Second Strand Synthesis Module (NEB, E6111L). The dsDNA was purified using RNAClean XP (Beckman Coulter, A63987) and was subjected to *in vitro* transcription (IVT) using HiScribe T7 High Yield RNA Synthesis Kit (NEB, E2040S). After ExoSAP-IT (Affymetrix, 78200) treatment, the IVT RNA was fragmented using RNA fragmentation reagents (Invitrogen, AM8740) and was subjected to the second round of reverse transcription using random hexamer: 5’-GCCTTGGCACCCGAGAATTCCANN NNNN-3’ The final library was amplified with indexed primers: RP1: 5’-AATGATACGGCGACCACCGAGATCTACACGTTCAGAGTTCTACAGTCCGA-3’ and RPI1: 5’-CAAGCAGAAGACGGCATACGAGATCGTGATGTGACTGGAGTTCCTTGGCACCCGAGAATTCCA-3’. After quality check, the purified libraries were sequenced by Illumina NovaSeq 6000.

### ATAC-Seq library preparation

The ATAC-Seq library for NK cells was constructed using the Hyperactive ATAC-Seq Library Prep Kit for Illumina (Vazyme, TD711). Briefly, the nuclei of sorted NK cells were isolated using a lysis buffer. Fragmentation buffer with Tn5 transposase was added, and the mixture was incubated at 37°C for 30 minutes. The released DNA fragments were then purified using ATAC DNA extraction beads. The library was subsequently amplified and purified with ATAC DNA clean beads.

### Cas9 RNP mediated knockout in NK cells

NK cells were isolated from PBMCs using EasySep human NK Cell isolation kit (STEMCELL, 17955) and were expanded in NK MACS medium (MACS, 130-114-429) for 10 days to render NK cells competent for electroporation. 200 pmol sgRNA (synthesized from GenScript) were mixed with 100 pmol Cas9 (IDT, 1081059) for 20 min in room temperature to form RNP. One million NK cells were resuspended in 20 ul 4D nucleofector master mix (82% P3 + 18% supplement 1; Lonza, V4XP-3032), and mixed with Cas9 RNP for electroporation using program CM137. The electroporated NK cells were ready for downstream experiments after culturing in NK MACS medium for 48 hrs. The target site for CD19 knockout (control) was 5’-CTAGGTCCGAAACATTCCAC-3’; for AREG knockout was 5’-GAGGACGGTTCACTACTAGA-3’; and for NR3C1 knockout was 5’-TTACATTGGTCGTACATGCA-3’, the corresponding sgRNAs were synthesized from IDT.

### scRNA-Seq data processing

An average 9043 cells per sample, 49,004 reads and 1243 median genes per cell were identified by cell ranger (10x Genomics, version 5.0.0) (Table S2). Following alignment, cells meeting the following criterias were retained using Seurat (Version 3.0) and were passed to downstream analysis: (1) nFeature range from 200 to 5900; (2) < 49000 UMIs; (3) < 35% UMIs of mitochondria genes. Potential doublets and multiplets were filtered by DoubletFinder, and 196,278 cells were integrated. The exclusive cell type specific markers were used to further exclude doublets and multiplets. CD45 negative non-hematopoietic cell clusters including keratinocytes (KRT1, KRT5, KRT10, KRT14), fibroblasts (MFAP5, PDPN, PDGFRA, COL1A1), endothelial cells (ACKR1, VWF, PECAM1) and pericytes (RGS5, ACTA2, TAGLN) were also excluded based on their cell type specific markers. Total 146,643 CD45 positive cells were re-clusterd into 8 unique clusters (res=0.5) and NK cell cluster was focused for further analysis.

To identify cluster highly expressed genes, the gene expression values for the cluster of interest was compared with the rest clusters using FindMarkers in Seurat by Model-based Analysis of Single-cell Transcriptomics (MAST) test, the average counts for each cluster were calculated by AverageExpression function.

### Gene sets score analysis

The AddModuleScore function in Seurat was used to calculate the gene set score of NK cell clusters, and the significance was determined by Wilcoxon rank-sum test. Signature genes that were used for gene sets score analysis were included in Table S5.

### Gene set correlation analysis

The AUCell score of each gene set, including GR pathway activity, tumor stress or NK cell activity, in each cell were calculated, the correlation among gene sets was calculated by Pearson correlation. Cells with extreme values (count = 0) were discarded from the analysis.

### Bulk RNA-Seq analysis

The transcriptome count matrix of all samples was generated using the default settings of CEL-Seq2 pipeline (https://github.com/yanailab/celseq2)^78^. Briefly, Read 2 was assigned to each sample based on their paired read 1 barcode and was mapped to hg19 using Bowie2. The UMIs for each sample were counted, and the generated count matrix was analyzed by DEseq2 package in R. The counts of transcripts per sample were normalized using variance stabilizing transformation (VST) method in DESeq2. The DEgenes were determined by DEseq2 (Log_2_FC> 0.5, *P*<0.01).

### ATAC-Seq analysis

Paired-end reads were filtered with Trimmomatic and aligned to the hg38 reference genome using Bowtie2. Duplicates were removed with Picard’s MarkDuplicates. Each aligned read was trimmed to the first 9 bases at the 5’ end to match the Tn5 transposase cut site. For peak smoothing, the start sites of trimmed reads were extended 10 bases upstream and downstream. Peaks were called with MACS2. Adjusted aligned reads were converted to TDF files for visualization using IGVTools.

### Killing assay

A375, A2058, Huh7, HePG2 or HCT116 cells were washed and resuspended in PBS at 10^6^ cells/ml. Calcein AM was added at a final concentration of 15 μM, and cells were labelled at 37°C in 5% CO_2_ for 30 min. After washing twice with PBS, the labelled cells were resuspended in RPMI 1640 and aliquoted in a V bottom 96 well plate at 1×10^4^ cells in 100 μl per well. The sorted NK cells were resuspended in RPMI 1640 and the concentration was adjusted at 20 fold of target cells (2×10^5^ cells) in 50μl. The effector and target cells were mixed and centrifuged at 50 × g for 0.5 min and were incubated at 37°C in 5% CO_2_ for 6 hrs. Cells were pelleted and 75 μl supernatant was transferred to a 96-well solid black microplate. The fluorescence released by labelled target cells were detected using BioTek Synergy H1 plate reader (excitation: 485 nm, emission: 528 nm). Specific lysis was determined as: [(test fluorescence release - spontaneous fluorescence release) / (maximum fluorescence release - spontaneous fluorescence release)] × 100.

### Survival probability analysis

Survival and RNA-seq data from TCGA (The Cancer Genome Atlas) PanCancer project were extracted from the UCSC XENA database (https://xena.ucsc.edu/). The Survival package was to compute Kaplan Meier curves for each tumor and to calculate the log-rank test to obtain the survival probability.

### Mouse experiments

All NCG mice (NOD/ShiLtJGpt-*Prkdc^em26Cd52^*Il2rg*^em26Cd22^/Gpt,* GemPharmatech, T001475) were kept in microisolator cages and provided with autoclaved food and acidified, autoclaved water within a specific pathogen-free facility. The use of animals followed the guidelines of the Institutional Animal Care and Use Committee of the Hospital for Skin Diseases, Institute of Dermatology, Chinese Academy of Medical Sciences and Peking Union Medical College. All experimental protocols were reviewed and approved by this committee.

### Statistical analysis

Statistical test was performed using GraphPad Prism 9. Wilcoxon matched-pairs signed rank test or two tailed paired t-test used in this study were specified in the figure legends. Variance was estimated by calculating the mean ± s.e.m. in each group. *P* < 0.05 was considered significant. For gene sets score analysis, the *P* value was determined by Wilcoxon rank-sum test using R software. P < 0.05 was considered significant.

## Supplemental Figure legends

**Figure S1.**
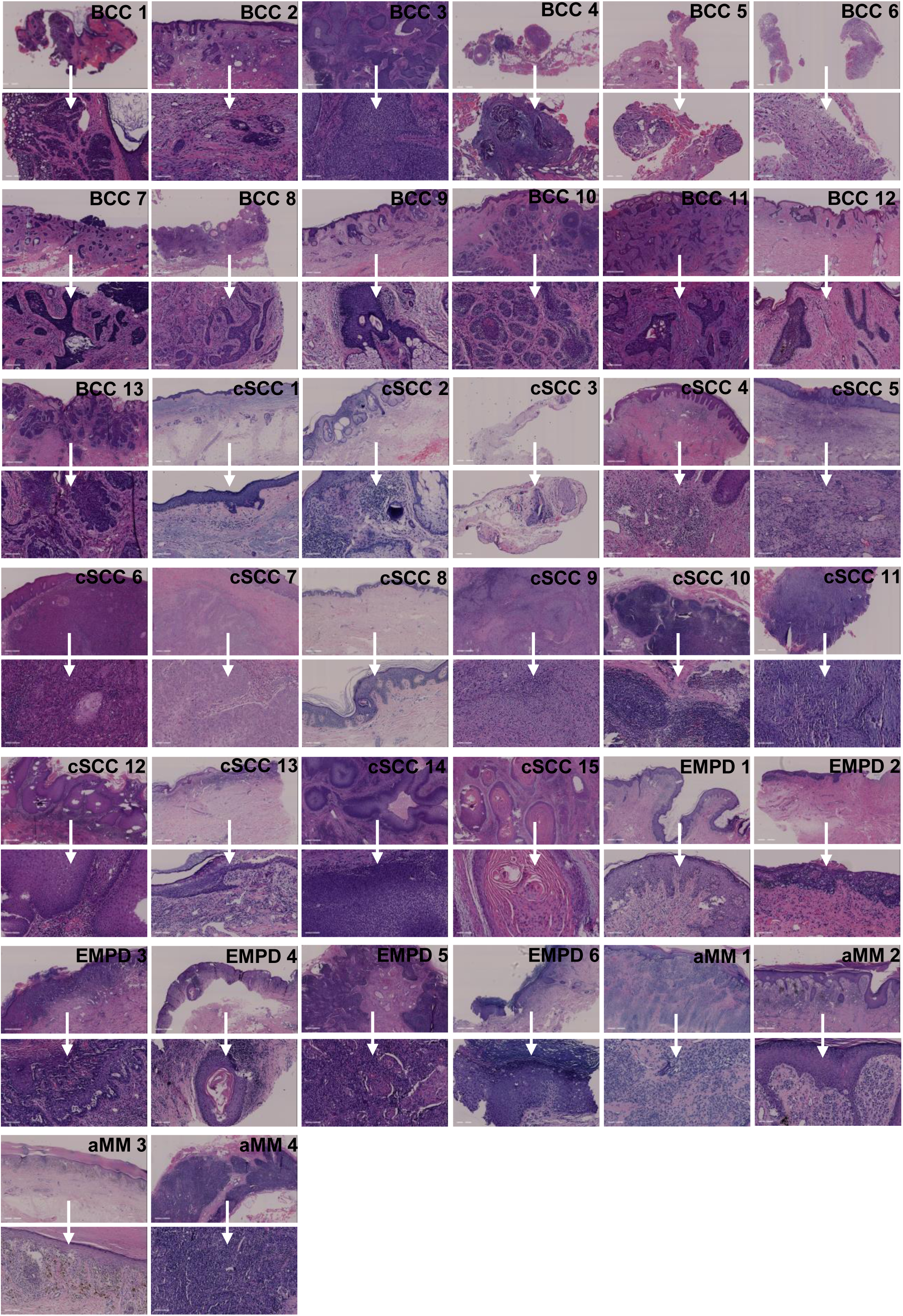
Histological pictures of skin cancer donors. Histological pictures of skin cancer donors for BCC (n=13), cSCC (n=15), EMPD (n=6) and aMM (n=4).

**Figure S2.**
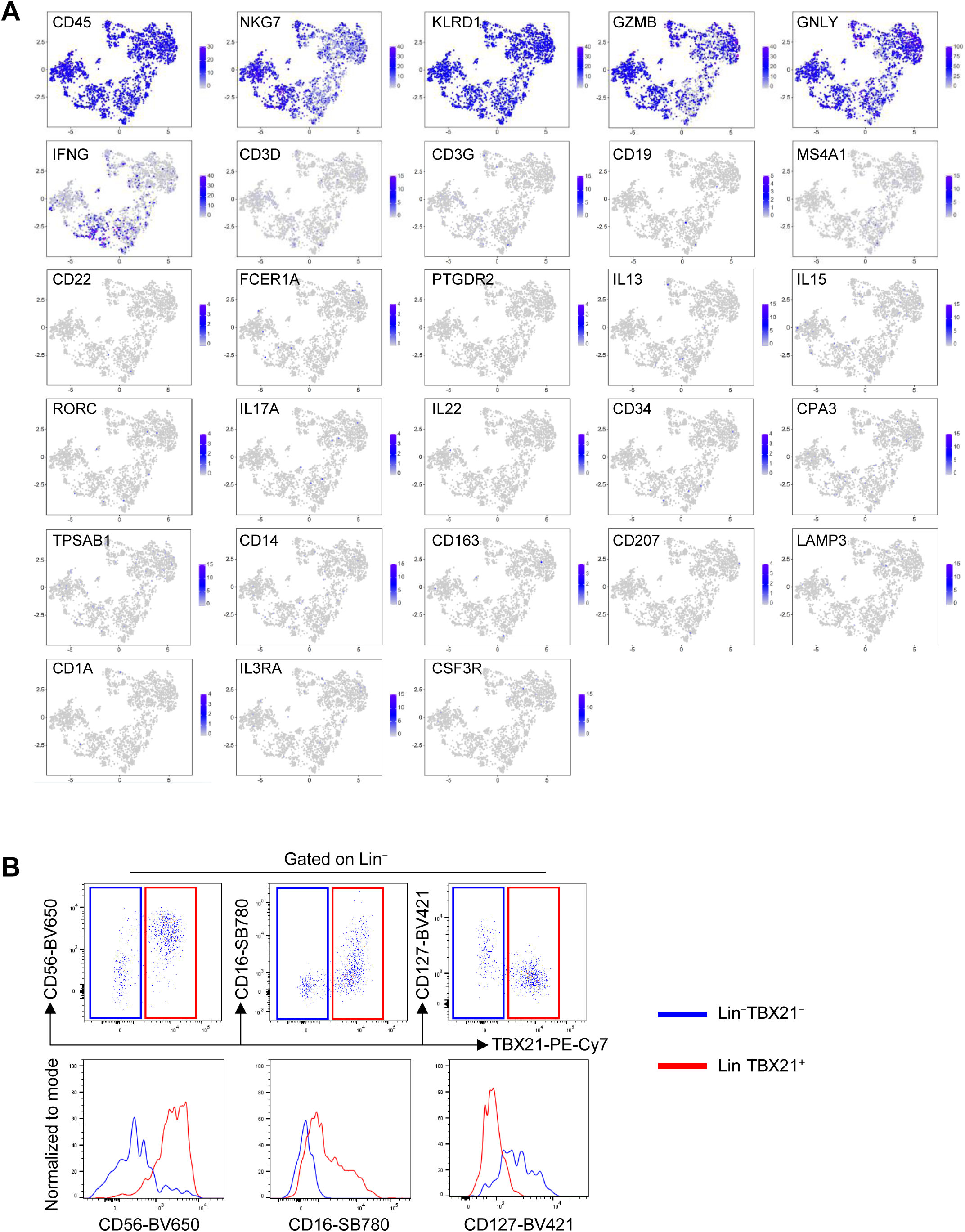
Detection of NK cells by scRNA-Seq. (A) The expression of markers for NK and other cell types in NK cells clusters. (B) CD56, CD16, and CD127 were detected in the Lin^-^TBX21^+^ or Lin^-^TBX21^-^ population in the BCC dermis by flow cytometry.

**Figure S3.**
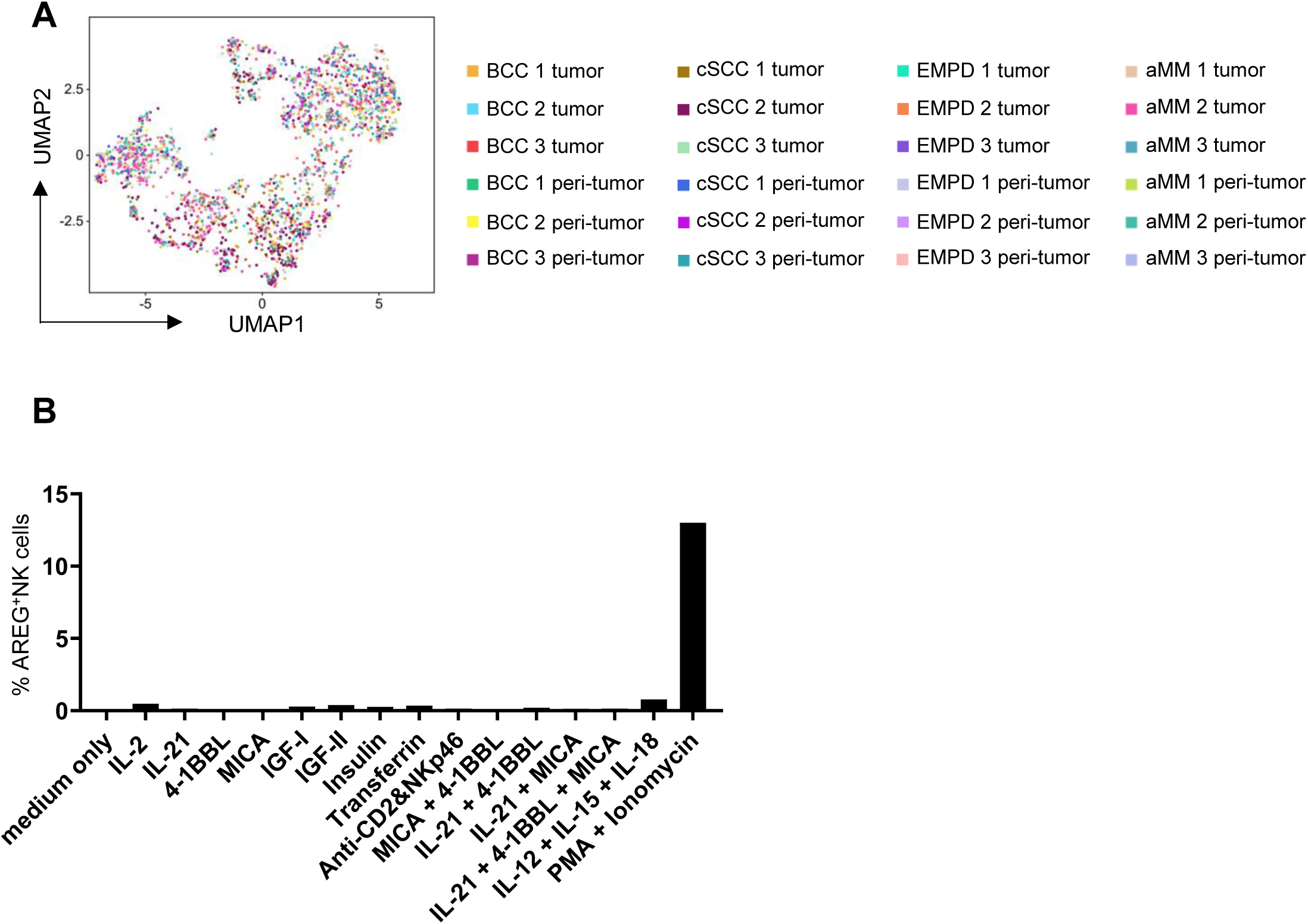
Cluster unique DEgenes of NK cells in skin cancers. **(A)** Distribution of NK cells in skin tumors and peri-tumor regions across donors. (B) PBMCs were stimulated in indicated conditions for 16 hrs, the AREG production from NK cells (Lin^-^TBX21^+^) were detected using flow cytometry.

**Figure S4.**
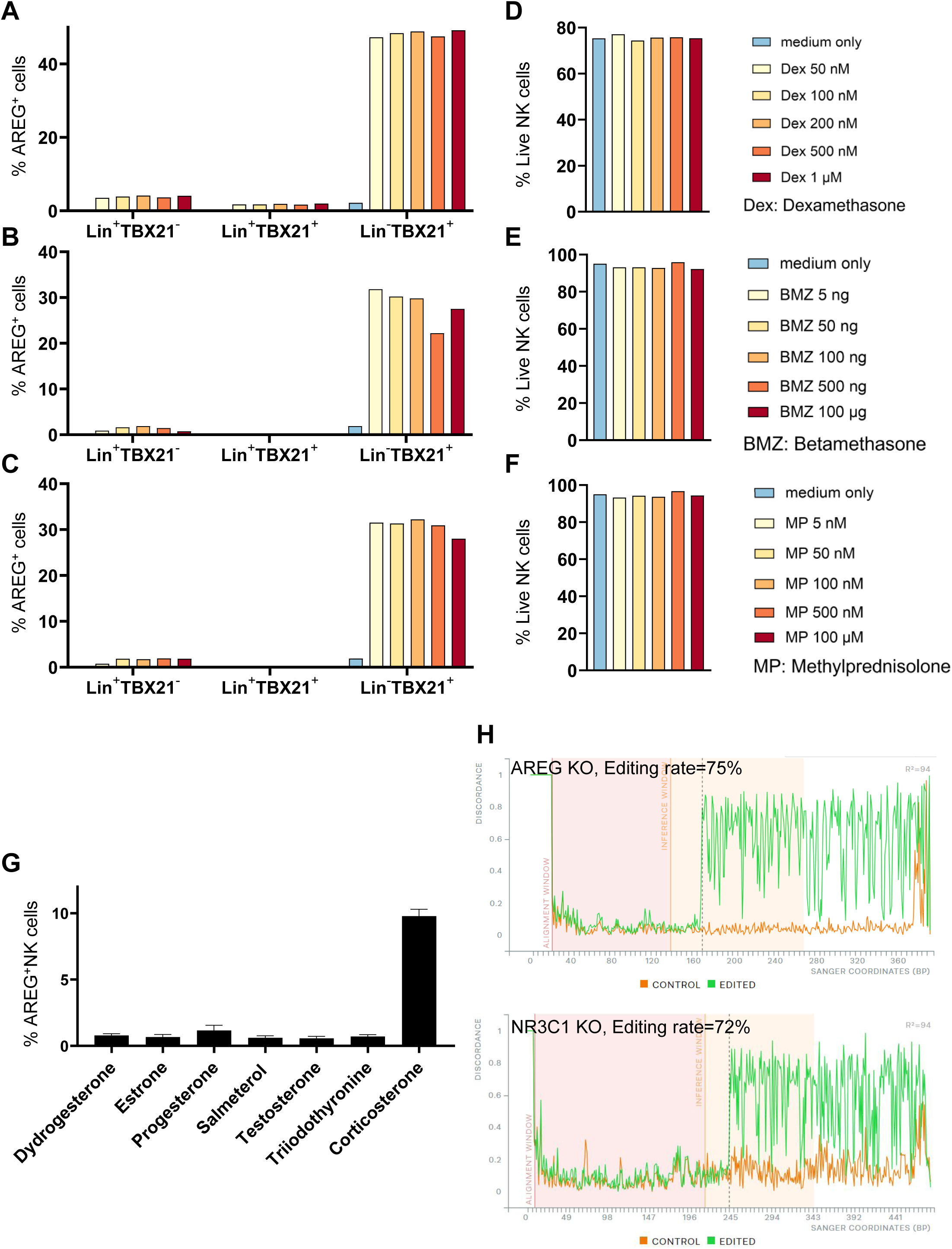
Glucocorticoid stimulation induces NK cell AREG production. (A-C) PBMCs were stimulated in different doses of dexamethasone (A), betamethasone (A) and methylprednisolone (C) for 16 hrs, NK cells produced AREG were detected using flow cytometry. (D-F) Proportion of live NK cells corresponding to (A–C). (G) PBMCs were treated with indicated hormones or hormone analogs for 16 hrs, NK cells produced AREG were detected by flow cytometry. (H) ICE analysis for knockout effect of AREG and NR3C1 in NK cells.

**Figure S5.**
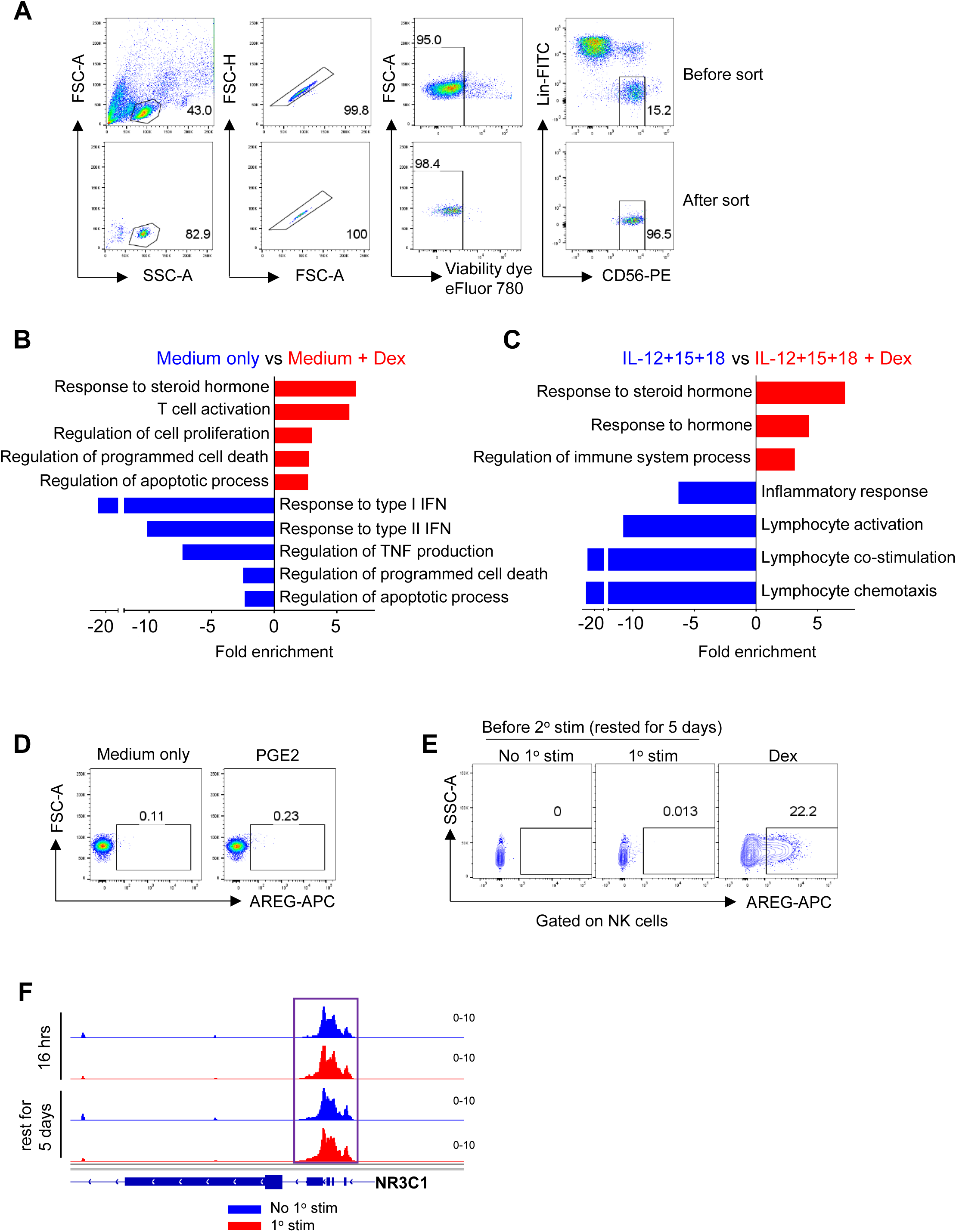
GR activation induces transcriptional alteration in NK cells. (A) Flow cytometry analysis of PBMC-derived NK cells before and after sorting. (B) Enriched pathways of Dex up (red) and down (blue) regulated genes by GO enrichment analysis. (C) Enriched pathways of Dex up (red) and down (blue) regulated genes in the condition of IL-12+IL-15+IL-18 stimulation by GO enrichment analysis. (D) PMBCs were treated with or without PGE2 for 16 hrs, AREG in the NK cell was detected by flow cytometry. (E) PBMCs with or without primary Dex stimulation were cultured in low dose IL-15 for 5 days, before secondary Dex stimulation, the production of AREG by NK cells were detected by flow cytometry. (F) ATAC-Seq analysis of NR3C1 loci of sorted blood NK cells with or without primary Dex stimulation for 16 hrs, and after rest for 5 days.

## References

1. Henrikson, N.B., Ivlev, I., Blasi, P.R., Nguyen, M.B., Senger, C.A., Perdue, L.A., and Lin, J.S. (2023). Skin Cancer Screening: Updated Evidence Report and Systematic Review for the US Preventive Services Task Force. JAMA 329, 1296–1307.

2. Tanese, K., Nakamura, Y., Hirai, I., and Funakoshi, T. (2019). Updates on the Systemic Treatment of Advanced Non-melanoma Skin Cancer. Front. Med. 6, 160.

3. Arnold, M., Singh, D., Laversanne, M., Vignat, J., Vaccarella, S., Meheus, F., Cust, A.E., de Vries, E., Whiteman, D.C., and Bray, F. (2022). Global Burden of Cutaneous Melanoma in 2020 and Projections to 2040. JAMA Dermatol. 158, 495–503.

4. Stonesifer, C.J., Djavid, A.R., Grimes, J.M., Khaleel, A.E., Soliman, Y.S., Maisel-Campbell, A., Garcia-Saleem, T.J., Geskin, L.J., and Carvajal, R.D. (2021). Immune Checkpoint Inhibition in Non-Melanoma Skin Cancer: A Review of Current Evidence. Front. Oncol. 11, 734354.

5. Jenkins, R.W., and Fisher, D.E. (2021). Treatment of Advanced Melanoma in 2020 and Beyond. J. Invest. Dermatol. 141, 23–31.

6. Ji, A.L., Rubin, A.J., Thrane, K., Jiang, S., Reynolds, D.L., Meyers, R.M., Guo, M.G., George, B.M., Mollbrink, A., Bergenstråhle, J., et al. (2020). Multimodal Analysis of Composition and Spatial Architecture in Human Squamous Cell Carcinoma. Cell 182, 1661–1662.

7. Ozbay Kurt, F.G., Lasser, S., Arkhypov, I., Utikal, J., and Umansky, V. (2023). Enhancing immunotherapy response in melanoma: myeloid-derived suppressor cells as a therapeutic target. J. Clin. Invest. 133. 10.1172/JCI170762.

8. Tjiu, J.-W., Chen, J.-S., Shun, C.-T., Lin, S.-J., Liao, Y.-H., Chu, C.-Y., Tsai, T.-F., Chiu, H.-C., Dai, Y.-S., Inoue, H., et al. (2009). Tumor-associated macrophage-induced invasion and angiogenesis of human basal cell carcinoma cells by cyclooxygenase-2 induction. J. Invest. Dermatol. 129, 1016–1025.

9. 9. Lorenzo-Sanz, L., Lopez-Cerda, M., da Silva-Diz, V., Artés, M.H., Llop, S., Penin, R.M., Bermejo, J.O., Gonzalez-Suarez, E., Esteller, M., Viñals, F., et al. (2024). Cancer cell plasticity defines response to immunotherapy in cutaneous squamous cell carcinoma. Nat. Commun. 15, 5352.

10. Zheng, L., Qin, S., Si, W., Wang, A., Xing, B., Gao, R., Ren, X., Wang, L., Wu, X., Zhang, J., et al. (2021). Pan-cancer single-cell landscape of tumor-infiltrating T cells. Science 374, abe6474.

11. Yost, K.E., Satpathy, A.T., Wells, D.K., Qi, Y., Wang, C., Kageyama, R., McNamara, K.L., Granja, J.M., Sarin, K.Y., Brown, R.A., et al. (2019). Clonal replacement of tumor-specific T cells following PD-1 blockade. Nat. Med. 25, 1251–1259.

12. Chiang, E., Stafford, H., Buell, J., Ramesh, U., Amit, M., Nagarajan, P., Migden, M., and Yaniv, D. (2023). Review of the tumor microenvironment in basal and squamous cell carcinoma. Cancers (Basel) 15. 10.3390/cancers15092453.

13. Kaporis, H.G., Guttman-Yassky, E., Lowes, M.A., Haider, A.S., Fuentes-Duculan, J., Darabi, K., Whynot-Ertelt, J., Khatcherian, A., Cardinale, I., Novitskaya, I., et al. (2007). Human basal cell carcinoma is associated with Foxp3+ T cells in a Th2 dominant microenvironment. J. Invest. Dermatol. 127, 2391–2398.

14. Jeon, S., Jeon, M., Choi, S., Yoo, S., Park, S., Lee, M., and Kim, I. (2023). Hypoxia in skin cancer: Molecular basis and clinical implications. Int. J. Mol. Sci. 24, 4430.

15. D’Aguanno, S., Mallone, F., Marenco, M., Del Bufalo, D., and Moramarco, A. (2021). Hypoxia-dependent drivers of melanoma progression. J. Exp. Clin. Cancer Res. 40, 159.

16. Zhou, L., Wang, Y., Zhou, M., Zhang, Y., Wang, P., Li, X., Yang, J., Wang, H., and Ding, Z. (2018). HOXA9 inhibits HIF-1α-mediated glycolysis through interacting with CRIP2 to repress cutaneous squamous cell carcinoma development. Nat. Commun. 9. 10.1038/s41467-018-03914-5.

17. Forsthuber, A., Aschenbrenner, B., Korosec, A., Jacob, T., Annusver, K., Krajic, N., Kholodniuk, D., Frech, S., Zhu, S., Purkhauser, K., et al. (2024). Cancer-associated fibroblast subtypes modulate the tumor-immune microenvironment and are associated with skin cancer malignancy. Nat. Commun. 15, 9678.

18. Barry, K.C., Hsu, J., Broz, M.L., Cueto, F.J., Binnewies, M., Combes, A.J., Nelson, A.E., Loo, K., Kumar, R., Rosenblum, M.D., et al. (2018). A natural killer–dendritic cell axis defines checkpoint therapy–responsive tumor microenvironments. Nat. Med. 24, 1178–1191.

19. Laskowski, T.J., Biederstädt, A., and Rezvani, K. (2022). Natural killer cells in antitumour adoptive cell immunotherapy. Nat. Rev. Cancer 22, 557–575.

20. Basar, R., Daher, M., and Rezvani, K. (2020). Next-generation cell therapies: the emerging role of CAR-NK cells. Hematology Am. Soc. Hematol. Educ. Program 2020, 570–578.

21. Luci, C., Bihl, F., Bourdely, P., Khou, S., Popa, A., Meghraoui-Kheddar, A., Vermeulen, O., Elaldi, R., Poissonnet, G., Sudaka, A., et al. (2021). Cutaneous Squamous Cell Carcinoma Development Is Associated with a Temporal Infiltration of ILC1 and NK Cells with Immune Dysfunctions. J. Invest. Dermatol. 141, 2369–2379.

22. Wu, S.-Y., Fu, T., Jiang, Y.-Z., and Shao, Z.-M. (2020). Natural killer cells in cancer biology and therapy. Mol. Cancer 19, 120.

23. Tang, F., Li, J., Qi, L., Liu, D., Bo, Y., Qin, S., Miao, Y., Yu, K., Hou, W., Li, J., et al. (2023). A pan-cancer single-cell panorama of human natural killer cells. Cell 0. 10.1016/j.cell.2023.07.034.

24. Böttcher, J.P., Bonavita, E., Chakravarty, P., Blees, H., Cabeza-Cabrerizo, M., Sammicheli, S., Rogers, N.C., Sahai, E., Zelenay, S., and Reis e Sousa, C. (2018). NK Cells Stimulate Recruitment of cDC1 into the Tumor Microenvironment Promoting Cancer Immune Control. Cell 172, 1022–1037.e14.

25. Zaiss, D.M.W., Gause, W.C., Osborne, L.C., and Artis, D. (2015). Emerging functions of amphiregulin in orchestrating immunity, inflammation, and tissue repair. Immunity 42, 216–226.

26. Wang, S., Zhang, Y., Wang, Y., Ye, P., Li, J., Li, H., Ding, Q., and Xia, J. (2016). Amphiregulin Confers Regulatory T Cell Suppressive Function and Tumor Invasion via the EGFR/GSK-3β/Foxp3 Axis. J. Biol. Chem. 291, 21085–21095.

27. Xu, Q., Long, Q., Zhu, D., Fu, D., Zhang, B., Han, L., Qian, M., Guo, J., Xu, J., Cao, L., et al. (2019). Targeting amphiregulin (AREG) derived from senescent stromal cells diminishes cancer resistance and averts programmed cell death 1 ligand (PD-L1)-mediated immunosuppression. Aging Cell 18, e13027.

28. Florentin, J., Zhao, J., Tai, Y.-Y., Sun, W., Ohayon, L.L., O’Neil, S.P., Arunkumar, A., Zhang, X., Zhu, J., Al Aaraj, Y., et al. (2022). Loss of Amphiregulin drives inflammation and endothelial apoptosis in pulmonary hypertension. Life Sci Alliance 5. 10.26508/lsa.202101264.

29. Lindzen, M., Ghosh, S., Noronha, A., Drago, D., Nataraj, N.B., Leitner, O., Carvalho, S., Zmora, E., Sapoznik, S., Shany, K.B., et al. (2021). Targeting autocrine amphiregulin robustly and reproducibly inhibits ovarian cancer in a syngeneic model: roles for wildtype p53. Oncogene 40, 3665–3679.

30. Hsieh, M.-J., Chen, Y.-H., Lee, I.-N., Huang, C., Ku, Y.-J., and Chen, J.-C. (2019). Secreted amphiregulin promotes vincristine resistance in oral squamous cell carcinoma. Int. J. Oncol. 55, 949–959.

31. Ishikawa, N., Daigo, Y., Takano, A., Taniwaki, M., Kato, T., Hayama, S., Murakami, H., Takeshima, Y., Inai, K., Nishimura, H., et al. (2005). Increases of amphiregulin and transforming growth factor-alpha in serum as predictors of poor response to gefitinib among patients with advanced non-small cell lung cancers. Cancer Res. 65, 9176–9184.

32. Wang, Y., Lifshitz, L., Silverstein, N.J., Mintzer, E., Luk, K., StLouis, P., Brehm, M.A., Wolfe, S.A., Deeks, S.G., and Luban, J. (2023). Transcriptional and chromatin profiling of human blood innate lymphoid cell subsets sheds light on HIV-1 pathogenesis. EMBO J., e114153.

33. Guerrero-Juarez, C.F., Lee, G.H., Liu, Y., Wang, S., Karikomi, M., Sha, Y., Chow, R.Y., Nguyen, T.T.L., Iglesias, V.S., Aasi, S., et al. (2022). Single-cell analysis of human basal cell carcinoma reveals novel regulators of tumor growth and the tumor microenvironment. Sci Adv 8, eabm7981.

34. Wang, Y., Lifshitz, L., Gellatly, K., Vinton, C.L., Busman-Sahay, K., McCauley, S., Vangala, P., Kim, K., Derr, A., Jaiswal, S., et al. (2020). HIV-1-induced cytokines deplete homeostatic innate lymphoid cells and expand TCF7-dependent memory NK cells. Nat. Immunol. 21,

35. Reynolds, G., Vegh, P., Fletcher, J., Poyner, E.F.M., Stephenson, E., Goh, I., Botting, R.A., Huang, N., Olabi, B., Dubois, A., et al. (2021). Developmental cell programs are co-opted in inflammatory skin disease. Science 371. 10.1126/science.aba6500.

36. Bal, S.M., Golebski, K., and Spits, H. (2020). Plasticity of innate lymphoid cell subsets. Nat. Rev. Immunol. 20, 552–565.

37. Feng, B., Bai, Z., Zhou, X., Zhao, Y., Xie, Y.-Q., Huang, X., Liu, Y., Enbar, T., Li, R., Wang, Y., et al. (2024). The type 2 cytokine Fc-IL-4 revitalizes exhausted CD8+ T cells against cancer. Nature 634, 712–720.

38. Fang, L., Ricketson, D., Getubig, L., and Darimont, B. (2006). Unliganded and hormone-bound glucocorticoid receptors interact with distinct hydrophobic sites in the Hsp90 C-terminal domain. Proc. Natl. Acad. Sci. U. S. A. 103, 18487–18492.

39. Kovacs, J.J., Murphy, P.J.M., Gaillard, S., Zhao, X., Wu, J.-T., Nicchitta, C.V., Yoshida, M., Toft, D.O., Pratt, W.B., and Yao, T.-P. (2005). HDAC6 regulates Hsp90 acetylation and chaperone-dependent activation of glucocorticoid receptor. Mol. Cell 18, 601–607.

40. Noddings, C.M., Johnson, J.L., and Agard, D.A. (2023). Cryo-EM reveals how Hsp90 and FKBP immunophilins co-regulate the glucocorticoid receptor. Nat. Struct. Mol. Biol. 30, 1867–1877.

41. Zhang, J., Li, H., Liu, Y., Zhao, K., Wei, S., Sugarman, E.T., Liu, L., and Zhang, G. (2022). Targeting HSP90 as a novel therapy for cancer: Mechanistic insights and translational relevance. Cells 11, 2778.

42. Lang, J.E., Forero-Torres, A., Yee, D., Yau, C., Wolf, D., Park, J., Parker, B.A., Chien, A.J., Wallace, A.M., Murthy, R., et al. (2022). Safety and efficacy of HSP90 inhibitor ganetespib for neoadjuvant treatment of stage II/III breast cancer. NPJ Breast Cancer 8, 128.

43. Agyeman, A.S., Jun, W.J., Proia, D.A., Kim, C.R., Skor, M.N., Kocherginsky, M., and Conzen, S.D. (2016). Hsp90 inhibition results in glucocorticoid receptor degradation in association with increased sensitivity to paclitaxel in triple-negative breast cancer. Horm. Cancer 7, 114–126.

44. Wang, L., Wang, L., Zhang, H., Lu, J., Zhang, Z., Wu, H., and Liang, Z. (2020). AREG mediates the epithelial-mesenchymal transition in pancreatic cancer cells via the EGFR/ERK/NF-κB signalling pathway. Oncol. Rep. 43, 1558–1568.

45. Collins, P.L., Cella, M., Porter, S.I., Li, S., Gurewitz, G.L., Hong, H.S., Johnson, R.P., Oltz, E.M., and Colonna, M. (2019). Gene Regulatory Programs Conferring Phenotypic Identities to Human NK Cells. Cell 176, 348–360.e12.

46. Stoll, S.W., Stuart, P.E., Lambert, S., Gandarillas, A., Rittié, L., Johnston, A., and Elder, J.T. (2016). Membrane-Tethered Intracellular Domain of Amphiregulin Promotes Keratinocyte Proliferation. J. Invest. Dermatol. 136, 444–452.

47. Zhao, X., Yang, W., Yu, T., Yu, Y., Cui, X., Zhou, Z., Yang, H., Yu, Y., Bilotta, A.J., Yao, S., et al. (2023). Th17 Cell-Derived Amphiregulin Promotes Colitis-Associated Intestinal Fibrosis Through Activation of mTOR and MEK in Intestinal Myofibroblasts. Gastroenterology 164,

48. Wang, J.-C., Derynck, M.K., Nonaka, D.F., Khodabakhsh, D.B., Haqq, C., and Yamamoto, K.R. (2004). Chromatin immunoprecipitation (ChIP) scanning identifies primary glucocorticoid receptor target genes. Proc. Natl. Acad. Sci. U. S. A. 101, 15603–15608.

49. Mostafa, M.M., Rider, C.F., Shah, S., Traves, S.L., Gordon, P.M.K., Miller-Larsson, A., Leigh, R., and Newton, R. (2019). Glucocorticoid-driven transcriptomes in human airway epithelial cells: commonalities, differences and functional insight from cell lines and primary cells. BMC Med. Genomics 12, 29.

50. Yang, H., Xia, L., Chen, J., Zhang, S., Martin, V., Li, Q., Lin, S., Chen, J., Calmette, J., Lu, M., et al. (2019). Stress–glucocorticoid–TSC22D3 axis compromises therapy-induced antitumor immunity. Nat. Med. 25, 1428–1441.

51. Obradović, M.M.S., Hamelin, B., Manevski, N., Couto, J.P., Sethi, A., Coissieux, M.-M., Münst, S., Okamoto, R., Kohler, H., Schmidt, A., et al. (2019). Glucocorticoids promote breast cancer metastasis. Nature 567, 540–544.

52. Uche, U.U., Piccirillo, A.R., Kataoka, S., Grebinoski, S.J., D’Cruz, L.M., and Kane, L.P. (2018). PIK3IP1/TrIP restricts activation of T cells through inhibition of PI3K/Akt. J. Exp. Med. 215, 3165–3179.

53. Chen, S., Bonifati, S., Qin, Z., St Gelais, C., Kodigepalli, K.M., Barrett, B.S., Kim, S.H., Antonucci, J.M., Ladner, K.J., Buzovetsky, O., et al. (2018). SAMHD1 suppresses innate immune responses to viral infections and inflammatory stimuli by inhibiting the NF-κB and interferon pathways. Proc. Natl. Acad. Sci. U. S. A. 115, E3798–E3807.

54. Ramasamy, S., Saez, B., Mukhopadhyay, S., Ding, D., Ahmed, A.M., Chen, X., Pucci, F., Yamin, R.’e, Wang, J., Pittet, M.J., et al. (2016). Tle1 tumor suppressor negatively regulates inflammation in vivo and modulates NF-κB inflammatory pathway. Proc. Natl. Acad. Sci. U. S. A. 113, 1871–1876.

55. Chodaparambil, J.V., Pate, K.T., Hepler, M.R.D., Tsai, B.P., Muthurajan, U.M., Luger, K., Waterman, M.L., and Weis, W.I. (2014). Molecular functions of the TLE tetramerization domain in Wnt target gene repression. EMBO J. 33, 719–731.

56. Kim, D.O., Byun, J.-E., Kim, W.S., Kim, M.J., Choi, J.H., Kim, H., Choi, E., Kim, T.-D., Yoon, S.R., Noh, J.-Y., et al. (2020). TXNIP Regulates Natural Killer Cell-Mediated Innate Immunity by Inhibiting IFN-γ Production during Bacterial Infection. Int. J. Mol. Sci. 21. 10.3390/ijms21249499.

57. Park, A., Lee, Y., Kim, M.S., Kang, Y.J., Park, Y.-J., Jung, H., Kim, T.-D., Lee, H.G., Choi, I., and Yoon, S.R. (2018). Prostaglandin E2 Secreted by Thyroid Cancer Cells Contributes to Immune Escape Through the Suppression of Natural Killer (NK) Cell Cytotoxicity and NK Cell Differentiation. Front. Immunol. 9, 1859.

58. Cook, M.E., Bradstreet, T.R., Webber, A.M., Kim, J., Santeford, A., Harris, K.M., Murphy, M.K., Tran, J., Abdalla, N.M., Schwarzkopf, E.A., et al. (2022). The ZFP36 family of RNA binding proteins regulates homeostatic and autoreactive T cell responses. Sci Immunol 7, eabo0981.

59. Bellone, S., Roque, D., Cocco, E., Gasparrini, S., Bortolomai, I., Buza, N., Abu-Khalaf, M., Silasi, D.-A., Ratner, E., Azodi, M., et al. (2012). Downregulation of membrane complement inhibitors CD55 and CD59 by siRNA sensitises uterine serous carcinoma overexpressing Her2/neu to complement and antibody-dependent cell cytotoxicity in vitro: implications for trastuzumab-based immunotherapy. Br. J. Cancer 106, 1543–1550.

60. Deng, Y., Kerdiles, Y., Chu, J., Yuan, S., Wang, Y., Chen, X., Mao, H., Zhang, L., Zhang, J., Hughes, T., et al. (2015). Transcription factor Foxo1 is a negative regulator of natural killer cell maturation and function. Immunity 42, 457–470.

61. Akman, B., Hu, X., Liu, X., Hatipoğlu, T., You, H., Chan, W.C., and Küçük, C. (2021). PRDM1 decreases sensitivity of human NK cells to IL2-induced cell expansion by directly repressing CD25 (IL2RA). J. Leukoc. Biol. 109, 901–914.

62. Smith, M.A., Maurin, M., Cho, H.I., Becknell, B., Freud, A.G., Yu, J., Wei, S., Djeu, J., Celis, E., Caligiuri, M.A., et al. (2010). PRDM1/Blimp-1 controls effector cytokine production in human NK cells. J. Immunol. 185, 6058–6067.

63. Franco, L.M., Gadkari, M., Howe, K.N., Sun, J., Kardava, L., Kumar, P., Kumari, S., Hu, Z., Fraser, I.D.C., Moir, S., et al. (2019). Immune regulation by glucocorticoids can be linked to cell type-dependent transcriptional responses. J. Exp. Med. 216, 384–406.

64. Bergeron, B.P., Barnett, K.R., Bhattarai, K.R., Mobley, R.J., Hansen, B.S., Brown, A., Kodali, K., High, A.A., Jeha, S., Pui, C.-H., et al. (2023). Mutual antagonism between glucocorticoid and canonical Wnt signaling pathways in B-cell acute lymphoblastic leukemia. Blood Adv. 10.1182/bloodadvances.2022009498.

65. Bayerl, F., Meiser, P., Donakonda, S., Hirschberger, A., Lacher, S.B., Pedde, A.-M., Hermann, C.D., Elewaut, A., Knolle, M., Ramsauer, L., et al. (2023). Tumor-derived prostaglandin E2 programs cDC1 dysfunction to impair intratumoral orchestration of anti-cancer T cell responses. Immunity 56, 1341–1358.e11.

66. Zelenay, S., van der Veen, A.G., Böttcher, J.P., Snelgrove, K.J., Rogers, N., Acton, S.E., Chakravarty, P., Girotti, M.R., Marais, R., Quezada, S.A., et al. (2015). Cyclooxygenase-Dependent Tumor Growth through Evasion of Immunity. Cell 162, 1257–1270.

67. Liu, S.-Y., Sanchez, D.J., Aliyari, R., Lu, S., and Cheng, G. (2012). Systematic identification of type I and type II interferon-induced antiviral factors. Proc. Natl. Acad. Sci. U. S. A. 109, 4239–4244.

68. Platanias, L.C. (2005). Mechanisms of type-I-and type-II-interferon-mediated signalling. Nat. Rev. Immunol. 5, 375–386.

69. Liu, W.M., and Zhang, X.A. (2006). KAI1/CD82, a tumor metastasis suppressor. Cancer Lett. 240, 183–194.

70. Jeong, S.-I., Kim, J.-W., Ko, K.-P., Ryu, B.-K., Lee, M.-G., Kim, H.-J., and Chi, S.-G. (2018). XAF1 forms a positive feedback loop with IRF-1 to drive apoptotic stress response and suppress tumorigenesis. Cell Death Dis. 9, 806.

71. Yang, R., Sun, L., Li, C.-F., Wang, Y.-H., Yao, J., Li, H., Yan, M., Chang, W.-C., Hsu, J.-M., Cha, J.-H., et al. (2021). Galectin-9 interacts with PD-1 and TIM-3 to regulate T cell death and is a target for cancer immunotherapy. Nat. Commun. 12, 832.

72. Du, X., de Almeida, P., Manieri, N., de Almeida Nagata, D., Wu, T.D., Harden Bowles, K., Arumugam, V., Schartner, J., Cubas, R., Mittman, S., et al. (2018). CD226 regulates natural killer cell antitumor responses via phosphorylation-mediated inactivation of transcription factor FOXO1. Proc. Natl. Acad. Sci. U. S. A. 115, E11731–E11740.

73. Schmidleithner, L., Thabet, Y., Schönfeld, E., Köhne, M., Sommer, D., Abdullah, Z., Sadlon, T., Osei-Sarpong, C., Subbaramaiah, K., Copperi, F., et al. (2019). Enzymatic Activity of HPGD in Treg Cells Suppresses Tconv Cells to Maintain Adipose Tissue Homeostasis and Prevent Metabolic Dysfunction. Immunity 50, 1232–1248.e14.

74. Li, H., Handsaker, B., Wysoker, A., Fennell, T., Ruan, J., Homer, N., Marth, G., Abecasis, G., Durbin, R., and 1000 Genome Project Data Processing Subgroup (2009). The Sequence Alignment/Map format and SAMtools. Bioinformatics 25, 2078–2079.

75. Love, M.I., Huber, W., and Anders, S. (2014). Moderated estimation of fold change and dispersion for RNA-seq data with DESeq2. Genome Biol. 15, 550.

76. Stuart, T., Butler, A., Hoffman, P., Hafemeister, C., Papalexi, E., Mauck, W.M., 3rd, Hao, Y., Stoeckius, M., Smibert, P., and Satija, R. (2019). Comprehensive Integration of Single-Cell Data. Cell 177, 1888–1902.e21.

77. Langmead, B., and Salzberg, S.L. (2012). Fast gapped-read alignment with Bowtie 2. Nat. Methods 9, 357–359.

78. Hashimshony, T., Senderovich, N., Avital, G., Klochendler, A., de Leeuw, Y., Anavy, L., Gennert, D., Li, S., Livak, K.J., Rozenblatt-Rosen, O., et al. (2016). CEL-Seq2: sensitive highly-multiplexed single-cell RNA-Seq. Genome Biol. 17, 77.

79. Zhang, Y., Liu, T., Meyer, C.A., Eeckhoute, J., Johnson, D.S., Bernstein, B.E., Nusbaum, C., Myers, R.M., Brown, M., Li, W., et al. (2008). Model-based analysis of ChIP-Seq (MACS). Genome Biol. 9, R137.

80. Bolger, A.M., Lohse, M., and Usadel, B. (2014). Trimmomatic: a flexible trimmer for Illumina sequence data. Bioinformatics 30, 2114–2120.

81. Robinson, J.T., Thorvaldsdóttir, H., Winckler, W., Guttman, M., Lander, E.S., Getz, G., and Mesirov, J.P. (2011). Integrative genomics viewer. Nat. Biotechnol. 29, 24–26.

