## Supplemental tables 1-6 for "Glucocorticoid receptor activation reprograms NK cells to drive AREG-mediated immunosuppression in skin cancer"

| Table S1 \| Skin cancer samples for scRNA-Seq and flow cytometry | | | | | |
|---|---|---|---|---|---|
| **Sample ID#** | **Sex** | **Age** | **Primary site** | **Experiment** | **Related Figure** |
| BCC 1 | Male | 41 | Face | scRNA-Seq, flow cytometry | Figure 1, 2, 3, 4, 5 |
| BCC 2 | Male | 71 | Retroauricular region | scRNA-Seq, flow cytometry | Figure 1, 2, 3, 4, 5 |
| BCC 3 | Female | 81 | Upper lip | scRNA-Seq, flow cytometry | Figure 1, 2, 3, 4, 5 |
| BCC 4 | Female | 82 | Ala of nose | flow cytometry | Figure 1, 4 |
| BCC 5 | Male | 82 | Root of nose | flow cytometry | Figure 1, 4 |
| BCC 6 | Female | 83 | Face | flow cytometry | Figure 1, 4 |
| BCC 7 | Female | 60 | Face | flow cytometry | Figure 1, 4 |
| BCC 8 | Female | 76 | Face | flow cytometry | Figure 1, 4 |
| BCC 9 | Male | 55 | Root of nose | flow cytometry | Figure 1, 4 |
| BCC 10 | Male | 69 | Posterior neck region | flow cytometry | Figure 1, 4 |
| BCC 11 | Female | 69 | Temporal region | flow cytometry | Figure 1, 4 |
| BCC 12 | Male | 59 | Axilla | flow cytometry | Figure 1, 4 |
| BCC 13 | Male | 68 | Lip | flow cytometry | Figure 1, 4 |
| cSCC 1 | Male | 86 | Vertex of head | scRNA-Seq, flow cytometry | Figure 1, 2, 3, 4, 5 |
| cSCC 2 | Male | 86 | Craniofacial region | scRNA-Seq, flow cytometry | Figure 1, 2, 3, 4, 5 |
| cSCC 3 | Female | 87 | Temporal region | scRNA-Seq, flow cytometry | Figure 1, 2, 3, 4, 5 |
| cSCC 4 | Female | 51 | Occipital region | flow cytometry | Figure 1, 4 |
| cSCC 5 | Male | 51 | Penis | flow cytometry | Figure 1, 4 |
| cSCC 6 | Male | 85 | Lower lip | flow cytometry | Figure 1, 4 |
| cSCC 7 | Male | 73 | Penis | flow cytometry | Figure 1, 4 |
| cSCC 8 | Male | 71 | Auricle | flow cytometry | Figure 1, 4 |
| cSCC 9 | Male | 71 | Face | flow cytometry | Figure 1, 4 |
| cSCC 10 | Female | 66 | Vulva | flow cytometry | Figure 1, 4 |
| cSCC 11 | Female | 83 | Nose | flow cytometry | Figure 1, 4 |
| cSCC 12 | Female | 86 | Upper eyelid | flow cytometry | Figure 1, 4 |
| cSCC 13 | Female | 67 | Upper arm | flow cytometry | Figure 1, 4 |
| cSCC 14 | Male | 87 | Vertex of head | flow cytometry | Figure 1, 4 |
| cSCC 15 | Male | 89 | Temporal region | flow cytometry | Figure 1, 4 |
| EMPD 1 | Male | 73 | Scrotum | scRNA-Seq , flow cytometry | Figure 1, 2, 3, 4, 5 |
| EMPD 2 | Female | 42 | Labium majus | scRNA-Seq | Figure 1, 2, 3, 5 |
| EMPD 3 | Male | 70 | Penis | scRNA-Seq | Figure 1, 2, 3, 5 |
| EMPD 4 | Female | 48 | Labium majus | flow cytometry | Figure 1, 4 |
| EMPD 5 | Male | 66 | Penis | flow cytometry | Figure 1, 4 |
| EMPD 6 | Female | 53 | Axilla | flow cytometry | Figure 1, 4 |
| aMM 1 | Female | 68 | Plantar region | scRNA-Seq, flow cytometry | Figure 1, 2, 3, 4, 5 |
| aMM 2 | Male | 81 | Hallux | scRNA-Seq | Figure 1, 2, 3, 5 |
| aMM 3 | Female | 74 | Toe | scRNA-Seq | Figure 1, 2, 3, 5 |
| aMM 4 | Male | 50 | Plantar region | flow cytometry | Figure 1, 4 |

| Table S2 \| Quality control for scRNA-Seq | | | |
|---|---|---|---|
| **Sample** | **Estimated Number of Cells** | **Fraction Reads in Cells** | **Mean Reads per Cell** |
| BCC1_Tumor | 6,048 | 83.10% | 77,529 |
| BCC2_Tumor | 7,561 | 91.30% | 48,144 |
| BCC3_Tumor | 14,042 | 89.20% | 32,057 |
| BCC1_Peri-tumor | 7,778 | 89.50% | 57,147 |
| BCC2_Peri-tumor | 6,793 | 93.00% | 53,915 |
| BCC3_Peri-tumor | 9,240 | 83.90% | 56,502 |
| Empd1_Tumor | 9,510 | 92.90% | 37,954 |
| Empd2_Tumor | 9,234 | 90.90% | 47,750 |
| Empd3_Tumor | 6,338 | 80.50% | 58,405 |
| Empd1_Peri-tumor | 10,055 | 95.50% | 43,816 |
| Empd2_Peri-tumor | 6,421 | 93.90% | 72,505 |
| Empd3_Peri-tumor | 10,311 | 84.90% | 39,961 |
| aMM1_Tumor | 10,538 | 93.60% | 38,782 |
| aMM2_Tumor | 10,100 | 94.20% | 36,491 |
| aMM3_Tumor | 8,237 | 86.30% | 61,182 |
| aMM1_Peri-tumor | 14,367 | 84.50% | 26,931 |
| aMM2_Peri-tumor | 10,906 | 92.10% | 42,746 |
| aMM3_Peri-tumor | 11,242 | 87.20% | 37,815 |
| cSCC1_Tumor | 7,542 | 90.20% | 56,883 |
| cSCC2_Tumor | 7,163 | 85.10% | 51,865 |
| cSCC3_Tumor | 8,007 | 90.30% | 46,905 |
| cSCC1_Peri-tumor | 10,643 | 92.40% | 38,657 |
| cSCC2_Peri-tumor | 7,472 | 85.40% | 57,480 |
| cSCC3_Peri-tumor | 7,489 | 88.80% | 54,668 |

| **Sample** | **Median Genes per Cell** | **Total Genes Detected** | **Median UMI Counts per Cell** |
|---|---|---|---|
| BCC1_Tumor | 1,057 | 24,822 | 2,643 |
| BCC2_Tumor | 1,364 | 24,078 | 3,781 |
| BCC3_Tumor | 1,098 | 24,904 | 2,996 |
| BCC1_Peri-tumor | 1,572 | 28,182 | 4,338 |
| BCC2_Peri-tumor | 1,266 | 22,840 | 3,841 |
| BCC3_Peri-tumor | 1,389 | 24,876 | 3,447 |
| Empd1_Tumor | 1,123 | 26,550 | 2,855 |
| Empd2_Tumor | 1,254 | 25,411 | 3,241 |
| Empd3_Tumor | 1,227 | 26,045 | 3,159 |
| Empd1_Peri-tumor | 1,612 | 27,531 | 4,546 |
| Empd2_Peri-tumor | 1,176 | 25,222 | 2,887 |
| Empd3_Peri-tumor | 1,093 | 27,491 | 2,613 |
| aMM1_Tumor | 1,303 | 25,560 | 3,623 |
| aMM2_Tumor | 1,288 | 25,834 | 3,963 |
| aMM3_Tumor | 1,122 | 27,273 | 3,811 |
| aMM1_Peri-tumor | 1,120 | 27,043 | 2,407 |
| aMM2_Peri-tumor | 1,370 | 28,223 | 3,882 |
| aMM3_Peri-tumor | 1,072 | 29,071 | 2,882 |
| cSCC1_Tumor | 1,085 | 23,422 | 3,109 |
| cSCC2_Tumor | 1,032 | 24,861 | 2,316 |
| cSCC3_Tumor | 1,526 | 24,816 | 4,647 |
| cSCC1_Peri-tumor | 1,250 | 27,538 | 3,551 |
| cSCC2_Peri-tumor | 1,056 | 24,628 | 2,979 |

| | | | |
|---|---|---|---|
| cSCC3_Peri-tumor | 1,372 | 25,074 | 4,437 |

| Sample | Reads Mapped Confidently to Genome | Reads Mapped Confidently to Intergenic Regions | Reads Mapped Confidently to Intronic Regions |
|---|---|---|---|
| BCC1_Tumor | 91.40% | 4.50% | 47.50% |
| BCC2_Tumor | 90.90% | 6.10% | 16.70% |
| BCC3_Tumor | 92.10% | 5.50% | 23.20% |
| BCC1_Peri-tumor | 90.40% | 6.10% | 40.30% |
| BCC2_Peri-tumor | 91.70% | 6.80% | 16.00% |
| BCC3_Peri-tumor | 91.10% | 5.50% | 17.60% |
| Empd1_Tumor | 91.80% | 5.20% | 22.00% |
| Empd2_Tumor | 91.90% | 4.30% | 23.10% |
| Empd3_Tumor | 90.90% | 5.30% | 32.70% |
| Empd1_Peri-tumor | 92.00% | 4.00% | 18.20% |
| Empd2_Peri-tumor | 92.60% | 5.00% | 20.80% |
| Empd3_Peri-tumor | 92.60% | 4.50% | 31.10% |
| aMM1_Tumor | 93.30% | 3.90% | 24.40% |
| aMM2_Tumor | 90.00% | 5.20% | 28.00% |
| aMM3_Tumor | 92.70% | 4.70% | 36.60% |
| aMM1_Peri-tumor | 90.50% | 4.30% | 15.80% |
| aMM2_Peri-tumor | 88.90% | 5.60% | 21.40% |
| aMM3_Peri-tumor | 88.30% | 6.80% | 40.00% |
| cSCC1_Tumor | 91.80% | 6.00% | 23.80% |
| cSCC2_Tumor | 90.80% | 6.60% | 25.80% |
| cSCC3_Tumor | 93.70% | 3.30% | 32.80% |
| cSCC1_Peri-tumor | 93.10% | 5.30% | 14.80% |
| cSCC2_Peri-tumor | 92.80% | 5.00% | 17.20% |
| cSCC3_Peri-tumor | 93.10% | 3.80% | 34.30% |

| Sample | Reads Mapped Confidently to Exonic Regions | Reads Mapped Confidently to Transcriptome |
|---|---|---|
| BCC1_Tumor | 39.40% | 33.40% |
| BCC2_Tumor | 68.10% | 63.00% |
| BCC3_Tumor | 63.40% | 58.30% |
| BCC1_Peri-tumor | 44.10% | 37.60% |
| BCC2_Peri-tumor | 68.90% | 63.80% |
| BCC3_Peri-tumor | 68.00% | 62.60% |
| Empd1_Tumor | 64.70% | 58.70% |
| Empd2_Tumor | 64.50% | 58.90% |
| Empd3_Tumor | 52.90% | 47.70% |
| Empd1_Peri-tumor | 69.80% | 63.70% |
| Empd2_Peri-tumor | 66.70% | 61.20% |
| Empd3_Peri-tumor | 57.00% | 50.40% |
| aMM1_Tumor | 65.00% | 59.00% |
| aMM2_Tumor | 56.70% | 50.50% |
| aMM3_Tumor | 51.50% | 45.30% |
| aMM1_Peri-tumor | 70.40% | 64.40% |
| aMM2_Peri-tumor | 61.90% | 55.90% |
| aMM3_Peri-tumor | 41.50% | 36.00% |
| cSCC1_Tumor | 61.90% | 56.00% |
| cSCC2_Tumor | 58.40% | 52.70% |

| | | |
|---|---|---|
| cSCC3_Tumor | 57.60% | 52.20% |
| cSCC1_Peri-tumor | 73.10% | 67.00% |
| cSCC2_Peri-tumor | 70.60% | 65.30% |
| cSCC3_Peri-tumor | 55.00% | 49.40% |

| Table S3 \| Skin tumor NK cells highly expressed genes | | | | | | | | |
|---|---|---|---|---|---|---|---|---|
| GeneName | Peri-tumor | Tumor | cells.1 | cells.2 | pct.1 | pct.2 | log2FC | p_value |
| GZMA | 2.37394802 | 6.88231292 | 549 | 1733 | 0.335 | 0.49 | 1.53560514 | 3.33E-14 |
| DUSP1 | 3.8074024 | 10.2655046 | 549 | 1733 | 0.485 | 0.66 | 1.43092559 | 4.23E-18 |
| GZMK | 1.01937565 | 2.61254397 | 549 | 1733 | 0.193 | 0.331 | 1.35776953 | 6.13E-12 |
| CCL3 | 3.83741468 | 8.89263658 | 549 | 1733 | 0.259 | 0.373 | 1.21247655 | 1.16E-07 |
| CCL4L2 | 3.5935138 | 8.02218912 | 549 | 1733 | 0.188 | 0.29 | 1.15860075 | 1.80E-06 |
| NR4A1 | 1.96685609 | 4.2034709 | 549 | 1733 | 0.268 | 0.398 | 1.09568969 | 8.44E-10 |
| DNAJB1 | 13.7464422 | 28.3640824 | 549 | 1733 | 0.514 | 0.698 | 1.04500692 | 2.52E-16 |
| RGS1 | 1.82462571 | 3.52305124 | 549 | 1733 | 0.237 | 0.403 | 0.94922491 | 2.40E-13 |
| CCL4 | 13.5545009 | 26.1518622 | 549 | 1733 | 0.464 | 0.579 | 0.94814169 | 5.91E-07 |
| IFNG | 4.18527632 | 7.96759147 | 549 | 1733 | 0.293 | 0.413 | 0.9288208 | 1.81E-06 |
| MIR23AHG | 1.99385372 | 3.76334104 | 549 | 1733 | 0.361 | 0.443 | 0.91645446 | 1.52E-05 |
| EMB | 0.53596628 | 1.00315018 | 549 | 1733 | 0.164 | 0.293 | 0.90432347 | 3.40E-09 |
| CTSD | 0.71422889 | 1.33224909 | 549 | 1733 | 0.209 | 0.327 | 0.89940546 | 6.08E-08 |
| AREG | 3.03869477 | 5.64697285 | 549 | 1733 | 0.313 | 0.419 | 0.89402593 | 2.31E-07 |
| EVL | 0.88822243 | 1.64175028 | 549 | 1733 | 0.22 | 0.367 | 0.88624179 | 3.53E-10 |
| PPP1R15A | 5.50945361 | 9.38606215 | 549 | 1733 | 0.579 | 0.697 | 0.76861076 | 5.25E-08 |
| DNAJB4 | 0.80546541 | 1.3458864 | 549 | 1733 | 0.175 | 0.253 | 0.74066212 | 4.91E-04 |
| DDX3Y | 1.68447835 | 2.81270006 | 549 | 1733 | 0.259 | 0.463 | 0.73965383 | 2.55E-17 |
| ACTB | 7.60056178 | 12.5133739 | 549 | 1733 | 0.78 | 0.88 | 0.71929287 | 7.85E-13 |
| ERN1 | 1.07070921 | 1.74744579 | 549 | 1733 | 0.226 | 0.321 | 0.70668098 | 6.47E-05 |
| TSC22D3 | 4.89605992 | 7.98639562 | 549 | 1733 | 0.523 | 0.719 | 0.70592332 | 5.66E-18 |
| SH3BGRL | 0.5831246 | 0.94333689 | 549 | 1733 | 0.166 | 0.279 | 0.69396889 | 1.82E-07 |
| RNF213 | 2.30794093 | 3.69097808 | 549 | 1733 | 0.45 | 0.589 | 0.67739687 | 4.49E-09 |
| LY6E | 0.7961081 | 1.26524577 | 549 | 1733 | 0.222 | 0.329 | 0.66838141 | 7.02E-06 |
| JUN | 6.18872063 | 9.78707524 | 549 | 1733 | 0.448 | 0.588 | 0.66123659 | 5.96E-09 |
| COTL1 | 1.62636851 | 2.57133882 | 549 | 1733 | 0.299 | 0.441 | 0.66086553 | 1.07E-08 |
| ALOX5AP | 1.04885935 | 1.65103454 | 549 | 1733 | 0.208 | 0.319 | 0.65454908 | 1.63E-06 |
| HAVCR2 | 1.02130445 | 1.6023851 | 549 | 1733 | 0.211 | 0.342 | 0.64980792 | 2.58E-08 |
| BCAS2 | 1.07468549 | 1.67998833 | 549 | 1733 | 0.279 | 0.37 | 0.6445367 | 2.80E-04 |
| FMNL1-DT | 0.91720939 | 1.39065038 | 549 | 1733 | 0.239 | 0.351 | 0.60043674 | 3.35E-06 |
| HLA-F | 1.49440673 | 2.26191423 | 549 | 1733 | 0.352 | 0.509 | 0.59797137 | 6.71E-10 |
| GBP5 | 1.03536311 | 1.55028312 | 549 | 1733 | 0.222 | 0.313 | 0.58239489 | 1.36E-04 |
| DDX27 | 0.68677486 | 1.01293698 | 549 | 1733 | 0.195 | 0.282 | 0.56063528 | 2.03E-04 |
| KLF6 | 6.35710117 | 9.34269225 | 549 | 1733 | 0.689 | 0.761 | 0.5554693 | 5.62E-08 |
| FOS | 8.23548385 | 12.0921447 | 549 | 1733 | 0.424 | 0.569 | 0.55414483 | 3.35E-09 |
| NFKBIA | 11.1144311 | 16.2850364 | 549 | 1733 | 0.778 | 0.842 | 0.55111284 | 2.18E-08 |
| USP16 | 0.61090606 | 0.89489702 | 549 | 1733 | 0.175 | 0.263 | 0.55077113 | 1.00E-04 |
| NFKBID | 0.98713586 | 1.44179499 | 549 | 1733 | 0.244 | 0.301 | 0.54654549 | 0.00472093 |
| IFITM1 | 3.10930492 | 4.45595974 | 549 | 1733 | 0.534 | 0.666 | 0.5191441 | 2.02E-08 |
| CALM3 | 0.68578978 | 0.98242984 | 549 | 1733 | 0.209 | 0.285 | 0.51858797 | 0.00169549 |
| MATK | 1.36638421 | 1.95037941 | 549 | 1733 | 0.341 | 0.462 | 0.51339159 | 2.27E-06 |
| ISG20 | 3.11961645 | 4.44981327 | 549 | 1733 | 0.505 | 0.604 | 0.51237613 | 1.26E-05 |
| KLHL6 | 0.75105806 | 1.06962538 | 549 | 1733 | 0.186 | 0.254 | 0.51010926 | 0.00328905 |
| CEBPD | 1.44636305 | 2.0502869 | 549 | 1733 | 0.255 | 0.312 | 0.50339608 | 0.01400605 |
| IL4R | 0.58591764 | 0.82843268 | 549 | 1733 | 0.166 | 0.253 | 0.49968659 | 3.14E-05 |
| HCST | 4.21131853 | 5.91227775 | 549 | 1733 | 0.627 | 0.754 | 0.48944205 | 1.26E-10 |
| VPS28 | 0.61534058 | 0.86328243 | 549 | 1733 | 0.173 | 0.256 | 0.48844748 | 1.81E-04 |
| PLP2 | 0.85956896 | 1.20119762 | 549 | 1733 | 0.246 | 0.336 | 0.48278822 | 3.11E-04 |
| LAT2 | 0.92003739 | 1.28505909 | 549 | 1733 | 0.237 | 0.327 | 0.48207031 | 2.74E-04 |
| GZMB | 10.8556317 | 15.1440735 | 549 | 1733 | 0.628 | 0.705 | 0.48030964 | 0.00373448 |
| TXK | 0.84817716 | 1.17967304 | 549 | 1733 | 0.208 | 0.278 | 0.47594952 | 0.00356026 |
| PARP8 | 1.18306293 | 1.64263195 | 549 | 1733 | 0.304 | 0.402 | 0.47348245 | 1.60E-04 |
| BIN2 | 0.93188136 | 1.29170766 | 549 | 1733 | 0.233 | 0.338 | 0.47106141 | 1.18E-05 |
| SYAP1 | 2.13292361 | 2.95164808 | 549 | 1733 | 0.426 | 0.526 | 0.46868843 | 2.87E-05 |
| CD69 | 9.38268978 | 12.9804097 | 549 | 1733 | 0.701 | 0.818 | 0.46826245 | 7.33E-09 |
| HSPA1A | 63.2868414 | 87.4251219 | 549 | 1733 | 0.763 | 0.85 | 0.46614234 | 1.03E-07 |
| CARD16 | 0.81044413 | 1.116349 | 549 | 1733 | 0.193 | 0.278 | 0.46200348 | 2.78E-04 |

| | | | | | | | | |
|---|---|---|---|---|---|---|---|---|
| TYROBP | 4.29654766 | 5.9070877 | 549 | 1733 | 0.628 | 0.784 | 0.45926914 | 8.56E-13 |
| BBLN | 0.98250373 | 1.34990977 | 549 | 1733 | 0.279 | 0.368 | 0.45832818 | 5.35E-04 |
| HSP90AA1 | 101.880215 | 139.762354 | 549 | 1733 | 0.913 | 0.977 | 0.4561019 | 7.58E-17 |
| FCER1G | 2.58251206 | 3.52933141 | 549 | 1733 | 0.399 | 0.548 | 0.45061983 | 7.76E-09 |
| IRF8 | 1.38398139 | 1.8856344 | 549 | 1733 | 0.253 | 0.373 | 0.44622544 | 5.55E-07 |
| KDM5B | 1.14582969 | 1.5592968 | 549 | 1733 | 0.288 | 0.362 | 0.44450293 | 0.00361016 |
| FOXN3 | 0.6008135 | 0.81695791 | 549 | 1733 | 0.175 | 0.253 | 0.44334452 | 3.97E-04 |
| NOP58 | 1.16878634 | 1.58709227 | 549 | 1733 | 0.301 | 0.394 | 0.44137478 | 3.33E-04 |
| ADGRE5 | 3.96858555 | 5.35940923 | 549 | 1733 | 0.648 | 0.735 | 0.43344907 | 2.81E-08 |
| SIAH2 | 0.64223152 | 0.86718606 | 549 | 1733 | 0.193 | 0.253 | 0.43324809 | 0.01438455 |
| BST2 | 1.2105129 | 1.62109303 | 549 | 1733 | 0.313 | 0.39 | 0.42134843 | 0.0024844 |
| RIN3 | 0.87679332 | 1.17276153 | 549 | 1733 | 0.246 | 0.329 | 0.41960097 | 7.68E-04 |
| NSG0000028288 | 0.80562622 | 1.07737187 | 549 | 1733 | 0.197 | 0.299 | 0.41933377 | 1.30E-06 |
| NAMPT | 3.01505579 | 4.02803994 | 549 | 1733 | 0.468 | 0.565 | 0.41789329 | 3.12E-04 |
| CORO1A | 2.27493389 | 3.038773 | 549 | 1733 | 0.486 | 0.593 | 0.41766429 | 2.62E-05 |
| ANAPC16 | 0.88542398 | 1.18167387 | 549 | 1733 | 0.235 | 0.317 | 0.41639156 | 9.83E-04 |
| STARD3NL | 0.69276715 | 0.92400614 | 549 | 1733 | 0.208 | 0.278 | 0.41553192 | 0.00374946 |
| IFITM3 | 1.56001422 | 2.08043497 | 549 | 1733 | 0.319 | 0.412 | 0.41532601 | 4.18E-04 |
| CRYBG1 | 1.02592335 | 1.36814847 | 549 | 1733 | 0.25 | 0.337 | 0.41530186 | 3.85E-04 |
| PAXX | 1.26820989 | 1.69079586 | 549 | 1733 | 0.333 | 0.396 | 0.41490895 | 0.01311021 |
| UBC | 19.9553414 | 26.6001303 | 549 | 1733 | 0.909 | 0.947 | 0.41465835 | 7.13E-06 |
| CRIP1 | 2.71517672 | 3.61505445 | 549 | 1733 | 0.395 | 0.555 | 0.41297128 | 2.65E-10 |
| ZBTB16 | 1.13875942 | 1.51577254 | 549 | 1733 | 0.211 | 0.295 | 0.41259029 | 3.71E-04 |
| NSG0000027225 | 1.0315028 | 1.3678913 | 549 | 1733 | 0.22 | 0.317 | 0.40720586 | 5.72E-05 |
| COX20 | 0.67885821 | 0.89999305 | 549 | 1733 | 0.208 | 0.269 | 0.40680358 | 0.0131369 |
| MYADM | 0.97535994 | 1.29180191 | 549 | 1733 | 0.224 | 0.317 | 0.40537823 | 7.97E-05 |
| HSPE1 | 16.7095421 | 22.1004467 | 549 | 1733 | 0.783 | 0.869 | 0.40340333 | 3.46E-06 |
| HSPA1B | 18.4223471 | 24.3565565 | 549 | 1733 | 0.475 | 0.594 | 0.4028533 | 2.08E-07 |
| RESF1 | 1.43188377 | 1.88964449 | 549 | 1733 | 0.344 | 0.398 | 0.40020045 | 0.01947754 |
| RPS4Y1 | 2.14446295 | 2.82930824 | 549 | 1733 | 0.319 | 0.525 | 0.39983297 | 3.57E-21 |
| CASP4 | 0.7147167 | 0.93910683 | 549 | 1733 | 0.222 | 0.291 | 0.39391778 | 0.00616202 |
| SLA2 | 0.98985915 | 1.29620705 | 549 | 1733 | 0.242 | 0.307 | 0.38900102 | 0.00834108 |
| VPS13C | 1.0654196 | 1.39434674 | 549 | 1733 | 0.259 | 0.345 | 0.38816764 | 3.63E-04 |
| FAM177A1 | 3.78798343 | 4.95004557 | 549 | 1733 | 0.503 | 0.63 | 0.38601179 | 9.21E-07 |
| LSP1 | 1.20102258 | 1.56916636 | 549 | 1733 | 0.286 | 0.388 | 0.38573504 | 5.32E-05 |
| TLN1 | 1.71558508 | 2.24070114 | 549 | 1733 | 0.393 | 0.511 | 0.38524956 | 9.52E-06 |
| CTSW | 3.34551612 | 4.36802389 | 549 | 1733 | 0.532 | 0.653 | 0.38475195 | 2.17E-06 |
| PDE4A | 1.44154898 | 1.88204786 | 549 | 1733 | 0.308 | 0.384 | 0.38468345 | 0.00344216 |
| JUNB | 8.33735788 | 10.8800854 | 549 | 1733 | 0.705 | 0.776 | 0.38402771 | 0.00157884 |
| GMFG | 1.00156708 | 1.30644483 | 549 | 1733 | 0.268 | 0.348 | 0.38338715 | 0.00196988 |
| GRK2 | 1.00985569 | 1.31713843 | 549 | 1733 | 0.27 | 0.362 | 0.38325783 | 1.53E-04 |
| ATP6V1F | 0.71384692 | 0.93041211 | 549 | 1733 | 0.224 | 0.297 | 0.38225514 | 0.0034022 |
| GZMM | 1.85344923 | 2.41548909 | 549 | 1733 | 0.353 | 0.481 | 0.38210274 | 6.87E-07 |
| IL2RB | 2.63601462 | 3.43064765 | 549 | 1733 | 0.483 | 0.609 | 0.38012259 | 1.31E-06 |
| ANKRD44 | 0.74785743 | 0.97085586 | 549 | 1733 | 0.191 | 0.279 | 0.37649385 | 1.93E-05 |
| ARID1B | 0.94910964 | 1.23144813 | 549 | 1733 | 0.246 | 0.323 | 0.3757092 | 0.00253769 |
| APOL6 | 1.00384767 | 1.30136036 | 549 | 1733 | 0.231 | 0.336 | 0.37448015 | 3.37E-06 |
| HSPH1 | 13.3515375 | 17.2114447 | 549 | 1733 | 0.627 | 0.752 | 0.36636232 | 1.50E-07 |

| Table S4 \| Highly expressed genes for each NK cell cluster | | | | | | | |
|---|---|---|---|---|---|---|---|
| Target Cluster | Gene Name | Target Cluster pct | Other Cluster pct | Target Cluster mean | Other Cluster mean | Log2FC | Pvalue |
| 0 | CD44 | 0.95 | 0.69 | 15.97 | 5.59 | 1.51 | 5.86E-161 |
| 0 | XCL1 | 0.89 | 0.40 | 32.55 | 11.26 | 1.53 | 1.45E-138 |
| 0 | IL7R | 0.70 | 0.28 | 15.95 | 2.82 | 2.50 | 2.71E-116 |
| 0 | GPR183 | 0.71 | 0.27 | 7.54 | 1.83 | 2.04 | 1.12E-111 |
| 0 | REL | 0.94 | 0.77 | 25.38 | 10.33 | 1.30 | 9.26E-110 |
| 0 | TNFRSF18 | 0.82 | 0.38 | 6.92 | 2.27 | 1.61 | 1.76E-109 |
| 0 | TCF7 | 0.68 | 0.25 | 4.54 | 1.15 | 1.98 | 7.10E-105 |
| 0 | XCL2 | 0.92 | 0.58 | 40.98 | 17.18 | 1.25 | 2.59E-98 |
| 0 | NFKB1 | 0.85 | 0.51 | 9.71 | 3.62 | 1.42 | 9.71E-96 |
| 0 | JAK1 | 0.93 | 0.75 | 12.87 | 6.04 | 1.09 | 2.77E-90 |
| 0 | BTG1 | 0.99 | 0.96 | 54.60 | 30.51 | 0.84 | 8.07E-86 |
| 0 | KLRC1 | 0.75 | 0.34 | 8.50 | 3.18 | 1.42 | 1.06E-81 |
| 0 | TPT1 | 0.99 | 0.99 | 71.66 | 48.84 | 0.55 | 3.94E-80 |
| 0 | AHI1 | 0.67 | 0.34 | 7.23 | 1.84 | 1.97 | 1.82E-75 |
| 0 | PABPC1 | 0.96 | 0.87 | 16.69 | 9.14 | 0.87 | 3.49E-72 |
| 0 | ITM2C | 0.51 | 0.17 | 2.65 | 0.70 | 1.92 | 1.94E-70 |
| 0 | SRGN | 0.97 | 0.96 | 54.53 | 28.77 | 0.92 | 2.42E-69 |
| 0 | VIM | 0.95 | 0.87 | 35.28 | 16.11 | 1.13 | 2.31E-68 |
| 0 | SPTBN1 | 0.62 | 0.31 | 4.15 | 1.31 | 1.66 | 1.98E-66 |
| 0 | IL2RA | 0.29 | 0.04 | 1.11 | 0.12 | 3.19 | 6.72E-66 |
| 0 | PDE4A | 0.56 | 0.24 | 3.03 | 0.95 | 1.68 | 1.40E-61 |
| 0 | NFE2L2 | 0.80 | 0.53 | 6.48 | 3.08 | 1.07 | 8.51E-60 |
| 0 | SNHG16 | 0.59 | 0.29 | 3.36 | 1.16 | 1.54 | 2.58E-58 |
| 0 | AGO2 | 0.60 | 0.29 | 3.20 | 1.14 | 1.49 | 2.98E-57 |
| 0 | BACH2 | 0.32 | 0.07 | 1.29 | 0.21 | 2.63 | 1.76E-55 |
| 0 | STAM | 0.50 | 0.20 | 2.03 | 0.72 | 1.49 | 6.50E-54 |
| 0 | TIAM1 | 0.30 | 0.07 | 1.22 | 0.22 | 2.45 | 3.81E-52 |
| 0 | IFITM3 | 0.57 | 0.27 | 3.17 | 1.15 | 1.46 | 1.64E-50 |
| 0 | IL12RB2 | 0.31 | 0.08 | 1.28 | 0.24 | 2.40 | 3.16E-50 |
| 0 | SMAP2 | 0.81 | 0.56 | 6.87 | 3.62 | 0.92 | 5.26E-50 |
| 0 | CD55 | 0.69 | 0.41 | 4.91 | 2.21 | 1.15 | 5.67E-50 |
| 0 | FCER1G | 0.70 | 0.39 | 4.71 | 2.37 | 0.99 | 6.70E-49 |
| 0 | LMNA | 0.79 | 0.50 | 8.11 | 4.82 | 0.75 | 4.57E-48 |
| 0 | CRTAM | 0.58 | 0.30 | 7.19 | 2.68 | 1.43 | 7.22E-48 |
| 0 | FES | 0.29 | 0.07 | 1.24 | 0.24 | 2.37 | 1.82E-47 |
| 0 | MYC | 0.32 | 0.09 | 1.76 | 0.51 | 1.79 | 8.66E-47 |
| 0 | NINJ1 | 0.56 | 0.27 | 2.72 | 1.12 | 1.28 | 3.78E-46 |
| 0 | SKP1 | 0.84 | 0.67 | 6.99 | 3.97 | 0.81 | 4.97E-46 |
| 0 | FURIN | 0.41 | 0.15 | 1.52 | 0.51 | 1.58 | 1.29E-44 |
| 0 | KDM6B | 0.73 | 0.48 | 5.72 | 2.89 | 0.98 | 8.19E-42 |
| 0 | METRNL | 0.86 | 0.64 | 10.44 | 6.07 | 0.78 | 1.03E-41 |
| 0 | ECE1 | 0.32 | 0.10 | 1.16 | 0.29 | 1.98 | 1.50E-41 |
| 0 | FMNL1-DT | 0.48 | 0.22 | 2.00 | 0.80 | 1.33 | 3.60E-41 |
| 0 | SELL | 0.31 | 0.09 | 1.30 | 0.33 | 1.96 | 5.32E-41 |
| 0 | RPLP1 | 0.99 | 0.99 | 70.09 | 54.23 | 0.37 | 4.09E-40 |
| 0 | SATB1 | 0.50 | 0.25 | 2.67 | 0.95 | 1.50 | 1.16E-39 |
| 0 | CXXC5 | 0.42 | 0.17 | 1.64 | 0.63 | 1.38 | 1.65E-39 |
| 0 | ABHD2 | 0.48 | 0.23 | 2.09 | 0.82 | 1.35 | 2.42E-39 |
| 0 | FAM177A1 | 0.73 | 0.51 | 6.84 | 3.24 | 1.08 | 3.08E-39 |
| 0 | BIRC3 | 0.77 | 0.55 | 15.92 | 7.50 | 1.09 | 3.27E-38 |
| 0 | GOLIM4 | 0.34 | 0.12 | 1.35 | 0.40 | 1.75 | 1.23E-37 |
| 0 | RPS24 | 0.99 | 0.98 | 40.59 | 31.47 | 0.37 | 1.26E-37 |
| 0 | PGK1 | 0.72 | 0.49 | 4.88 | 2.61 | 0.90 | 1.86E-37 |
| 0 | FOSL2 | 0.76 | 0.54 | 5.50 | 3.02 | 0.86 | 2.37E-37 |

| | | | | | | | |
|---|---|---|---|---|---|---|---|
| 0 | ANXA11 | 0.53 | 0.28 | 2.32 | 0.99 | 1.22 | 2.70E-37 |
| 0 | SERBP1 | 0.84 | 0.65 | 6.33 | 3.92 | 0.69 | 2.12E-36 |
| 0 | NFKB2 | 0.59 | 0.35 | 2.97 | 1.45 | 1.04 | 2.01E-35 |
| 0 | RPS5 | 0.95 | 0.91 | 15.15 | 10.98 | 0.46 | 2.57E-35 |
| 0 | LEPROTL1 | 0.68 | 0.45 | 4.09 | 2.05 | 1.00 | 2.92E-35 |
| 0 | TMEM123 | 0.51 | 0.27 | 2.37 | 1.01 | 1.23 | 3.43E-35 |
| 0 | TGFB1 | 0.84 | 0.69 | 8.31 | 4.91 | 0.76 | 4.46E-35 |
| 0 | RAMP1 | 0.27 | 0.08 | 1.20 | 0.34 | 1.81 | 4.79E-35 |
| 0 | PDE4B | 0.63 | 0.36 | 3.40 | 1.91 | 0.83 | 1.17E-34 |
| 0 | PAK1 | 0.26 | 0.08 | 0.79 | 0.22 | 1.86 | 2.38E-34 |
| 0 | SPAG1 | 0.29 | 0.10 | 0.97 | 0.29 | 1.75 | 5.08E-34 |
| 0 | RPL9 | 0.98 | 0.96 | 22.33 | 17.02 | 0.39 | 1.18E-33 |
| 0 | LPXN | 0.51 | 0.29 | 2.49 | 1.04 | 1.26 | 3.80E-33 |
| 0 | TAMALIN | 0.44 | 0.20 | 2.15 | 1.00 | 1.10 | 4.67E-33 |
| 0 | WSB1 | 0.69 | 0.50 | 5.62 | 2.64 | 1.09 | 5.98E-33 |
| 0 | KLHL6 | 0.37 | 0.15 | 1.49 | 0.66 | 1.17 | 6.73E-33 |
| 0 | ZFPM1 | 0.29 | 0.10 | 1.04 | 0.30 | 1.78 | 7.47E-33 |
| 0 | IRAG2 | 0.50 | 0.26 | 2.19 | 1.03 | 1.09 | 1.04E-32 |
| 0 | AREG | 0.55 | 0.29 | 6.84 | 3.82 | 0.84 | 1.44E-32 |
| 0 | ERGIC1 | 0.36 | 0.15 | 1.30 | 0.49 | 1.40 | 9.54E-32 |
| 0 | PPP1CB | 0.71 | 0.49 | 4.04 | 2.32 | 0.80 | 1.53E-31 |
| 0 | ATP1B1 | 0.42 | 0.19 | 2.26 | 0.95 | 1.25 | 1.67E-31 |
| 0 | TAGLN2 | 0.77 | 0.61 | 5.94 | 3.55 | 0.74 | 1.72E-31 |
| 0 | MRPS6 | 0.61 | 0.39 | 3.90 | 1.92 | 1.02 | 1.93E-31 |
| 0 | CREM | 0.86 | 0.69 | 12.29 | 8.38 | 0.55 | 3.10E-31 |
| 0 | DUSP4 | 0.47 | 0.22 | 2.55 | 1.44 | 0.82 | 2.08E-30 |
| 0 | CD7 | 0.84 | 0.70 | 8.98 | 5.65 | 0.67 | 5.65E-30 |
| 0 | RIN3 | 0.44 | 0.22 | 1.65 | 0.74 | 1.17 | 1.63E-29 |
| 0 | PRKX | 0.51 | 0.29 | 2.48 | 1.19 | 1.06 | 2.22E-29 |
| 0 | B4GALT1 | 0.70 | 0.48 | 4.03 | 2.41 | 0.74 | 2.38E-29 |
| 0 | FGFR1OP2 | 0.75 | 0.56 | 5.15 | 3.05 | 0.75 | 3.03E-29 |
| 0 | ZEB1 | 0.28 | 0.10 | 1.02 | 0.31 | 1.70 | 5.43E-29 |
| 0 | RPS2 | 0.97 | 0.97 | 33.35 | 24.99 | 0.42 | 5.79E-29 |
| 0 | RHBDF2 | 0.33 | 0.14 | 1.22 | 0.46 | 1.41 | 6.05E-29 |
| 0 | ZBTB16 | 0.40 | 0.19 | 2.23 | 0.89 | 1.32 | 6.31E-29 |
| 0 | RAB21 | 0.63 | 0.42 | 3.33 | 1.81 | 0.88 | 7.80E-29 |
| 0 | PTMA | 0.99 | 0.99 | 57.45 | 44.70 | 0.36 | 2.14E-28 |
| 0 | SYPL1 | 0.34 | 0.15 | 1.14 | 0.44 | 1.37 | 2.22E-28 |
| 0 | EML4 | 0.65 | 0.42 | 3.48 | 2.00 | 0.80 | 2.39E-28 |
| 0 | TRAF1 | 0.36 | 0.16 | 1.47 | 0.56 | 1.40 | 2.50E-28 |
| 0 | DENND4A | 0.47 | 0.26 | 2.21 | 0.94 | 1.24 | 2.95E-28 |
| 0 | SARAF | 0.86 | 0.75 | 8.59 | 5.87 | 0.55 | 1.21E-27 |
| 0 | STAG2 | 0.63 | 0.42 | 3.27 | 1.85 | 0.82 | 1.29E-27 |
| 0 | B3GNT7 | 0.43 | 0.22 | 1.69 | 0.86 | 0.97 | 1.49E-27 |
| 0 | SLC16A3 | 0.31 | 0.12 | 1.01 | 0.39 | 1.37 | 2.98E-27 |
| 0 | SEC14L1 | 0.48 | 0.27 | 2.14 | 0.99 | 1.11 | 4.16E-27 |
| 0 | ZFP36L2 | 0.90 | 0.80 | 15.96 | 10.27 | 0.64 | 9.52E-27 |
| 0 | MKNK2 | 0.62 | 0.41 | 3.42 | 1.91 | 0.84 | 1.43E-26 |
| 0 | TNFRSF9 | 0.44 | 0.24 | 4.41 | 1.72 | 1.36 | 2.61E-26 |
| 0 | GPX4 | 0.71 | 0.51 | 4.21 | 2.61 | 0.69 | 3.03E-26 |
| 0 | RALGAPA1 | 0.57 | 0.37 | 3.03 | 1.58 | 0.94 | 6.93E-26 |
| 0 | NFAT5 | 0.56 | 0.37 | 3.62 | 1.76 | 1.04 | 2.38E-25 |
| 0 | IKZF2 | 0.29 | 0.12 | 1.21 | 0.47 | 1.37 | 2.39E-25 |
| 0 | RPLP0 | 0.91 | 0.84 | 11.82 | 8.49 | 0.48 | 2.93E-25 |
| 0 | KDM5B | 0.47 | 0.26 | 2.04 | 1.07 | 0.93 | 2.99E-25 |
| 0 | SIPA1L1 | 0.29 | 0.12 | 1.11 | 0.38 | 1.56 | 3.51E-25 |
| 0 | SNX9 | 0.26 | 0.10 | 0.94 | 0.32 | 1.53 | 6.68E-25 |
| 0 | SESN1 | 0.30 | 0.13 | 1.23 | 0.49 | 1.33 | 8.65E-25 |
| 0 | WHRN | 0.36 | 0.18 | 1.75 | 0.74 | 1.24 | 1.08E-24 |

| | | | | | | | |
|---|---|---|---|---|---|---|---|
| 0 | FYN | 0.81 | 0.65 | 6.26 | 4.21 | 0.57 | 5.40E-24 |
| 0 | IFITM2 | 0.89 | 0.83 | 13.37 | 9.12 | 0.55 | 6.39E-24 |
| 0 | PIK3R1 | 0.84 | 0.71 | 11.88 | 7.91 | 0.59 | 6.41E-24 |
| 0 | ADGRE5 | 0.80 | 0.66 | 6.34 | 4.15 | 0.61 | 6.44E-24 |
| 0 | IL2RB | 0.68 | 0.51 | 4.17 | 2.63 | 0.67 | 6.57E-24 |
| 0 | RBPJ | 0.49 | 0.29 | 2.48 | 1.19 | 1.06 | 7.00E-24 |
| 0 | RERE | 0.31 | 0.14 | 1.08 | 0.47 | 1.21 | 1.03E-23 |
| 0 | LRRFIP1 | 0.78 | 0.65 | 7.89 | 4.90 | 0.69 | 1.27E-23 |
| 0 | CAPG | 0.27 | 0.11 | 0.97 | 0.40 | 1.29 | 2.46E-23 |
| 0 | VDAC1 | 0.48 | 0.28 | 1.81 | 0.96 | 0.92 | 2.87E-23 |
| 0 | EMD | 0.50 | 0.30 | 1.91 | 1.06 | 0.86 | 3.32E-23 |
| 0 | SLC38A1 | 0.75 | 0.59 | 5.01 | 3.32 | 0.59 | 3.52E-23 |
| 0 | SMAD3 | 0.27 | 0.11 | 0.93 | 0.34 | 1.47 | 6.23E-23 |
| 0 | PSMD13 | 0.41 | 0.22 | 1.56 | 0.74 | 1.07 | 6.94E-23 |
| 0 | PPP1R14B | 0.43 | 0.23 | 1.76 | 0.97 | 0.86 | 1.23E-22 |
| 0 | CTNNB1 | 0.53 | 0.33 | 2.60 | 1.36 | 0.94 | 2.04E-22 |
| 0 | ZHX2 | 0.28 | 0.12 | 0.96 | 0.36 | 1.43 | 3.39E-22 |
| 0 | TMSB4X | 0.99 | 0.99 | 118.30 | 87.21 | 0.44 | 4.49E-22 |
| 0 | FBXO34 | 0.43 | 0.24 | 1.69 | 0.84 | 1.01 | 4.67E-22 |
| 0 | BID | 0.31 | 0.14 | 0.94 | 0.43 | 1.11 | 5.99E-22 |
| 0 | EZR | 0.87 | 0.74 | 11.80 | 8.33 | 0.50 | 7.33E-22 |
| 0 | KLRD1 | 0.91 | 0.87 | 13.01 | 9.47 | 0.46 | 1.34E-21 |
| 0 | SKIL | 0.56 | 0.36 | 2.65 | 1.51 | 0.82 | 2.01E-21 |
| 0 | OGT | 0.58 | 0.38 | 2.77 | 1.65 | 0.75 | 2.24E-21 |
| 0 | FOXP1 | 0.46 | 0.27 | 2.37 | 1.11 | 1.09 | 2.49E-21 |
| 0 | ANKRD11 | 0.56 | 0.38 | 3.25 | 1.77 | 0.87 | 3.09E-21 |
| 0 | BICDL1 | 0.40 | 0.22 | 1.91 | 0.89 | 1.10 | 3.40E-21 |
| 0 | CHMP4B | 0.30 | 0.14 | 0.94 | 0.40 | 1.23 | 5.93E-21 |
| 0 | IL4R | 0.34 | 0.16 | 1.11 | 0.55 | 1.01 | 6.06E-21 |
| 0 | SYTL3 | 0.70 | 0.55 | 5.15 | 3.24 | 0.67 | 8.59E-21 |
| 0 | CLDND1 | 0.59 | 0.40 | 3.59 | 2.11 | 0.77 | 1.01E-20 |
| 0 | NCK2 | 0.30 | 0.14 | 0.93 | 0.42 | 1.15 | 1.01E-20 |
| 0 | GSPT1 | 0.56 | 0.36 | 2.78 | 1.66 | 0.74 | 1.49E-20 |
| 0 | BRD1 | 0.46 | 0.27 | 1.91 | 0.99 | 0.95 | 1.59E-20 |
| 0 | BCL2A1 | 0.49 | 0.31 | 3.24 | 1.79 | 0.86 | 2.00E-20 |
| 0 | SLC7A5 | 0.52 | 0.31 | 2.38 | 1.49 | 0.68 | 2.33E-20 |
| 0 | USP47 | 0.46 | 0.28 | 1.89 | 0.99 | 0.93 | 2.36E-20 |
| 0 | CXCR4 | 0.85 | 0.72 | 14.66 | 9.63 | 0.61 | 2.87E-20 |
| 0 | PGRMC2 | 0.27 | 0.12 | 0.95 | 0.40 | 1.24 | 6.93E-20 |
| 0 | BZW1 | 0.82 | 0.69 | 6.29 | 4.43 | 0.51 | 7.68E-20 |
| 0 | H3-3A | 0.94 | 0.87 | 12.52 | 9.57 | 0.39 | 8.46E-20 |
| 0 | HNRNPC | 0.86 | 0.72 | 6.72 | 5.01 | 0.43 | 1.33E-19 |
| 0 | TMEM120B | 0.47 | 0.29 | 1.82 | 0.98 | 0.89 | 1.43E-19 |
| 0 | TRGC1 | 0.52 | 0.33 | 2.66 | 1.42 | 0.90 | 1.46E-19 |
| 0 | CYLD | 0.70 | 0.52 | 4.19 | 2.66 | 0.65 | 1.74E-19 |
| 0 | CYSTM1 | 0.36 | 0.19 | 1.24 | 0.61 | 1.02 | 2.15E-19 |
| 0 | MSN | 0.79 | 0.69 | 6.80 | 4.59 | 0.57 | 2.64E-19 |
| 0 | CNOT6L | 0.83 | 0.68 | 7.65 | 5.43 | 0.49 | 3.03E-19 |
| 0 | SERPINB9 | 0.64 | 0.49 | 6.09 | 3.32 | 0.87 | 3.04E-19 |
| 0 | TNFRSF4 | 0.27 | 0.12 | 1.29 | 0.50 | 1.37 | 3.50E-19 |
| 0 | RHOF | 0.58 | 0.41 | 2.71 | 1.61 | 0.75 | 6.14E-19 |
| 0 | CPNE1 | 0.43 | 0.25 | 1.57 | 0.92 | 0.78 | 6.36E-19 |
| 0 | ASAP1 | 0.26 | 0.11 | 0.83 | 0.33 | 1.33 | 7.39E-19 |
| 0 | SYAP1 | 0.62 | 0.43 | 3.55 | 2.23 | 0.67 | 8.22E-19 |
| 0 | RANBP2 | 0.55 | 0.36 | 2.63 | 1.61 | 0.71 | 1.22E-18 |
| 0 | ELL2 | 0.48 | 0.29 | 2.00 | 1.23 | 0.69 | 1.73E-18 |
| 0 | AGPAT4 | 0.32 | 0.16 | 1.15 | 0.58 | 0.98 | 1.80E-18 |
| 0 | LYST | 0.62 | 0.44 | 3.75 | 2.50 | 0.59 | 2.06E-18 |
| 0 | PIM2 | 0.36 | 0.19 | 1.26 | 0.67 | 0.91 | 2.07E-18 |

| | | | | | | | |
|---|---|---|---|---|---|---|---|
| 0 | RUNX3 | 0.87 | 0.74 | 8.22 | 5.99 | 0.46 | 3.90E-18 |
| 0 | OTULIN | 0.58 | 0.42 | 3.17 | 1.79 | 0.83 | 4.76E-18 |
| 0 | FTH1 | 0.97 | 0.96 | 62.43 | 34.80 | 0.84 | 6.39E-18 |
| 0 | NAP1L1 | 0.72 | 0.57 | 4.56 | 3.02 | 0.59 | 1.37E-17 |
| 0 | SIK3 | 0.51 | 0.33 | 2.26 | 1.36 | 0.73 | 1.64E-17 |
| 0 | FXYD5 | 0.68 | 0.52 | 3.61 | 2.36 | 0.62 | 1.68E-17 |
| 0 | JARID2 | 0.50 | 0.32 | 2.12 | 1.22 | 0.80 | 3.85E-17 |
| 0 | MCTP2 | 0.50 | 0.32 | 2.31 | 1.43 | 0.70 | 4.14E-17 |
| 0 | TRAF4 | 0.33 | 0.18 | 1.10 | 0.60 | 0.88 | 9.35E-17 |
| 0 | IL21R | 0.37 | 0.21 | 1.36 | 0.75 | 0.86 | 1.07E-16 |
| 0 | STK4 | 0.91 | 0.84 | 10.39 | 7.88 | 0.40 | 2.06E-16 |
| 0 | GNA13 | 0.54 | 0.37 | 2.39 | 1.50 | 0.67 | 2.50E-16 |
| 0 | MRPL14 | 0.33 | 0.18 | 1.08 | 0.59 | 0.87 | 2.81E-16 |
| 0 | KMT2E | 0.87 | 0.77 | 7.85 | 6.02 | 0.38 | 3.25E-16 |
| 0 | COTL1 | 0.51 | 0.34 | 2.99 | 1.92 | 0.64 | 4.73E-16 |
| 0 | SKI | 0.36 | 0.20 | 1.24 | 0.72 | 0.78 | 6.63E-16 |
| 0 | PLP2 | 0.41 | 0.25 | 1.50 | 0.87 | 0.79 | 7.00E-16 |
| 0 | EIF5B | 0.51 | 0.35 | 2.43 | 1.46 | 0.74 | 7.23E-16 |
| 0 | CHD4 | 0.56 | 0.41 | 2.76 | 1.76 | 0.65 | 9.44E-16 |
| 0 | IRF8 | 0.45 | 0.28 | 2.30 | 1.41 | 0.70 | 9.53E-16 |
| 0 | LARP1 | 0.39 | 0.23 | 1.49 | 0.76 | 0.97 | 1.12E-15 |
| 0 | FAM107B | 0.65 | 0.51 | 4.68 | 2.96 | 0.66 | 1.72E-15 |
| 0 | AKNA | 0.64 | 0.49 | 3.63 | 2.44 | 0.57 | 2.43E-15 |
| 0 | PAK2 | 0.53 | 0.37 | 2.47 | 1.56 | 0.66 | 2.45E-15 |
| 0 | HIVEP2 | 0.34 | 0.19 | 1.18 | 0.65 | 0.85 | 2.64E-15 |
| 0 | QKI | 0.50 | 0.34 | 2.21 | 1.31 | 0.75 | 2.80E-15 |
| 0 | GABARAPL | 0.77 | 0.63 | 5.67 | 4.02 | 0.50 | 4.20E-15 |
| 0 | UBE2F | 0.42 | 0.28 | 1.87 | 1.03 | 0.86 | 4.51E-15 |
| 0 | GOLGB1 | 0.65 | 0.50 | 4.34 | 2.90 | 0.58 | 5.51E-15 |
| 0 | GALNT11 | 0.31 | 0.17 | 1.07 | 0.58 | 0.88 | 6.83E-15 |
| 0 | RPL23 | 0.87 | 0.79 | 8.99 | 6.69 | 0.43 | 7.94E-15 |
| 0 | HDGF | 0.51 | 0.34 | 2.05 | 1.32 | 0.63 | 1.10E-14 |
| 0 | RAP1B | 0.81 | 0.69 | 5.99 | 4.57 | 0.39 | 1.10E-14 |
| 0 | HBS1L | 0.31 | 0.17 | 1.05 | 0.55 | 0.94 | 1.53E-14 |
| 0 | TNIP1 | 0.39 | 0.24 | 1.50 | 0.87 | 0.79 | 2.19E-14 |
| 0 | TAB2 | 0.28 | 0.15 | 0.87 | 0.47 | 0.87 | 2.25E-14 |
| 0 | RALA | 0.35 | 0.20 | 1.19 | 0.69 | 0.78 | 3.63E-14 |
| 0 | CDC37 | 0.66 | 0.53 | 3.81 | 2.59 | 0.56 | 4.36E-14 |
| 0 | NOTCH2NL | 0.28 | 0.15 | 0.94 | 0.51 | 0.88 | 4.39E-14 |
| 0 | MBP | 0.69 | 0.55 | 4.55 | 3.26 | 0.48 | 4.50E-14 |
| 0 | CBLB | 0.49 | 0.32 | 2.13 | 1.37 | 0.64 | 4.92E-14 |
| 0 | STAT4 | 0.53 | 0.38 | 2.58 | 1.68 | 0.62 | 5.07E-14 |
| 0 | BRAF | 0.39 | 0.24 | 1.50 | 0.86 | 0.80 | 6.60E-14 |
| 0 | HDLBP | 0.28 | 0.15 | 0.97 | 0.51 | 0.93 | 6.85E-14 |
| 0 | PRRC2C | 0.84 | 0.75 | 7.14 | 5.43 | 0.40 | 6.95E-14 |
| 0 | ZC3H18 | 0.35 | 0.21 | 1.13 | 0.63 | 0.85 | 9.29E-14 |
| 0 | HIF1A | 0.59 | 0.42 | 2.83 | 1.98 | 0.51 | 9.40E-14 |
| 0 | ID2 | 0.74 | 0.64 | 8.41 | 5.74 | 0.55 | 9.48E-14 |
| 0 | CNOT2 | 0.53 | 0.40 | 2.87 | 1.73 | 0.73 | 1.23E-13 |
| 0 | TES | 0.48 | 0.33 | 2.20 | 1.37 | 0.69 | 1.82E-13 |
| 0 | BCL6 | 0.34 | 0.20 | 1.29 | 0.74 | 0.80 | 2.38E-13 |
| 0 | NFKBIZ | 0.51 | 0.36 | 2.93 | 1.95 | 0.59 | 2.76E-13 |
| 0 | STK17A | 0.85 | 0.75 | 7.69 | 5.80 | 0.41 | 3.12E-13 |
| 0 | ARID4B | 0.76 | 0.63 | 5.16 | 3.88 | 0.41 | 3.17E-13 |
| 0 | TRAF5 | 0.30 | 0.17 | 1.13 | 0.62 | 0.86 | 4.19E-13 |
| 0 | NR4A3 | 0.27 | 0.15 | 1.00 | 0.53 | 0.93 | 5.11E-13 |
| 0 | LDHA | 0.79 | 0.67 | 6.54 | 4.60 | 0.51 | 5.75E-13 |
| 0 | DHRS3 | 0.27 | 0.15 | 0.93 | 0.53 | 0.80 | 5.93E-13 |
| 0 | PBXIP1 | 0.36 | 0.23 | 1.52 | 0.86 | 0.83 | 8.92E-13 |

| | | | | | | | |
|---|---|---|---|---|---|---|---|
| 0 | CMTM6 | 0.51 | 0.36 | 2.00 | 1.32 | 0.60 | 9.87E-13 |
| 0 | TUBA4A | 0.79 | 0.65 | 9.35 | 6.90 | 0.44 | 9.91E-13 |
| 0 | RPL13A | 0.91 | 0.86 | 21.63 | 16.41 | 0.40 | 1.15E-12 |
| 0 | YBX3 | 0.37 | 0.23 | 2.27 | 1.33 | 0.78 | 1.19E-12 |
| 0 | DAAM1 | 0.27 | 0.15 | 1.12 | 0.58 | 0.94 | 1.53E-12 |
| 0 | ZNF331 | 0.63 | 0.47 | 4.63 | 3.46 | 0.42 | 1.60E-12 |
| 0 | PRDX6 | 0.57 | 0.43 | 2.82 | 1.79 | 0.66 | 1.61E-12 |
| 0 | GPR65 | 0.60 | 0.45 | 3.33 | 2.26 | 0.56 | 1.84E-12 |
| 0 | RAB8B | 0.81 | 0.71 | 6.76 | 5.12 | 0.40 | 2.02E-12 |
| 0 | ACTN4 | 0.64 | 0.48 | 3.20 | 2.21 | 0.53 | 2.03E-12 |
| 0 | RAD21 | 0.57 | 0.42 | 2.61 | 1.77 | 0.56 | 2.37E-12 |
| 0 | RBM17 | 0.41 | 0.26 | 1.50 | 0.93 | 0.69 | 2.53E-12 |
| 0 | RHOG | 0.56 | 0.40 | 2.32 | 1.61 | 0.52 | 2.53E-12 |
| 0 | RALGDS | 0.35 | 0.22 | 1.29 | 0.74 | 0.80 | 3.00E-12 |
| 0 | LPIN1 | 0.34 | 0.21 | 1.24 | 0.72 | 0.78 | 3.23E-12 |
| 0 | S100A11 | 0.69 | 0.57 | 4.92 | 3.58 | 0.46 | 3.49E-12 |
| 0 | PIM3 | 0.62 | 0.46 | 3.32 | 2.40 | 0.47 | 3.69E-12 |
| 0 | AKAP13 | 0.79 | 0.72 | 6.68 | 4.99 | 0.42 | 3.88E-12 |
| 0 | SACM1L | 0.42 | 0.29 | 1.56 | 1.00 | 0.64 | 4.79E-12 |
| 0 | CRYBG1 | 0.40 | 0.26 | 1.73 | 0.99 | 0.81 | 4.96E-12 |
| 0 | SLFN13 | 0.32 | 0.19 | 1.11 | 0.70 | 0.66 | 5.02E-12 |
| 0 | PABPC4 | 0.36 | 0.22 | 1.21 | 0.74 | 0.71 | 5.38E-12 |
| 0 | PHF20 | 0.58 | 0.45 | 3.02 | 2.06 | 0.55 | 7.14E-12 |
| 0 | USP12 | 0.35 | 0.22 | 1.31 | 0.76 | 0.78 | 9.31E-12 |
| 0 | RSL1D1 | 0.50 | 0.37 | 1.98 | 1.31 | 0.60 | 9.51E-12 |
| 0 | ST3GAL1 | 0.34 | 0.21 | 1.26 | 0.72 | 0.80 | 1.08E-11 |
| 0 | ETS1 | 0.76 | 0.63 | 5.16 | 3.91 | 0.40 | 1.21E-11 |
| 0 | ABHD17B | 0.25 | 0.14 | 0.78 | 0.40 | 0.96 | 1.36E-11 |
| 0 | SEC11A | 0.55 | 0.40 | 2.16 | 1.54 | 0.48 | 1.37E-11 |
| 0 | CTSZ | 0.25 | 0.14 | 0.79 | 0.44 | 0.86 | 1.82E-11 |
| 0 | SURF4 | 0.44 | 0.29 | 1.63 | 1.06 | 0.63 | 2.61E-11 |
| 0 | POLR2L | 0.54 | 0.40 | 2.35 | 1.58 | 0.57 | 2.72E-11 |
| 0 | KDM2B | 0.25 | 0.14 | 0.76 | 0.43 | 0.81 | 2.97E-11 |
| 0 | SPRY1 | 0.28 | 0.16 | 1.40 | 1.09 | 0.36 | 2.97E-11 |
| 0 | CD63 | 0.75 | 0.63 | 4.64 | 3.59 | 0.37 | 3.78E-11 |
| 0 | IPO7 | 0.29 | 0.18 | 0.95 | 0.54 | 0.81 | 4.49E-11 |
| 0 | TUT4 | 0.49 | 0.35 | 2.00 | 1.37 | 0.55 | 4.90E-11 |
| 0 | MARCHF7 | 0.37 | 0.24 | 1.33 | 0.84 | 0.66 | 5.41E-11 |
| 0 | CARD19 | 0.32 | 0.20 | 1.13 | 0.71 | 0.67 | 5.99E-11 |
| 0 | TIPARP | 0.51 | 0.37 | 2.31 | 1.69 | 0.46 | 6.37E-11 |
| 0 | CSNK1D | 0.51 | 0.35 | 1.93 | 1.38 | 0.48 | 7.01E-11 |
| 0 | PTK2B | 0.32 | 0.20 | 1.41 | 0.72 | 0.96 | 8.29E-11 |
| 0 | OTUD5 | 0.31 | 0.19 | 1.06 | 0.63 | 0.75 | 9.34E-11 |
| 0 | CCDC93 | 0.29 | 0.18 | 1.06 | 0.61 | 0.81 | 1.06E-10 |
| 0 | WDR48 | 0.29 | 0.18 | 0.92 | 0.53 | 0.80 | 1.09E-10 |
| 0 | KTN1 | 0.61 | 0.48 | 3.11 | 2.20 | 0.50 | 1.10E-10 |
| 0 | HOOK3 | 0.29 | 0.17 | 0.99 | 0.58 | 0.77 | 1.14E-10 |
| 0 | USP15 | 0.56 | 0.44 | 2.84 | 1.94 | 0.55 | 1.27E-10 |
| 0 | PDXK | 0.28 | 0.16 | 0.87 | 0.50 | 0.78 | 1.32E-10 |
| 0 | PCMTD1 | 0.37 | 0.23 | 1.31 | 0.84 | 0.65 | 1.41E-10 |
| 0 | PDCD4 | 0.57 | 0.42 | 2.85 | 2.11 | 0.44 | 1.81E-10 |
| 0 | AHR | 0.32 | 0.20 | 1.18 | 0.87 | 0.44 | 2.10E-10 |
| 0 | IRF2BP2 | 0.49 | 0.34 | 2.04 | 1.47 | 0.47 | 2.12E-10 |
| 0 | EDF1 | 0.76 | 0.67 | 4.62 | 3.54 | 0.39 | 2.21E-10 |
| 0 | ARRDC2 | 0.27 | 0.16 | 0.88 | 0.53 | 0.74 | 2.34E-10 |
| 0 | CMIP | 0.40 | 0.27 | 1.62 | 1.16 | 0.48 | 2.43E-10 |
| 0 | PSMD7 | 0.33 | 0.20 | 1.09 | 0.67 | 0.71 | 2.51E-10 |
| 0 | SOS2 | 0.32 | 0.20 | 1.09 | 0.68 | 0.69 | 2.66E-10 |
| 0 | LRPAP1 | 0.31 | 0.18 | 0.91 | 0.59 | 0.63 | 2.93E-10 |

| | | | | | | | |
|---|---|---|---|---|---|---|---|
| 0 | RAB9A | 0.49 | 0.37 | 2.44 | 1.54 | 0.66 | 3.30E-10 |
| 0 | SINHCAF | 0.29 | 0.18 | 0.95 | 0.58 | 0.71 | 4.25E-10 |
| 0 | LAPTM4A | 0.42 | 0.28 | 1.55 | 1.06 | 0.55 | 4.71E-10 |
| 0 | STT3B | 0.37 | 0.24 | 1.20 | 0.80 | 0.59 | 5.81E-10 |
| 0 | EHD1 | 0.38 | 0.25 | 1.43 | 0.95 | 0.59 | 5.92E-10 |
| 0 | SLC25A36 | 0.35 | 0.23 | 1.21 | 0.77 | 0.66 | 6.52E-10 |
| 0 | HCG18 | 0.33 | 0.21 | 1.20 | 0.78 | 0.63 | 6.60E-10 |
| 0 | SPAG9 | 0.48 | 0.35 | 2.08 | 1.39 | 0.58 | 6.94E-10 |
| 0 | NR3C1 | 0.67 | 0.55 | 3.97 | 3.04 | 0.39 | 7.09E-10 |
| 0 | BIRC2 | 0.44 | 0.31 | 1.77 | 1.26 | 0.50 | 7.12E-10 |
| 0 | EIF4G2 | 0.73 | 0.62 | 4.27 | 3.26 | 0.39 | 9.11E-10 |
| 0 | FNBP1 | 0.69 | 0.57 | 4.72 | 3.48 | 0.44 | 9.66E-10 |
| 0 | PRMT1 | 0.38 | 0.26 | 1.37 | 0.89 | 0.62 | 1.10E-09 |
| 0 | LFNG | 0.30 | 0.19 | 1.03 | 0.67 | 0.62 | 1.32E-09 |
| 0 | LAT2 | 0.38 | 0.26 | 1.48 | 1.01 | 0.54 | 1.74E-09 |
| 0 | IL18RAP | 0.30 | 0.19 | 1.24 | 0.76 | 0.70 | 1.98E-09 |
| 0 | RPS6KA3 | 0.38 | 0.27 | 1.54 | 1.01 | 0.61 | 2.00E-09 |
| 0 | SEPTIN11 | 0.27 | 0.16 | 0.95 | 0.54 | 0.80 | 2.16E-09 |
| 0 | ZCCHC2 | 0.36 | 0.24 | 1.23 | 0.86 | 0.51 | 2.17E-09 |
| 0 | CSNK2A1 | 0.28 | 0.17 | 0.92 | 0.59 | 0.65 | 2.35E-09 |
| 0 | CLPP | 0.26 | 0.16 | 0.82 | 0.45 | 0.85 | 2.53E-09 |
| 0 | APMAP | 0.61 | 0.46 | 2.88 | 2.24 | 0.37 | 3.30E-09 |
| 0 | MAPKAPK2 | 0.46 | 0.33 | 1.73 | 1.22 | 0.50 | 4.34E-09 |
| 0 | DDX21 | 0.64 | 0.53 | 3.80 | 2.88 | 0.40 | 4.34E-09 |
| 0 | NPEPPS | 0.33 | 0.22 | 1.06 | 0.73 | 0.53 | 4.40E-09 |
| 0 | TRABD | 0.40 | 0.27 | 1.48 | 1.02 | 0.54 | 4.72E-09 |
| 0 | TMBIM6 | 0.70 | 0.58 | 3.70 | 2.88 | 0.36 | 5.18E-09 |
| 0 | IFNGR1 | 0.46 | 0.33 | 1.99 | 1.45 | 0.46 | 5.30E-09 |
| 0 | GAS5 | 0.64 | 0.53 | 3.36 | 2.59 | 0.38 | 5.46E-09 |
| 0 | ZNF267 | 0.39 | 0.26 | 1.47 | 0.96 | 0.62 | 6.49E-09 |
| 0 | CD300A | 0.46 | 0.35 | 2.46 | 1.59 | 0.63 | 6.65E-09 |
| 0 | RPS17 | 0.52 | 0.40 | 2.16 | 1.58 | 0.45 | 7.08E-09 |
| 0 | CEBPD | 0.37 | 0.25 | 2.30 | 1.64 | 0.49 | 7.67E-09 |
| 0 | UBE2M | 0.42 | 0.29 | 1.44 | 1.02 | 0.50 | 1.12E-08 |
| 0 | NDUFS5 | 0.60 | 0.47 | 2.77 | 2.08 | 0.41 | 1.16E-08 |
| 0 | UBE2I | 0.51 | 0.37 | 1.85 | 1.42 | 0.38 | 1.22E-08 |
| 0 | ARHGEF2 | 0.32 | 0.21 | 1.11 | 0.69 | 0.68 | 1.25E-08 |
| 0 | PDE7A | 0.41 | 0.29 | 1.61 | 1.10 | 0.54 | 1.45E-08 |
| 0 | KMT2C | 0.50 | 0.38 | 2.28 | 1.69 | 0.43 | 1.76E-08 |
| 0 | NARF | 0.32 | 0.21 | 1.15 | 0.69 | 0.73 | 1.76E-08 |
| 0 | RELA | 0.27 | 0.17 | 0.92 | 0.54 | 0.76 | 1.89E-08 |
| 0 | HMGA1 | 0.33 | 0.22 | 1.05 | 0.72 | 0.54 | 2.07E-08 |
| 0 | POLR2K | 0.48 | 0.36 | 2.03 | 1.47 | 0.46 | 2.29E-08 |
| 0 | NBEAL1 | 0.26 | 0.16 | 0.87 | 0.53 | 0.71 | 2.43E-08 |
| 0 | FLNA | 0.70 | 0.59 | 4.51 | 3.35 | 0.43 | 2.50E-08 |
| 0 | IL10RA | 0.48 | 0.37 | 2.23 | 1.49 | 0.58 | 2.58E-08 |
| 0 | ZC3H7A | 0.30 | 0.19 | 0.95 | 0.63 | 0.61 | 3.12E-08 |
| 0 | FMNL1 | 0.64 | 0.51 | 3.24 | 2.38 | 0.44 | 3.42E-08 |
| 0 | MAFF | 0.36 | 0.24 | 1.43 | 1.10 | 0.38 | 3.58E-08 |
| 0 | ARFGAP3 | 0.42 | 0.31 | 1.66 | 1.16 | 0.52 | 4.24E-08 |
| 0 | ZNF706 | 0.44 | 0.33 | 1.69 | 1.17 | 0.53 | 4.37E-08 |
| 0 | ELOVL5 | 0.48 | 0.37 | 2.05 | 1.48 | 0.47 | 4.40E-08 |
| 0 | CLINT1 | 0.39 | 0.27 | 1.44 | 0.96 | 0.58 | 4.43E-08 |
| 0 | NIBAN1 | 0.28 | 0.19 | 1.23 | 0.72 | 0.79 | 5.14E-08 |
| 0 | AKAP17A | 0.38 | 0.28 | 1.44 | 0.90 | 0.67 | 5.18E-08 |
| 0 | CEBPZ | 0.42 | 0.31 | 1.53 | 1.10 | 0.48 | 5.31E-08 |
| 0 | EIF3E | 0.61 | 0.49 | 2.78 | 2.15 | 0.37 | 5.32E-08 |
| 0 | N4BP1 | 0.35 | 0.24 | 1.27 | 0.91 | 0.48 | 5.35E-08 |
| 0 | MORF4L2 | 0.48 | 0.35 | 1.85 | 1.39 | 0.41 | 5.87E-08 |

| | | | | | | | |
|---|---|---|---|---|---|---|---|
| 0 | NFKBID | 0.35 | 0.25 | 1.65 | 1.13 | 0.55 | 6.71E-08 |
| 0 | ATP1B3 | 0.66 | 0.56 | 6.78 | 4.92 | 0.46 | 7.57E-08 |
| 0 | TUBB | 0.60 | 0.47 | 2.91 | 2.24 | 0.38 | 7.95E-08 |
| 0 | TM9SF3 | 0.44 | 0.33 | 1.65 | 1.18 | 0.49 | 8.67E-08 |
| 0 | CAST | 0.61 | 0.51 | 4.16 | 2.81 | 0.57 | 9.28E-08 |
| 0 | RNF145 | 0.35 | 0.25 | 1.22 | 0.84 | 0.54 | 9.39E-08 |
| 0 | GSTO1 | 0.38 | 0.26 | 1.35 | 0.98 | 0.46 | 9.43E-08 |
| 0 | FTX | 0.28 | 0.18 | 1.01 | 0.62 | 0.70 | 9.70E-08 |
| 0 | CIAO2B | 0.43 | 0.32 | 1.63 | 1.11 | 0.56 | 1.02E-07 |
| 0 | SEMA4D | 0.46 | 0.34 | 1.90 | 1.35 | 0.50 | 1.03E-07 |
| 0 | ASH1L | 0.48 | 0.36 | 2.11 | 1.55 | 0.45 | 1.04E-07 |
| 0 | TACC1 | 0.44 | 0.33 | 1.79 | 1.26 | 0.51 | 1.06E-07 |
| 0 | LUC7L | 0.32 | 0.21 | 1.01 | 0.72 | 0.49 | 1.13E-07 |
| 0 | NIPBL | 0.47 | 0.36 | 1.95 | 1.44 | 0.43 | 1.30E-07 |
| 0 | THRAP3 | 0.48 | 0.35 | 1.86 | 1.40 | 0.41 | 1.48E-07 |
| 0 | USP1 | 0.27 | 0.18 | 0.90 | 0.59 | 0.59 | 1.48E-07 |
| 0 | ITM2A | 0.45 | 0.35 | 2.60 | 1.80 | 0.53 | 1.62E-07 |
| 0 | CBX6 | 0.26 | 0.16 | 0.85 | 0.54 | 0.65 | 1.78E-07 |
| 0 | MIR23AHG | 0.48 | 0.38 | 4.31 | 2.70 | 0.68 | 1.84E-07 |
| 0 | SETD5 | 0.42 | 0.31 | 1.70 | 1.20 | 0.50 | 1.86E-07 |
| 0 | VAPA | 0.56 | 0.46 | 2.39 | 1.79 | 0.41 | 2.27E-07 |
| 0 | BCLAF1 | 0.55 | 0.44 | 2.67 | 2.04 | 0.38 | 2.29E-07 |
| 0 | UBA6 | 0.28 | 0.18 | 0.87 | 0.57 | 0.59 | 2.48E-07 |
| 0 | EIF1AX | 0.53 | 0.42 | 2.63 | 1.95 | 0.43 | 2.85E-07 |
| 0 | GYPC | 0.45 | 0.34 | 1.91 | 1.42 | 0.43 | 2.93E-07 |
| 0 | PHF1 | 0.40 | 0.29 | 1.39 | 1.00 | 0.48 | 2.93E-07 |
| 0 | CAB39 | 0.26 | 0.17 | 0.81 | 0.54 | 0.58 | 2.95E-07 |
| 0 | OXSR1 | 0.27 | 0.18 | 0.83 | 0.60 | 0.47 | 3.62E-07 |
| 0 | SMARCC1 | 0.27 | 0.18 | 0.86 | 0.60 | 0.51 | 3.66E-07 |
| 0 | HSP90B1 | 0.75 | 0.67 | 6.02 | 4.60 | 0.39 | 4.08E-07 |
| 0 | ANP32A | 0.47 | 0.36 | 1.86 | 1.36 | 0.45 | 4.25E-07 |
| 0 | PA2G4 | 0.50 | 0.39 | 2.11 | 1.56 | 0.44 | 4.42E-07 |
| 0 | APLP2 | 0.37 | 0.27 | 1.41 | 0.99 | 0.51 | 4.53E-07 |
| 0 | EIF5A | 0.57 | 0.45 | 2.50 | 1.91 | 0.39 | 5.82E-07 |
| 0 | RWDD1 | 0.40 | 0.29 | 1.55 | 1.08 | 0.52 | 6.04E-07 |
| 0 | PUM1 | 0.38 | 0.28 | 1.30 | 0.93 | 0.49 | 6.23E-07 |
| 0 | EIF3J | 0.43 | 0.33 | 1.89 | 1.33 | 0.51 | 7.41E-07 |
| 0 | PCMT1 | 0.32 | 0.22 | 1.00 | 0.72 | 0.47 | 7.64E-07 |
| 0 | FBL | 0.35 | 0.25 | 1.13 | 0.78 | 0.52 | 7.82E-07 |
| 0 | HUWE1 | 0.43 | 0.32 | 1.69 | 1.26 | 0.43 | 8.05E-07 |
| 0 | RNPS1 | 0.43 | 0.32 | 1.57 | 1.13 | 0.48 | 8.63E-07 |
| 0 | H2AJ | 0.50 | 0.40 | 2.87 | 1.96 | 0.55 | 8.90E-07 |
| 0 | BEX4 | 0.27 | 0.17 | 0.83 | 0.60 | 0.47 | 9.18E-07 |
| 0 | SLC39A10 | 0.34 | 0.24 | 1.37 | 0.95 | 0.53 | 9.55E-07 |
| 0 | SAFB2 | 0.37 | 0.27 | 1.31 | 0.95 | 0.46 | 1.01E-06 |
| 0 | PRKAR1A | 0.45 | 0.35 | 1.76 | 1.29 | 0.45 | 1.06E-06 |
| 0 | NPAT | 0.28 | 0.19 | 0.87 | 0.65 | 0.42 | 1.08E-06 |
| 0 | DHX9 | 0.32 | 0.23 | 1.09 | 0.78 | 0.48 | 1.21E-06 |
| 0 | CSNK1G3 | 0.34 | 0.24 | 1.08 | 0.81 | 0.41 | 1.27E-06 |
| 0 | TXNRD1 | 0.32 | 0.23 | 1.32 | 0.90 | 0.56 | 1.34E-06 |
| 0 | NAP1L4 | 0.52 | 0.41 | 2.32 | 1.78 | 0.38 | 1.37E-06 |
| 0 | TENT4B | 0.39 | 0.29 | 1.53 | 1.13 | 0.43 | 1.53E-06 |
| 0 | SEPTIN2 | 0.35 | 0.26 | 1.20 | 0.85 | 0.50 | 1.57E-06 |
| 0 | MED13 | 0.40 | 0.29 | 1.42 | 1.05 | 0.43 | 1.58E-06 |
| 0 | YARS1 | 0.32 | 0.22 | 1.07 | 0.73 | 0.55 | 1.63E-06 |
| 0 | MAT2A | 0.37 | 0.27 | 1.41 | 1.06 | 0.42 | 1.70E-06 |
| 0 | RPL7L1 | 0.33 | 0.24 | 1.09 | 0.77 | 0.51 | 1.76E-06 |
| 0 | DDAH2 | 0.30 | 0.21 | 1.08 | 0.73 | 0.56 | 1.97E-06 |
| 0 | UBE2R2 | 0.36 | 0.26 | 1.16 | 0.89 | 0.39 | 2.33E-06 |

| | | | | | | | |
|---|---|---|---|---|---|---|---|
| 0 | PSME4 | 0.36 | 0.26 | 1.26 | 0.94 | 0.41 | 2.37E-06 |
| 0 | CLK4 | 0.26 | 0.17 | 0.76 | 0.55 | 0.46 | 2.61E-06 |
| 0 | CSNK1A1 | 0.41 | 0.31 | 1.44 | 1.05 | 0.45 | 2.63E-06 |
| 0 | CUL3 | 0.35 | 0.25 | 1.21 | 0.85 | 0.52 | 2.76E-06 |
| 0 | KAT6A | 0.42 | 0.32 | 1.56 | 1.19 | 0.39 | 2.92E-06 |
| 0 | KCTD20 | 0.27 | 0.19 | 0.89 | 0.63 | 0.50 | 2.96E-06 |
| 0 | AHNAK | 0.58 | 0.50 | 3.68 | 2.75 | 0.42 | 2.97E-06 |
| 0 | PLEKHO1 | 0.32 | 0.22 | 1.07 | 0.81 | 0.40 | 3.02E-06 |
| 0 | GNL3 | 0.32 | 0.23 | 1.07 | 0.82 | 0.38 | 3.24E-06 |
| 0 | RANGAP1 | 0.30 | 0.21 | 1.05 | 0.73 | 0.52 | 3.60E-06 |
| 0 | MAPK1IP1L | 0.28 | 0.19 | 0.85 | 0.60 | 0.52 | 3.77E-06 |
| 0 | SETD2 | 0.45 | 0.36 | 1.89 | 1.41 | 0.42 | 4.12E-06 |
| 0 | RNF19A | 0.72 | 0.66 | 6.23 | 4.85 | 0.36 | 4.33E-06 |
| 0 | TAX1BP1 | 0.59 | 0.50 | 3.01 | 2.32 | 0.38 | 4.57E-06 |
| 0 | MAT2B | 0.29 | 0.20 | 0.95 | 0.68 | 0.48 | 5.50E-06 |
| 0 | TNRC6B | 0.48 | 0.39 | 2.37 | 1.76 | 0.43 | 5.58E-06 |
| 0 | SELENOS | 0.37 | 0.27 | 1.37 | 1.03 | 0.40 | 5.73E-06 |
| 0 | COPB1 | 0.31 | 0.22 | 1.08 | 0.75 | 0.53 | 8.60E-06 |
| 0 | IL27RA | 0.27 | 0.18 | 0.86 | 0.62 | 0.48 | 1.07E-05 |
| 0 | TTC39C | 0.30 | 0.22 | 1.10 | 0.74 | 0.56 | 1.12E-05 |
| 0 | LINC01138 | 0.27 | 0.19 | 1.03 | 0.78 | 0.41 | 1.12E-05 |
| 0 | OFD1 | 0.54 | 0.44 | 3.01 | 2.08 | 0.53 | 1.19E-05 |
| 0 | CRCP | 0.28 | 0.19 | 0.83 | 0.62 | 0.44 | 1.21E-05 |
| 0 | RELB | 0.45 | 0.35 | 1.80 | 1.36 | 0.40 | 1.23E-05 |
| 0 | SRPK1 | 0.29 | 0.21 | 0.97 | 0.71 | 0.46 | 1.35E-05 |
| 0 | EIF2AK3 | 0.27 | 0.19 | 1.04 | 0.74 | 0.49 | 1.37E-05 |
| 0 | CCNT1 | 0.30 | 0.22 | 1.02 | 0.71 | 0.52 | 1.39E-05 |
| 0 | G3BP2 | 0.59 | 0.50 | 3.09 | 2.35 | 0.39 | 1.46E-05 |
| 0 | MACROH2A | 0.25 | 0.18 | 0.80 | 0.55 | 0.55 | 1.67E-05 |
| 0 | LAMTOR5 | 0.34 | 0.25 | 1.09 | 0.82 | 0.40 | 1.92E-05 |
| 0 | DYNLT1 | 0.29 | 0.20 | 0.98 | 0.71 | 0.46 | 1.96E-05 |
| 0 | PIK3AP1 | 0.30 | 0.22 | 1.10 | 0.73 | 0.60 | 2.07E-05 |
| 0 | BDP1 | 0.42 | 0.33 | 1.83 | 1.42 | 0.37 | 2.12E-05 |
| 0 | KPNA4 | 0.44 | 0.35 | 1.70 | 1.27 | 0.42 | 2.26E-05 |
| 0 | DNAJC1 | 0.47 | 0.37 | 1.90 | 1.46 | 0.38 | 2.36E-05 |
| 0 | ZC3H15 | 0.42 | 0.33 | 1.62 | 1.22 | 0.40 | 2.67E-05 |
| 0 | RLF | 0.33 | 0.25 | 1.18 | 0.90 | 0.39 | 2.80E-05 |
| 0 | FLOT1 | 0.26 | 0.18 | 0.81 | 0.59 | 0.46 | 3.10E-05 |
| 0 | ATP11B | 0.33 | 0.25 | 1.14 | 0.81 | 0.50 | 3.45E-05 |
| 0 | PCNP | 0.36 | 0.27 | 1.17 | 0.89 | 0.39 | 3.87E-05 |
| 0 | TMED2 | 0.26 | 0.19 | 0.80 | 0.59 | 0.44 | 4.77E-05 |
| 0 | WDR82 | 0.38 | 0.29 | 1.31 | 1.00 | 0.39 | 4.98E-05 |
| 0 | ETF1 | 0.41 | 0.33 | 1.51 | 1.17 | 0.37 | 5.43E-05 |
| 0 | SCFD1 | 0.28 | 0.20 | 0.93 | 0.70 | 0.41 | 6.30E-05 |
| 0 | POLE3 | 0.29 | 0.21 | 0.97 | 0.73 | 0.40 | 6.34E-05 |
| 0 | C1QBP | 0.27 | 0.19 | 0.88 | 0.63 | 0.49 | 6.70E-05 |
| 0 | EPB41L4A-A | 0.34 | 0.25 | 1.05 | 0.81 | 0.38 | 6.84E-05 |
| 0 | C11orf58 | 0.46 | 0.37 | 1.83 | 1.42 | 0.36 | 7.03E-05 |
| 0 | PPA1 | 0.32 | 0.24 | 1.10 | 0.83 | 0.40 | 7.44E-05 |
| 0 | SMC4 | 0.29 | 0.21 | 1.06 | 0.80 | 0.39 | 7.57E-05 |
| 0 | FYTTD1 | 0.33 | 0.25 | 1.10 | 0.82 | 0.41 | 7.77E-05 |
| 0 | CELF1 | 0.32 | 0.24 | 1.08 | 0.84 | 0.37 | 7.85E-05 |
| 0 | CCT8 | 0.36 | 0.28 | 1.23 | 0.95 | 0.38 | 8.01E-05 |
| 0 | RSF1 | 0.38 | 0.30 | 1.47 | 1.13 | 0.38 | 8.60E-05 |
| 0 | IRF7 | 0.26 | 0.19 | 0.98 | 0.66 | 0.57 | 9.03E-05 |
| 0 | TNRC6C | 0.36 | 0.28 | 1.38 | 1.04 | 0.41 | 9.57E-05 |
| 0 | PPM1G | 0.41 | 0.33 | 1.60 | 1.22 | 0.40 | 9.85E-05 |
| 0 | ENO1 | 0.57 | 0.50 | 2.88 | 2.24 | 0.36 | 0.00010431 |
| 0 | H1-4 | 0.33 | 0.25 | 1.40 | 1.07 | 0.39 | 0.00011201 |

| | | | | | | | |
|---|---|---|---|---|---|---|---|
| 0 | DDX27 | 0.31 | 0.23 | 1.12 | 0.82 | 0.45 | 0.00012746 |
| 0 | MRFAP1 | 0.47 | 0.38 | 1.75 | 1.34 | 0.39 | 0.00013098 |
| 0 | RTF2 | 0.27 | 0.20 | 0.86 | 0.63 | 0.46 | 0.00015176 |
| 0 | NT5C3A | 0.26 | 0.19 | 0.79 | 0.61 | 0.37 | 0.00015349 |
| 0 | ZFC3H1 | 0.40 | 0.32 | 1.62 | 1.23 | 0.39 | 0.00017484 |
| 0 | BRD4 | 0.44 | 0.35 | 1.65 | 1.28 | 0.37 | 0.00022065 |
| 0 | BNIP3L | 0.30 | 0.24 | 1.15 | 0.82 | 0.48 | 0.00022671 |
| 0 | TOR1AIP2 | 0.29 | 0.21 | 0.91 | 0.71 | 0.36 | 0.00023246 |
| 0 | SIVA1 | 0.32 | 0.25 | 1.07 | 0.80 | 0.41 | 0.00025112 |
| 0 | CPD | 0.33 | 0.26 | 1.28 | 0.97 | 0.40 | 0.0003038 |
| 0 | CDK17 | 0.28 | 0.21 | 0.98 | 0.70 | 0.47 | 0.00031372 |
| 0 | TMEM243 | 0.30 | 0.22 | 1.00 | 0.76 | 0.39 | 0.00034123 |
| 0 | USP9X | 0.27 | 0.20 | 0.85 | 0.66 | 0.37 | 0.0004343 |
| 0 | RB1CC1 | 0.29 | 0.22 | 1.02 | 0.77 | 0.39 | 0.00046941 |
| 0 | COX5A | 0.37 | 0.31 | 1.30 | 0.99 | 0.40 | 0.00060783 |
| 0 | SAR1B | 0.28 | 0.22 | 1.00 | 0.73 | 0.46 | 0.00060854 |
| 0 | CD83 | 0.28 | 0.22 | 1.79 | 1.31 | 0.45 | 0.00062001 |
| 0 | SH2D1A | 0.43 | 0.37 | 2.18 | 1.58 | 0.46 | 0.00069527 |
| 0 | SERINC3 | 0.26 | 0.19 | 0.81 | 0.63 | 0.37 | 0.00085305 |
| 0 | INPP5D | 0.32 | 0.25 | 1.17 | 0.90 | 0.38 | 0.00093172 |
| 0 | RANBP1 | 0.27 | 0.21 | 0.93 | 0.71 | 0.39 | 0.00136047 |
| 0 | UBR5 | 0.30 | 0.23 | 1.01 | 0.78 | 0.37 | 0.00144087 |
| 0 | UQCRC2 | 0.29 | 0.23 | 0.97 | 0.76 | 0.36 | 0.00144527 |
| 0 | HSH2D | 0.28 | 0.22 | 1.13 | 0.87 | 0.38 | 0.00148124 |
| 0 | INSIG1 | 0.39 | 0.33 | 2.46 | 1.59 | 0.63 | 0.00189876 |
| 0 | CWC25 | 0.28 | 0.22 | 0.92 | 0.71 | 0.37 | 0.00206462 |
| 0 | SLA2 | 0.33 | 0.27 | 1.41 | 1.10 | 0.37 | 0.0039543 |
| 0 | CEMIP2 | 0.55 | 0.52 | 4.59 | 3.39 | 0.44 | 0.00517124 |
| 0 | RAB5A | 0.25 | 0.20 | 0.81 | 0.60 | 0.44 | 0.00558094 |
| 1 | FGFBP2 | 0.71 | 0.14 | 6.11 | 1.10 | 2.47 | 2.58E-139 |
| 1 | NKG7 | 0.99 | 0.91 | 44.83 | 18.03 | 1.31 | 8.41E-111 |
| 1 | FCGR3A | 0.75 | 0.25 | 5.76 | 1.53 | 1.91 | 1.53E-97 |
| 1 | GZMH | 0.78 | 0.29 | 7.48 | 2.05 | 1.87 | 4.94E-96 |
| 1 | S1PR5 | 0.53 | 0.13 | 2.64 | 0.53 | 2.32 | 2.78E-81 |
| 1 | RORA | 0.80 | 0.44 | 7.86 | 2.89 | 1.44 | 2.05E-64 |
| 1 | PPP2R5C | 0.83 | 0.53 | 7.35 | 2.83 | 1.38 | 1.26E-63 |
| 1 | SYNE2 | 0.75 | 0.36 | 5.90 | 2.17 | 1.44 | 2.06E-59 |
| 1 | CCL5 | 0.96 | 0.78 | 36.09 | 18.12 | 0.99 | 4.15E-59 |
| 1 | GZMB | 0.96 | 0.62 | 20.72 | 12.50 | 0.73 | 5.53E-59 |
| 1 | B2M | 1.00 | 1.00 | 131.57 | 89.35 | 0.56 | 6.95E-59 |
| 1 | PRDM1 | 0.56 | 0.20 | 3.11 | 1.01 | 1.62 | 1.04E-57 |
| 1 | DSTN | 0.62 | 0.28 | 4.22 | 1.16 | 1.86 | 1.42E-57 |
| 1 | TGFBR3 | 0.42 | 0.12 | 2.14 | 0.43 | 2.32 | 4.37E-55 |
| 1 | PYHIN1 | 0.54 | 0.20 | 3.02 | 0.85 | 1.83 | 7.73E-55 |
| 1 | CST7 | 0.93 | 0.76 | 15.05 | 6.97 | 1.11 | 4.68E-53 |
| 1 | HLA-C | 0.99 | 0.97 | 29.55 | 18.82 | 0.65 | 1.75E-52 |
| 1 | PRF1 | 0.80 | 0.52 | 8.30 | 3.21 | 1.37 | 3.73E-52 |
| 1 | HLA-E | 0.97 | 0.91 | 17.87 | 10.51 | 0.77 | 6.70E-52 |
| 1 | HLA-B | 1.00 | 0.99 | 48.36 | 33.05 | 0.55 | 5.54E-50 |
| 1 | IGF2R | 0.58 | 0.26 | 3.07 | 0.95 | 1.69 | 2.41E-49 |
| 1 | ZEB2 | 0.83 | 0.57 | 9.34 | 4.09 | 1.19 | 5.49E-49 |
| 1 | GNG2 | 0.75 | 0.41 | 6.17 | 2.47 | 1.32 | 4.30E-48 |
| 1 | GTF3C1 | 0.46 | 0.17 | 2.28 | 0.61 | 1.89 | 6.54E-47 |
| 1 | PTPRC | 0.98 | 0.91 | 22.45 | 13.37 | 0.75 | 3.25E-45 |
| 1 | ENC1 | 0.34 | 0.09 | 1.48 | 0.36 | 2.02 | 5.54E-44 |
| 1 | ARL4C | 0.87 | 0.64 | 9.65 | 5.09 | 0.92 | 9.49E-43 |
| 1 | SPON2 | 0.37 | 0.10 | 1.93 | 0.57 | 1.75 | 2.46E-42 |
| 1 | PLEK | 0.68 | 0.39 | 5.01 | 2.08 | 1.27 | 3.98E-40 |
| 1 | TNFRSF1B | 0.80 | 0.56 | 6.15 | 3.00 | 1.04 | 1.10E-39 |

| | | | | | | | |
|---|---|---|---|---|---|---|---|
| 1 | FTL | 0.98 | 0.97 | 46.89 | 25.33 | 0.89 | 9.95E-39 |
| 1 | S100A6 | 0.84 | 0.66 | 9.88 | 5.01 | 0.98 | 1.43E-38 |
| 1 | CCL4 | 0.82 | 0.49 | 32.30 | 20.88 | 0.63 | 3.16E-37 |
| 1 | ABHD17A | 0.75 | 0.51 | 4.78 | 2.28 | 1.07 | 3.69E-34 |
| 1 | ZBTB38 | 0.38 | 0.14 | 1.64 | 0.51 | 1.70 | 3.79E-33 |
| 1 | EFHD2 | 0.77 | 0.55 | 5.98 | 2.97 | 1.01 | 4.74E-33 |
| 1 | TFDP2 | 0.37 | 0.14 | 1.60 | 0.48 | 1.74 | 5.23E-33 |
| 1 | CYBA | 0.89 | 0.75 | 11.04 | 6.27 | 0.82 | 1.11E-31 |
| 1 | C12orf75 | 0.48 | 0.23 | 2.38 | 0.89 | 1.42 | 1.79E-31 |
| 1 | ABI3 | 0.33 | 0.12 | 1.37 | 0.40 | 1.79 | 3.82E-31 |
| 1 | DTHD1 | 0.28 | 0.09 | 1.39 | 0.31 | 2.17 | 1.03E-30 |
| 1 | CTSC | 0.65 | 0.40 | 4.19 | 2.02 | 1.05 | 3.16E-30 |
| 1 | CD48 | 0.74 | 0.55 | 4.99 | 2.62 | 0.93 | 5.09E-29 |
| 1 | CHST12 | 0.55 | 0.30 | 2.96 | 1.28 | 1.21 | 9.19E-29 |
| 1 | HLA-A | 0.98 | 0.96 | 29.21 | 21.01 | 0.48 | 1.09E-28 |
| 1 | LGALS1 | 0.85 | 0.64 | 10.61 | 6.23 | 0.77 | 1.41E-28 |
| 1 | ARPC2 | 0.92 | 0.83 | 10.21 | 6.47 | 0.66 | 2.83E-28 |
| 1 | IFI16 | 0.77 | 0.58 | 6.10 | 3.23 | 0.92 | 8.56E-28 |
| 1 | PTMS | 0.39 | 0.17 | 2.02 | 0.72 | 1.49 | 1.59E-27 |
| 1 | S100A4 | 0.78 | 0.61 | 9.18 | 4.81 | 0.93 | 8.59E-27 |
| 1 | CCL4L2 | 0.47 | 0.22 | 11.80 | 5.78 | 1.03 | 2.83E-26 |
| 1 | ITGB2 | 0.56 | 0.32 | 3.09 | 1.38 | 1.16 | 1.01E-25 |
| 1 | PTP4A2 | 0.72 | 0.54 | 4.70 | 2.43 | 0.95 | 2.33E-25 |
| 1 | GK5 | 0.33 | 0.14 | 1.58 | 0.46 | 1.79 | 3.47E-25 |
| 1 | FGL2 | 0.26 | 0.09 | 1.39 | 0.44 | 1.67 | 1.17E-24 |
| 1 | CELF2 | 0.53 | 0.32 | 3.07 | 1.26 | 1.28 | 1.22E-24 |
| 1 | CLEC2D | 0.65 | 0.44 | 5.58 | 2.66 | 1.07 | 2.83E-24 |
| 1 | ORAI1 | 0.49 | 0.28 | 2.40 | 0.98 | 1.30 | 2.93E-24 |
| 1 | PRKCH | 0.57 | 0.36 | 2.84 | 1.35 | 1.07 | 3.06E-24 |
| 1 | BCL11B | 0.39 | 0.18 | 1.75 | 0.71 | 1.31 | 6.34E-24 |
| 1 | SLC15A4 | 0.39 | 0.19 | 1.71 | 0.64 | 1.41 | 9.22E-24 |
| 1 | F2R | 0.29 | 0.11 | 1.09 | 0.37 | 1.58 | 1.23E-23 |
| 1 | TMSB10 | 0.98 | 0.96 | 28.93 | 20.28 | 0.51 | 1.81E-23 |
| 1 | SYNE1 | 0.46 | 0.24 | 2.48 | 1.00 | 1.31 | 4.03E-23 |
| 1 | MT2A | 0.65 | 0.44 | 10.88 | 5.31 | 1.03 | 4.88E-23 |
| 1 | C1orf21 | 0.45 | 0.24 | 1.99 | 0.83 | 1.26 | 9.37E-23 |
| 1 | ADGRG1 | 0.27 | 0.10 | 1.17 | 0.35 | 1.72 | 1.50E-22 |
| 1 | KLF2 | 0.62 | 0.37 | 4.81 | 3.03 | 0.67 | 1.65E-22 |
| 1 | ISG20 | 0.72 | 0.55 | 6.23 | 3.62 | 0.78 | 1.97E-22 |
| 1 | GCLM | 0.40 | 0.21 | 2.14 | 0.78 | 1.45 | 4.35E-22 |
| 1 | RPL3 | 0.98 | 0.98 | 36.82 | 28.46 | 0.37 | 3.94E-21 |
| 1 | IRF1-AS1 | 0.33 | 0.15 | 1.49 | 0.49 | 1.62 | 4.12E-21 |
| 1 | MAF | 0.32 | 0.14 | 1.54 | 0.61 | 1.33 | 2.02E-20 |
| 1 | FKBP11 | 0.32 | 0.15 | 1.32 | 0.48 | 1.47 | 2.57E-20 |
| 1 | LITAF | 0.85 | 0.69 | 8.19 | 5.26 | 0.64 | 2.76E-20 |
| 1 | MCOLN2 | 0.25 | 0.10 | 1.18 | 0.38 | 1.62 | 5.09E-20 |
| 1 | PRKCB | 0.26 | 0.10 | 1.02 | 0.31 | 1.71 | 6.75E-20 |
| 1 | GNPTAB | 0.43 | 0.23 | 2.23 | 0.95 | 1.22 | 9.72E-20 |
| 1 | LPCAT1 | 0.28 | 0.12 | 1.03 | 0.39 | 1.40 | 1.44E-19 |
| 1 | SLFN12L | 0.46 | 0.26 | 2.38 | 1.09 | 1.13 | 1.77E-19 |
| 1 | CD247 | 0.76 | 0.63 | 7.04 | 4.20 | 0.75 | 2.51E-19 |
| 1 | IKZF3 | 0.54 | 0.33 | 3.13 | 1.84 | 0.76 | 1.76E-18 |
| 1 | STAT5B | 0.29 | 0.13 | 1.22 | 0.42 | 1.55 | 3.43E-18 |
| 1 | ARID5B | 0.55 | 0.36 | 3.42 | 1.90 | 0.85 | 6.06E-18 |
| 1 | PBX4 | 0.27 | 0.12 | 1.15 | 0.41 | 1.47 | 8.85E-18 |
| 1 | PCED1B-AS | 0.49 | 0.31 | 2.34 | 1.12 | 1.06 | 9.13E-18 |
| 1 | ERP29 | 0.48 | 0.31 | 2.00 | 1.00 | 1.00 | 1.44E-17 |
| 1 | LYAR | 0.36 | 0.18 | 1.48 | 0.67 | 1.13 | 2.77E-17 |
| 1 | ADRB2 | 0.30 | 0.14 | 1.21 | 0.56 | 1.12 | 7.06E-17 |

| | | | | | | | |
|---|---|---|---|---|---|---|---|
| 1 | MYL12B | 0.83 | 0.72 | 7.03 | 4.73 | 0.57 | 8.77E-17 |
| 1 | RAPGEF1 | 0.28 | 0.13 | 1.22 | 0.42 | 1.53 | 1.69E-16 |
| 1 | TLE5 | 0.64 | 0.47 | 3.22 | 1.99 | 0.70 | 3.75E-16 |
| 1 | SH3KBP1 | 0.42 | 0.26 | 1.81 | 0.89 | 1.03 | 1.15E-15 |
| 1 | IQGAP2 | 0.52 | 0.36 | 2.79 | 1.39 | 1.01 | 1.54E-15 |
| 1 | SLA | 0.57 | 0.40 | 3.38 | 1.98 | 0.78 | 2.40E-15 |
| 1 | SH3BGRL3 | 0.82 | 0.74 | 8.04 | 5.46 | 0.56 | 3.60E-15 |
| 1 | MGAT4A | 0.32 | 0.17 | 1.49 | 0.58 | 1.36 | 3.89E-15 |
| 1 | ARL6IP5 | 0.78 | 0.65 | 6.53 | 4.21 | 0.64 | 9.20E-15 |
| 1 | APMAP | 0.65 | 0.49 | 3.65 | 2.21 | 0.72 | 1.16E-14 |
| 1 | CAPN2 | 0.27 | 0.13 | 1.04 | 0.42 | 1.29 | 1.24E-14 |
| 1 | CALM1 | 0.93 | 0.88 | 15.19 | 10.80 | 0.49 | 1.26E-14 |
| 1 | LIMD2 | 0.58 | 0.41 | 2.75 | 1.63 | 0.75 | 2.67E-14 |
| 1 | EPG5 | 0.27 | 0.13 | 1.16 | 0.46 | 1.33 | 2.87E-14 |
| 1 | ARPC5 | 0.50 | 0.35 | 2.39 | 1.25 | 0.93 | 3.72E-14 |
| 1 | MYO1F | 0.32 | 0.17 | 1.36 | 0.65 | 1.07 | 7.39E-14 |
| 1 | SELPLG | 0.26 | 0.13 | 0.98 | 0.39 | 1.32 | 8.60E-14 |
| 1 | MAN1A1 | 0.29 | 0.15 | 1.06 | 0.48 | 1.14 | 8.88E-14 |
| 1 | CEP78 | 0.28 | 0.14 | 1.41 | 0.49 | 1.52 | 1.05E-13 |
| 1 | SAMD3 | 0.62 | 0.46 | 3.59 | 2.23 | 0.69 | 1.19E-13 |
| 1 | ITGAL | 0.42 | 0.26 | 1.92 | 0.97 | 0.98 | 1.98E-13 |
| 1 | TERF1 | 0.35 | 0.20 | 1.52 | 0.71 | 1.09 | 2.14E-13 |
| 1 | NFATC2 | 0.33 | 0.19 | 1.18 | 0.59 | 0.98 | 2.77E-13 |
| 1 | PFN1 | 0.87 | 0.84 | 10.42 | 7.41 | 0.49 | 3.13E-13 |
| 1 | CYFIP2 | 0.31 | 0.17 | 1.21 | 0.53 | 1.18 | 3.25E-13 |
| 1 | LBH | 0.42 | 0.26 | 2.02 | 1.11 | 0.87 | 3.85E-13 |
| 1 | GZMM | 0.58 | 0.42 | 3.30 | 2.03 | 0.70 | 6.01E-13 |
| 1 | TRBC1 | 0.52 | 0.38 | 3.31 | 1.80 | 0.88 | 7.86E-13 |
| 1 | HLA-DPB1 | 0.56 | 0.41 | 2.67 | 1.73 | 0.63 | 8.57E-13 |
| 1 | PDIA3 | 0.71 | 0.60 | 4.41 | 2.97 | 0.57 | 8.80E-13 |
| 1 | LCP1 | 0.86 | 0.77 | 8.80 | 6.57 | 0.42 | 9.00E-13 |
| 1 | ARRB2 | 0.30 | 0.16 | 1.13 | 0.50 | 1.18 | 1.01E-12 |
| 1 | HAVCR2 | 0.43 | 0.28 | 2.40 | 1.23 | 0.96 | 1.59E-12 |
| 1 | PPIB | 0.64 | 0.49 | 3.27 | 2.13 | 0.62 | 1.65E-12 |
| 1 | RASGRP1 | 0.33 | 0.19 | 1.61 | 0.66 | 1.30 | 1.75E-12 |
| 1 | TRG-AS1 | 0.46 | 0.31 | 2.35 | 1.32 | 0.84 | 2.14E-12 |
| 1 | RNF213 | 0.66 | 0.53 | 4.48 | 3.09 | 0.54 | 2.45E-12 |
| 1 | PTPRE | 0.34 | 0.20 | 1.47 | 0.70 | 1.07 | 3.22E-12 |
| 1 | RAP1B | 0.83 | 0.72 | 6.59 | 4.78 | 0.46 | 7.61E-12 |
| 1 | APLP2 | 0.42 | 0.28 | 1.81 | 1.00 | 0.85 | 8.86E-12 |
| 1 | ENSG00000 | 0.30 | 0.16 | 1.34 | 0.86 | 0.63 | 1.29E-11 |
| 1 | YBX1 | 0.84 | 0.78 | 7.92 | 5.91 | 0.42 | 1.83E-11 |
| 1 | LYN | 0.35 | 0.22 | 1.64 | 0.73 | 1.17 | 2.04E-11 |
| 1 | KLF3 | 0.45 | 0.31 | 2.17 | 1.20 | 0.85 | 2.46E-11 |
| 1 | SUB1 | 0.80 | 0.73 | 5.86 | 4.30 | 0.45 | 3.69E-11 |
| 1 | MXRA7 | 0.32 | 0.20 | 1.36 | 0.64 | 1.10 | 4.43E-11 |
| 1 | DGKD | 0.33 | 0.21 | 1.50 | 0.68 | 1.14 | 4.69E-11 |
| 1 | RASA3 | 0.25 | 0.14 | 0.96 | 0.41 | 1.23 | 5.55E-11 |
| 1 | ARNTL | 0.26 | 0.14 | 1.00 | 0.48 | 1.06 | 5.75E-11 |
| 1 | ARAP2 | 0.49 | 0.36 | 2.58 | 1.60 | 0.68 | 7.37E-11 |
| 1 | LCK | 0.40 | 0.28 | 1.66 | 0.92 | 0.85 | 8.64E-11 |
| 1 | TMEM50A | 0.55 | 0.44 | 2.70 | 1.68 | 0.68 | 9.25E-11 |
| 1 | GAPDH | 0.91 | 0.91 | 18.63 | 14.49 | 0.36 | 1.11E-10 |
| 1 | FLNA | 0.74 | 0.61 | 4.67 | 3.60 | 0.37 | 1.17E-10 |
| 1 | PDIA6 | 0.39 | 0.26 | 1.60 | 0.88 | 0.85 | 1.19E-10 |
| 1 | GIMAP7 | 0.37 | 0.23 | 1.66 | 0.98 | 0.76 | 1.71E-10 |
| 1 | DIAPH1 | 0.56 | 0.42 | 2.61 | 1.71 | 0.61 | 2.03E-10 |
| 1 | PGAM1 | 0.37 | 0.24 | 1.45 | 0.79 | 0.87 | 2.05E-10 |
| 1 | CD99 | 0.60 | 0.48 | 3.26 | 2.18 | 0.58 | 2.57E-10 |

| | | | | | | | |
|---|---|---|---|---|---|---|---|
| 1 | PTPN22 | 0.64 | 0.50 | 3.85 | 2.55 | 0.59 | 3.09E-10 |
| 1 | PTPRA | 0.29 | 0.17 | 1.05 | 0.59 | 0.83 | 3.13E-10 |
| 1 | MYO1G | 0.28 | 0.16 | 1.12 | 0.54 | 1.05 | 3.27E-10 |
| 1 | FYN | 0.79 | 0.69 | 6.45 | 4.68 | 0.46 | 3.82E-10 |
| 1 | RRBP1 | 0.49 | 0.37 | 2.30 | 1.42 | 0.70 | 5.20E-10 |
| 1 | AAK1 | 0.55 | 0.42 | 2.59 | 1.71 | 0.60 | 5.51E-10 |
| 1 | HMGB2 | 0.58 | 0.45 | 3.85 | 2.56 | 0.59 | 5.54E-10 |
| 1 | ARHGAP25 | 0.30 | 0.18 | 1.19 | 0.56 | 1.08 | 6.27E-10 |
| 1 | TGFBR2 | 0.43 | 0.30 | 1.95 | 1.14 | 0.77 | 7.17E-10 |
| 1 | PRR5L | 0.30 | 0.18 | 1.25 | 0.62 | 1.01 | 8.53E-10 |
| 1 | TBX21 | 0.34 | 0.22 | 1.45 | 0.79 | 0.88 | 1.12E-09 |
| 1 | LBR | 0.45 | 0.32 | 2.01 | 1.22 | 0.72 | 1.19E-09 |
| 1 | MYH9 | 0.66 | 0.55 | 3.78 | 2.71 | 0.48 | 1.60E-09 |
| 1 | GGA2 | 0.30 | 0.18 | 1.14 | 0.59 | 0.96 | 1.82E-09 |
| 1 | CD53 | 0.69 | 0.59 | 4.19 | 2.90 | 0.53 | 2.06E-09 |
| 1 | BCL7C | 0.26 | 0.15 | 0.94 | 0.47 | 1.00 | 2.06E-09 |
| 1 | DEK | 0.60 | 0.48 | 3.29 | 2.19 | 0.59 | 2.38E-09 |
| 1 | CEBPB | 0.36 | 0.23 | 1.51 | 1.01 | 0.58 | 2.74E-09 |
| 1 | GIMAP4 | 0.35 | 0.23 | 1.71 | 0.91 | 0.91 | 3.07E-09 |
| 1 | APBB1IP | 0.37 | 0.26 | 1.73 | 0.90 | 0.94 | 3.68E-09 |
| 1 | VTI1B | 0.25 | 0.15 | 0.98 | 0.42 | 1.20 | 3.88E-09 |
| 1 | STK38 | 0.32 | 0.20 | 1.22 | 0.64 | 0.94 | 4.72E-09 |
| 1 | METRNL | 0.80 | 0.71 | 9.54 | 7.39 | 0.37 | 5.58E-09 |
| 1 | PSAP | 0.32 | 0.20 | 1.12 | 0.60 | 0.90 | 6.12E-09 |
| 1 | EIF4G3 | 0.31 | 0.20 | 1.14 | 0.63 | 0.84 | 7.70E-09 |
| 1 | STK10 | 0.37 | 0.25 | 1.47 | 0.88 | 0.75 | 1.11E-08 |
| 1 | DUSP5 | 0.42 | 0.28 | 1.81 | 1.39 | 0.38 | 1.54E-08 |
| 1 | GLRX | 0.37 | 0.25 | 1.58 | 0.99 | 0.67 | 1.88E-08 |
| 1 | UPP1 | 0.40 | 0.29 | 1.76 | 1.12 | 0.65 | 2.14E-08 |
| 1 | ITK | 0.30 | 0.20 | 1.29 | 0.64 | 1.02 | 2.19E-08 |
| 1 | TNIP2 | 0.28 | 0.17 | 1.01 | 0.55 | 0.89 | 2.49E-08 |
| 1 | LAPTM5 | 0.52 | 0.41 | 2.21 | 1.50 | 0.56 | 2.76E-08 |
| 1 | DHRS7 | 0.39 | 0.28 | 1.67 | 0.99 | 0.75 | 3.33E-08 |
| 1 | UQCRB | 0.74 | 0.67 | 4.90 | 3.57 | 0.46 | 3.70E-08 |
| 1 | WNK1 | 0.45 | 0.34 | 1.99 | 1.27 | 0.64 | 4.46E-08 |
| 1 | RAB27A | 0.45 | 0.33 | 2.04 | 1.26 | 0.69 | 5.09E-08 |
| 1 | SYTL3 | 0.70 | 0.59 | 5.00 | 3.76 | 0.41 | 5.18E-08 |
| 1 | SH2D2A | 0.41 | 0.29 | 1.92 | 1.16 | 0.72 | 5.54E-08 |
| 1 | APOL6 | 0.40 | 0.29 | 1.83 | 1.08 | 0.76 | 5.59E-08 |
| 1 | GLIPR1 | 0.42 | 0.29 | 1.86 | 1.30 | 0.52 | 5.68E-08 |
| 1 | PSMB8 | 0.52 | 0.42 | 2.37 | 1.57 | 0.60 | 6.07E-08 |
| 1 | ADD3 | 0.26 | 0.16 | 1.02 | 0.53 | 0.95 | 6.17E-08 |
| 1 | TUT7 | 0.36 | 0.25 | 1.40 | 0.84 | 0.73 | 9.10E-08 |
| 1 | PITPNC1 | 0.38 | 0.27 | 1.59 | 1.02 | 0.64 | 1.02E-07 |
| 1 | CCDC88C | 0.36 | 0.25 | 1.56 | 0.90 | 0.79 | 1.14E-07 |
| 1 | SDCBP | 0.55 | 0.45 | 2.98 | 2.07 | 0.53 | 1.18E-07 |
| 1 | ATP1B3 | 0.67 | 0.58 | 6.98 | 5.34 | 0.39 | 1.33E-07 |
| 1 | EMP3 | 0.68 | 0.58 | 4.24 | 3.09 | 0.46 | 1.34E-07 |
| 1 | ENSG0000( | 0.36 | 0.25 | 1.48 | 0.91 | 0.71 | 1.40E-07 |
| 1 | SSR2 | 0.62 | 0.54 | 3.14 | 2.21 | 0.51 | 1.52E-07 |
| 1 | SLC9A3R1 | 0.37 | 0.25 | 1.44 | 0.97 | 0.58 | 2.13E-07 |
| 1 | SRPK2 | 0.28 | 0.18 | 1.05 | 0.56 | 0.92 | 2.23E-07 |
| 1 | OSBPL8 | 0.43 | 0.32 | 1.94 | 1.26 | 0.62 | 2.75E-07 |
| 1 | MORC3 | 0.38 | 0.27 | 1.43 | 0.90 | 0.67 | 2.90E-07 |
| 1 | HCLS1 | 0.41 | 0.30 | 1.61 | 1.04 | 0.63 | 2.94E-07 |
| 1 | TNFRSF14 | 0.27 | 0.17 | 1.03 | 0.54 | 0.94 | 3.00E-07 |
| 1 | IRF2 | 0.25 | 0.16 | 0.95 | 0.55 | 0.78 | 3.23E-07 |
| 1 | MYL12A | 0.76 | 0.67 | 6.09 | 4.67 | 0.38 | 4.10E-07 |
| 1 | CD47 | 0.36 | 0.26 | 1.54 | 0.88 | 0.80 | 4.13E-07 |

| | | | | | | | |
|---|---|---|---|---|---|---|---|
| 1 | SLA2 | 0.38 | 0.27 | 1.66 | 1.12 | 0.57 | 4.15E-07 |
| 1 | PTPN12 | 0.28 | 0.19 | 1.14 | 0.59 | 0.94 | 4.94E-07 |
| 1 | DIP2A | 0.30 | 0.21 | 1.44 | 0.76 | 0.91 | 5.07E-07 |
| 1 | SP140 | 0.27 | 0.17 | 0.92 | 0.55 | 0.74 | 5.55E-07 |
| 1 | TMEM59 | 0.53 | 0.44 | 2.41 | 1.60 | 0.59 | 5.93E-07 |
| 1 | RO60 | 0.31 | 0.22 | 1.28 | 0.70 | 0.87 | 6.03E-07 |
| 1 | SPCS2 | 0.46 | 0.35 | 1.92 | 1.27 | 0.60 | 6.26E-07 |
| 1 | ANKRD12 | 0.65 | 0.57 | 4.92 | 3.52 | 0.48 | 6.82E-07 |
| 1 | MOB2 | 0.29 | 0.19 | 1.05 | 0.58 | 0.84 | 7.15E-07 |
| 1 | STAT5A | 0.33 | 0.23 | 1.45 | 0.92 | 0.66 | 7.36E-07 |
| 1 | SPATA13 | 0.25 | 0.16 | 0.90 | 0.48 | 0.89 | 7.49E-07 |
| 1 | TMA7 | 0.77 | 0.70 | 5.42 | 4.18 | 0.38 | 8.80E-07 |
| 1 | CCNI | 0.73 | 0.67 | 4.88 | 3.69 | 0.40 | 9.41E-07 |
| 1 | CD3E | 0.31 | 0.21 | 1.40 | 0.80 | 0.80 | 9.65E-07 |
| 1 | UBL3 | 0.31 | 0.21 | 1.12 | 0.67 | 0.75 | 1.16E-06 |
| 1 | PIP4K2A | 0.53 | 0.45 | 2.74 | 1.78 | 0.62 | 1.33E-06 |
| 1 | LSM14A | 0.40 | 0.30 | 1.60 | 1.00 | 0.69 | 1.34E-06 |
| 1 | OPTN | 0.53 | 0.45 | 2.74 | 1.91 | 0.52 | 1.47E-06 |
| 1 | CMC1 | 0.59 | 0.50 | 5.13 | 3.62 | 0.51 | 1.49E-06 |
| 1 | NSD3 | 0.60 | 0.53 | 3.36 | 2.48 | 0.44 | 1.77E-06 |
| 1 | ZBTB1 | 0.49 | 0.40 | 2.22 | 1.55 | 0.52 | 1.80E-06 |
| 1 | TCF25 | 0.62 | 0.55 | 3.65 | 2.54 | 0.52 | 1.81E-06 |
| 1 | MANF | 0.33 | 0.24 | 1.35 | 0.77 | 0.81 | 1.84E-06 |
| 1 | RAP2B | 0.35 | 0.25 | 1.43 | 0.88 | 0.70 | 1.89E-06 |
| 1 | SEC11C | 0.27 | 0.18 | 1.04 | 0.60 | 0.81 | 1.90E-06 |
| 1 | ARHGEF3 | 0.29 | 0.20 | 1.15 | 0.70 | 0.72 | 2.11E-06 |
| 1 | SIRT2 | 0.33 | 0.23 | 1.28 | 0.79 | 0.69 | 2.20E-06 |
| 1 | PKM | 0.44 | 0.35 | 2.23 | 1.44 | 0.63 | 2.24E-06 |
| 1 | PTPN4 | 0.26 | 0.17 | 1.09 | 0.57 | 0.94 | 2.27E-06 |
| 1 | MBP | 0.68 | 0.59 | 4.61 | 3.57 | 0.37 | 2.32E-06 |
| 1 | ISCU | 0.37 | 0.27 | 1.47 | 0.91 | 0.69 | 2.42E-06 |
| 1 | PPP1R16B | 0.36 | 0.26 | 1.38 | 0.89 | 0.64 | 2.57E-06 |
| 1 | CMIP | 0.40 | 0.30 | 1.86 | 1.22 | 0.61 | 2.72E-06 |
| 1 | PSMA3 | 0.34 | 0.24 | 1.25 | 0.79 | 0.66 | 2.90E-06 |
| 1 | BLOC1S1 | 0.27 | 0.18 | 0.93 | 0.57 | 0.70 | 3.03E-06 |
| 1 | SLAMF7 | 0.40 | 0.30 | 1.81 | 1.22 | 0.57 | 3.06E-06 |
| 1 | AUTS2 | 0.29 | 0.20 | 1.25 | 0.75 | 0.74 | 3.18E-06 |
| 1 | MAPRE2 | 0.47 | 0.37 | 2.26 | 1.56 | 0.54 | 3.68E-06 |
| 1 | MTPN | 0.50 | 0.41 | 2.25 | 1.56 | 0.53 | 4.21E-06 |
| 1 | TRAPPC10 | 0.42 | 0.34 | 1.86 | 1.19 | 0.65 | 5.20E-06 |
| 1 | CCND3 | 0.34 | 0.24 | 1.34 | 0.85 | 0.66 | 5.30E-06 |
| 1 | PRMT2 | 0.37 | 0.28 | 1.69 | 1.02 | 0.72 | 5.39E-06 |
| 1 | HNRNPR | 0.53 | 0.46 | 2.56 | 1.77 | 0.53 | 5.86E-06 |
| 1 | ADAM10 | 0.27 | 0.19 | 1.03 | 0.57 | 0.85 | 6.19E-06 |
| 1 | ATP6V0E1 | 0.59 | 0.52 | 2.93 | 2.12 | 0.47 | 6.40E-06 |
| 1 | ANXA2 | 0.41 | 0.32 | 1.96 | 1.29 | 0.60 | 6.53E-06 |
| 1 | MGAT1 | 0.33 | 0.23 | 1.27 | 0.80 | 0.66 | 6.70E-06 |
| 1 | MYBL1 | 0.37 | 0.27 | 1.86 | 1.21 | 0.63 | 7.36E-06 |
| 1 | TRAM1 | 0.38 | 0.30 | 1.59 | 1.03 | 0.63 | 8.51E-06 |
| 1 | WASF2 | 0.43 | 0.35 | 1.85 | 1.22 | 0.60 | 9.46E-06 |
| 1 | C1orf35 | 0.30 | 0.21 | 1.18 | 0.73 | 0.70 | 1.02E-05 |
| 1 | RNF166 | 0.31 | 0.23 | 1.27 | 0.77 | 0.72 | 1.16E-05 |
| 1 | KLRF1 | 0.41 | 0.31 | 1.97 | 1.43 | 0.46 | 1.23E-05 |
| 1 | NAP1L4 | 0.52 | 0.44 | 2.56 | 1.86 | 0.46 | 1.36E-05 |
| 1 | RHOC | 0.31 | 0.22 | 1.10 | 0.69 | 0.66 | 1.64E-05 |
| 1 | NDFIP1 | 0.37 | 0.28 | 1.25 | 0.88 | 0.50 | 1.70E-05 |
| 1 | IL2RG | 0.63 | 0.56 | 3.34 | 2.50 | 0.42 | 1.78E-05 |
| 1 | ATXN1 | 0.37 | 0.29 | 1.70 | 1.09 | 0.64 | 1.80E-05 |
| 1 | PPDPF | 0.48 | 0.40 | 2.10 | 1.47 | 0.52 | 2.13E-05 |

| | | | | | | | |
|---|---|---|---|---|---|---|---|
| 1 | SPN | 0.38 | 0.29 | 1.59 | 1.07 | 0.57 | 2.25E-05 |
| 1 | OSTF1 | 0.43 | 0.35 | 1.85 | 1.27 | 0.54 | 2.25E-05 |
| 1 | CCNDBP1 | 0.31 | 0.23 | 1.17 | 0.69 | 0.77 | 2.37E-05 |
| 1 | ODC1 | 0.32 | 0.23 | 1.50 | 0.93 | 0.69 | 2.39E-05 |
| 1 | HMGN2 | 0.51 | 0.44 | 2.46 | 1.78 | 0.47 | 2.89E-05 |
| 1 | RNF168 | 0.42 | 0.33 | 1.75 | 1.24 | 0.50 | 3.02E-05 |
| 1 | SETX | 0.35 | 0.26 | 1.46 | 0.96 | 0.61 | 3.12E-05 |
| 1 | CDC42SE2 | 0.70 | 0.67 | 4.73 | 3.62 | 0.39 | 3.14E-05 |
| 1 | EBP | 0.31 | 0.23 | 1.23 | 0.83 | 0.57 | 3.47E-05 |
| 1 | SF3B2 | 0.45 | 0.38 | 1.82 | 1.27 | 0.53 | 3.72E-05 |
| 1 | ORMDL1 | 0.40 | 0.32 | 1.60 | 1.09 | 0.55 | 3.73E-05 |
| 1 | CIB1 | 0.50 | 0.42 | 2.22 | 1.62 | 0.46 | 3.79E-05 |
| 1 | STAU1 | 0.38 | 0.30 | 1.41 | 0.99 | 0.51 | 3.80E-05 |
| 1 | CAPZA1 | 0.51 | 0.43 | 2.12 | 1.59 | 0.41 | 3.85E-05 |
| 1 | CCSER2 | 0.54 | 0.46 | 2.61 | 2.00 | 0.38 | 3.91E-05 |
| 1 | ENSG00000 | 0.33 | 0.24 | 1.38 | 0.95 | 0.54 | 3.98E-05 |
| 1 | TOMM20 | 0.55 | 0.47 | 2.39 | 1.82 | 0.39 | 4.24E-05 |
| 1 | MECP2 | 0.44 | 0.37 | 2.04 | 1.33 | 0.61 | 4.40E-05 |
| 1 | N4BP2L2 | 0.64 | 0.57 | 3.93 | 2.80 | 0.49 | 4.53E-05 |
| 1 | CMPK1 | 0.29 | 0.21 | 1.04 | 0.64 | 0.69 | 4.55E-05 |
| 1 | PSMB9 | 0.43 | 0.35 | 1.96 | 1.33 | 0.56 | 4.60E-05 |
| 1 | DYNLT1 | 0.30 | 0.22 | 1.12 | 0.74 | 0.58 | 4.87E-05 |
| 1 | PLAC8 | 0.38 | 0.29 | 1.77 | 1.23 | 0.53 | 5.09E-05 |
| 1 | EIF4B | 0.52 | 0.47 | 2.44 | 1.79 | 0.44 | 5.37E-05 |
| 1 | ZNF292 | 0.40 | 0.31 | 1.84 | 1.27 | 0.53 | 5.60E-05 |
| 1 | ERBIN | 0.28 | 0.20 | 1.07 | 0.65 | 0.72 | 5.87E-05 |
| 1 | HLA-DPA1 | 0.47 | 0.38 | 2.14 | 1.63 | 0.40 | 6.20E-05 |
| 1 | LYST | 0.56 | 0.50 | 4.02 | 2.75 | 0.55 | 7.48E-05 |
| 1 | PRELID1 | 0.51 | 0.44 | 2.38 | 1.71 | 0.48 | 7.81E-05 |
| 1 | RASA2 | 0.42 | 0.35 | 1.90 | 1.28 | 0.57 | 8.76E-05 |
| 1 | UBE2R2 | 0.36 | 0.28 | 1.35 | 0.91 | 0.57 | 8.89E-05 |
| 1 | TECR | 0.32 | 0.24 | 1.22 | 0.83 | 0.56 | 9.16E-05 |
| 1 | BRD7 | 0.38 | 0.31 | 1.69 | 1.14 | 0.57 | 9.27E-05 |
| 1 | BTN3A2 | 0.37 | 0.29 | 1.53 | 1.03 | 0.56 | 9.80E-05 |
| 1 | WDR1 | 0.31 | 0.23 | 1.18 | 0.73 | 0.68 | 9.90E-05 |
| 1 | EPS15 | 0.27 | 0.20 | 0.95 | 0.57 | 0.74 | 0.00010693 |
| 1 | IKZF1 | 0.55 | 0.49 | 2.82 | 2.13 | 0.41 | 0.00010742 |
| 1 | CSGALNAC | 0.30 | 0.23 | 1.09 | 0.70 | 0.64 | 0.00011168 |
| 1 | PLAAT4 | 0.41 | 0.33 | 1.87 | 1.30 | 0.53 | 0.00011487 |
| 1 | SPOCK2 | 0.39 | 0.33 | 2.48 | 1.45 | 0.78 | 0.00011618 |
| 1 | MACF1 | 0.59 | 0.55 | 3.95 | 2.99 | 0.40 | 0.00011952 |
| 1 | PSME1 | 0.59 | 0.51 | 2.93 | 2.27 | 0.37 | 0.00012012 |
| 1 | ENSG00000 | 0.44 | 0.38 | 2.03 | 1.43 | 0.51 | 0.0001335 |
| 1 | UTRN | 0.49 | 0.43 | 2.69 | 1.87 | 0.53 | 0.00015015 |
| 1 | SLTM | 0.43 | 0.37 | 1.96 | 1.37 | 0.52 | 0.00015138 |
| 1 | PRKACB | 0.34 | 0.26 | 1.40 | 0.97 | 0.54 | 0.0001523 |
| 1 | CSNK1G2 | 0.40 | 0.33 | 1.75 | 1.20 | 0.54 | 0.00015367 |
| 1 | ADAR | 0.40 | 0.32 | 1.60 | 1.17 | 0.45 | 0.00015435 |
| 1 | GHITM | 0.42 | 0.34 | 1.63 | 1.21 | 0.43 | 0.00015481 |
| 1 | KLRC2 | 0.41 | 0.35 | 2.64 | 1.68 | 0.65 | 0.0001604 |
| 1 | TMEM160 | 0.26 | 0.19 | 0.94 | 0.61 | 0.62 | 0.00016284 |
| 1 | SMC3 | 0.39 | 0.32 | 1.70 | 1.20 | 0.51 | 0.00020393 |
| 1 | RFC1 | 0.32 | 0.26 | 1.35 | 0.85 | 0.67 | 0.00023674 |
| 1 | CCDC186 | 0.40 | 0.32 | 1.82 | 1.24 | 0.55 | 0.00023682 |
| 1 | RABGAP1L | 0.30 | 0.23 | 1.26 | 0.76 | 0.73 | 0.00027201 |
| 1 | SPEN | 0.43 | 0.37 | 2.19 | 1.49 | 0.56 | 0.00027289 |
| 1 | LNPEP | 0.37 | 0.31 | 1.71 | 1.15 | 0.57 | 0.00027292 |
| 1 | STAT3 | 0.59 | 0.54 | 3.39 | 2.61 | 0.38 | 0.00027306 |
| 1 | RIPOR2 | 0.36 | 0.30 | 1.97 | 1.22 | 0.70 | 0.00027686 |

| | | | | | | | |
|---|---|---|---|---|---|---|---|
| 1 | EID1 | 0.33 | 0.26 | 1.28 | 0.88 | 0.54 | 0.00028027 |
| 1 | MSL1 | 0.26 | 0.19 | 0.91 | 0.58 | 0.65 | 0.00029683 |
| 1 | SRSF9 | 0.50 | 0.45 | 2.25 | 1.73 | 0.38 | 0.00031647 |
| 1 | SRP9 | 0.31 | 0.25 | 1.16 | 0.78 | 0.58 | 0.00034183 |
| 1 | TMED9 | 0.28 | 0.22 | 1.10 | 0.68 | 0.71 | 0.00036091 |
| 1 | TXNRD1 | 0.32 | 0.25 | 1.45 | 0.97 | 0.58 | 0.00041471 |
| 1 | CLTA | 0.32 | 0.24 | 1.14 | 0.80 | 0.50 | 0.00041781 |
| 1 | BPTF | 0.45 | 0.40 | 2.31 | 1.72 | 0.42 | 0.0004487 |
| 1 | MVD | 0.29 | 0.22 | 1.03 | 0.75 | 0.46 | 0.00046199 |
| 1 | LINC-PINT | 0.59 | 0.54 | 3.51 | 2.73 | 0.36 | 0.00048982 |
| 1 | ENSG00000 | 0.40 | 0.34 | 2.10 | 1.42 | 0.57 | 0.00050313 |
| 1 | ARIH2 | 0.33 | 0.26 | 1.15 | 0.87 | 0.41 | 0.00053001 |
| 1 | NSMCE3 | 0.32 | 0.25 | 1.16 | 0.86 | 0.44 | 0.00056076 |
| 1 | GCC2 | 0.56 | 0.51 | 3.29 | 2.53 | 0.38 | 0.00058421 |
| 1 | SH3BGRL | 0.30 | 0.24 | 1.18 | 0.78 | 0.60 | 0.00060557 |
| 1 | CRBN | 0.30 | 0.23 | 1.21 | 0.76 | 0.66 | 0.00061465 |
| 1 | RPS27L | 0.41 | 0.35 | 1.88 | 1.31 | 0.52 | 0.00061544 |
| 1 | KMT2A | 0.46 | 0.40 | 2.05 | 1.60 | 0.36 | 0.00069906 |
| 1 | EIF3A | 0.52 | 0.47 | 2.51 | 1.95 | 0.37 | 0.00073418 |
| 1 | NFIL3 | 0.32 | 0.25 | 1.42 | 1.02 | 0.48 | 0.00073516 |
| 1 | PTPN7 | 0.37 | 0.30 | 1.53 | 1.09 | 0.49 | 0.00076054 |
| 1 | THRAP3 | 0.45 | 0.39 | 1.93 | 1.50 | 0.36 | 0.00076458 |
| 1 | LASP1 | 0.35 | 0.28 | 1.23 | 0.92 | 0.43 | 0.00079551 |
| 1 | CYTH1 | 0.46 | 0.41 | 2.33 | 1.74 | 0.43 | 0.0008159 |
| 1 | IGBP1 | 0.26 | 0.20 | 0.90 | 0.59 | 0.60 | 0.00081769 |
| 1 | TP53BP2 | 0.25 | 0.19 | 0.95 | 0.58 | 0.73 | 0.00084223 |
| 1 | RAC2 | 0.60 | 0.57 | 3.72 | 2.84 | 0.39 | 0.00084864 |
| 1 | CREBRF | 0.45 | 0.40 | 2.18 | 1.64 | 0.41 | 0.00091898 |
| 1 | LPIN2 | 0.26 | 0.20 | 1.00 | 0.69 | 0.54 | 0.00092231 |
| 1 | ARHGAP30 | 0.37 | 0.31 | 1.47 | 1.07 | 0.46 | 0.00096256 |
| 1 | CD300A | 0.45 | 0.38 | 2.42 | 1.82 | 0.41 | 0.00101143 |
| 1 | FYB1 | 0.36 | 0.30 | 1.77 | 1.25 | 0.50 | 0.00105403 |
| 1 | CDK2AP2 | 0.43 | 0.38 | 1.84 | 1.36 | 0.43 | 0.0010722 |
| 1 | ATP2B1 | 0.39 | 0.33 | 1.82 | 1.37 | 0.40 | 0.00120002 |
| 1 | CYRIB | 0.40 | 0.33 | 1.59 | 1.21 | 0.40 | 0.001216 |
| 1 | RAP1A | 0.41 | 0.37 | 1.81 | 1.24 | 0.55 | 0.00122792 |
| 1 | YPEL3 | 0.43 | 0.39 | 2.12 | 1.53 | 0.47 | 0.00123289 |
| 1 | RNF149 | 0.38 | 0.34 | 1.75 | 1.24 | 0.50 | 0.0013228 |
| 1 | FAM117A | 0.25 | 0.19 | 0.86 | 0.62 | 0.48 | 0.00135771 |
| 1 | SMARCA2 | 0.30 | 0.24 | 1.14 | 0.82 | 0.47 | 0.00137302 |
| 1 | ZC3H13 | 0.25 | 0.19 | 1.01 | 0.67 | 0.59 | 0.00137707 |
| 1 | PTPN1 | 0.26 | 0.20 | 1.03 | 0.61 | 0.76 | 0.00140424 |
| 1 | TAOK3 | 0.35 | 0.28 | 1.39 | 1.07 | 0.38 | 0.00142403 |
| 1 | HP1BP3 | 0.48 | 0.43 | 2.17 | 1.67 | 0.38 | 0.0014479 |
| 1 | SPCS3 | 0.39 | 0.33 | 1.65 | 1.18 | 0.49 | 0.00149539 |
| 1 | JADE2 | 0.27 | 0.21 | 1.02 | 0.68 | 0.59 | 0.00168434 |
| 1 | ATP5MK | 0.39 | 0.33 | 1.57 | 1.18 | 0.41 | 0.00177005 |
| 1 | UBE2K | 0.30 | 0.24 | 1.15 | 0.79 | 0.55 | 0.00180393 |
| 1 | TIAL1 | 0.35 | 0.30 | 1.61 | 1.03 | 0.65 | 0.00183819 |
| 1 | USP7 | 0.29 | 0.24 | 1.01 | 0.70 | 0.53 | 0.00191207 |
| 1 | ARL8A | 0.25 | 0.20 | 0.99 | 0.62 | 0.66 | 0.00195257 |
| 1 | EPB41 | 0.29 | 0.23 | 1.05 | 0.77 | 0.44 | 0.00198978 |
| 1 | NDUFB2 | 0.43 | 0.38 | 1.96 | 1.48 | 0.41 | 0.00202167 |
| 1 | UQCRFS1 | 0.32 | 0.26 | 1.14 | 0.83 | 0.46 | 0.0022053 |
| 1 | HERPUD2 | 0.29 | 0.23 | 1.15 | 0.80 | 0.53 | 0.00235153 |
| 1 | XRN1 | 0.35 | 0.30 | 1.38 | 1.02 | 0.44 | 0.00235359 |
| 1 | ATP1A1 | 0.42 | 0.37 | 1.95 | 1.48 | 0.40 | 0.00243385 |
| 1 | CREBBP | 0.30 | 0.25 | 1.19 | 0.78 | 0.62 | 0.00256062 |
| 1 | ST8SIA4 | 0.27 | 0.21 | 1.06 | 0.77 | 0.46 | 0.00281847 |

| | | | | | | | |
|---|---|---|---|---|---|---|---|
| 1 | XRN2 | 0.35 | 0.30 | 1.39 | 1.00 | 0.48 | 0.00287647 |
| 1 | PHF20L1 | 0.37 | 0.31 | 1.53 | 1.13 | 0.44 | 0.00289084 |
| 1 | TMEM258 | 0.35 | 0.29 | 1.37 | 1.03 | 0.41 | 0.0029457 |
| 1 | ANXA6 | 0.29 | 0.24 | 1.10 | 0.73 | 0.60 | 0.00312489 |
| 1 | TGOLN2 | 0.41 | 0.36 | 1.75 | 1.32 | 0.41 | 0.00334636 |
| 1 | SPSB3 | 0.33 | 0.28 | 1.31 | 0.96 | 0.45 | 0.00344732 |
| 1 | UQCR10 | 0.35 | 0.30 | 1.44 | 1.07 | 0.43 | 0.0036636 |
| 1 | KIF2A | 0.41 | 0.36 | 1.72 | 1.34 | 0.36 | 0.00391588 |
| 1 | KIAA1109 | 0.32 | 0.28 | 1.37 | 0.97 | 0.49 | 0.00398201 |
| 1 | RNF7 | 0.41 | 0.37 | 1.71 | 1.32 | 0.38 | 0.00405259 |
| 1 | ARPC4 | 0.29 | 0.24 | 1.01 | 0.76 | 0.41 | 0.00448107 |
| 1 | TRIM22 | 0.32 | 0.26 | 1.29 | 0.98 | 0.40 | 0.00454269 |
| 1 | ATP6V0B | 0.40 | 0.35 | 1.58 | 1.22 | 0.37 | 0.00459941 |
| 1 | DCP2 | 0.29 | 0.24 | 1.12 | 0.78 | 0.53 | 0.00462152 |
| 1 | FBXW7 | 0.27 | 0.22 | 1.09 | 0.69 | 0.65 | 0.00465303 |
| 1 | DGKZ | 0.37 | 0.32 | 1.52 | 1.15 | 0.41 | 0.00465955 |
| 1 | SRPK1 | 0.28 | 0.23 | 1.12 | 0.74 | 0.61 | 0.00471112 |
| 1 | BAX | 0.26 | 0.21 | 0.98 | 0.67 | 0.54 | 0.00500572 |
| 1 | P4HB | 0.34 | 0.28 | 1.33 | 0.95 | 0.48 | 0.00515277 |
| 1 | RAB14 | 0.37 | 0.32 | 1.43 | 1.08 | 0.40 | 0.00519948 |
| 1 | ARPC1B | 0.37 | 0.32 | 1.56 | 1.13 | 0.47 | 0.00547615 |
| 1 | RAB18 | 0.29 | 0.24 | 1.04 | 0.74 | 0.49 | 0.00582041 |
| 1 | RNF115 | 0.38 | 0.34 | 1.67 | 1.20 | 0.47 | 0.00584525 |
| 1 | ADNP | 0.26 | 0.21 | 0.84 | 0.63 | 0.41 | 0.00612887 |
| 1 | MRPS34 | 0.26 | 0.21 | 0.93 | 0.65 | 0.52 | 0.0061957 |
| 1 | PTBP3 | 0.32 | 0.27 | 1.20 | 0.91 | 0.40 | 0.00626365 |
| 1 | ATP5IF1 | 0.38 | 0.34 | 1.69 | 1.29 | 0.38 | 0.00671242 |
| 1 | GABARAPL | 0.39 | 0.35 | 1.57 | 1.14 | 0.47 | 0.00726552 |
| 1 | ISCA1 | 0.40 | 0.35 | 1.54 | 1.18 | 0.38 | 0.00733717 |
| 1 | RAB10 | 0.30 | 0.25 | 1.07 | 0.81 | 0.39 | 0.00733989 |
| 1 | ADD1 | 0.28 | 0.23 | 1.12 | 0.76 | 0.56 | 0.00774527 |
| 1 | ZBTB7A | 0.37 | 0.32 | 1.57 | 1.15 | 0.45 | 0.00812841 |
| 1 | CAP1 | 0.44 | 0.40 | 2.03 | 1.56 | 0.38 | 0.00822662 |
| 1 | RBM38 | 0.36 | 0.31 | 1.49 | 1.15 | 0.37 | 0.00832976 |
| 1 | NDUFA12 | 0.32 | 0.27 | 1.22 | 0.90 | 0.43 | 0.00856461 |
| 1 | CDK13 | 0.33 | 0.28 | 1.17 | 0.90 | 0.38 | 0.00895537 |
| 1 | FRG1 | 0.27 | 0.22 | 0.91 | 0.69 | 0.40 | 0.00912962 |
| 1 | POLR1D | 0.42 | 0.38 | 1.67 | 1.30 | 0.37 | 0.0092056 |
| 1 | YAF2 | 0.26 | 0.22 | 1.03 | 0.74 | 0.48 | 0.00980904 |
| 1 | KDELR2 | 0.25 | 0.21 | 0.88 | 0.61 | 0.52 | 0.0098433 |
| 1 | PSMB6 | 0.36 | 0.33 | 1.50 | 1.10 | 0.45 | 0.01010445 |
| 1 | COX14 | 0.32 | 0.27 | 1.14 | 0.85 | 0.42 | 0.0105363 |
| 1 | TMOD3 | 0.35 | 0.30 | 1.28 | 0.97 | 0.40 | 0.01058846 |
| 1 | KCTD20 | 0.26 | 0.21 | 0.95 | 0.68 | 0.49 | 0.01064416 |
| 1 | TBCA | 0.34 | 0.30 | 1.40 | 1.05 | 0.40 | 0.01098646 |
| 1 | TRPM7 | 0.26 | 0.21 | 0.89 | 0.65 | 0.46 | 0.01119904 |
| 1 | CAPZB | 0.37 | 0.33 | 1.49 | 1.11 | 0.42 | 0.01123709 |
| 1 | SNRNP200 | 0.29 | 0.24 | 1.11 | 0.78 | 0.52 | 0.01151479 |
| 1 | SYF2 | 0.53 | 0.52 | 2.78 | 2.14 | 0.38 | 0.01182973 |
| 1 | EIF2S3 | 0.37 | 0.34 | 1.52 | 1.15 | 0.40 | 0.0120087 |
| 1 | PSMB3 | 0.27 | 0.23 | 1.01 | 0.69 | 0.56 | 0.01223052 |
| 1 | TNFAIP8 | 0.26 | 0.22 | 1.16 | 0.86 | 0.44 | 0.01253144 |
| 1 | EIF3L | 0.28 | 0.24 | 0.99 | 0.73 | 0.45 | 0.0125859 |
| 1 | ATP2B4 | 0.25 | 0.21 | 1.06 | 0.74 | 0.53 | 0.01343483 |
| 1 | MDM4 | 0.30 | 0.26 | 1.24 | 0.93 | 0.42 | 0.01352462 |
| 1 | PDCD7 | 0.26 | 0.22 | 0.97 | 0.69 | 0.49 | 0.01360785 |
| 1 | SSH2 | 0.26 | 0.22 | 1.06 | 0.72 | 0.55 | 0.01408768 |
| 1 | ERP44 | 0.26 | 0.22 | 0.85 | 0.64 | 0.41 | 0.01418818 |
| 1 | NDUFA1 | 0.43 | 0.39 | 1.84 | 1.41 | 0.39 | 0.01471309 |

| | | | | | | | |
|---|---|---|---|---|---|---|---|
| 1 | MED10 | 0.34 | 0.30 | 1.34 | 0.98 | 0.46 | 0.01474221 |
| 1 | UHMK1 | 0.26 | 0.21 | 0.95 | 0.69 | 0.46 | 0.016979 |
| 1 | SAFB | 0.28 | 0.24 | 1.05 | 0.79 | 0.40 | 0.01729355 |
| 1 | AURKAIP1 | 0.37 | 0.33 | 1.37 | 1.07 | 0.36 | 0.01772174 |
| 1 | HSH2D | 0.28 | 0.23 | 1.21 | 0.92 | 0.40 | 0.01815482 |
| 1 | CXCL8 | 0.26 | 0.22 | 1.23 | 0.86 | 0.52 | 0.01947126 |
| 1 | SERPINB1 | 0.36 | 0.32 | 1.74 | 1.34 | 0.38 | 0.02110081 |
| 1 | PPP1CA | 0.27 | 0.23 | 0.97 | 0.74 | 0.39 | 0.02140138 |
| 1 | CSDE1 | 0.48 | 0.45 | 2.11 | 1.62 | 0.38 | 0.02170729 |
| 1 | RAB29 | 0.30 | 0.25 | 1.21 | 0.89 | 0.44 | 0.02253878 |
| 1 | MAP2K2 | 0.36 | 0.33 | 1.39 | 1.05 | 0.40 | 0.02319055 |
| 1 | CHD3 | 0.33 | 0.30 | 1.63 | 1.08 | 0.60 | 0.02320679 |
| 1 | PRDX5 | 0.31 | 0.27 | 1.28 | 0.93 | 0.45 | 0.02656727 |
| 1 | TRBC2 | 0.41 | 0.39 | 2.07 | 1.56 | 0.41 | 0.03058906 |
| 1 | ANXA5 | 0.25 | 0.21 | 1.06 | 0.78 | 0.44 | 0.03075889 |
| 1 | MAT2B | 0.26 | 0.23 | 0.97 | 0.74 | 0.40 | 0.03274525 |
| 1 | ESYT2 | 0.26 | 0.23 | 1.08 | 0.73 | 0.57 | 0.03375164 |
| 1 | GTF3A | 0.35 | 0.31 | 1.45 | 1.11 | 0.39 | 0.03444681 |
| 1 | HNRNPL | 0.33 | 0.30 | 1.28 | 0.97 | 0.41 | 0.03489004 |
| 1 | SAMD9 | 0.26 | 0.22 | 1.00 | 0.78 | 0.36 | 0.04058665 |
| 1 | BCL9L | 0.26 | 0.24 | 1.14 | 0.82 | 0.47 | 0.04159875 |
| 1 | DR1 | 0.32 | 0.28 | 1.20 | 0.94 | 0.36 | 0.04387401 |
| 1 | STARD3NL | 0.29 | 0.25 | 1.06 | 0.82 | 0.37 | 0.04523085 |
| 1 | GBP4 | 0.28 | 0.25 | 1.26 | 0.98 | 0.36 | 0.04970632 |
| 2 | DNAJB1 | 0.96 | 0.59 | 70.06 | 15.30 | 2.20 | 4.01E-109 |
| 2 | HSPA1B | 0.92 | 0.49 | 68.65 | 13.27 | 2.37 | 5.93E-107 |
| 2 | NR4A1 | 0.77 | 0.28 | 12.48 | 1.80 | 2.79 | 5.76E-103 |
| 2 | PPP1R15A | 0.93 | 0.61 | 22.15 | 5.56 | 1.99 | 1.92E-90 |
| 2 | HSPA1A | 0.98 | 0.80 | 200.73 | 56.46 | 1.83 | 6.09E-90 |
| 2 | FOSB | 0.64 | 0.26 | 7.40 | 1.32 | 2.48 | 3.71E-62 |
| 2 | FOS | 0.81 | 0.48 | 33.01 | 6.55 | 2.33 | 1.30E-61 |
| 2 | SERPINH1 | 0.45 | 0.12 | 4.02 | 0.65 | 2.62 | 9.97E-61 |
| 2 | HSPA6 | 0.66 | 0.31 | 42.36 | 8.63 | 2.29 | 1.20E-58 |
| 2 | HSPH1 | 0.92 | 0.68 | 34.93 | 12.34 | 1.50 | 1.39E-58 |
| 2 | HSP90AA1 | 0.99 | 0.96 | 240.15 | 107.52 | 1.16 | 1.92E-57 |
| 2 | JUN | 0.82 | 0.50 | 21.59 | 6.25 | 1.79 | 7.12E-57 |
| 2 | CD69 | 0.93 | 0.76 | 25.71 | 9.24 | 1.48 | 2.54E-55 |
| 2 | JUNB | 0.90 | 0.73 | 24.28 | 7.31 | 1.73 | 8.81E-51 |
| 2 | ZFP36 | 0.83 | 0.63 | 16.45 | 4.93 | 1.74 | 7.45E-49 |
| 2 | DUSP1 | 0.80 | 0.58 | 23.66 | 5.55 | 2.09 | 7.99E-49 |
| 2 | BAG3 | 0.41 | 0.12 | 2.74 | 0.46 | 2.56 | 2.00E-48 |
| 2 | DNAJA1 | 0.88 | 0.78 | 18.12 | 7.60 | 1.25 | 4.67E-44 |
| 2 | CLK1 | 0.86 | 0.64 | 9.71 | 3.90 | 1.32 | 1.14E-43 |
| 2 | HSPA8 | 0.95 | 0.89 | 37.40 | 17.45 | 1.10 | 5.49E-43 |
| 2 | ZFAND2A | 0.56 | 0.27 | 5.33 | 1.27 | 2.07 | 3.59E-40 |
| 2 | EGR1 | 0.27 | 0.06 | 1.57 | 0.22 | 2.86 | 2.20E-39 |
| 2 | UBC | 0.97 | 0.93 | 44.48 | 20.89 | 1.09 | 2.93E-38 |
| 2 | RGCC | 0.61 | 0.32 | 6.48 | 1.66 | 1.96 | 3.95E-38 |
| 2 | DEDD2 | 0.54 | 0.26 | 3.06 | 0.98 | 1.64 | 2.17E-37 |
| 2 | RGS1 | 0.58 | 0.32 | 8.91 | 1.89 | 2.24 | 7.00E-36 |
| 2 | KLF6 | 0.88 | 0.72 | 15.80 | 7.11 | 1.15 | 8.71E-35 |
| 2 | HSPE1 | 0.94 | 0.83 | 34.07 | 18.00 | 0.92 | 4.07E-34 |
| 2 | HSPB1 | 0.75 | 0.52 | 13.40 | 4.84 | 1.47 | 8.48E-33 |
| 2 | CACYBP | 0.76 | 0.57 | 10.58 | 4.52 | 1.23 | 1.57E-32 |
| 2 | DNAJB4 | 0.44 | 0.19 | 2.86 | 0.87 | 1.72 | 7.84E-30 |
| 2 | DUSP2 | 0.92 | 0.88 | 32.83 | 14.81 | 1.15 | 5.77E-29 |
| 2 | ATF3 | 0.37 | 0.15 | 2.69 | 0.69 | 1.97 | 6.33E-29 |
| 2 | JUND | 0.99 | 0.97 | 52.58 | 33.59 | 0.65 | 7.39E-27 |
| 2 | IER5 | 0.68 | 0.49 | 8.63 | 2.78 | 1.64 | 9.17E-27 |

| | | | | | | | |
|---|---|---|---|---|---|---|---|
| 2 | HSP90AB1 | 0.95 | 0.95 | 72.05 | 41.18 | 0.81 | 6.35E-26 |
| 2 | SLC2A3 | 0.68 | 0.48 | 6.84 | 2.78 | 1.30 | 9.13E-26 |
| 2 | TRA2B | 0.74 | 0.57 | 6.60 | 2.79 | 1.24 | 1.21E-25 |
| 2 | TNFSF14 | 0.57 | 0.35 | 4.98 | 1.60 | 1.64 | 2.04E-25 |
| 2 | AREG | 0.59 | 0.35 | 10.20 | 3.92 | 1.38 | 4.53E-25 |
| 2 | NFKBIA | 0.90 | 0.81 | 27.42 | 12.43 | 1.14 | 4.77E-25 |
| 2 | UBB | 0.94 | 0.85 | 23.03 | 12.05 | 0.93 | 5.06E-25 |
| 2 | HSPD1 | 0.93 | 0.88 | 58.44 | 29.11 | 1.01 | 1.86E-24 |
| 2 | RHOB | 0.30 | 0.11 | 1.54 | 0.47 | 1.72 | 2.53E-24 |
| 2 | MCL1 | 0.85 | 0.79 | 13.31 | 6.91 | 0.95 | 2.04E-23 |
| 2 | CCNL1 | 0.77 | 0.64 | 6.70 | 3.46 | 0.95 | 2.18E-23 |
| 2 | UBE2S | 0.62 | 0.44 | 6.01 | 2.11 | 1.51 | 2.70E-23 |
| 2 | BTG2 | 0.70 | 0.52 | 6.82 | 2.87 | 1.25 | 3.96E-23 |
| 2 | SPRY2 | 0.32 | 0.13 | 1.41 | 0.44 | 1.70 | 1.09E-22 |
| 2 | ZC3H12A | 0.53 | 0.32 | 3.24 | 1.21 | 1.42 | 1.94E-22 |
| 2 | IER5L | 0.36 | 0.17 | 3.39 | 0.75 | 2.19 | 1.30E-21 |
| 2 | TENT5A | 0.36 | 0.17 | 1.88 | 0.56 | 1.74 | 1.61E-21 |
| 2 | NKRF | 0.27 | 0.10 | 1.11 | 0.30 | 1.87 | 2.32E-21 |
| 2 | GADD45B | 0.76 | 0.62 | 20.37 | 6.92 | 1.56 | 3.04E-21 |
| 2 | SPRY1 | 0.37 | 0.17 | 2.99 | 0.83 | 1.84 | 4.64E-21 |
| 2 | GADD45G | 0.29 | 0.12 | 2.73 | 0.54 | 2.35 | 6.22E-20 |
| 2 | SRSF3 | 0.79 | 0.67 | 7.70 | 3.90 | 0.98 | 7.99E-20 |
| 2 | TSEN34 | 0.33 | 0.15 | 1.54 | 0.50 | 1.62 | 1.07E-18 |
| 2 | ARL5B | 0.37 | 0.19 | 2.34 | 0.68 | 1.78 | 1.27E-18 |
| 2 | RASGEF1B | 0.36 | 0.19 | 2.06 | 0.68 | 1.61 | 2.40E-17 |
| 2 | NEU1 | 0.42 | 0.24 | 2.73 | 0.95 | 1.53 | 7.38E-17 |
| 2 | YPEL5 | 0.66 | 0.53 | 5.13 | 2.35 | 1.13 | 1.24E-16 |
| 2 | IER2 | 0.79 | 0.74 | 13.92 | 6.09 | 1.19 | 1.26E-16 |
| 2 | NR4A2 | 0.71 | 0.55 | 8.05 | 3.92 | 1.04 | 1.81E-16 |
| 2 | MRPL18 | 0.47 | 0.28 | 2.88 | 1.20 | 1.27 | 2.00E-16 |
| 2 | ZBTB10 | 0.39 | 0.22 | 2.11 | 0.85 | 1.30 | 2.28E-16 |
| 2 | H3-3B | 0.98 | 0.98 | 33.71 | 21.48 | 0.65 | 2.91E-16 |
| 2 | MYLIP | 0.34 | 0.17 | 1.81 | 0.65 | 1.47 | 5.52E-16 |
| 2 | ANKRD37 | 0.30 | 0.14 | 1.88 | 0.55 | 1.77 | 7.24E-16 |
| 2 | MAFF | 0.43 | 0.26 | 2.40 | 0.98 | 1.29 | 1.97E-15 |
| 2 | TXNIP | 0.69 | 0.58 | 9.70 | 4.66 | 1.06 | 4.72E-15 |
| 2 | CHASERR | 0.75 | 0.63 | 5.85 | 3.41 | 0.78 | 5.08E-15 |
| 2 | CCT4 | 0.66 | 0.52 | 4.99 | 2.64 | 0.92 | 6.90E-15 |
| 2 | CKS2 | 0.46 | 0.28 | 2.45 | 1.19 | 1.04 | 7.01E-15 |
| 2 | RGS2 | 0.45 | 0.28 | 3.42 | 1.32 | 1.37 | 1.71E-14 |
| 2 | LMNA | 0.71 | 0.60 | 10.89 | 5.13 | 1.09 | 1.78E-14 |
| 2 | BTG1-DT | 0.34 | 0.18 | 2.07 | 0.73 | 1.50 | 2.65E-14 |
| 2 | TAGAP | 0.64 | 0.54 | 5.90 | 2.86 | 1.05 | 2.35E-13 |
| 2 | DNAJB6 | 0.79 | 0.73 | 9.14 | 5.60 | 0.71 | 3.62E-13 |
| 2 | BCAS2 | 0.48 | 0.32 | 2.89 | 1.25 | 1.21 | 4.04E-13 |
| 2 | ENSG00000 | 0.43 | 0.27 | 2.36 | 1.06 | 1.16 | 5.08E-13 |
| 2 | HNRNPU | 0.86 | 0.79 | 10.14 | 6.64 | 0.61 | 9.36E-13 |
| 2 | XCL1 | 0.77 | 0.56 | 28.98 | 17.77 | 0.71 | 1.13E-12 |
| 2 | TCP1 | 0.60 | 0.48 | 3.75 | 2.12 | 0.82 | 2.48E-12 |
| 2 | EIF4A2 | 0.80 | 0.74 | 7.17 | 4.79 | 0.58 | 4.05E-12 |
| 2 | CSRNP1 | 0.44 | 0.29 | 2.11 | 0.98 | 1.11 | 4.52E-12 |
| 2 | ZC3HAV1 | 0.66 | 0.53 | 5.32 | 3.12 | 0.77 | 1.10E-11 |
| 2 | CHORDC1 | 0.59 | 0.50 | 6.44 | 3.52 | 0.87 | 4.54E-11 |
| 2 | IFRD1 | 0.42 | 0.28 | 2.27 | 0.98 | 1.21 | 5.68E-11 |
| 2 | MYADM | 0.40 | 0.27 | 2.38 | 0.97 | 1.30 | 6.69E-11 |
| 2 | ZFP36L1 | 0.67 | 0.60 | 8.76 | 4.15 | 1.08 | 2.62E-10 |
| 2 | CHD2 | 0.70 | 0.58 | 5.28 | 3.35 | 0.66 | 3.00E-10 |
| 2 | SNHG8 | 0.66 | 0.57 | 4.97 | 2.97 | 0.74 | 3.72E-10 |
| 2 | MIR23AHG | 0.54 | 0.40 | 5.14 | 2.96 | 0.80 | 3.85E-10 |

| | | | | | | | |
|---|---|---|---|---|---|---|---|
| 2 | PHLDA1 | 0.41 | 0.28 | 2.55 | 1.13 | 1.17 | 4.36E-10 |
| 2 | TSPYL2 | 0.64 | 0.56 | 8.39 | 4.25 | 0.98 | 5.48E-10 |
| 2 | CD83 | 0.35 | 0.22 | 2.64 | 1.26 | 1.07 | 7.35E-10 |
| 2 | TAF7 | 0.64 | 0.53 | 3.91 | 2.45 | 0.67 | 7.98E-10 |
| 2 | INTS6 | 0.42 | 0.29 | 2.31 | 1.16 | 0.99 | 1.09E-09 |
| 2 | DDX3X | 0.77 | 0.71 | 7.83 | 4.84 | 0.69 | 1.28E-09 |
| 2 | GPBP1 | 0.85 | 0.82 | 9.44 | 6.91 | 0.45 | 1.44E-09 |
| 2 | SERTAD1 | 0.39 | 0.27 | 2.77 | 1.04 | 1.41 | 1.64E-09 |
| 2 | NFKBID | 0.40 | 0.26 | 2.19 | 1.15 | 0.92 | 2.75E-09 |
| 2 | AHSA1 | 0.59 | 0.47 | 3.41 | 2.25 | 0.60 | 2.76E-09 |
| 2 | PPP1R15B | 0.49 | 0.38 | 2.72 | 1.43 | 0.92 | 2.98E-09 |
| 2 | PPP1R2 | 0.61 | 0.52 | 5.17 | 2.58 | 1.00 | 3.54E-09 |
| 2 | MXD1 | 0.37 | 0.25 | 1.79 | 0.93 | 0.93 | 3.57E-09 |
| 2 | CITED2 | 0.34 | 0.21 | 1.99 | 0.91 | 1.13 | 4.14E-09 |
| 2 | SFPQ | 0.73 | 0.65 | 5.18 | 3.51 | 0.56 | 8.68E-09 |
| 2 | H2AX | 0.37 | 0.25 | 1.75 | 0.87 | 1.01 | 1.34E-08 |
| 2 | PPP1R10 | 0.41 | 0.30 | 2.71 | 1.18 | 1.19 | 1.45E-08 |
| 2 | TUBA1A | 0.56 | 0.48 | 4.94 | 2.27 | 1.12 | 1.85E-08 |
| 2 | DYNLL1 | 0.64 | 0.56 | 6.36 | 3.29 | 0.95 | 2.75E-08 |
| 2 | GATA3 | 0.35 | 0.24 | 1.66 | 0.83 | 1.00 | 2.98E-08 |
| 2 | RAB11FIP1 | 0.50 | 0.39 | 2.70 | 1.64 | 0.72 | 4.11E-08 |
| 2 | DOK2 | 0.61 | 0.51 | 4.62 | 2.95 | 0.65 | 5.10E-08 |
| 2 | NABP1 | 0.34 | 0.23 | 2.01 | 0.83 | 1.27 | 6.37E-08 |
| 2 | CGAS | 0.29 | 0.18 | 1.22 | 0.68 | 0.85 | 6.86E-08 |
| 2 | HMGCS1 | 0.27 | 0.16 | 1.32 | 0.57 | 1.20 | 8.56E-08 |
| 2 | PDE4B | 0.57 | 0.44 | 3.65 | 2.26 | 0.69 | 9.00E-08 |
| 2 | PRMT9 | 0.26 | 0.16 | 1.26 | 0.55 | 1.20 | 1.26E-07 |
| 2 | ABHD3 | 0.36 | 0.24 | 1.45 | 0.88 | 0.72 | 1.53E-07 |
| 2 | MIDEAS | 0.31 | 0.20 | 1.58 | 0.74 | 1.10 | 2.36E-07 |
| 2 | EIF4A3 | 0.37 | 0.28 | 2.11 | 0.93 | 1.17 | 4.07E-07 |
| 2 | ATP1B1 | 0.38 | 0.26 | 2.15 | 1.32 | 0.70 | 4.19E-07 |
| 2 | HEXIM1 | 0.35 | 0.25 | 1.77 | 0.90 | 0.97 | 4.42E-07 |
| 2 | ZNF331 | 0.60 | 0.51 | 5.76 | 3.54 | 0.70 | 4.77E-07 |
| 2 | PMAIP1 | 0.45 | 0.35 | 3.41 | 2.05 | 0.73 | 4.92E-07 |
| 2 | AHR | 0.34 | 0.23 | 1.66 | 0.85 | 0.97 | 5.44E-07 |
| 2 | AZIN1 | 0.39 | 0.28 | 1.58 | 0.94 | 0.74 | 5.98E-07 |
| 2 | CYCS | 0.61 | 0.53 | 4.14 | 2.61 | 0.66 | 9.60E-07 |
| 2 | AMD1 | 0.57 | 0.48 | 3.54 | 2.21 | 0.68 | 1.05E-06 |
| 2 | HSPA9 | 0.56 | 0.47 | 3.07 | 1.85 | 0.73 | 1.12E-06 |
| 2 | DUSP10 | 0.29 | 0.19 | 1.39 | 0.62 | 1.15 | 1.24E-06 |
| 2 | TSC22D3 | 0.72 | 0.66 | 10.35 | 6.59 | 0.65 | 1.47E-06 |
| 2 | TRA2A | 0.58 | 0.49 | 3.34 | 2.12 | 0.65 | 2.15E-06 |
| 2 | SBDS | 0.41 | 0.30 | 1.77 | 1.08 | 0.71 | 2.53E-06 |
| 2 | EIF5 | 0.68 | 0.66 | 5.55 | 3.65 | 0.60 | 2.77E-06 |
| 2 | PIK3IP1 | 0.27 | 0.17 | 1.01 | 0.56 | 0.86 | 4.15E-06 |
| 2 | FUS | 0.69 | 0.65 | 6.32 | 3.78 | 0.74 | 4.18E-06 |
| 2 | NOP58 | 0.45 | 0.36 | 2.26 | 1.32 | 0.77 | 5.56E-06 |
| 2 | MOB4 | 0.49 | 0.42 | 3.16 | 1.77 | 0.84 | 5.66E-06 |
| 2 | DDIT4 | 0.61 | 0.56 | 6.61 | 3.65 | 0.86 | 7.57E-06 |
| 2 | ZNF394 | 0.35 | 0.26 | 1.32 | 0.78 | 0.75 | 8.00E-06 |
| 2 | SLC38A2 | 0.63 | 0.58 | 4.70 | 3.15 | 0.58 | 1.21E-05 |
| 2 | REX1BD | 0.26 | 0.17 | 0.90 | 0.57 | 0.66 | 1.32E-05 |
| 2 | MIDN | 0.47 | 0.39 | 2.67 | 1.58 | 0.76 | 1.69E-05 |
| 2 | THAP9-AS1 | 0.32 | 0.23 | 1.37 | 0.76 | 0.85 | 1.79E-05 |
| 2 | NFKBIZ | 0.50 | 0.40 | 3.17 | 2.16 | 0.55 | 1.86E-05 |
| 2 | GADD45A | 0.35 | 0.27 | 2.80 | 1.20 | 1.22 | 2.01E-05 |
| 2 | IER3 | 0.31 | 0.22 | 2.72 | 1.02 | 1.42 | 2.11E-05 |
| 2 | NUDT4 | 0.27 | 0.18 | 1.11 | 0.61 | 0.86 | 3.40E-05 |
| 2 | JMJD6 | 0.44 | 0.37 | 2.34 | 1.36 | 0.79 | 3.41E-05 |

| | | | | | | | |
|---|---|---|---|---|---|---|---|
| 2 | TNFAIP3 | 0.80 | 0.78 | 12.55 | 8.01 | 0.65 | 3.76E-05 |
| 2 | PNPLA8 | 0.41 | 0.32 | 1.82 | 1.14 | 0.68 | 3.98E-05 |
| 2 | DUSP4 | 0.39 | 0.30 | 2.89 | 1.67 | 0.79 | 4.28E-05 |
| 2 | KAT6B | 0.30 | 0.21 | 1.12 | 0.72 | 0.65 | 4.31E-05 |
| 2 | IRF1 | 0.71 | 0.66 | 6.23 | 4.53 | 0.46 | 4.72E-05 |
| 2 | FKBP4 | 0.54 | 0.46 | 4.77 | 3.16 | 0.59 | 5.59E-05 |
| 2 | BRD2 | 0.68 | 0.67 | 6.95 | 3.87 | 0.85 | 5.78E-05 |
| 2 | PRR7 | 0.32 | 0.24 | 1.33 | 0.81 | 0.72 | 6.70E-05 |
| 2 | ERN1 | 0.37 | 0.28 | 2.10 | 1.48 | 0.51 | 7.18E-05 |
| 2 | NUFIP2 | 0.54 | 0.48 | 3.17 | 2.14 | 0.57 | 8.02E-05 |
| 2 | HES4 | 0.25 | 0.17 | 2.09 | 0.92 | 1.18 | 8.22E-05 |
| 2 | COTL1 | 0.48 | 0.39 | 2.99 | 2.21 | 0.44 | 8.37E-05 |
| 2 | ARL4A | 0.30 | 0.22 | 1.54 | 0.83 | 0.89 | 8.74E-05 |
| 2 | CDKN1A | 0.33 | 0.25 | 1.64 | 0.99 | 0.72 | 0.00012032 |
| 2 | MAP3K8 | 0.51 | 0.44 | 2.96 | 2.09 | 0.50 | 0.00012178 |
| 2 | NAMPT | 0.60 | 0.53 | 5.47 | 3.43 | 0.67 | 0.0001579 |
| 2 | CHMP1B | 0.35 | 0.28 | 1.79 | 1.01 | 0.82 | 0.00016684 |
| 2 | KLHL6 | 0.30 | 0.22 | 1.48 | 0.89 | 0.73 | 0.00017163 |
| 2 | ACAP1 | 0.53 | 0.46 | 2.85 | 1.98 | 0.52 | 0.00034697 |
| 2 | ENSG00000 | 0.27 | 0.20 | 1.16 | 0.80 | 0.54 | 0.00035819 |
| 2 | XCL2 | 0.79 | 0.70 | 33.20 | 25.26 | 0.39 | 0.0003693 |
| 2 | IFNG | 0.44 | 0.37 | 10.86 | 6.25 | 0.80 | 0.0004254 |
| 2 | SOCS1 | 0.34 | 0.27 | 1.75 | 1.07 | 0.71 | 0.00042889 |
| 2 | SIAH2 | 0.30 | 0.23 | 1.15 | 0.74 | 0.64 | 0.00044051 |
| 2 | EVL | 0.40 | 0.32 | 1.86 | 1.38 | 0.44 | 0.00063532 |
| 2 | BUD31 | 0.48 | 0.41 | 2.27 | 1.54 | 0.56 | 0.00078899 |
| 2 | TOPORS | 0.30 | 0.23 | 1.20 | 0.76 | 0.66 | 0.00079162 |
| 2 | YTHDC1 | 0.49 | 0.46 | 2.92 | 1.96 | 0.57 | 0.00093325 |
| 2 | ARRDC3 | 0.40 | 0.33 | 1.97 | 1.53 | 0.37 | 0.00099543 |
| 2 | PER1 | 0.32 | 0.26 | 1.46 | 0.87 | 0.75 | 0.00121235 |
| 2 | B3GNT2 | 0.35 | 0.29 | 1.58 | 0.95 | 0.73 | 0.00139369 |
| 2 | NBEAL1 | 0.26 | 0.19 | 0.91 | 0.62 | 0.57 | 0.00158119 |
| 2 | LINC01138 | 0.28 | 0.21 | 1.30 | 0.79 | 0.72 | 0.00203782 |
| 2 | FYB1 | 0.36 | 0.30 | 1.90 | 1.24 | 0.62 | 0.00214257 |
| 2 | TOB1 | 0.27 | 0.20 | 1.27 | 0.88 | 0.52 | 0.00243773 |
| 2 | PIM3 | 0.55 | 0.52 | 3.63 | 2.58 | 0.49 | 0.00294377 |
| 2 | CDKN1B | 0.51 | 0.47 | 2.84 | 2.01 | 0.50 | 0.00304296 |
| 2 | TAMALIN | 0.34 | 0.28 | 2.05 | 1.33 | 0.62 | 0.00323512 |
| 2 | NXT1 | 0.27 | 0.21 | 0.88 | 0.64 | 0.47 | 0.00325762 |
| 2 | TSC22D2 | 0.25 | 0.19 | 0.97 | 0.64 | 0.60 | 0.00327752 |
| 2 | COPA | 0.53 | 0.50 | 3.35 | 2.32 | 0.53 | 0.00382784 |
| 2 | AFF4 | 0.35 | 0.29 | 1.50 | 1.01 | 0.57 | 0.00395277 |
| 2 | DDIT3 | 0.27 | 0.21 | 1.21 | 0.78 | 0.63 | 0.00504361 |
| 2 | TIPARP | 0.46 | 0.41 | 2.66 | 1.78 | 0.58 | 0.0064962 |
| 2 | LUZP1 | 0.36 | 0.32 | 1.78 | 1.17 | 0.61 | 0.00709554 |
| 2 | BHLHE40 | 0.49 | 0.45 | 2.96 | 2.18 | 0.44 | 0.00710718 |
| 2 | RBBP6 | 0.36 | 0.30 | 1.45 | 1.05 | 0.46 | 0.00727427 |
| 2 | GPX1 | 0.34 | 0.28 | 1.50 | 0.95 | 0.66 | 0.0081507 |
| 2 | SPTY2D1 | 0.40 | 0.35 | 2.00 | 1.31 | 0.61 | 0.00900135 |
| 2 | CEBPD | 0.34 | 0.29 | 2.87 | 1.70 | 0.75 | 0.00994878 |
| 2 | BEX4 | 0.25 | 0.20 | 0.94 | 0.64 | 0.54 | 0.01033171 |
| 2 | ENSG00000 | 0.32 | 0.26 | 1.32 | 0.95 | 0.48 | 0.0106645 |
| 2 | SAT1 | 0.82 | 0.84 | 14.37 | 9.87 | 0.54 | 0.01364193 |
| 2 | KDM2A | 0.41 | 0.36 | 1.68 | 1.23 | 0.44 | 0.01525502 |
| 2 | EIF4A1 | 0.27 | 0.22 | 1.04 | 0.68 | 0.62 | 0.01533483 |
| 2 | RPL22L1 | 0.47 | 0.43 | 2.47 | 1.89 | 0.38 | 0.01613334 |
| 2 | RSRC2 | 0.59 | 0.61 | 4.16 | 3.06 | 0.44 | 0.01756043 |
| 2 | ZFAND5 | 0.41 | 0.39 | 2.20 | 1.46 | 0.59 | 0.020316 |
| 2 | VMP1 | 0.44 | 0.38 | 2.30 | 1.75 | 0.40 | 0.02067312 |

| | | | | | | | |
|---|---|---|---|---|---|---|---|
| 2 | HECA | 0.39 | 0.35 | 1.74 | 1.28 | 0.45 | 0.02105793 |
| 2 | PAF1 | 0.25 | 0.20 | 0.90 | 0.63 | 0.53 | 0.02113271 |
| 2 | CCDC59 | 0.32 | 0.26 | 1.22 | 0.91 | 0.42 | 0.02238155 |
| 2 | SRSF7 | 0.78 | 0.81 | 9.33 | 6.81 | 0.46 | 0.02322384 |
| 2 | CDKN2D | 0.41 | 0.36 | 2.11 | 1.46 | 0.54 | 0.02327651 |
| 2 | STIP1 | 0.46 | 0.43 | 2.36 | 1.77 | 0.42 | 0.02374072 |
| 2 | BCL2 | 0.29 | 0.24 | 1.35 | 0.88 | 0.62 | 0.0242222 |
| 2 | TRMT10C | 0.28 | 0.24 | 1.02 | 0.75 | 0.43 | 0.02606188 |
| 2 | ZNF326 | 0.25 | 0.21 | 0.91 | 0.68 | 0.42 | 0.02741796 |
| 2 | HERPUD1 | 0.47 | 0.44 | 2.83 | 2.00 | 0.50 | 0.03448755 |
| 2 | GLA | 0.35 | 0.31 | 2.04 | 1.33 | 0.62 | 0.03600398 |
| 2 | SLC39A10 | 0.31 | 0.27 | 1.37 | 1.06 | 0.37 | 0.04173611 |
| 2 | NECAP2 | 0.31 | 0.27 | 1.16 | 0.88 | 0.40 | 0.04469873 |
| 3 | DNAJB1 | 0.93 | 0.60 | 57.37 | 18.90 | 1.60 | 1.43E-78 |
| 3 | HSPA1B | 0.87 | 0.51 | 49.96 | 17.98 | 1.47 | 1.81E-66 |
| 3 | FGFBP2 | 0.61 | 0.18 | 5.27 | 1.50 | 1.81 | 3.48E-65 |
| 3 | HSPA1A | 0.95 | 0.81 | 171.38 | 65.19 | 1.39 | 2.55E-60 |
| 3 | CCL4L2 | 0.59 | 0.21 | 20.80 | 4.42 | 2.23 | 4.72E-59 |
| 3 | CACYBP | 0.86 | 0.56 | 11.40 | 4.51 | 1.34 | 7.62E-56 |
| 3 | UBC | 0.99 | 0.93 | 47.91 | 20.81 | 1.20 | 1.79E-54 |
| 3 | IFNG | 0.69 | 0.33 | 21.19 | 4.47 | 2.24 | 1.39E-53 |
| 3 | CCL4 | 0.87 | 0.49 | 51.37 | 17.95 | 1.52 | 5.89E-53 |
| 3 | HSPA8 | 0.97 | 0.89 | 38.86 | 17.65 | 1.14 | 7.81E-53 |
| 3 | HSP90AA1 | 0.99 | 0.96 | 251.67 | 108.50 | 1.21 | 3.42E-52 |
| 3 | GZMH | 0.73 | 0.32 | 6.39 | 2.51 | 1.35 | 1.10E-48 |
| 3 | DNAJA1 | 0.93 | 0.77 | 17.48 | 7.96 | 1.13 | 3.06E-48 |
| 3 | HSPA6 | 0.66 | 0.32 | 36.47 | 10.50 | 1.80 | 3.26E-47 |
| 3 | FCGR3A | 0.67 | 0.29 | 4.82 | 1.91 | 1.34 | 4.05E-46 |
| 3 | UBB | 0.96 | 0.85 | 23.84 | 12.16 | 0.97 | 1.25E-43 |
| 3 | NKG7 | 0.99 | 0.91 | 38.21 | 20.55 | 0.89 | 2.70E-43 |
| 3 | CCL3 | 0.64 | 0.29 | 19.24 | 5.56 | 1.79 | 1.75E-41 |
| 3 | SPON2 | 0.38 | 0.11 | 2.28 | 0.58 | 1.99 | 1.11E-38 |
| 3 | JUN | 0.80 | 0.51 | 17.49 | 7.35 | 1.25 | 1.04E-37 |
| 3 | ACTB | 0.94 | 0.84 | 20.77 | 9.60 | 1.11 | 1.29E-35 |
| 3 | UBE2S | 0.73 | 0.43 | 5.01 | 2.38 | 1.07 | 5.85E-35 |
| 3 | ENSG00000 | 0.41 | 0.14 | 2.50 | 0.67 | 1.90 | 8.43E-35 |
| 3 | CCL5 | 0.97 | 0.79 | 32.40 | 19.67 | 0.72 | 9.49E-35 |
| 3 | S100A4 | 0.84 | 0.61 | 10.06 | 4.86 | 1.05 | 1.04E-32 |
| 3 | HSPH1 | 0.87 | 0.69 | 27.92 | 14.15 | 0.98 | 2.11E-32 |
| 3 | TXNIP | 0.81 | 0.56 | 10.42 | 4.65 | 1.16 | 3.73E-32 |
| 3 | DEDD2 | 0.56 | 0.26 | 2.67 | 1.10 | 1.28 | 3.80E-31 |
| 3 | TNF | 0.33 | 0.11 | 2.21 | 0.55 | 2.00 | 3.14E-29 |
| 3 | SYNE1 | 0.52 | 0.24 | 2.55 | 1.06 | 1.27 | 3.60E-29 |
| 3 | GZMA | 0.71 | 0.41 | 8.59 | 5.29 | 0.70 | 7.09E-29 |
| 3 | PRDM1 | 0.51 | 0.23 | 2.93 | 1.14 | 1.36 | 1.03E-28 |
| 3 | CTSC | 0.67 | 0.40 | 4.53 | 2.07 | 1.13 | 1.95E-27 |
| 3 | BAG3 | 0.37 | 0.13 | 1.64 | 0.72 | 1.19 | 3.80E-27 |
| 3 | LDLR | 0.34 | 0.12 | 1.46 | 0.47 | 1.64 | 2.56E-26 |
| 3 | APOL6 | 0.54 | 0.27 | 2.31 | 1.03 | 1.16 | 3.23E-25 |
| 3 | SYNE2 | 0.69 | 0.39 | 4.44 | 2.62 | 0.76 | 7.99E-25 |
| 3 | RGS2 | 0.53 | 0.27 | 3.08 | 1.43 | 1.10 | 1.89E-24 |
| 3 | ENSG00000 | 0.28 | 0.09 | 1.13 | 0.45 | 1.33 | 2.11E-24 |
| 3 | KLF2 | 0.65 | 0.38 | 6.40 | 2.83 | 1.18 | 2.47E-24 |
| 3 | CGAS | 0.38 | 0.16 | 1.85 | 0.58 | 1.69 | 6.91E-24 |
| 3 | PPP1R15A | 0.83 | 0.64 | 13.86 | 7.46 | 0.89 | 7.79E-24 |
| 3 | ZFAND2A | 0.53 | 0.28 | 3.51 | 1.70 | 1.05 | 8.52E-24 |
| 3 | HSPE1 | 0.92 | 0.84 | 32.37 | 18.69 | 0.79 | 1.52E-23 |
| 3 | C12orf75 | 0.49 | 0.24 | 2.19 | 1.00 | 1.13 | 9.91E-23 |
| 3 | PMAIP1 | 0.56 | 0.33 | 4.94 | 1.80 | 1.46 | 2.05E-22 |

| | | | | | | | |
|---|---|---|---|---|---|---|---|
| 3 | TMSB10 | 0.99 | 0.96 | 29.84 | 20.53 | 0.54 | 8.51E-22 |
| 3 | HCST | 0.86 | 0.70 | 8.59 | 4.94 | 0.80 | 8.63E-22 |
| 3 | MRPL18 | 0.50 | 0.28 | 2.81 | 1.25 | 1.16 | 4.76E-21 |
| 3 | HSP90AB1 | 0.99 | 0.95 | 69.88 | 42.30 | 0.72 | 1.34E-20 |
| 3 | TOB1 | 0.39 | 0.18 | 2.12 | 0.73 | 1.53 | 1.97E-20 |
| 3 | GZMM | 0.66 | 0.41 | 3.59 | 2.04 | 0.81 | 2.12E-20 |
| 3 | BIN2 | 0.51 | 0.28 | 2.07 | 1.05 | 0.98 | 6.05E-20 |
| 3 | UAP1 | 0.33 | 0.14 | 1.28 | 0.47 | 1.45 | 7.36E-20 |
| 3 | DNAJB6 | 0.87 | 0.72 | 8.97 | 5.72 | 0.65 | 2.08E-19 |
| 3 | CCL3L1 | 0.28 | 0.10 | 2.20 | 0.90 | 1.29 | 4.06E-19 |
| 3 | DOK2 | 0.70 | 0.50 | 5.36 | 2.85 | 0.91 | 4.89E-19 |
| 3 | PCED1B-AS | 0.54 | 0.31 | 2.37 | 1.17 | 1.01 | 1.06E-18 |
| 3 | ITGB2 | 0.56 | 0.33 | 2.99 | 1.48 | 1.01 | 1.08E-18 |
| 3 | ATF3 | 0.35 | 0.15 | 2.09 | 0.84 | 1.31 | 1.67E-18 |
| 3 | SH3BGRL3 | 0.88 | 0.74 | 8.61 | 5.48 | 0.65 | 2.49E-18 |
| 3 | NEU1 | 0.45 | 0.24 | 2.35 | 1.06 | 1.15 | 6.12E-18 |
| 3 | IRF1 | 0.81 | 0.64 | 7.31 | 4.38 | 0.74 | 8.17E-18 |
| 3 | HSPB1 | 0.75 | 0.53 | 9.14 | 5.82 | 0.65 | 1.14E-17 |
| 3 | SLC20A1 | 0.42 | 0.22 | 2.03 | 0.77 | 1.40 | 1.30E-17 |
| 3 | CALM1 | 0.96 | 0.88 | 15.25 | 11.00 | 0.47 | 2.96E-17 |
| 3 | PFN1 | 0.93 | 0.83 | 11.52 | 7.35 | 0.65 | 3.96E-17 |
| 3 | CKS2 | 0.48 | 0.28 | 2.81 | 1.15 | 1.28 | 7.28E-17 |
| 3 | CDC42EP3 | 0.46 | 0.26 | 2.27 | 1.01 | 1.16 | 1.20E-16 |
| 3 | H3-3B | 0.99 | 0.97 | 30.79 | 22.30 | 0.47 | 1.60E-16 |
| 3 | ADGRG1 | 0.28 | 0.11 | 0.94 | 0.44 | 1.12 | 5.18E-16 |
| 3 | PYHIN1 | 0.45 | 0.23 | 2.05 | 1.13 | 0.86 | 6.13E-16 |
| 3 | DNAJB4 | 0.40 | 0.21 | 2.23 | 1.03 | 1.12 | 6.45E-16 |
| 3 | CD69 | 0.89 | 0.77 | 17.86 | 11.06 | 0.69 | 1.20E-15 |
| 3 | PLEK | 0.62 | 0.41 | 4.03 | 2.40 | 0.75 | 5.09E-15 |
| 3 | LYAR | 0.37 | 0.19 | 1.50 | 0.71 | 1.08 | 6.58E-15 |
| 3 | ERN1 | 0.46 | 0.27 | 3.07 | 1.31 | 1.22 | 1.10E-14 |
| 3 | GBP5 | 0.46 | 0.26 | 2.24 | 1.28 | 0.81 | 2.55E-14 |
| 3 | RNF213 | 0.71 | 0.53 | 4.71 | 3.11 | 0.60 | 3.83E-14 |
| 3 | SERPINH1 | 0.32 | 0.15 | 2.23 | 1.06 | 1.08 | 5.02E-14 |
| 3 | IER5L | 0.35 | 0.17 | 1.88 | 1.08 | 0.79 | 5.03E-14 |
| 3 | KLF6 | 0.84 | 0.73 | 12.92 | 7.84 | 0.72 | 5.48E-14 |
| 3 | ZC3HAV1 | 0.70 | 0.53 | 5.21 | 3.19 | 0.71 | 5.84E-14 |
| 3 | AHSA1 | 0.64 | 0.46 | 3.88 | 2.19 | 0.83 | 7.78E-14 |
| 3 | IDI1 | 0.58 | 0.42 | 3.26 | 1.74 | 0.90 | 2.29E-13 |
| 3 | JUNB | 0.83 | 0.75 | 14.15 | 9.56 | 0.57 | 2.36E-13 |
| 3 | ANXA1 | 0.83 | 0.70 | 17.27 | 10.17 | 0.76 | 2.58E-13 |
| 3 | CHORDC1 | 0.64 | 0.49 | 6.55 | 3.57 | 0.88 | 2.80E-13 |
| 3 | ENC1 | 0.25 | 0.11 | 1.26 | 0.46 | 1.46 | 4.62E-13 |
| 3 | GADD45B | 0.76 | 0.62 | 13.75 | 8.44 | 0.70 | 6.58E-13 |
| 3 | GNG2 | 0.66 | 0.44 | 4.43 | 2.97 | 0.58 | 6.74E-13 |
| 3 | CYBA | 0.90 | 0.75 | 9.68 | 6.75 | 0.52 | 9.35E-13 |
| 3 | KMT2E-AS1 | 0.30 | 0.14 | 1.19 | 0.61 | 0.96 | 1.65E-12 |
| 3 | ADRB2 | 0.30 | 0.15 | 1.20 | 0.59 | 1.03 | 1.73E-12 |
| 3 | BCL11B | 0.37 | 0.19 | 1.50 | 0.80 | 0.90 | 1.87E-12 |
| 3 | DYNLL1 | 0.72 | 0.55 | 5.29 | 3.56 | 0.57 | 2.71E-12 |
| 3 | SRSF3 | 0.81 | 0.67 | 6.07 | 4.29 | 0.50 | 3.75E-12 |
| 3 | NEDD9 | 0.28 | 0.13 | 1.06 | 0.50 | 1.10 | 4.57E-12 |
| 3 | RIPK2 | 0.26 | 0.12 | 0.86 | 0.40 | 1.13 | 4.69E-12 |
| 3 | MYLIP | 0.33 | 0.18 | 1.61 | 0.72 | 1.17 | 5.94E-12 |
| 3 | MYBL1 | 0.45 | 0.26 | 2.06 | 1.20 | 0.77 | 6.54E-12 |
| 3 | AMD1 | 0.62 | 0.47 | 3.89 | 2.18 | 0.84 | 7.54E-12 |
| 3 | TLE5 | 0.65 | 0.48 | 3.23 | 2.05 | 0.66 | 8.49E-12 |
| 3 | NCR3 | 0.25 | 0.12 | 0.94 | 0.39 | 1.27 | 1.09E-11 |
| 3 | MXD1 | 0.42 | 0.24 | 1.60 | 0.99 | 0.69 | 1.74E-11 |

| | | | | | | | |
|---|---|---|---|---|---|---|---|
| 3 | ARPC5L | 0.64 | 0.48 | 3.24 | 2.05 | 0.66 | 1.93E-11 |
| 3 | GNPTAB | 0.41 | 0.24 | 1.86 | 1.08 | 0.78 | 2.12E-11 |
| 3 | ARRDC3 | 0.48 | 0.31 | 2.83 | 1.38 | 1.04 | 2.36E-11 |
| 3 | CCT4 | 0.68 | 0.52 | 4.35 | 2.81 | 0.63 | 3.04E-11 |
| 3 | TBC1D10C | 0.31 | 0.15 | 1.08 | 0.61 | 0.83 | 3.08E-11 |
| 3 | REX1BD | 0.31 | 0.16 | 1.12 | 0.54 | 1.05 | 3.72E-11 |
| 3 | TRBC1 | 0.55 | 0.38 | 3.24 | 1.88 | 0.79 | 3.75E-11 |
| 3 | CLK1 | 0.80 | 0.66 | 6.52 | 4.62 | 0.50 | 4.53E-11 |
| 3 | PPP1R18 | 0.46 | 0.31 | 2.03 | 1.09 | 0.90 | 5.77E-11 |
| 3 | TCP1 | 0.62 | 0.48 | 3.44 | 2.21 | 0.64 | 5.95E-11 |
| 3 | DNAJA4 | 0.29 | 0.16 | 1.73 | 0.72 | 1.26 | 7.78E-11 |
| 3 | NR4A2 | 0.69 | 0.55 | 6.53 | 4.29 | 0.61 | 1.13E-10 |
| 3 | SBDS | 0.46 | 0.30 | 1.90 | 1.07 | 0.82 | 1.29E-10 |
| 3 | S100A6 | 0.83 | 0.67 | 7.38 | 5.70 | 0.37 | 1.77E-10 |
| 3 | ACTG1 | 0.93 | 0.87 | 14.80 | 10.85 | 0.45 | 2.81E-10 |
| 3 | DDIT3 | 0.34 | 0.20 | 1.51 | 0.74 | 1.04 | 3.94E-10 |
| 3 | TRBC2 | 0.54 | 0.36 | 2.36 | 1.53 | 0.63 | 4.33E-10 |
| 3 | AZIN1 | 0.44 | 0.28 | 1.64 | 0.95 | 0.80 | 4.44E-10 |
| 3 | DUSP5 | 0.44 | 0.29 | 2.29 | 1.32 | 0.80 | 5.31E-10 |
| 3 | CYCS | 0.67 | 0.52 | 3.97 | 2.68 | 0.57 | 5.90E-10 |
| 3 | MAF | 0.29 | 0.15 | 1.20 | 0.72 | 0.74 | 6.00E-10 |
| 3 | CYTOR | 0.47 | 0.31 | 2.01 | 1.27 | 0.66 | 7.45E-10 |
| 3 | ITGAL | 0.42 | 0.27 | 1.74 | 1.05 | 0.73 | 7.49E-10 |
| 3 | EFHD2 | 0.71 | 0.57 | 4.89 | 3.32 | 0.56 | 8.13E-10 |
| 3 | GP6-AS1 | 0.27 | 0.14 | 0.99 | 0.47 | 1.06 | 8.64E-10 |
| 3 | HSPD1 | 0.92 | 0.88 | 44.18 | 32.41 | 0.45 | 1.06E-09 |
| 3 | MYL12A | 0.75 | 0.67 | 7.47 | 4.49 | 0.73 | 1.44E-09 |
| 3 | UPP1 | 0.44 | 0.29 | 1.92 | 1.12 | 0.77 | 1.48E-09 |
| 3 | TRA2B | 0.71 | 0.58 | 4.41 | 3.27 | 0.43 | 2.14E-09 |
| 3 | UCP2 | 0.30 | 0.17 | 1.04 | 0.55 | 0.91 | 2.43E-09 |
| 3 | CEP78 | 0.28 | 0.15 | 1.08 | 0.59 | 0.86 | 2.96E-09 |
| 3 | NFIL3 | 0.38 | 0.24 | 1.90 | 0.96 | 0.99 | 2.99E-09 |
| 3 | LGALS1 | 0.81 | 0.66 | 8.99 | 6.74 | 0.42 | 3.47E-09 |
| 3 | CFL1 | 0.91 | 0.83 | 9.71 | 7.43 | 0.39 | 4.40E-09 |
| 3 | CCND3 | 0.39 | 0.24 | 1.49 | 0.85 | 0.81 | 4.73E-09 |
| 3 | PTPN4 | 0.30 | 0.17 | 1.05 | 0.60 | 0.81 | 5.02E-09 |
| 3 | TNFSF9 | 0.26 | 0.14 | 1.20 | 0.68 | 0.81 | 5.68E-09 |
| 3 | CARD16 | 0.38 | 0.23 | 1.61 | 0.94 | 0.77 | 6.27E-09 |
| 3 | PPP2R5C | 0.72 | 0.56 | 4.80 | 3.52 | 0.45 | 8.16E-09 |
| 3 | VMP1 | 0.51 | 0.37 | 2.78 | 1.67 | 0.73 | 9.03E-09 |
| 3 | EIF5 | 0.77 | 0.64 | 5.06 | 3.78 | 0.42 | 1.42E-08 |
| 3 | KLRF1 | 0.45 | 0.31 | 2.17 | 1.42 | 0.62 | 1.48E-08 |
| 3 | IL32 | 0.71 | 0.55 | 7.90 | 5.78 | 0.45 | 2.08E-08 |
| 3 | PLAAT4 | 0.48 | 0.32 | 1.90 | 1.32 | 0.52 | 2.35E-08 |
| 3 | CAPN2 | 0.25 | 0.14 | 0.87 | 0.48 | 0.85 | 4.08E-08 |
| 3 | CD99 | 0.62 | 0.48 | 3.24 | 2.24 | 0.54 | 4.35E-08 |
| 3 | PPP1R2 | 0.64 | 0.51 | 4.04 | 2.84 | 0.50 | 5.63E-08 |
| 3 | ODC1 | 0.36 | 0.22 | 1.61 | 0.93 | 0.78 | 6.70E-08 |
| 3 | CIB1 | 0.57 | 0.41 | 2.32 | 1.63 | 0.51 | 8.77E-08 |
| 3 | CCDC88C | 0.39 | 0.25 | 1.50 | 0.94 | 0.67 | 9.69E-08 |
| 3 | BCAS2 | 0.46 | 0.33 | 2.20 | 1.41 | 0.64 | 1.15E-07 |
| 3 | ARHGEF3 | 0.32 | 0.20 | 1.21 | 0.71 | 0.78 | 1.61E-07 |
| 3 | NDUFB7 | 0.26 | 0.15 | 0.92 | 0.51 | 0.84 | 1.62E-07 |
| 3 | BTG2 | 0.64 | 0.54 | 5.29 | 3.25 | 0.70 | 1.96E-07 |
| 3 | BCL2 | 0.36 | 0.23 | 1.40 | 0.88 | 0.67 | 2.09E-07 |
| 3 | PRNP | 0.47 | 0.35 | 2.28 | 1.45 | 0.65 | 2.26E-07 |
| 3 | ARL4C | 0.78 | 0.67 | 7.69 | 5.67 | 0.44 | 2.26E-07 |
| 3 | MYO1F | 0.31 | 0.18 | 1.04 | 0.74 | 0.49 | 2.66E-07 |
| 3 | TBCB | 0.28 | 0.17 | 0.91 | 0.56 | 0.70 | 2.66E-07 |

| | | | | | | | |
|---|---|---|---|---|---|---|---|
| 3 | ANKRD37 | 0.26 | 0.15 | 1.17 | 0.71 | 0.72 | 3.05E-07 |
| 3 | PAXX | 0.50 | 0.36 | 2.14 | 1.49 | 0.52 | 3.19E-07 |
| 3 | CAPZB | 0.45 | 0.31 | 1.60 | 1.11 | 0.53 | 3.60E-07 |
| 3 | SAP18 | 0.66 | 0.54 | 3.74 | 2.37 | 0.66 | 3.94E-07 |
| 3 | C12orf57 | 0.50 | 0.37 | 2.10 | 1.45 | 0.53 | 4.69E-07 |
| 3 | C1orf52 | 0.30 | 0.19 | 0.98 | 0.61 | 0.69 | 1.27E-06 |
| 3 | DHRS7 | 0.41 | 0.28 | 1.58 | 1.04 | 0.59 | 1.29E-06 |
| 3 | MARCKSL1 | 0.29 | 0.18 | 1.14 | 0.68 | 0.76 | 1.31E-06 |
| 3 | SIAH2 | 0.33 | 0.22 | 1.29 | 0.73 | 0.83 | 1.60E-06 |
| 3 | HMGB2 | 0.58 | 0.46 | 3.57 | 2.67 | 0.42 | 2.22E-06 |
| 3 | ST8SIA4 | 0.32 | 0.20 | 1.11 | 0.78 | 0.50 | 2.22E-06 |
| 3 | CITED2 | 0.32 | 0.22 | 1.83 | 0.96 | 0.94 | 2.36E-06 |
| 3 | WAS | 0.26 | 0.16 | 0.85 | 0.52 | 0.72 | 2.49E-06 |
| 3 | PLIN2 | 0.50 | 0.40 | 4.06 | 2.40 | 0.76 | 2.56E-06 |
| 3 | TENT5A | 0.29 | 0.18 | 1.14 | 0.73 | 0.64 | 3.07E-06 |
| 3 | ZBTB11 | 0.32 | 0.22 | 1.21 | 0.72 | 0.76 | 4.13E-06 |
| 3 | SNHG15 | 0.48 | 0.36 | 2.15 | 1.49 | 0.53 | 4.18E-06 |
| 3 | GCH1 | 0.26 | 0.16 | 0.88 | 0.55 | 0.69 | 4.28E-06 |
| 3 | TNFRSF14 | 0.28 | 0.18 | 0.85 | 0.59 | 0.52 | 4.43E-06 |
| 3 | SIGIRR | 0.31 | 0.20 | 0.95 | 0.66 | 0.52 | 4.85E-06 |
| 3 | IVNS1ABP | 0.50 | 0.37 | 2.35 | 1.65 | 0.51 | 4.89E-06 |
| 3 | PTMS | 0.30 | 0.19 | 1.24 | 0.93 | 0.42 | 6.69E-06 |
| 3 | ZAP70 | 0.30 | 0.20 | 1.08 | 0.75 | 0.52 | 7.43E-06 |
| 3 | OTUD6B-AS | 0.29 | 0.19 | 1.07 | 0.67 | 0.67 | 7.74E-06 |
| 3 | CCDC85B | 0.47 | 0.35 | 1.97 | 1.40 | 0.49 | 7.77E-06 |
| 3 | CDKN2D | 0.46 | 0.35 | 2.11 | 1.47 | 0.52 | 1.13E-05 |
| 3 | OAT | 0.27 | 0.17 | 0.94 | 0.60 | 0.64 | 1.16E-05 |
| 3 | HNRNPU | 0.87 | 0.79 | 8.92 | 6.95 | 0.36 | 1.34E-05 |
| 3 | CORO1A | 0.66 | 0.55 | 3.78 | 2.68 | 0.50 | 1.43E-05 |
| 3 | H2AZ1 | 0.77 | 0.72 | 8.65 | 6.14 | 0.49 | 1.54E-05 |
| 3 | BTG3 | 0.42 | 0.32 | 1.98 | 1.33 | 0.57 | 1.70E-05 |
| 3 | JPT1 | 0.34 | 0.25 | 1.42 | 0.84 | 0.76 | 1.95E-05 |
| 3 | TAF7 | 0.62 | 0.54 | 3.70 | 2.52 | 0.55 | 2.06E-05 |
| 3 | PRKACB | 0.37 | 0.26 | 1.38 | 0.99 | 0.48 | 2.21E-05 |
| 3 | HERPUD1 | 0.52 | 0.43 | 3.00 | 1.99 | 0.59 | 2.29E-05 |
| 3 | EBP | 0.33 | 0.23 | 1.32 | 0.83 | 0.67 | 2.65E-05 |
| 3 | RASSF1 | 0.25 | 0.16 | 0.86 | 0.53 | 0.69 | 2.88E-05 |
| 3 | RBM22 | 0.30 | 0.20 | 1.00 | 0.64 | 0.63 | 3.15E-05 |
| 3 | ABHD3 | 0.35 | 0.25 | 1.35 | 0.91 | 0.57 | 3.42E-05 |
| 3 | DSTN | 0.45 | 0.33 | 2.18 | 1.68 | 0.37 | 3.72E-05 |
| 3 | TBX21 | 0.33 | 0.23 | 1.25 | 0.86 | 0.53 | 3.99E-05 |
| 3 | SKAP1 | 0.31 | 0.22 | 1.10 | 0.75 | 0.54 | 4.35E-05 |
| 3 | VPS29 | 0.26 | 0.17 | 0.83 | 0.57 | 0.53 | 4.40E-05 |
| 3 | HSPA5 | 0.69 | 0.65 | 5.55 | 4.03 | 0.46 | 4.58E-05 |
| 3 | GMFG | 0.41 | 0.31 | 1.73 | 1.14 | 0.60 | 4.95E-05 |
| 3 | POLR1F | 0.29 | 0.20 | 1.38 | 0.84 | 0.71 | 5.12E-05 |
| 3 | CDKN1A | 0.36 | 0.25 | 1.42 | 1.05 | 0.44 | 5.66E-05 |
| 3 | EPC1 | 0.45 | 0.35 | 1.86 | 1.34 | 0.47 | 5.76E-05 |
| 3 | MOB4 | 0.52 | 0.42 | 2.56 | 1.91 | 0.42 | 5.83E-05 |
| 3 | TGFBR3 | 0.26 | 0.16 | 0.94 | 0.73 | 0.37 | 5.84E-05 |
| 3 | DPP7 | 0.31 | 0.21 | 0.96 | 0.71 | 0.43 | 5.88E-05 |
| 3 | HLA-DPB1 | 0.53 | 0.42 | 2.55 | 1.80 | 0.50 | 6.09E-05 |
| 3 | FKBP11 | 0.26 | 0.17 | 0.80 | 0.61 | 0.40 | 6.11E-05 |
| 3 | CD3E | 0.31 | 0.22 | 1.26 | 0.86 | 0.56 | 6.15E-05 |
| 3 | SUN2 | 0.44 | 0.34 | 1.72 | 1.24 | 0.47 | 6.22E-05 |
| 3 | ITK | 0.30 | 0.20 | 1.01 | 0.72 | 0.49 | 9.06E-05 |
| 3 | TECR | 0.35 | 0.24 | 1.22 | 0.85 | 0.52 | 9.88E-05 |
| 3 | ELOC | 0.36 | 0.27 | 1.25 | 0.87 | 0.53 | 9.95E-05 |
| 3 | HLA-DPA1 | 0.48 | 0.38 | 2.32 | 1.62 | 0.52 | 0.00010411 |

| | | | | | | | |
|---|---|---|---|---|---|---|---|
| 3 | NDUFB8 | 0.40 | 0.30 | 1.46 | 1.04 | 0.49 | 0.00010763 |
| 3 | HSPA4 | 0.45 | 0.36 | 1.99 | 1.45 | 0.46 | 0.00010837 |
| 3 | TERF1 | 0.31 | 0.21 | 1.17 | 0.81 | 0.53 | 0.00010894 |
| 3 | IER2 | 0.79 | 0.74 | 9.17 | 7.14 | 0.36 | 0.00010977 |
| 3 | ARPC4 | 0.33 | 0.23 | 1.17 | 0.74 | 0.66 | 0.00011099 |
| 3 | FKBP4 | 0.54 | 0.47 | 4.89 | 3.18 | 0.62 | 0.00011313 |
| 3 | NSD1 | 0.27 | 0.18 | 0.85 | 0.62 | 0.45 | 0.00011344 |
| 3 | TAOK3 | 0.38 | 0.28 | 1.44 | 1.07 | 0.43 | 0.00011841 |
| 3 | RNASEK | 0.26 | 0.17 | 0.85 | 0.61 | 0.48 | 0.00012361 |
| 3 | IER5 | 0.58 | 0.51 | 5.31 | 3.52 | 0.59 | 0.00012847 |
| 3 | MBNL2 | 0.28 | 0.19 | 1.01 | 0.70 | 0.54 | 0.00013426 |
| 3 | GSTK1 | 0.28 | 0.19 | 0.88 | 0.65 | 0.46 | 0.00015415 |
| 3 | CASP8 | 0.39 | 0.29 | 1.51 | 1.07 | 0.49 | 0.00015675 |
| 3 | MLLT6 | 0.37 | 0.27 | 1.38 | 1.03 | 0.42 | 0.0001946 |
| 3 | HERPUD2 | 0.31 | 0.22 | 1.16 | 0.81 | 0.51 | 0.00026455 |
| 3 | PNPLA8 | 0.42 | 0.32 | 1.55 | 1.20 | 0.37 | 0.00035991 |
| 3 | CCNH | 0.46 | 0.37 | 2.12 | 1.50 | 0.50 | 0.00037409 |
| 3 | NABP1 | 0.32 | 0.24 | 1.35 | 0.98 | 0.46 | 0.00041159 |
| 3 | GTF3C1 | 0.29 | 0.21 | 1.37 | 0.86 | 0.67 | 0.00044168 |
| 3 | LAMTOR4 | 0.34 | 0.25 | 1.18 | 0.85 | 0.47 | 0.0004437 |
| 3 | LBH | 0.38 | 0.28 | 1.61 | 1.22 | 0.40 | 0.00046124 |
| 3 | COX8A | 0.52 | 0.43 | 2.18 | 1.64 | 0.41 | 0.00046225 |
| 3 | SEC11C | 0.26 | 0.19 | 0.93 | 0.64 | 0.54 | 0.00047905 |
| 3 | SLC3A2 | 0.61 | 0.55 | 3.41 | 2.58 | 0.40 | 0.00049014 |
| 3 | STIP1 | 0.48 | 0.43 | 2.50 | 1.76 | 0.51 | 0.00057561 |
| 3 | COQ10B | 0.33 | 0.26 | 1.15 | 0.85 | 0.43 | 0.00069139 |
| 3 | CNN2 | 0.43 | 0.34 | 1.79 | 1.35 | 0.41 | 0.00081182 |
| 3 | ARHGEF1 | 0.41 | 0.33 | 1.68 | 1.26 | 0.42 | 0.00086887 |
| 3 | UGP2 | 0.33 | 0.25 | 1.21 | 0.93 | 0.38 | 0.00088648 |
| 3 | HMGN2 | 0.51 | 0.44 | 2.38 | 1.83 | 0.38 | 0.00094328 |
| 3 | CYRIB | 0.42 | 0.33 | 1.61 | 1.22 | 0.40 | 0.00103401 |
| 3 | CYFIP2 | 0.26 | 0.19 | 0.84 | 0.63 | 0.42 | 0.00105311 |
| 3 | SNHG9 | 0.26 | 0.19 | 1.07 | 0.79 | 0.44 | 0.00114406 |
| 3 | ENSG00000 | 0.27 | 0.20 | 1.22 | 0.80 | 0.62 | 0.00116483 |
| 3 | ABI1 | 0.32 | 0.23 | 1.02 | 0.78 | 0.39 | 0.00119741 |
| 3 | UQCR11 | 0.34 | 0.26 | 1.25 | 0.94 | 0.41 | 0.00119926 |
| 3 | CD52 | 0.42 | 0.36 | 2.39 | 1.52 | 0.66 | 0.00120802 |
| 3 | PTGER4 | 0.51 | 0.46 | 3.39 | 2.34 | 0.54 | 0.00126454 |
| 3 | OSER1 | 0.27 | 0.20 | 0.84 | 0.60 | 0.47 | 0.00140749 |
| 3 | GLUL | 0.30 | 0.23 | 1.27 | 0.88 | 0.53 | 0.00153683 |
| 3 | TNFAIP8 | 0.29 | 0.21 | 1.18 | 0.87 | 0.44 | 0.00154824 |
| 3 | VAMP2 | 0.51 | 0.44 | 2.40 | 1.81 | 0.41 | 0.0015762 |
| 3 | H2AX | 0.33 | 0.26 | 1.31 | 0.97 | 0.43 | 0.00193414 |
| 3 | COPE | 0.35 | 0.28 | 1.30 | 0.97 | 0.43 | 0.00195086 |
| 3 | IER3 | 0.30 | 0.23 | 1.77 | 1.23 | 0.53 | 0.00201245 |
| 3 | TAF10 | 0.29 | 0.22 | 0.87 | 0.67 | 0.37 | 0.00231477 |
| 3 | DYNLL2 | 0.31 | 0.23 | 1.02 | 0.75 | 0.45 | 0.0023634 |
| 3 | ENSG00000 | 0.33 | 0.26 | 1.33 | 0.96 | 0.46 | 0.0024156 |
| 3 | RASAL3 | 0.26 | 0.18 | 0.83 | 0.64 | 0.37 | 0.002529 |
| 3 | SS18L2 | 0.31 | 0.24 | 1.12 | 0.77 | 0.54 | 0.00255546 |
| 3 | SNHG5 | 0.59 | 0.53 | 3.56 | 2.59 | 0.46 | 0.00314941 |
| 3 | DAD1 | 0.29 | 0.22 | 0.99 | 0.77 | 0.37 | 0.00333506 |
| 3 | SEM1 | 0.27 | 0.21 | 0.92 | 0.68 | 0.43 | 0.00409162 |
| 3 | BAX | 0.28 | 0.21 | 0.92 | 0.70 | 0.40 | 0.00415708 |
| 3 | TRAPPC4 | 0.25 | 0.19 | 0.82 | 0.60 | 0.44 | 0.00486694 |
| 3 | MT2A | 0.56 | 0.47 | 8.58 | 6.01 | 0.51 | 0.0051235 |
| 3 | TSC22D4 | 0.26 | 0.20 | 0.92 | 0.68 | 0.45 | 0.00539738 |
| 3 | THAP9-AS1 | 0.31 | 0.24 | 1.07 | 0.83 | 0.37 | 0.00568421 |
| 3 | NDUFA12 | 0.34 | 0.27 | 1.22 | 0.92 | 0.41 | 0.00622453 |

| | | | | | | | |
|---|---|---|---|---|---|---|---|
| 3 | NFKBIB | 0.26 | 0.20 | 0.86 | 0.65 | 0.40 | 0.00642927 |
| 3 | C1orf21 | 0.33 | 0.27 | 1.32 | 1.01 | 0.39 | 0.00652712 |
| 3 | UHMK1 | 0.28 | 0.21 | 0.92 | 0.71 | 0.37 | 0.00674031 |
| 3 | DERL1 | 0.31 | 0.24 | 0.98 | 0.76 | 0.37 | 0.00702283 |
| 3 | NSMCE3 | 0.31 | 0.25 | 1.18 | 0.87 | 0.44 | 0.00737554 |
| 3 | ARL6IP4 | 0.27 | 0.21 | 0.94 | 0.69 | 0.45 | 0.00810109 |
| 3 | BANF1 | 0.28 | 0.22 | 0.96 | 0.70 | 0.46 | 0.00940342 |
| 3 | SELENOK | 0.74 | 0.71 | 6.40 | 4.81 | 0.41 | 0.0108514 |
| 3 | HCP5 | 0.25 | 0.20 | 0.91 | 0.64 | 0.52 | 0.01280983 |
| 3 | ATP5PO | 0.26 | 0.21 | 0.86 | 0.63 | 0.44 | 0.01523144 |
| 3 | POLR1H | 0.31 | 0.25 | 1.11 | 0.85 | 0.39 | 0.01557322 |
| 3 | PRR13 | 0.33 | 0.27 | 1.15 | 0.89 | 0.38 | 0.01677422 |
| 3 | SNRPB2 | 0.33 | 0.27 | 1.22 | 0.92 | 0.40 | 0.01712549 |
| 3 | CHMP1B | 0.34 | 0.28 | 1.48 | 1.09 | 0.44 | 0.02469938 |
| 3 | PRMT2 | 0.34 | 0.29 | 1.44 | 1.10 | 0.40 | 0.02742686 |
| 3 | SMIM26 | 0.26 | 0.21 | 0.85 | 0.66 | 0.37 | 0.02825133 |
| 3 | LPIN2 | 0.25 | 0.21 | 0.99 | 0.71 | 0.48 | 0.02979539 |
| 3 | CMC2 | 0.31 | 0.27 | 1.13 | 0.87 | 0.37 | 0.03360671 |
| 3 | GTF2B | 0.28 | 0.24 | 1.03 | 0.79 | 0.38 | 0.0469592 |
| 3 | TUBA1C | 0.30 | 0.26 | 1.21 | 0.87 | 0.47 | 0.04772087 |
| 4 | ITGA1 | 0.53 | 0.10 | 3.74 | 0.49 | 2.94 | 3.88E-55 |
| 4 | ACP5 | 0.26 | 0.03 | 1.08 | 0.08 | 3.69 | 1.55E-45 |
| 4 | LINC02446 | 0.46 | 0.12 | 6.54 | 1.05 | 2.64 | 1.03E-34 |
| 4 | CXCR6 | 0.38 | 0.08 | 1.89 | 0.32 | 2.56 | 7.74E-34 |
| 4 | LGALS3 | 0.51 | 0.17 | 2.48 | 0.61 | 2.03 | 2.19E-27 |
| 4 | HLA-DQA1 | 0.52 | 0.20 | 3.62 | 0.88 | 2.03 | 3.74E-22 |
| 4 | CD74 | 0.95 | 0.72 | 10.66 | 5.21 | 1.03 | 7.63E-22 |
| 4 | CCL5 | 0.96 | 0.81 | 39.51 | 20.38 | 0.96 | 7.75E-22 |
| 4 | HLA-DRB1 | 0.83 | 0.67 | 16.54 | 5.42 | 1.61 | 6.56E-21 |
| 4 | HLA-DRA | 0.74 | 0.47 | 7.03 | 2.34 | 1.59 | 3.24E-20 |
| 4 | CSF1 | 0.28 | 0.07 | 1.68 | 0.41 | 2.02 | 7.98E-20 |
| 4 | ENTPD1 | 0.32 | 0.09 | 2.13 | 0.44 | 2.26 | 1.07E-19 |
| 4 | LDLRAD4 | 0.35 | 0.11 | 1.80 | 0.46 | 1.96 | 1.44E-18 |
| 4 | HLA-DPA1 | 0.68 | 0.38 | 3.70 | 1.59 | 1.22 | 2.42E-17 |
| 4 | LINC01871 | 0.73 | 0.43 | 7.25 | 2.89 | 1.33 | 2.98E-17 |
| 4 | ALOX5AP | 0.57 | 0.27 | 3.58 | 1.36 | 1.40 | 3.46E-16 |
| 4 | GZMA | 0.70 | 0.44 | 22.77 | 4.60 | 2.31 | 1.28E-15 |
| 4 | SRGAP3 | 0.27 | 0.08 | 1.23 | 0.31 | 1.97 | 3.90E-15 |
| 4 | TBCD | 0.26 | 0.08 | 1.14 | 0.26 | 2.11 | 1.12E-14 |
| 4 | GZMK | 0.54 | 0.28 | 6.67 | 1.91 | 1.80 | 1.22E-14 |
| 4 | CD7 | 0.91 | 0.74 | 11.66 | 6.65 | 0.81 | 3.58E-14 |
| 4 | GZMB | 0.90 | 0.67 | 25.17 | 13.33 | 0.92 | 3.93E-14 |
| 4 | COTL1 | 0.68 | 0.39 | 4.44 | 2.20 | 1.01 | 8.30E-14 |
| 4 | HLA-DMA | 0.31 | 0.11 | 1.14 | 0.33 | 1.80 | 1.13E-13 |
| 4 | C4orf48 | 0.26 | 0.08 | 0.79 | 0.25 | 1.64 | 1.43E-13 |
| 4 | TTN | 0.34 | 0.13 | 1.29 | 0.53 | 1.28 | 3.07E-13 |
| 4 | GAPDH | 0.97 | 0.90 | 23.29 | 14.73 | 0.66 | 3.97E-13 |
| 4 | LAG3 | 0.29 | 0.10 | 1.17 | 0.39 | 1.57 | 7.65E-13 |
| 4 | ATP8B4 | 0.36 | 0.14 | 1.25 | 0.49 | 1.35 | 1.57E-12 |
| 4 | HLA-DQB1 | 0.47 | 0.23 | 2.34 | 0.89 | 1.40 | 2.25E-12 |
| 4 | PRF1 | 0.82 | 0.55 | 7.24 | 4.00 | 0.86 | 4.24E-12 |
| 4 | SAMSN1 | 0.76 | 0.53 | 5.52 | 2.97 | 0.89 | 9.15E-12 |
| 4 | PTMS | 0.41 | 0.20 | 2.40 | 0.87 | 1.46 | 1.56E-11 |
| 4 | KLRC2 | 0.60 | 0.35 | 3.56 | 1.74 | 1.03 | 3.49E-11 |
| 4 | IKZF3 | 0.61 | 0.35 | 4.37 | 1.93 | 1.18 | 3.55E-11 |
| 4 | TNFRSF9 | 0.52 | 0.31 | 6.05 | 2.56 | 1.24 | 3.73E-11 |
| 4 | SH2D1B | 0.55 | 0.30 | 2.52 | 1.32 | 0.93 | 7.23E-11 |
| 4 | YWHAB | 0.82 | 0.60 | 5.64 | 3.18 | 0.83 | 8.52E-11 |
| 4 | IRF4 | 0.33 | 0.14 | 1.65 | 0.56 | 1.56 | 1.05E-10 |

| | | | | | | | |
|---|---|---|---|---|---|---|---|
| 4 | PIN1 | 0.26 | 0.09 | 0.80 | 0.28 | 1.51 | 1.47E-10 |
| 4 | CARD16 | 0.47 | 0.24 | 2.03 | 0.97 | 1.06 | 1.88E-10 |
| 4 | CKLF | 0.46 | 0.24 | 2.36 | 0.90 | 1.39 | 3.09E-10 |
| 4 | HLA-DPB1 | 0.64 | 0.43 | 3.41 | 1.81 | 0.91 | 5.48E-10 |
| 4 | MXD4 | 0.58 | 0.34 | 2.51 | 1.33 | 0.92 | 6.51E-10 |
| 4 | STAT5A | 0.46 | 0.24 | 1.85 | 0.97 | 0.93 | 6.81E-10 |
| 4 | GALNT2 | 0.27 | 0.11 | 1.03 | 0.32 | 1.66 | 1.20E-09 |
| 4 | SLA2 | 0.48 | 0.28 | 2.43 | 1.14 | 1.09 | 1.85E-09 |
| 4 | TRDC | 0.70 | 0.50 | 4.98 | 2.74 | 0.86 | 5.30E-09 |
| 4 | OST4 | 0.83 | 0.63 | 4.85 | 3.16 | 0.62 | 8.07E-09 |
| 4 | OSTF1 | 0.58 | 0.35 | 2.42 | 1.31 | 0.88 | 8.88E-09 |
| 4 | GNLY | 0.89 | 0.85 | 100.65 | 53.09 | 0.92 | 1.03E-08 |
| 4 | CD96 | 0.58 | 0.38 | 3.14 | 1.72 | 0.86 | 1.18E-08 |
| 4 | PPP1CA | 0.43 | 0.22 | 1.36 | 0.74 | 0.87 | 1.36E-08 |
| 4 | TTC1 | 0.30 | 0.13 | 0.83 | 0.39 | 1.10 | 1.89E-08 |
| 4 | BST2 | 0.56 | 0.36 | 2.79 | 1.43 | 0.96 | 1.99E-08 |
| 4 | RASAL3 | 0.36 | 0.18 | 1.34 | 0.62 | 1.11 | 2.16E-08 |
| 4 | LCP1 | 0.90 | 0.78 | 9.87 | 6.80 | 0.54 | 3.43E-08 |
| 4 | SURF4 | 0.54 | 0.33 | 2.06 | 1.23 | 0.74 | 3.52E-08 |
| 4 | PLAAT4 | 0.54 | 0.33 | 2.61 | 1.33 | 0.98 | 3.64E-08 |
| 4 | RBX1 | 0.48 | 0.27 | 1.81 | 0.94 | 0.94 | 3.65E-08 |
| 4 | IFI27L2 | 0.28 | 0.12 | 0.93 | 0.42 | 1.16 | 4.45E-08 |
| 4 | APOBEC3G | 0.71 | 0.51 | 4.30 | 2.67 | 0.69 | 5.36E-08 |
| 4 | CYTOR | 0.52 | 0.32 | 2.88 | 1.28 | 1.17 | 6.36E-08 |
| 4 | SUMO2 | 0.89 | 0.77 | 6.71 | 4.81 | 0.48 | 8.10E-08 |
| 4 | AKAP5 | 0.29 | 0.13 | 0.97 | 0.48 | 1.02 | 1.21E-07 |
| 4 | KIR2DL4 | 0.26 | 0.11 | 1.05 | 0.42 | 1.31 | 1.22E-07 |
| 4 | COX6C | 0.60 | 0.40 | 2.53 | 1.58 | 0.68 | 1.50E-07 |
| 4 | ITGAE | 0.31 | 0.15 | 1.22 | 0.54 | 1.17 | 2.36E-07 |
| 4 | TIGIT | 0.46 | 0.29 | 2.75 | 1.36 | 1.02 | 2.40E-07 |
| 4 | GIMAP4 | 0.42 | 0.24 | 1.87 | 1.01 | 0.89 | 2.59E-07 |
| 4 | SMC3 | 0.52 | 0.32 | 2.14 | 1.24 | 0.79 | 3.42E-07 |
| 4 | NDUFC1 | 0.25 | 0.11 | 0.76 | 0.35 | 1.14 | 4.62E-07 |
| 4 | CD52 | 0.55 | 0.36 | 2.62 | 1.58 | 0.73 | 6.02E-07 |
| 4 | RALA | 0.42 | 0.25 | 1.69 | 0.84 | 1.02 | 6.42E-07 |
| 4 | NCK1 | 0.37 | 0.20 | 1.35 | 0.64 | 1.07 | 6.76E-07 |
| 4 | CALM3 | 0.43 | 0.26 | 1.68 | 0.86 | 0.97 | 7.09E-07 |
| 4 | ITM2A | 0.58 | 0.37 | 3.46 | 2.03 | 0.77 | 7.18E-07 |
| 4 | COX7A2 | 0.67 | 0.47 | 3.01 | 1.92 | 0.65 | 7.77E-07 |
| 4 | ARPC3 | 0.76 | 0.60 | 4.61 | 2.96 | 0.64 | 9.26E-07 |
| 4 | ELOB | 0.72 | 0.49 | 2.97 | 2.09 | 0.51 | 9.44E-07 |
| 4 | RGS10 | 0.35 | 0.19 | 1.30 | 0.62 | 1.07 | 9.61E-07 |
| 4 | IL32 | 0.72 | 0.56 | 9.01 | 5.90 | 0.61 | 1.05E-06 |
| 4 | UQCRB | 0.79 | 0.67 | 5.42 | 3.71 | 0.55 | 1.12E-06 |
| 4 | HDAC1 | 0.31 | 0.15 | 0.78 | 0.45 | 0.80 | 1.28E-06 |
| 4 | PSMB9 | 0.54 | 0.35 | 2.13 | 1.40 | 0.60 | 1.28E-06 |
| 4 | SURF2 | 0.27 | 0.13 | 0.86 | 0.40 | 1.11 | 1.47E-06 |
| 4 | CD2 | 0.74 | 0.59 | 6.58 | 4.02 | 0.71 | 1.49E-06 |
| 4 | GGA2 | 0.35 | 0.19 | 1.31 | 0.65 | 1.01 | 1.91E-06 |
| 4 | CARD19 | 0.42 | 0.24 | 1.50 | 0.83 | 0.85 | 1.91E-06 |
| 4 | CAP1 | 0.58 | 0.40 | 2.36 | 1.60 | 0.56 | 2.14E-06 |
| 4 | PMF1 | 0.27 | 0.13 | 0.84 | 0.40 | 1.07 | 2.36E-06 |
| 4 | CORO1A | 0.74 | 0.56 | 4.20 | 2.76 | 0.61 | 2.39E-06 |
| 4 | ARPC1B | 0.52 | 0.32 | 1.88 | 1.17 | 0.69 | 2.47E-06 |
| 4 | APOBEC3C | 0.33 | 0.18 | 1.40 | 0.63 | 1.16 | 2.48E-06 |
| 4 | SERPINB1 | 0.48 | 0.31 | 2.37 | 1.35 | 0.82 | 2.76E-06 |
| 4 | GPRIN3 | 0.56 | 0.36 | 2.61 | 1.53 | 0.77 | 2.88E-06 |
| 4 | PRDX1 | 0.66 | 0.49 | 4.55 | 2.57 | 0.82 | 2.91E-06 |
| 4 | ARPC4 | 0.39 | 0.24 | 1.43 | 0.77 | 0.90 | 3.03E-06 |

| | | | | | | | |
|---|---|---|---|---|---|---|---|
| 4 | BAX | 0.37 | 0.21 | 1.21 | 0.70 | 0.80 | 3.45E-06 |
| 4 | GLIPR1 | 0.49 | 0.31 | 2.14 | 1.36 | 0.66 | 3.58E-06 |
| 4 | NDUFB4 | 0.49 | 0.32 | 1.85 | 1.08 | 0.77 | 3.58E-06 |
| 4 | TRG-AS1 | 0.50 | 0.32 | 2.34 | 1.46 | 0.68 | 3.79E-06 |
| 4 | NDUFA13 | 0.51 | 0.32 | 1.86 | 1.15 | 0.69 | 3.87E-06 |
| 4 | C19orf53 | 0.59 | 0.40 | 2.38 | 1.51 | 0.66 | 4.38E-06 |
| 4 | NEDD8 | 0.44 | 0.26 | 1.42 | 0.88 | 0.69 | 4.61E-06 |
| 4 | FABP5 | 0.28 | 0.14 | 1.05 | 0.57 | 0.88 | 4.85E-06 |
| 4 | SPCS3 | 0.53 | 0.33 | 1.93 | 1.22 | 0.66 | 5.13E-06 |
| 4 | LMAN2 | 0.40 | 0.23 | 1.28 | 0.77 | 0.73 | 5.43E-06 |
| 4 | GBP5 | 0.44 | 0.28 | 2.42 | 1.36 | 0.84 | 5.61E-06 |
| 4 | LAT2 | 0.46 | 0.29 | 2.13 | 1.13 | 0.92 | 5.70E-06 |
| 4 | SLC7A5 | 0.54 | 0.39 | 3.21 | 1.75 | 0.88 | 5.91E-06 |
| 4 | PTTG1 | 0.29 | 0.15 | 0.99 | 0.59 | 0.74 | 6.40E-06 |
| 4 | PARK7 | 0.61 | 0.46 | 3.21 | 1.98 | 0.70 | 6.57E-06 |
| 4 | FAS | 0.36 | 0.20 | 1.34 | 0.77 | 0.81 | 8.13E-06 |
| 4 | ATP5MG | 0.82 | 0.70 | 5.75 | 3.94 | 0.54 | 8.14E-06 |
| 4 | EP300 | 0.40 | 0.24 | 1.24 | 0.78 | 0.66 | 8.41E-06 |
| 4 | SPCS1 | 0.41 | 0.25 | 1.37 | 0.82 | 0.74 | 8.45E-06 |
| 4 | UBL5 | 0.77 | 0.63 | 4.30 | 3.08 | 0.48 | 9.69E-06 |
| 4 | UQCRC2 | 0.40 | 0.24 | 1.44 | 0.80 | 0.85 | 1.00E-05 |
| 4 | CLEC2D | 0.62 | 0.47 | 4.96 | 3.11 | 0.67 | 1.06E-05 |
| 4 | LSM10 | 0.29 | 0.15 | 0.85 | 0.46 | 0.90 | 1.15E-05 |
| 4 | CHCHD10 | 0.27 | 0.14 | 0.87 | 0.43 | 1.03 | 1.16E-05 |
| 4 | SEC61G | 0.53 | 0.35 | 2.24 | 1.34 | 0.74 | 1.25E-05 |
| 4 | PRDX5 | 0.44 | 0.27 | 1.57 | 0.96 | 0.71 | 1.34E-05 |
| 4 | GIMAP7 | 0.41 | 0.24 | 1.72 | 1.07 | 0.68 | 1.43E-05 |
| 4 | APBB1IP | 0.44 | 0.27 | 1.59 | 1.02 | 0.64 | 1.48E-05 |
| 4 | ATP2A2 | 0.34 | 0.19 | 1.18 | 0.63 | 0.91 | 1.64E-05 |
| 4 | NDUFAB1 | 0.34 | 0.19 | 1.02 | 0.59 | 0.79 | 1.78E-05 |
| 4 | GZMH | 0.56 | 0.37 | 4.42 | 3.02 | 0.55 | 1.94E-05 |
| 4 | LSM2 | 0.31 | 0.17 | 0.95 | 0.53 | 0.83 | 2.04E-05 |
| 4 | SMC4 | 0.38 | 0.23 | 1.92 | 0.83 | 1.21 | 2.20E-05 |
| 4 | SLFN12L | 0.46 | 0.29 | 2.18 | 1.28 | 0.76 | 2.27E-05 |
| 4 | PBX4 | 0.27 | 0.14 | 0.99 | 0.53 | 0.91 | 2.27E-05 |
| 4 | EVL | 0.48 | 0.32 | 2.51 | 1.39 | 0.86 | 2.29E-05 |
| 4 | TRAF5 | 0.35 | 0.21 | 1.38 | 0.78 | 0.82 | 2.55E-05 |
| 4 | PGK1 | 0.72 | 0.57 | 4.81 | 3.42 | 0.49 | 2.55E-05 |
| 4 | HOPX | 0.62 | 0.46 | 3.81 | 2.60 | 0.55 | 2.77E-05 |
| 4 | TBC1D10C | 0.31 | 0.17 | 1.25 | 0.64 | 0.97 | 2.82E-05 |
| 4 | SRSF9 | 0.61 | 0.45 | 2.56 | 1.78 | 0.53 | 3.07E-05 |
| 4 | CFL1 | 0.93 | 0.84 | 9.98 | 7.63 | 0.39 | 3.20E-05 |
| 4 | PSMA7 | 0.74 | 0.59 | 4.00 | 3.01 | 0.41 | 3.25E-05 |
| 4 | MYL12B | 0.85 | 0.74 | 6.72 | 5.07 | 0.41 | 3.42E-05 |
| 4 | SYNGR2 | 0.28 | 0.15 | 0.89 | 0.49 | 0.86 | 3.43E-05 |
| 4 | ROMO1 | 0.47 | 0.30 | 1.53 | 1.06 | 0.53 | 3.51E-05 |
| 4 | SNRPF | 0.44 | 0.29 | 1.66 | 1.00 | 0.73 | 3.56E-05 |
| 4 | USP48 | 0.25 | 0.14 | 0.87 | 0.41 | 1.07 | 3.80E-05 |
| 4 | TSPO | 0.37 | 0.24 | 1.46 | 0.78 | 0.89 | 3.94E-05 |
| 4 | PTPN22 | 0.66 | 0.52 | 4.00 | 2.72 | 0.56 | 3.94E-05 |
| 4 | ASXL2 | 0.27 | 0.14 | 0.99 | 0.48 | 1.06 | 4.18E-05 |
| 4 | MYH9 | 0.75 | 0.56 | 3.91 | 2.85 | 0.46 | 4.38E-05 |
| 4 | EDF1 | 0.80 | 0.70 | 5.28 | 3.88 | 0.44 | 4.43E-05 |
| 4 | PKM | 0.52 | 0.35 | 2.24 | 1.55 | 0.53 | 4.61E-05 |
| 4 | SLFN5 | 0.44 | 0.28 | 2.06 | 1.29 | 0.67 | 4.64E-05 |
| 4 | UBE2L3 | 0.44 | 0.28 | 1.47 | 0.92 | 0.67 | 4.82E-05 |
| 4 | SRI | 0.39 | 0.24 | 1.30 | 0.80 | 0.69 | 5.21E-05 |
| 4 | GSTO1 | 0.46 | 0.30 | 1.58 | 1.09 | 0.53 | 5.22E-05 |
| 4 | PPP1R14B | 0.45 | 0.30 | 2.13 | 1.23 | 0.80 | 5.44E-05 |

| | | | | | | | |
|---|---|---|---|---|---|---|---|
| 4 | OCIAD2 | 0.26 | 0.14 | 0.91 | 0.42 | 1.11 | 5.96E-05 |
| 4 | PRRC2A | 0.28 | 0.15 | 0.79 | 0.43 | 0.87 | 6.05E-05 |
| 4 | PIK3AP1 | 0.40 | 0.24 | 1.26 | 0.85 | 0.57 | 6.10E-05 |
| 4 | TNFSF10 | 0.28 | 0.16 | 1.21 | 0.63 | 0.94 | 6.32E-05 |
| 4 | GSPT1 | 0.58 | 0.43 | 3.36 | 2.02 | 0.74 | 6.42E-05 |
| 4 | PPP1R16B | 0.41 | 0.27 | 1.60 | 0.94 | 0.77 | 6.43E-05 |
| 4 | COPS6 | 0.28 | 0.16 | 0.87 | 0.49 | 0.82 | 6.86E-05 |
| 4 | ABI3 | 0.28 | 0.15 | 0.96 | 0.56 | 0.77 | 6.92E-05 |
| 4 | LPIN2 | 0.34 | 0.20 | 1.15 | 0.73 | 0.66 | 7.17E-05 |
| 4 | ITM2C | 0.44 | 0.29 | 2.45 | 1.41 | 0.80 | 7.49E-05 |
| 4 | SYTL2 | 0.27 | 0.15 | 1.12 | 0.55 | 1.01 | 7.53E-05 |
| 4 | CD81 | 0.72 | 0.59 | 4.05 | 2.89 | 0.49 | 7.66E-05 |
| 4 | WNK1 | 0.50 | 0.35 | 2.11 | 1.36 | 0.63 | 7.76E-05 |
| 4 | HMGB2 | 0.62 | 0.47 | 4.02 | 2.72 | 0.56 | 7.82E-05 |
| 4 | CHST12 | 0.50 | 0.34 | 2.31 | 1.56 | 0.57 | 9.29E-05 |
| 4 | TMA7 | 0.79 | 0.71 | 6.12 | 4.30 | 0.51 | 9.42E-05 |
| 4 | UBA2 | 0.32 | 0.19 | 1.03 | 0.58 | 0.84 | 9.63E-05 |
| 4 | MDFIC | 0.26 | 0.14 | 0.99 | 0.50 | 1.00 | 9.66E-05 |
| 4 | SEPTIN7 | 0.87 | 0.73 | 6.03 | 4.59 | 0.39 | 9.83E-05 |
| 4 | PRPF4B | 0.66 | 0.49 | 3.22 | 2.40 | 0.42 | 9.89E-05 |
| 4 | DUSP4 | 0.44 | 0.31 | 3.00 | 1.80 | 0.73 | 0.00010251 |
| 4 | PSMB8 | 0.57 | 0.43 | 2.57 | 1.67 | 0.63 | 0.00010969 |
| 4 | RBPJ | 0.50 | 0.36 | 2.61 | 1.64 | 0.67 | 0.00011781 |
| 4 | CHST11 | 0.31 | 0.18 | 1.03 | 0.59 | 0.79 | 0.00012105 |
| 4 | CDC42SE2 | 0.80 | 0.67 | 4.98 | 3.76 | 0.41 | 0.00012224 |
| 4 | MGST3 | 0.29 | 0.17 | 0.93 | 0.53 | 0.81 | 0.00012328 |
| 4 | TMCO1 | 0.30 | 0.18 | 1.11 | 0.59 | 0.91 | 0.00012346 |
| 4 | CLDND1 | 0.60 | 0.47 | 4.16 | 2.59 | 0.68 | 0.00013671 |
| 4 | COX8A | 0.58 | 0.43 | 2.42 | 1.67 | 0.53 | 0.00013839 |
| 4 | PSMB1 | 0.55 | 0.41 | 2.33 | 1.56 | 0.58 | 0.00013921 |
| 4 | OAZ1 | 0.93 | 0.91 | 17.28 | 13.15 | 0.39 | 0.00014032 |
| 4 | PPP1CC | 0.37 | 0.22 | 1.03 | 0.72 | 0.53 | 0.00015789 |
| 4 | FIS1 | 0.33 | 0.20 | 1.09 | 0.64 | 0.76 | 0.00017022 |
| 4 | HDLBP | 0.33 | 0.20 | 1.17 | 0.66 | 0.84 | 0.00017686 |
| 4 | NDUFS5 | 0.67 | 0.51 | 3.09 | 2.30 | 0.43 | 0.00019252 |
| 4 | STUB1 | 0.38 | 0.24 | 1.20 | 0.77 | 0.64 | 0.00019427 |
| 4 | IL2RB | 0.71 | 0.57 | 4.36 | 3.16 | 0.47 | 0.00020183 |
| 4 | PSMA4 | 0.39 | 0.26 | 1.43 | 0.86 | 0.73 | 0.00020729 |
| 4 | SRP14 | 0.87 | 0.80 | 7.84 | 6.06 | 0.37 | 0.00021505 |
| 4 | CD99 | 0.62 | 0.50 | 3.42 | 2.32 | 0.56 | 0.00021616 |
| 4 | EMC7 | 0.39 | 0.24 | 1.06 | 0.75 | 0.50 | 0.000218 |
| 4 | CIAO1 | 0.31 | 0.18 | 0.88 | 0.58 | 0.59 | 0.00022292 |
| 4 | ITGB1 | 0.82 | 0.74 | 7.61 | 5.77 | 0.40 | 0.0002322 |
| 4 | PRDX2 | 0.25 | 0.14 | 0.77 | 0.45 | 0.78 | 0.00023606 |
| 4 | MPC2 | 0.36 | 0.22 | 0.96 | 0.67 | 0.52 | 0.00024025 |
| 4 | PET100 | 0.39 | 0.25 | 1.31 | 0.89 | 0.57 | 0.00025622 |
| 4 | DNAJB14 | 0.44 | 0.30 | 1.43 | 1.00 | 0.52 | 0.00025962 |
| 4 | SEM1 | 0.34 | 0.21 | 1.08 | 0.69 | 0.64 | 0.00026181 |
| 4 | REEP5 | 0.41 | 0.28 | 1.46 | 0.90 | 0.70 | 0.00026462 |
| 4 | ANXA2 | 0.49 | 0.33 | 1.88 | 1.39 | 0.43 | 0.00026496 |
| 4 | RABAC1 | 0.52 | 0.38 | 2.17 | 1.45 | 0.58 | 0.00027194 |
| 4 | PPM1G | 0.52 | 0.35 | 1.83 | 1.34 | 0.45 | 0.00027232 |
| 4 | CLTC | 0.29 | 0.17 | 0.92 | 0.50 | 0.87 | 0.00028048 |
| 4 | PPP4C | 0.35 | 0.22 | 1.12 | 0.74 | 0.61 | 0.00028814 |
| 4 | ANXA5 | 0.34 | 0.21 | 1.41 | 0.80 | 0.82 | 0.00029443 |
| 4 | NDUFB11 | 0.56 | 0.43 | 2.23 | 1.59 | 0.49 | 0.00029747 |
| 4 | LAMTOR5 | 0.42 | 0.27 | 1.33 | 0.90 | 0.57 | 0.00030108 |
| 4 | NUDC | 0.59 | 0.44 | 2.72 | 1.93 | 0.49 | 0.00030367 |
| 4 | PDCD4 | 0.61 | 0.47 | 3.57 | 2.32 | 0.62 | 0.00031108 |

| | | | | | | | |
|---|---|---|---|---|---|---|---|
| 4 | PGAM1 | 0.38 | 0.26 | 1.39 | 0.89 | 0.65 | 0.00031187 |
| 4 | PPP1R35 | 0.28 | 0.16 | 0.79 | 0.52 | 0.60 | 0.00031795 |
| 4 | ZBTB38 | 0.31 | 0.18 | 1.18 | 0.70 | 0.76 | 0.00033565 |
| 4 | MSI2 | 0.27 | 0.16 | 0.83 | 0.48 | 0.79 | 0.00033776 |
| 4 | DNAJA2 | 0.55 | 0.39 | 1.96 | 1.36 | 0.53 | 0.0003491 |
| 4 | NAMPT | 0.66 | 0.53 | 4.79 | 3.71 | 0.37 | 0.00035462 |
| 4 | RPS27L | 0.52 | 0.35 | 1.88 | 1.39 | 0.44 | 0.00035789 |
| 4 | PRKACB | 0.40 | 0.27 | 1.49 | 1.02 | 0.55 | 0.00038278 |
| 4 | NOL7 | 0.41 | 0.27 | 1.30 | 0.93 | 0.49 | 0.00040295 |
| 4 | N4BP1 | 0.41 | 0.28 | 1.49 | 1.02 | 0.55 | 0.00041693 |
| 4 | NDUFA12 | 0.41 | 0.27 | 1.38 | 0.94 | 0.55 | 0.0004435 |
| 4 | CASP4 | 0.40 | 0.27 | 1.31 | 0.85 | 0.62 | 0.00044509 |
| 4 | FASLG | 0.42 | 0.28 | 1.66 | 1.14 | 0.55 | 0.00046621 |
| 4 | CIAO2B | 0.48 | 0.35 | 1.78 | 1.28 | 0.47 | 0.00046651 |
| 4 | SUB1 | 0.81 | 0.74 | 5.81 | 4.52 | 0.36 | 0.00049231 |
| 4 | ATP5MK | 0.47 | 0.33 | 1.83 | 1.22 | 0.59 | 0.00049362 |
| 4 | RNF213 | 0.70 | 0.55 | 4.92 | 3.25 | 0.60 | 0.00050155 |
| 4 | SHFL | 0.31 | 0.19 | 1.09 | 0.61 | 0.83 | 0.00050512 |
| 4 | PSME2 | 0.50 | 0.38 | 2.19 | 1.45 | 0.59 | 0.00051491 |
| 4 | TMEM248 | 0.31 | 0.18 | 0.82 | 0.56 | 0.55 | 0.00052424 |
| 4 | DR1 | 0.42 | 0.28 | 1.37 | 0.96 | 0.51 | 0.00052705 |
| 4 | HUWE1 | 0.48 | 0.36 | 2.25 | 1.37 | 0.71 | 0.00053102 |
| 4 | PAIP2 | 0.66 | 0.50 | 3.26 | 2.31 | 0.50 | 0.00054209 |
| 4 | SRP19 | 0.38 | 0.25 | 1.15 | 0.81 | 0.51 | 0.00055271 |
| 4 | CAPG | 0.27 | 0.16 | 1.08 | 0.59 | 0.87 | 0.00055378 |
| 4 | NDUFA6 | 0.43 | 0.30 | 1.50 | 1.01 | 0.57 | 0.0005816 |
| 4 | ANXA7 | 0.27 | 0.16 | 0.76 | 0.46 | 0.72 | 0.00060142 |
| 4 | LCK | 0.42 | 0.29 | 1.55 | 1.03 | 0.59 | 0.00060619 |
| 4 | TXK | 0.36 | 0.25 | 2.00 | 1.04 | 0.95 | 0.00062237 |
| 4 | SPN | 0.45 | 0.30 | 1.66 | 1.14 | 0.54 | 0.00062721 |
| 4 | ATP5IF1 | 0.48 | 0.34 | 1.86 | 1.33 | 0.48 | 0.00062801 |
| 4 | ACTN4 | 0.66 | 0.54 | 3.46 | 2.54 | 0.45 | 0.0006389 |
| 4 | RTF2 | 0.36 | 0.22 | 0.98 | 0.70 | 0.47 | 0.00065278 |
| 4 | METAP2 | 0.29 | 0.19 | 1.16 | 0.62 | 0.89 | 0.00066531 |
| 4 | COMMD6 | 0.74 | 0.59 | 3.80 | 2.87 | 0.40 | 0.00067979 |
| 4 | FAM3C | 0.31 | 0.19 | 0.97 | 0.60 | 0.70 | 0.00068286 |
| 4 | NDUFB8 | 0.44 | 0.31 | 1.55 | 1.08 | 0.52 | 0.00068925 |
| 4 | LSP1 | 0.49 | 0.35 | 2.11 | 1.44 | 0.55 | 0.00069654 |
| 4 | DOCK10 | 0.35 | 0.22 | 1.10 | 0.74 | 0.57 | 0.00071228 |
| 4 | CCT7 | 0.32 | 0.20 | 0.90 | 0.60 | 0.59 | 0.00072134 |
| 4 | ATP5MF | 0.41 | 0.29 | 1.55 | 1.05 | 0.56 | 0.00072606 |
| 4 | CCT6A | 0.47 | 0.34 | 1.58 | 1.18 | 0.43 | 0.00074671 |
| 4 | CREBZF | 0.46 | 0.32 | 1.71 | 1.24 | 0.47 | 0.00076839 |
| 4 | LY6E | 0.42 | 0.30 | 1.86 | 1.10 | 0.75 | 0.00077539 |
| 4 | SELENOW | 0.50 | 0.37 | 2.27 | 1.55 | 0.55 | 0.0007986 |
| 4 | SAMD3 | 0.62 | 0.48 | 3.27 | 2.44 | 0.42 | 0.00082486 |
| 4 | TPR | 0.52 | 0.40 | 2.48 | 1.70 | 0.54 | 0.00082826 |
| 4 | UQCR10 | 0.45 | 0.30 | 1.63 | 1.11 | 0.56 | 0.00083871 |
| 4 | NIBAN1 | 0.33 | 0.22 | 1.41 | 0.89 | 0.66 | 0.00084009 |
| 4 | PDCL3 | 0.29 | 0.18 | 0.96 | 0.60 | 0.69 | 0.00086409 |
| 4 | MAP2K2 | 0.46 | 0.32 | 1.53 | 1.09 | 0.49 | 0.00087134 |
| 4 | TIAL1 | 0.44 | 0.30 | 1.58 | 1.11 | 0.51 | 0.00088609 |
| 4 | FKBP4 | 0.58 | 0.47 | 4.80 | 3.35 | 0.52 | 0.00089409 |
| 4 | PSMB2 | 0.32 | 0.21 | 1.06 | 0.64 | 0.73 | 0.00092309 |
| 4 | RAB27A | 0.47 | 0.35 | 2.21 | 1.36 | 0.70 | 0.00092802 |
| 4 | PSMB4 | 0.40 | 0.28 | 1.29 | 0.87 | 0.57 | 0.00094249 |
| 4 | DYNLT1 | 0.35 | 0.23 | 1.20 | 0.79 | 0.60 | 0.0009462 |
| 4 | PPDPF | 0.52 | 0.41 | 2.21 | 1.55 | 0.52 | 0.0009568 |
| 4 | BLOC1S1 | 0.31 | 0.19 | 0.88 | 0.62 | 0.50 | 0.00096875 |

| | | | | | | | |
|---|---|---|---|---|---|---|---|
| 4 | COPB2 | 0.26 | 0.15 | 0.71 | 0.46 | 0.63 | 0.00097198 |
| 4 | STAT5B | 0.26 | 0.16 | 0.83 | 0.56 | 0.57 | 0.00097538 |
| 4 | GPR174 | 0.36 | 0.24 | 1.39 | 0.88 | 0.67 | 0.00102291 |
| 4 | ATP5F1C | 0.36 | 0.24 | 1.28 | 0.82 | 0.64 | 0.00103677 |
| 4 | MZT2B | 0.64 | 0.48 | 2.45 | 1.88 | 0.38 | 0.00104248 |
| 4 | OTUB1 | 0.27 | 0.16 | 0.90 | 0.50 | 0.86 | 0.00105361 |
| 4 | MAGOH | 0.40 | 0.28 | 1.37 | 0.95 | 0.54 | 0.0010704 |
| 4 | TNIP2 | 0.31 | 0.19 | 0.96 | 0.61 | 0.64 | 0.0010885 |
| 4 | GSTP1 | 0.59 | 0.45 | 2.84 | 2.05 | 0.47 | 0.00110985 |
| 4 | ZMAT2 | 0.29 | 0.18 | 0.84 | 0.53 | 0.66 | 0.00114701 |
| 4 | ACTB | 0.91 | 0.85 | 14.30 | 11.12 | 0.36 | 0.00125095 |
| 4 | EIF3G | 0.68 | 0.57 | 3.71 | 2.65 | 0.49 | 0.00130466 |
| 4 | SNRPG | 0.57 | 0.44 | 2.23 | 1.72 | 0.37 | 0.00131534 |
| 4 | LPIN1 | 0.36 | 0.25 | 1.34 | 0.89 | 0.59 | 0.00135455 |
| 4 | ATP5PF | 0.38 | 0.27 | 1.38 | 0.90 | 0.62 | 0.0013644 |
| 4 | CSTB | 0.38 | 0.26 | 1.31 | 0.92 | 0.52 | 0.00143397 |
| 4 | ATP6V1F | 0.38 | 0.27 | 1.32 | 0.85 | 0.64 | 0.00145744 |
| 4 | TTC39C | 0.36 | 0.24 | 1.26 | 0.86 | 0.55 | 0.00152949 |
| 4 | GADD45GIF | 0.33 | 0.21 | 1.03 | 0.69 | 0.59 | 0.00154616 |
| 4 | NCOA7 | 0.29 | 0.18 | 1.12 | 0.71 | 0.65 | 0.00155413 |
| 4 | COX7A2L | 0.37 | 0.25 | 1.30 | 0.86 | 0.58 | 0.00156238 |
| 4 | METTL9 | 0.33 | 0.21 | 0.93 | 0.68 | 0.46 | 0.00158021 |
| 4 | SOD1 | 0.70 | 0.66 | 6.92 | 5.00 | 0.47 | 0.00165047 |
| 4 | TYROBP | 0.81 | 0.74 | 7.04 | 5.41 | 0.38 | 0.00167004 |
| 4 | DYNLRB1 | 0.32 | 0.21 | 0.98 | 0.67 | 0.56 | 0.00168501 |
| 4 | PRELID1 | 0.56 | 0.44 | 2.46 | 1.80 | 0.45 | 0.00171543 |
| 4 | HEBP2 | 0.27 | 0.16 | 0.72 | 0.49 | 0.55 | 0.0017328 |
| 4 | MAP4 | 0.29 | 0.19 | 1.04 | 0.61 | 0.77 | 0.00174124 |
| 4 | STOM | 0.40 | 0.29 | 1.49 | 1.03 | 0.53 | 0.00178867 |
| 4 | CALCOCO2 | 0.35 | 0.24 | 1.14 | 0.76 | 0.57 | 0.00183659 |
| 4 | STXBP3 | 0.31 | 0.19 | 0.92 | 0.61 | 0.59 | 0.00184712 |
| 4 | TRAPPC1 | 0.31 | 0.19 | 0.95 | 0.61 | 0.65 | 0.00197596 |
| 4 | HSBP1 | 0.26 | 0.16 | 0.68 | 0.48 | 0.50 | 0.00204121 |
| 4 | IK | 0.52 | 0.39 | 2.13 | 1.53 | 0.48 | 0.00205276 |
| 4 | LSM8 | 0.43 | 0.30 | 1.48 | 1.07 | 0.47 | 0.00210252 |
| 4 | HCLS1 | 0.42 | 0.31 | 1.57 | 1.12 | 0.49 | 0.0021403 |
| 4 | SSU72 | 0.32 | 0.21 | 1.07 | 0.66 | 0.70 | 0.00218991 |
| 4 | NDUFA1 | 0.50 | 0.39 | 1.98 | 1.46 | 0.44 | 0.00225024 |
| 4 | SPCS2 | 0.50 | 0.36 | 1.77 | 1.37 | 0.37 | 0.0022747 |
| 4 | PAK2 | 0.56 | 0.43 | 2.48 | 1.88 | 0.40 | 0.00229788 |
| 4 | MDH2 | 0.25 | 0.16 | 0.76 | 0.48 | 0.65 | 0.00243536 |
| 4 | RASGRP1 | 0.32 | 0.21 | 1.20 | 0.82 | 0.55 | 0.00244259 |
| 4 | PSMB3 | 0.33 | 0.23 | 1.14 | 0.73 | 0.65 | 0.00245132 |
| 4 | TIMM13 | 0.26 | 0.16 | 0.68 | 0.48 | 0.50 | 0.00250163 |
| 4 | COPE | 0.39 | 0.28 | 1.56 | 0.98 | 0.67 | 0.00257974 |
| 4 | EIF2S3 | 0.45 | 0.34 | 1.61 | 1.20 | 0.42 | 0.00268673 |
| 4 | C21orf91 | 0.27 | 0.18 | 0.92 | 0.57 | 0.69 | 0.00271509 |
| 4 | LDHB | 0.41 | 0.29 | 1.44 | 1.02 | 0.50 | 0.00277367 |
| 4 | VPS28 | 0.34 | 0.23 | 1.17 | 0.78 | 0.58 | 0.00278208 |
| 4 | SF3B6 | 0.45 | 0.33 | 1.64 | 1.16 | 0.51 | 0.00289528 |
| 4 | NDUFB1 | 0.33 | 0.22 | 1.36 | 0.78 | 0.80 | 0.00289656 |
| 4 | SYNRG | 0.40 | 0.29 | 1.44 | 1.03 | 0.48 | 0.00294891 |
| 4 | GSTK1 | 0.31 | 0.20 | 0.99 | 0.66 | 0.58 | 0.00297767 |
| 4 | ATP5MC3 | 0.53 | 0.43 | 2.25 | 1.67 | 0.43 | 0.00306648 |
| 4 | TBL1XR1 | 0.37 | 0.26 | 1.23 | 0.91 | 0.43 | 0.00309668 |
| 4 | PSMD13 | 0.39 | 0.29 | 1.51 | 1.04 | 0.54 | 0.00328867 |
| 4 | ABCB1 | 0.27 | 0.17 | 1.06 | 0.60 | 0.82 | 0.00333687 |
| 4 | ANXA6 | 0.34 | 0.24 | 1.19 | 0.77 | 0.62 | 0.00333808 |
| 4 | MRPL57 | 0.31 | 0.20 | 0.95 | 0.63 | 0.60 | 0.00340319 |

| | | | | | | | |
|---|---|---|---|---|---|---|---|
| 4 | BATF | 0.25 | 0.16 | 0.87 | 0.57 | 0.60 | 0.00345672 |
| 4 | TMEM258 | 0.40 | 0.29 | 1.47 | 1.07 | 0.46 | 0.00346615 |
| 4 | MZT2A | 0.39 | 0.28 | 1.32 | 0.90 | 0.54 | 0.0035223 |
| 4 | PIK3CG | 0.28 | 0.18 | 1.00 | 0.60 | 0.75 | 0.00355014 |
| 4 | GID8 | 0.26 | 0.16 | 0.68 | 0.48 | 0.50 | 0.00358396 |
| 4 | PTPN1 | 0.31 | 0.20 | 0.97 | 0.67 | 0.53 | 0.00365414 |
| 4 | SUPT16H | 0.31 | 0.21 | 1.02 | 0.69 | 0.55 | 0.00374661 |
| 4 | BRK1 | 0.40 | 0.28 | 1.21 | 0.90 | 0.43 | 0.00380131 |
| 4 | MAN1A1 | 0.26 | 0.17 | 0.98 | 0.57 | 0.78 | 0.00385305 |
| 4 | PDCD10 | 0.34 | 0.24 | 1.19 | 0.79 | 0.59 | 0.00390731 |
| 4 | UBA6 | 0.32 | 0.21 | 0.89 | 0.68 | 0.39 | 0.00406429 |
| 4 | HARS1 | 0.25 | 0.16 | 0.65 | 0.47 | 0.45 | 0.0040682 |
| 4 | PDIA6 | 0.40 | 0.28 | 1.38 | 1.00 | 0.47 | 0.00415131 |
| 4 | PSMD8 | 0.50 | 0.36 | 1.60 | 1.21 | 0.40 | 0.0042155 |
| 4 | SERINC1 | 0.48 | 0.37 | 1.79 | 1.34 | 0.42 | 0.00422789 |
| 4 | RAB10 | 0.34 | 0.26 | 1.34 | 0.83 | 0.69 | 0.00424373 |
| 4 | MYDGF | 0.30 | 0.20 | 0.94 | 0.69 | 0.43 | 0.00460767 |
| 4 | NCBP3 | 0.29 | 0.19 | 0.97 | 0.60 | 0.71 | 0.00469313 |
| 4 | MRPL33 | 0.36 | 0.26 | 1.41 | 0.94 | 0.59 | 0.00485459 |
| 4 | COX14 | 0.39 | 0.27 | 1.18 | 0.89 | 0.41 | 0.00488572 |
| 4 | PSMA5 | 0.38 | 0.29 | 1.58 | 1.01 | 0.64 | 0.00490216 |
| 4 | RESF1 | 0.50 | 0.38 | 2.33 | 1.74 | 0.42 | 0.0049066 |
| 4 | POMP | 0.54 | 0.44 | 2.45 | 1.81 | 0.44 | 0.00514296 |
| 4 | ROCK1 | 0.53 | 0.42 | 2.29 | 1.70 | 0.43 | 0.00516584 |
| 4 | TMEM160 | 0.29 | 0.20 | 0.93 | 0.66 | 0.51 | 0.00530237 |
| 4 | GTF2I | 0.41 | 0.30 | 1.48 | 1.07 | 0.47 | 0.00542312 |
| 4 | CYSTM1 | 0.36 | 0.25 | 1.11 | 0.84 | 0.40 | 0.0054688 |
| 4 | CCND2 | 0.55 | 0.44 | 2.66 | 2.01 | 0.41 | 0.00566979 |
| 4 | DHRS7 | 0.40 | 0.29 | 1.55 | 1.10 | 0.50 | 0.00570529 |
| 4 | TYMP | 0.26 | 0.17 | 0.87 | 0.61 | 0.51 | 0.00581002 |
| 4 | CIB1 | 0.54 | 0.43 | 2.24 | 1.70 | 0.40 | 0.00585516 |
| 4 | NCKAP1L | 0.31 | 0.22 | 1.15 | 0.73 | 0.65 | 0.00591042 |
| 4 | NDUFB10 | 0.37 | 0.26 | 1.18 | 0.89 | 0.41 | 0.00591809 |
| 4 | MRPL52 | 0.29 | 0.19 | 0.83 | 0.59 | 0.49 | 0.00595367 |
| 4 | ZNF217 | 0.31 | 0.21 | 1.12 | 0.73 | 0.62 | 0.00597755 |
| 4 | STAM | 0.41 | 0.31 | 1.65 | 1.21 | 0.45 | 0.00601749 |
| 4 | SPSB3 | 0.40 | 0.29 | 1.48 | 0.99 | 0.58 | 0.00619557 |
| 4 | BACH1 | 0.27 | 0.18 | 0.84 | 0.57 | 0.55 | 0.00620829 |
| 4 | VTI1B | 0.25 | 0.16 | 0.71 | 0.52 | 0.46 | 0.00627224 |
| 4 | NT5C | 0.30 | 0.20 | 0.98 | 0.64 | 0.61 | 0.00631349 |
| 4 | TMF1 | 0.48 | 0.36 | 1.77 | 1.35 | 0.39 | 0.00635146 |
| 4 | EIF3M | 0.38 | 0.26 | 1.13 | 0.85 | 0.42 | 0.00639672 |
| 4 | HERC1 | 0.28 | 0.19 | 1.05 | 0.61 | 0.78 | 0.00644101 |
| 4 | MIS18BP1 | 0.35 | 0.24 | 1.20 | 0.87 | 0.46 | 0.00653551 |
| 4 | MRPL51 | 0.31 | 0.21 | 0.96 | 0.68 | 0.50 | 0.00665161 |
| 4 | LSM7 | 0.38 | 0.28 | 1.31 | 0.93 | 0.50 | 0.00699596 |
| 4 | ITM2B | 0.77 | 0.64 | 4.70 | 3.52 | 0.42 | 0.00722192 |
| 4 | NIPBL | 0.50 | 0.39 | 2.24 | 1.60 | 0.48 | 0.00730384 |
| 4 | PITPNC1 | 0.38 | 0.28 | 1.52 | 1.11 | 0.46 | 0.00733861 |
| 4 | CTSD | 0.38 | 0.29 | 1.93 | 1.13 | 0.78 | 0.00742603 |
| 4 | RB1CC1 | 0.35 | 0.24 | 1.11 | 0.85 | 0.37 | 0.00754924 |
| 4 | SH2D2A | 0.42 | 0.31 | 1.72 | 1.28 | 0.42 | 0.0075648 |
| 4 | MRPS21 | 0.27 | 0.18 | 0.85 | 0.59 | 0.53 | 0.00764233 |
| 4 | ARF5 | 0.29 | 0.20 | 1.01 | 0.65 | 0.63 | 0.00768453 |
| 4 | SELENOF | 0.29 | 0.20 | 0.89 | 0.60 | 0.56 | 0.00822381 |
| 4 | COPS9 | 0.37 | 0.26 | 1.16 | 0.88 | 0.41 | 0.00827777 |
| 4 | DGUOK | 0.26 | 0.17 | 0.86 | 0.55 | 0.64 | 0.00833127 |
| 4 | RWDD1 | 0.44 | 0.33 | 1.61 | 1.24 | 0.38 | 0.00837968 |
| 4 | TBCB | 0.26 | 0.18 | 0.90 | 0.59 | 0.60 | 0.00838109 |

| | | | | | | | |
|---|---|---|---|---|---|---|---|
| 4 | DCXR | 0.34 | 0.25 | 1.22 | 0.86 | 0.50 | 0.00853132 |
| 4 | SLAMF7 | 0.41 | 0.31 | 1.71 | 1.30 | 0.39 | 0.00870085 |
| 4 | LIMS1 | 0.26 | 0.17 | 0.83 | 0.56 | 0.58 | 0.00890903 |
| 4 | PLP2 | 0.41 | 0.31 | 1.60 | 1.09 | 0.56 | 0.00892853 |
| 4 | BIRC3 | 0.70 | 0.64 | 14.74 | 10.57 | 0.48 | 0.00901912 |
| 4 | LSM1 | 0.27 | 0.17 | 0.69 | 0.53 | 0.39 | 0.00902074 |
| 4 | ADAM8 | 0.35 | 0.27 | 1.57 | 0.98 | 0.68 | 0.00913478 |
| 4 | CANX | 0.45 | 0.34 | 1.60 | 1.23 | 0.38 | 0.00927481 |
| 4 | HERPUD2 | 0.33 | 0.23 | 1.16 | 0.85 | 0.45 | 0.00938434 |
| 4 | CASP3 | 0.28 | 0.19 | 0.94 | 0.65 | 0.53 | 0.00962934 |
| 4 | PRR13 | 0.36 | 0.27 | 1.34 | 0.90 | 0.58 | 0.00967358 |
| 4 | AP2M1 | 0.29 | 0.20 | 0.90 | 0.62 | 0.54 | 0.01003678 |
| 4 | BRWD1 | 0.27 | 0.20 | 1.21 | 0.62 | 0.96 | 0.01007472 |
| 4 | SNRPD3 | 0.36 | 0.27 | 1.17 | 0.86 | 0.44 | 0.01023766 |
| 4 | HES4 | 0.26 | 0.18 | 1.96 | 1.06 | 0.88 | 0.0104737 |
| 4 | RAB6A | 0.28 | 0.19 | 0.83 | 0.60 | 0.48 | 0.01055574 |
| 4 | EVI2B | 0.48 | 0.37 | 1.94 | 1.50 | 0.37 | 0.01060708 |
| 4 | ACADVL | 0.34 | 0.24 | 1.17 | 0.83 | 0.50 | 0.01100993 |
| 4 | MBP | 0.66 | 0.60 | 4.93 | 3.69 | 0.42 | 0.01104086 |
| 4 | UBE2K | 0.34 | 0.25 | 1.13 | 0.84 | 0.43 | 0.01127594 |
| 4 | XRN1 | 0.39 | 0.30 | 1.54 | 1.06 | 0.54 | 0.01167414 |
| 4 | MBD2 | 0.33 | 0.23 | 1.17 | 0.78 | 0.58 | 0.01236188 |
| 4 | PREX1 | 0.46 | 0.35 | 2.04 | 1.42 | 0.53 | 0.0123733 |
| 4 | DEGS1 | 0.27 | 0.19 | 0.78 | 0.58 | 0.44 | 0.01237859 |
| 4 | ANAPC16 | 0.39 | 0.29 | 1.41 | 1.09 | 0.37 | 0.01283787 |
| 4 | TRBC2 | 0.48 | 0.38 | 2.17 | 1.62 | 0.42 | 0.012914 |
| 4 | ATP5F1D | 0.51 | 0.42 | 2.24 | 1.62 | 0.47 | 0.01329637 |
| 4 | EIF6 | 0.31 | 0.22 | 0.93 | 0.69 | 0.42 | 0.01347247 |
| 4 | ERGIC1 | 0.33 | 0.23 | 1.03 | 0.80 | 0.37 | 0.01372851 |
| 4 | SEC61B | 0.59 | 0.52 | 2.86 | 2.22 | 0.36 | 0.01397658 |
| 4 | TUFM | 0.34 | 0.26 | 1.27 | 0.87 | 0.56 | 0.01422354 |
| 4 | EIF2S2 | 0.42 | 0.31 | 1.62 | 1.25 | 0.37 | 0.01425459 |
| 4 | ATP5PD | 0.26 | 0.18 | 0.77 | 0.56 | 0.47 | 0.01472497 |
| 4 | NDUFA4 | 0.51 | 0.43 | 2.21 | 1.66 | 0.41 | 0.01516973 |
| 4 | SUZ12 | 0.34 | 0.24 | 1.15 | 0.82 | 0.49 | 0.01530448 |
| 4 | CMTM3 | 0.27 | 0.19 | 0.85 | 0.55 | 0.65 | 0.01609143 |
| 4 | BBLN | 0.42 | 0.34 | 1.89 | 1.22 | 0.63 | 0.01696338 |
| 4 | RPS19BP1 | 0.36 | 0.28 | 1.33 | 0.91 | 0.55 | 0.01718764 |
| 4 | UBE2N | 0.34 | 0.26 | 1.05 | 0.81 | 0.37 | 0.01747458 |
| 4 | TCEA1 | 0.38 | 0.31 | 1.51 | 1.06 | 0.51 | 0.01759372 |
| 4 | PHF12 | 0.27 | 0.19 | 0.84 | 0.61 | 0.45 | 0.01831399 |
| 4 | PPIB | 0.62 | 0.52 | 3.10 | 2.30 | 0.43 | 0.01891244 |
| 4 | EMB | 0.34 | 0.26 | 1.22 | 0.87 | 0.49 | 0.01939378 |
| 4 | ATP5MJ | 0.36 | 0.29 | 1.41 | 1.00 | 0.49 | 0.01945041 |
| 4 | ZNHIT1 | 0.25 | 0.17 | 0.74 | 0.57 | 0.39 | 0.01967854 |
| 4 | ATP2B4 | 0.29 | 0.21 | 1.08 | 0.78 | 0.46 | 0.01987596 |
| 4 | ERH | 0.34 | 0.25 | 1.12 | 0.82 | 0.45 | 0.02014978 |
| 4 | USP10 | 0.27 | 0.19 | 0.74 | 0.55 | 0.43 | 0.02126953 |
| 4 | S100A10 | 0.76 | 0.73 | 8.58 | 6.47 | 0.41 | 0.02144656 |
| 4 | CEP350 | 0.38 | 0.28 | 1.39 | 1.04 | 0.43 | 0.02205117 |
| 4 | PSMA1 | 0.36 | 0.28 | 1.48 | 0.95 | 0.63 | 0.02239738 |
| 4 | GHITM | 0.45 | 0.35 | 1.66 | 1.27 | 0.38 | 0.02293712 |
| 4 | SMARCC1 | 0.29 | 0.21 | 0.97 | 0.68 | 0.51 | 0.02332595 |
| 4 | FAM118A | 0.29 | 0.21 | 1.14 | 0.69 | 0.72 | 0.02340784 |
| 4 | IL16 | 0.25 | 0.18 | 0.98 | 0.63 | 0.63 | 0.02344701 |
| 4 | TXNL4A | 0.29 | 0.21 | 0.90 | 0.64 | 0.48 | 0.02378257 |
| 4 | SIPA1 | 0.29 | 0.20 | 0.94 | 0.67 | 0.47 | 0.02401977 |
| 4 | TRIP12 | 0.29 | 0.20 | 0.93 | 0.66 | 0.49 | 0.02429582 |
| 4 | RPS6KA3 | 0.37 | 0.31 | 1.81 | 1.18 | 0.62 | 0.02460827 |

| | | | | | | | |
|---|---|---|---|---|---|---|---|
| 4 | NDUFB9 | 0.29 | 0.21 | 0.87 | 0.66 | 0.40 | 0.02561679 |
| 4 | SLK | 0.27 | 0.20 | 0.92 | 0.67 | 0.45 | 0.02581951 |
| 4 | NEMF | 0.25 | 0.18 | 0.93 | 0.58 | 0.69 | 0.02641916 |
| 4 | KRAS | 0.44 | 0.36 | 1.82 | 1.30 | 0.49 | 0.02647177 |
| 4 | SETD2 | 0.47 | 0.39 | 2.05 | 1.57 | 0.39 | 0.02684038 |
| 4 | CELF1 | 0.34 | 0.27 | 1.25 | 0.91 | 0.46 | 0.02698476 |
| 4 | CDC26 | 0.31 | 0.22 | 0.95 | 0.71 | 0.41 | 0.02753176 |
| 4 | C4orf3 | 0.38 | 0.30 | 1.48 | 1.10 | 0.42 | 0.02776644 |
| 4 | CD58 | 0.27 | 0.18 | 0.79 | 0.60 | 0.40 | 0.02874781 |
| 4 | SKAP1 | 0.31 | 0.23 | 1.12 | 0.79 | 0.51 | 0.02895219 |
| 4 | MTHFD2 | 0.30 | 0.22 | 0.95 | 0.73 | 0.38 | 0.02944947 |
| 4 | TAF15 | 0.36 | 0.28 | 1.28 | 0.97 | 0.40 | 0.03144895 |
| 4 | FERMT3 | 0.29 | 0.22 | 1.02 | 0.74 | 0.47 | 0.03200437 |
| 4 | JPT1 | 0.33 | 0.26 | 1.26 | 0.90 | 0.48 | 0.0325232 |
| 4 | FYB1 | 0.40 | 0.31 | 1.74 | 1.33 | 0.39 | 0.03434884 |
| 4 | RAC1 | 0.56 | 0.48 | 2.57 | 1.96 | 0.39 | 0.03457494 |
| 4 | SACM1L | 0.41 | 0.34 | 1.57 | 1.20 | 0.39 | 0.03474303 |
| 4 | ACTR2 | 0.52 | 0.45 | 2.36 | 1.78 | 0.40 | 0.03511883 |
| 4 | NDUFA3 | 0.26 | 0.19 | 0.88 | 0.66 | 0.41 | 0.0357787 |
| 4 | M6PR | 0.33 | 0.25 | 1.10 | 0.82 | 0.42 | 0.03615685 |
| 4 | PHB2 | 0.33 | 0.25 | 1.05 | 0.77 | 0.44 | 0.03702691 |
| 4 | KCTD20 | 0.29 | 0.22 | 0.95 | 0.71 | 0.41 | 0.0374375 |
| 4 | RAB5C | 0.28 | 0.21 | 0.90 | 0.63 | 0.52 | 0.03861334 |
| 4 | TRGC1 | 0.48 | 0.40 | 2.59 | 1.87 | 0.48 | 0.04071784 |
| 4 | TANK | 0.46 | 0.37 | 2.07 | 1.59 | 0.38 | 0.04145336 |
| 4 | MED28 | 0.25 | 0.18 | 0.74 | 0.56 | 0.42 | 0.04184654 |
| 4 | AHR | 0.31 | 0.24 | 1.39 | 0.96 | 0.53 | 0.04258731 |
| 4 | RAC2 | 0.62 | 0.57 | 3.80 | 2.96 | 0.36 | 0.04454672 |
| 4 | NKTR | 0.51 | 0.45 | 2.85 | 2.02 | 0.50 | 0.04519438 |
| 4 | SLC4A7 | 0.27 | 0.20 | 0.99 | 0.69 | 0.52 | 0.04571954 |
| 4 | CHIC2 | 0.31 | 0.25 | 1.15 | 0.76 | 0.59 | 0.04620798 |
| 4 | DNAJC3 | 0.33 | 0.25 | 1.11 | 0.85 | 0.37 | 0.04744715 |
| 4 | NDUFV2 | 0.26 | 0.20 | 0.81 | 0.61 | 0.42 | 0.04819614 |
| 4 | PRKD3 | 0.27 | 0.20 | 0.84 | 0.65 | 0.37 | 0.04882629 |
| 4 | CAPRIN1 | 0.31 | 0.24 | 0.98 | 0.75 | 0.39 | 0.04969352 |
| 5 | IFIT1 | 0.52 | 0.01 | 3.16 | 0.02 | 7.04 | 2.61E-108 |
| 5 | IFIT3 | 0.72 | 0.03 | 7.54 | 0.14 | 5.74 | 1.60E-76 |
| 5 | MX2 | 0.60 | 0.05 | 5.31 | 0.16 | 5.10 | 2.79E-36 |
| 5 | IFIT2 | 0.52 | 0.04 | 16.62 | 0.23 | 6.20 | 2.07E-35 |
| 5 | IFI44 | 0.52 | 0.04 | 3.19 | 0.12 | 4.74 | 1.54E-31 |
| 5 | MX1 | 0.60 | 0.06 | 4.20 | 0.20 | 4.41 | 4.78E-29 |
| 5 | EPSTI1 | 0.64 | 0.09 | 3.82 | 0.30 | 3.65 | 3.00E-22 |
| 5 | OAS1 | 0.28 | 0.02 | 1.61 | 0.05 | 4.89 | 7.91E-20 |
| 5 | DDX58 | 0.36 | 0.03 | 1.45 | 0.10 | 3.82 | 1.38E-18 |
| 5 | IFI6 | 0.52 | 0.08 | 4.56 | 0.27 | 4.06 | 2.92E-17 |
| 5 | HERC5 | 0.52 | 0.08 | 6.25 | 0.25 | 4.65 | 8.81E-17 |
| 5 | SAMD9L | 0.48 | 0.08 | 2.16 | 0.30 | 2.86 | 1.10E-12 |
| 5 | ISG15 | 0.92 | 0.49 | 32.57 | 3.40 | 3.26 | 1.37E-12 |
| 5 | OAS2 | 0.32 | 0.05 | 1.17 | 0.14 | 3.05 | 1.02E-09 |
| 5 | DDX60L | 0.28 | 0.04 | 1.16 | 0.12 | 3.24 | 3.20E-09 |
| 5 | PARP14 | 0.72 | 0.28 | 4.08 | 1.05 | 1.95 | 4.09E-09 |
| 5 | IFI44L | 0.28 | 0.04 | 2.09 | 0.17 | 3.59 | 1.15E-08 |
| 5 | PLSCR1 | 0.44 | 0.11 | 1.96 | 0.33 | 2.58 | 1.75E-08 |
| 5 | IFITM1 | 0.92 | 0.63 | 13.64 | 4.03 | 1.76 | 4.47E-08 |
| 5 | RNF213 | 0.92 | 0.55 | 10.54 | 3.28 | 1.68 | 7.20E-08 |
| 5 | LY6E | 0.68 | 0.30 | 5.44 | 1.10 | 2.30 | 9.02E-08 |
| 5 | ISG20 | 0.88 | 0.58 | 15.87 | 4.00 | 1.99 | 1.20E-07 |
| 5 | XAF1 | 0.52 | 0.16 | 2.75 | 0.58 | 2.24 | 1.39E-07 |
| 5 | PARP9 | 0.28 | 0.05 | 1.12 | 0.17 | 2.73 | 6.39E-07 |

| | | | | | | | |
|---|---|---|---|---|---|---|---|
| 5 | DRAP1 | 0.76 | 0.40 | 6.10 | 1.56 | 1.97 | 9.25E-07 |
| 5 | ANGEL2 | 0.28 | 0.05 | 0.89 | 0.18 | 2.29 | 9.94E-07 |
| 5 | IRF1-AS1 | 0.52 | 0.18 | 2.67 | 0.66 | 2.01 | 1.15E-06 |
| 5 | TRIM22 | 0.60 | 0.26 | 4.42 | 1.00 | 2.14 | 1.59E-06 |
| 5 | EIF2AK2 | 0.44 | 0.15 | 3.05 | 0.50 | 2.61 | 3.09E-06 |
| 5 | SP110 | 0.52 | 0.19 | 1.99 | 0.66 | 1.60 | 9.22E-06 |
| 5 | KLF13 | 0.68 | 0.33 | 3.49 | 1.19 | 1.55 | 1.79E-05 |
| 5 | HLA-E | 1.00 | 0.93 | 22.27 | 11.84 | 0.91 | 2.60E-05 |
| 5 | SP100 | 0.76 | 0.44 | 4.65 | 1.85 | 1.33 | 2.62E-05 |
| 5 | CCL5 | 0.96 | 0.81 | 44.73 | 21.39 | 1.06 | 2.69E-05 |
| 5 | NT5C3A | 0.52 | 0.22 | 2.44 | 0.67 | 1.87 | 2.81E-05 |
| 5 | SAMD9 | 0.52 | 0.23 | 3.01 | 0.80 | 1.91 | 4.29E-05 |
| 5 | STAT2 | 0.32 | 0.09 | 1.57 | 0.28 | 2.48 | 5.94E-05 |
| 5 | LAP3 | 0.36 | 0.11 | 1.39 | 0.33 | 2.09 | 6.45E-05 |
| 5 | HLA-C | 1.00 | 0.97 | 32.93 | 20.79 | 0.66 | 8.02E-05 |
| 5 | PPM1K | 0.44 | 0.18 | 2.71 | 0.58 | 2.23 | 0.00010072 |
| 5 | SYTL3 | 0.88 | 0.61 | 10.39 | 3.93 | 1.40 | 0.00010098 |
| 5 | PSME1 | 0.84 | 0.53 | 4.76 | 2.37 | 1.01 | 0.00010344 |
| 5 | UBTF | 0.36 | 0.12 | 1.37 | 0.39 | 1.82 | 0.00012609 |
| 5 | ZBP1 | 0.28 | 0.08 | 1.08 | 0.27 | 2.01 | 0.0001409 |
| 5 | PML | 0.32 | 0.10 | 1.10 | 0.28 | 1.96 | 0.00015083 |
| 5 | TYROBP | 0.92 | 0.75 | 9.86 | 5.47 | 0.85 | 0.00019845 |
| 5 | IFITM3 | 0.68 | 0.39 | 5.88 | 1.91 | 1.62 | 0.00020583 |
| 5 | RBIS | 0.56 | 0.26 | 2.02 | 0.88 | 1.21 | 0.00033347 |
| 5 | LAG3 | 0.32 | 0.11 | 2.00 | 0.43 | 2.23 | 0.00035011 |
| 5 | B2M | 1.00 | 1.00 | 143.08 | 97.12 | 0.56 | 0.00038179 |
| 5 | TRAPPC2L | 0.28 | 0.09 | 0.94 | 0.25 | 1.92 | 0.00040828 |
| 5 | TRANK1 | 0.28 | 0.09 | 1.17 | 0.27 | 2.12 | 0.00041419 |
| 5 | BST2 | 0.64 | 0.37 | 3.26 | 1.50 | 1.11 | 0.0004723 |
| 5 | XIST | 0.48 | 0.19 | 2.38 | 1.22 | 0.97 | 0.00047774 |
| 5 | ADAR | 0.56 | 0.34 | 3.50 | 1.23 | 1.51 | 0.0005099 |
| 5 | REEP5 | 0.56 | 0.28 | 2.01 | 0.92 | 1.12 | 0.000617 |
| 5 | RARS2 | 0.32 | 0.11 | 1.21 | 0.36 | 1.74 | 0.00063238 |
| 5 | DUSP12 | 0.28 | 0.09 | 0.80 | 0.25 | 1.65 | 0.00071275 |
| 5 | NDUFB1 | 0.48 | 0.23 | 2.29 | 0.80 | 1.51 | 0.0008769 |
| 5 | PRCC | 0.28 | 0.10 | 1.08 | 0.26 | 2.06 | 0.00110184 |
| 5 | LNPEP | 0.60 | 0.32 | 2.49 | 1.25 | 1.00 | 0.00125106 |
| 5 | CCL4 | 0.72 | 0.55 | 80.84 | 22.48 | 1.85 | 0.0013257 |
| 5 | PLAAT4 | 0.60 | 0.35 | 2.83 | 1.40 | 1.02 | 0.00141666 |
| 5 | HSH2D | 0.48 | 0.24 | 2.59 | 0.96 | 1.44 | 0.00146336 |
| 5 | SLFN5 | 0.52 | 0.28 | 3.46 | 1.32 | 1.39 | 0.00156463 |
| 5 | CCL4L2 | 0.48 | 0.26 | 30.68 | 6.69 | 2.20 | 0.00203793 |
| 5 | BTN2A1 | 0.40 | 0.18 | 1.53 | 0.57 | 1.43 | 0.00204582 |
| 5 | CCL3L1 | 0.32 | 0.13 | 9.41 | 1.01 | 3.23 | 0.00207679 |
| 5 | CWC22 | 0.28 | 0.10 | 0.99 | 0.32 | 1.64 | 0.00211218 |
| 5 | PSMB9 | 0.60 | 0.36 | 4.23 | 1.42 | 1.57 | 0.00216195 |
| 5 | ITGAE | 0.36 | 0.16 | 1.80 | 0.57 | 1.65 | 0.00217013 |
| 5 | MED29 | 0.28 | 0.10 | 0.91 | 0.33 | 1.48 | 0.00218253 |
| 5 | STAT1 | 0.32 | 0.13 | 1.91 | 0.42 | 2.19 | 0.00248674 |
| 5 | SLA | 0.68 | 0.43 | 4.34 | 2.23 | 0.96 | 0.00248713 |
| 5 | ADD3 | 0.40 | 0.18 | 1.41 | 0.61 | 1.20 | 0.00250908 |
| 5 | GNG5 | 0.52 | 0.31 | 2.27 | 1.05 | 1.11 | 0.00284682 |
| 5 | KLRC4 | 0.40 | 0.18 | 1.32 | 0.64 | 1.03 | 0.00293582 |
| 5 | FIP1L1 | 0.36 | 0.16 | 1.28 | 0.47 | 1.44 | 0.0029733 |
| 5 | STRN3 | 0.28 | 0.10 | 0.75 | 0.27 | 1.46 | 0.00297763 |
| 5 | TXNDC15 | 0.28 | 0.11 | 1.16 | 0.33 | 1.81 | 0.00307725 |
| 5 | TMSB10 | 1.00 | 0.96 | 42.34 | 21.74 | 0.96 | 0.00310679 |
| 5 | ANXA5 | 0.44 | 0.22 | 2.18 | 0.82 | 1.41 | 0.00327448 |
| 5 | AUTS2 | 0.44 | 0.22 | 2.18 | 0.83 | 1.39 | 0.00337196 |

| | | | | | | | |
|---|---|---|---|---|---|---|---|
| 5 | IRF9 | 0.32 | 0.13 | 1.36 | 0.44 | 1.63 | 0.00342307 |
| 5 | SAMD3 | 0.68 | 0.49 | 4.73 | 2.47 | 0.93 | 0.00344494 |
| 5 | CCDC85B | 0.60 | 0.37 | 3.05 | 1.47 | 1.05 | 0.00396113 |
| 5 | MT2A | 0.68 | 0.48 | 18.13 | 6.27 | 1.53 | 0.00401804 |
| 5 | NKG7 | 0.96 | 0.92 | 33.29 | 23.17 | 0.52 | 0.00435568 |
| 5 | TMEM123 | 0.56 | 0.37 | 3.74 | 1.53 | 1.29 | 0.00442176 |
| 5 | MYL12A | 0.88 | 0.68 | 6.59 | 4.93 | 0.42 | 0.0046333 |
| 5 | IFITM2 | 1.00 | 0.85 | 16.20 | 10.75 | 0.59 | 0.00496123 |
| 5 | ATP5MC2 | 0.72 | 0.57 | 4.66 | 2.65 | 0.82 | 0.00568215 |
| 5 | IRF7 | 0.40 | 0.21 | 2.31 | 0.77 | 1.59 | 0.00591938 |
| 5 | PMAIP1 | 0.60 | 0.36 | 6.58 | 2.24 | 1.55 | 0.00637191 |
| 5 | CYBC1 | 0.28 | 0.11 | 1.00 | 0.39 | 1.34 | 0.00657246 |
| 5 | SRSF1 | 0.36 | 0.16 | 0.96 | 0.47 | 1.01 | 0.00690884 |
| 5 | MED15 | 0.36 | 0.17 | 1.09 | 0.51 | 1.10 | 0.00719508 |
| 5 | RNMT | 0.56 | 0.39 | 3.16 | 1.47 | 1.10 | 0.0074314 |
| 5 | TXNIP | 0.80 | 0.59 | 8.64 | 5.50 | 0.65 | 0.00817275 |
| 5 | JAK3 | 0.28 | 0.12 | 1.23 | 0.36 | 1.78 | 0.00830034 |
| 5 | GZMB | 0.88 | 0.69 | 20.16 | 14.05 | 0.52 | 0.00830315 |
| 5 | KLF9 | 0.28 | 0.11 | 1.00 | 0.40 | 1.32 | 0.00853567 |
| 5 | SEPTIN7 | 0.88 | 0.74 | 6.95 | 4.66 | 0.58 | 0.00891406 |
| 5 | C1orf21 | 0.48 | 0.28 | 2.08 | 1.05 | 0.99 | 0.0089335 |
| 5 | CD47 | 0.48 | 0.28 | 2.43 | 1.00 | 1.29 | 0.00896098 |
| 5 | SECISBP2L | 0.28 | 0.12 | 0.87 | 0.36 | 1.28 | 0.00954935 |
| 5 | GZMA | 0.60 | 0.45 | 13.01 | 5.72 | 1.19 | 0.00958919 |
| 5 | PPP2R5C | 0.76 | 0.59 | 6.08 | 3.69 | 0.72 | 0.00967723 |
| 5 | CCL3 | 0.52 | 0.34 | 34.95 | 7.37 | 2.24 | 0.01026725 |
| 5 | SH2D2A | 0.52 | 0.31 | 2.44 | 1.30 | 0.91 | 0.01063901 |
| 5 | PARP8 | 0.56 | 0.38 | 3.25 | 1.51 | 1.10 | 0.01066448 |
| 5 | GBP4 | 0.44 | 0.25 | 2.31 | 1.02 | 1.18 | 0.01067814 |
| 5 | ME2 | 0.32 | 0.15 | 1.05 | 0.50 | 1.08 | 0.01071643 |
| 5 | RPA2 | 0.28 | 0.12 | 0.97 | 0.37 | 1.38 | 0.0107429 |
| 5 | GOLGA8B | 0.36 | 0.19 | 1.43 | 0.62 | 1.22 | 0.0111486 |
| 5 | ZNF652 | 0.28 | 0.12 | 1.10 | 0.41 | 1.42 | 0.01130212 |
| 5 | EVL | 0.52 | 0.33 | 3.42 | 1.44 | 1.25 | 0.01134767 |
| 5 | GFOD1 | 0.40 | 0.22 | 1.97 | 0.76 | 1.38 | 0.01163881 |
| 5 | IL18R1 | 0.36 | 0.19 | 2.22 | 0.73 | 1.60 | 0.01170651 |
| 5 | TNFSF10 | 0.32 | 0.16 | 2.72 | 0.64 | 2.08 | 0.01191496 |
| 5 | OASL | 0.48 | 0.34 | 9.53 | 1.49 | 2.67 | 0.01232207 |
| 5 | SHFL | 0.36 | 0.20 | 1.50 | 0.64 | 1.24 | 0.01275841 |
| 5 | CHMP5 | 0.28 | 0.13 | 1.49 | 0.39 | 1.95 | 0.0133866 |
| 5 | PRKACB | 0.52 | 0.27 | 1.59 | 1.05 | 0.60 | 0.01364711 |
| 5 | PLCG2 | 0.32 | 0.16 | 1.73 | 0.54 | 1.69 | 0.01364985 |
| 5 | GSTK1 | 0.36 | 0.21 | 1.74 | 0.67 | 1.38 | 0.01547689 |
| 5 | MACROH2A | 0.40 | 0.21 | 1.37 | 0.64 | 1.10 | 0.01582996 |
| 5 | DHRS7 | 0.48 | 0.30 | 2.39 | 1.11 | 1.10 | 0.01626586 |
| 5 | SNHG3 | 0.32 | 0.16 | 1.08 | 0.47 | 1.19 | 0.01640023 |
| 5 | ZBTB38 | 0.36 | 0.19 | 1.56 | 0.72 | 1.12 | 0.01652054 |
| 5 | ARL4C | 0.84 | 0.68 | 8.56 | 5.95 | 0.52 | 0.01663925 |
| 5 | AAK1 | 0.60 | 0.44 | 3.18 | 1.86 | 0.77 | 0.0168149 |
| 5 | RER1 | 0.28 | 0.13 | 1.19 | 0.40 | 1.57 | 0.0169295 |
| 5 | PSMB2 | 0.40 | 0.21 | 1.21 | 0.66 | 0.87 | 0.01698422 |
| 5 | PSMC3 | 0.28 | 0.13 | 0.91 | 0.38 | 1.27 | 0.01700811 |
| 5 | IFI16 | 0.72 | 0.62 | 6.53 | 3.76 | 0.79 | 0.01737506 |
| 5 | GON4L | 0.36 | 0.19 | 1.39 | 0.62 | 1.16 | 0.01832576 |
| 5 | NEDD8 | 0.44 | 0.27 | 2.03 | 0.91 | 1.17 | 0.01890343 |
| 5 | COX7C | 0.88 | 0.76 | 6.30 | 4.57 | 0.46 | 0.01939348 |
| 5 | RNF149 | 0.48 | 0.34 | 2.95 | 1.32 | 1.16 | 0.01964138 |
| 5 | TRIM56 | 0.28 | 0.13 | 1.00 | 0.43 | 1.20 | 0.01987152 |
| 5 | N4BP1 | 0.44 | 0.28 | 2.41 | 1.04 | 1.22 | 0.02035313 |

| | | | | | | | |
|---|---|---|---|---|---|---|---|
| 5 | UBE2J1 | 0.28 | 0.14 | 1.80 | 0.40 | 2.17 | 0.02059144 |
| 5 | TOB1 | 0.40 | 0.21 | 1.78 | 0.94 | 0.92 | 0.02125886 |
| 5 | PDCD7 | 0.40 | 0.23 | 1.47 | 0.74 | 0.98 | 0.02264428 |
| 5 | LINC01871 | 0.60 | 0.45 | 6.45 | 3.14 | 1.04 | 0.0232307 |
| 5 | DAZAP2 | 0.48 | 0.34 | 2.42 | 1.18 | 1.04 | 0.02428588 |
| 5 | PHF11 | 0.28 | 0.14 | 1.25 | 0.46 | 1.45 | 0.02430256 |
| 5 | CEP78 | 0.32 | 0.17 | 1.47 | 0.66 | 1.16 | 0.02456967 |
| 5 | PSMB8 | 0.60 | 0.44 | 2.88 | 1.71 | 0.75 | 0.02532449 |
| 5 | F2R | 0.28 | 0.14 | 1.71 | 0.49 | 1.79 | 0.02756173 |
| 5 | PLCL2 | 0.28 | 0.13 | 0.72 | 0.38 | 0.92 | 0.02786248 |
| 5 | CPNE1 | 0.48 | 0.32 | 2.10 | 1.17 | 0.85 | 0.02866445 |
| 5 | SMDT1 | 0.48 | 0.30 | 1.75 | 0.98 | 0.84 | 0.0292149 |
| 5 | UXT | 0.64 | 0.46 | 2.71 | 1.74 | 0.63 | 0.02938044 |
| 5 | ANKIB1 | 0.28 | 0.14 | 0.83 | 0.40 | 1.05 | 0.02965203 |
| 5 | KLRK1 | 0.28 | 0.13 | 0.82 | 0.41 | 1.01 | 0.02971231 |
| 5 | NFATC2 | 0.36 | 0.22 | 1.47 | 0.70 | 1.07 | 0.03347188 |
| 5 | SH3BGRL3 | 0.84 | 0.76 | 9.32 | 5.93 | 0.65 | 0.03412786 |
| 5 | ETS1 | 0.84 | 0.68 | 6.30 | 4.38 | 0.52 | 0.03415483 |
| 5 | SP3 | 0.36 | 0.21 | 1.35 | 0.68 | 0.98 | 0.03439554 |
| 5 | GBP5 | 0.44 | 0.29 | 2.52 | 1.41 | 0.83 | 0.03494941 |
| 5 | SIPA1 | 0.36 | 0.21 | 1.20 | 0.69 | 0.81 | 0.03619195 |
| 5 | ARMCX3 | 0.44 | 0.31 | 2.27 | 1.11 | 1.03 | 0.03685695 |
| 5 | IK | 0.56 | 0.40 | 2.84 | 1.55 | 0.87 | 0.03798962 |
| 5 | BTN3A1 | 0.28 | 0.15 | 1.27 | 0.50 | 1.35 | 0.0385377 |
| 5 | TMEM87A | 0.28 | 0.14 | 0.91 | 0.43 | 1.09 | 0.03934942 |
| 5 | GLUL | 0.40 | 0.24 | 1.69 | 0.93 | 0.86 | 0.03938409 |
| 5 | EIF3H | 0.56 | 0.46 | 2.88 | 1.77 | 0.71 | 0.03967732 |
| 5 | KLRC2 | 0.52 | 0.36 | 3.80 | 1.84 | 1.04 | 0.03984173 |
| 5 | ARFGAP2 | 0.28 | 0.15 | 1.01 | 0.40 | 1.32 | 0.03993781 |
| 5 | TBRG1 | 0.28 | 0.14 | 1.06 | 0.48 | 1.14 | 0.0420227 |
| 5 | USP16 | 0.40 | 0.24 | 1.41 | 0.82 | 0.78 | 0.04233353 |
| 5 | H2AZ2 | 0.76 | 0.57 | 3.87 | 2.69 | 0.53 | 0.04384626 |
| 5 | S100A4 | 0.72 | 0.64 | 11.46 | 5.60 | 1.03 | 0.04450403 |
| 5 | ABHD2 | 0.48 | 0.33 | 2.24 | 1.32 | 0.77 | 0.04811866 |
| 5 | HADHA | 0.40 | 0.26 | 1.64 | 0.82 | 1.00 | 0.04851733 |
| 5 | ZBTB20 | 0.28 | 0.15 | 1.20 | 0.51 | 1.24 | 0.04906389 |
| 5 | PHF3 | 0.52 | 0.36 | 2.12 | 1.34 | 0.67 | 0.04952757 |

| Table S5 \| Signature genes for gene set analysis | | |
|---|---|---|
| Tumor stress signatures | NK cell activation signatures | GR target genes |
| DNAJB1 | CCL3 | FN1 |
| DNAJB4 | CCL4 | KLF9 |
| DNAJC1 | CCL5 | ANKRD1 |
| DNAJC21 | CXCL13 | MT2A |
| DNAJC3 | CXCL8 | SNAI2 |
| DNAJC7 | GNLY | VIM |
| DNAJC8 | GZMA | ID3 |
| HIF1A | GZMB | POU5F1 |
| HSP90AA1 | GZMH | WNT5A |
| HSPA1A | GZMK | NFKBIA |
| HSPA1B | GZMM | DUSP1 |
| HSPA4 | PRF1 | IRS2 |
| HSPA5 | LTB | RAMP1 |
| HSPA6 | NKG7 | TXNIP |
| HSPA8 | CST7 | CD55 |
| HSPA9 | ZAP70 | RGS2 |
| HSPB1 | TYROBP | GPR183 |
| HSPD1 | XCL1 | FOSL2 |
| HSPE1 | XCL2 | ANXA1 |
| HSPH1 | CSF1 | ZFP36 |
| GADD45A | EOMES | TNFAIP3 |
| GADD45B | IL32 | GADD45A |
| | FCGR3A | GADD45B |
| | FCER1G | BCL6 |
| | IFNG | FKBP5 |
| | | TSC22D3 |

| Table S6 \| DEgenes of Dex treatment | | |
|---|---|---|
| *P* <0.01, \|Log2FC\|>0.5 | | |
| gene name | meidum only vs Dex log2FC | medium only vs Dex p value |
| AREG | 6.296576447 | 2.43834E-10 |
| RN7SL832P | 5.210629067 | 0.001879597 |
| GFPT2 | 4.810620631 | 0.00063653 |
| INSR | 4.461524704 | 0.00167963 |
| MGAM | 4.455843882 | 0.002424043 |
| FKBP5 | 3.941482143 | 3.13686E-35 |
| PCSK2 | 3.598273837 | 0.005597773 |
| FXYD7 | 3.566302999 | 0.004036219 |
| DDIT4 | 3.193880578 | 3.436E-19 |
| CXXC11 | 3.106175043 | 0.003419284 |
| XYLT1 | 3.099577252 | 9.36173E-07 |
| TSC22D3 | 3.053089618 | 9.65441E-14 |
| RASGRP2 | 3.011115083 | 1.13665E-06 |
| BCAT1 | 2.687097696 | 0.007938813 |
| BNC2 | 2.642550028 | 0.000889196 |
| DUSP1 | 2.500157601 | 0.000797199 |
| ZBTB16 | 2.385264276 | 3.39505E-05 |
| SMAP2 | 2.297516335 | 1.24983E-11 |
| RGS1 | 2.253618047 | 4.33317E-05 |
| LEF1 | 2.251291234 | 0.005572039 |
| CXCR2 | 2.2484651 | 3.10227E-05 |
| HPGD | 2.024432499 | 0.000314993 |
| OSBPL5 | 1.97528331 | 4.20811E-07 |
| MMAB | 1.9475172 | 0.00618255 |
| HS3ST3B1 | 1.93584726 | 0.004774214 |
| TLE1 | 1.93405308 | 0.000470141 |
| RAP1GDS1 | 1.893067408 | 9.2697E-06 |
| TXNIP | 1.88384567 | 5.9727E-07 |
| ADAM19 | 1.870519777 | 0.008521582 |
| FAM129A | 1.813705377 | 2.69383E-09 |
| ZBED4 | 1.732642406 | 0.00094722 |
| ANKRD9 | 1.720528185 | 0.009924738 |
| KIR2DL3 | 1.715492261 | 0.003546276 |
| PRDM1 | 1.695643438 | 2.12152E-06 |
| SYTL3 | 1.693898383 | 3.74144E-06 |
| WHSC1 | 1.690150068 | 0.004939435 |
| TMEM173 | 1.684527313 | 0.003996836 |
| LRRC8C | 1.683081692 | 3.10618E-05 |
| OGFRL1 | 1.670464485 | 0.006288269 |
| TUBA4A | 1.64776039 | 0.000507423 |
| RNF130 | 1.647065318 | 0.009131806 |
| SLC1A5 | 1.617881181 | 0.00027273 |
| CD55 | 1.590484603 | 0.000247571 |
| IL6ST | 1.579284368 | 1.13692E-05 |
| KIR2DL4 | 1.572710396 | 0.000486699 |
| KLF9 | 1.571710066 | 0.000122539 |
| SLC7A5 | 1.555160068 | 0.000187089 |
| TMEM2 | 1.513898945 | 1.28525E-05 |
| TNFAIP8 | 1.506216713 | 0.001380955 |
| GPATCH8 | 1.439119777 | 8.50353E-06 |
| CASC7 | 1.422664589 | 5.59182E-05 |
| SPON2 | 1.416311037 | 0.002112859 |
| ELOVL6 | 1.414324879 | 0.003698467 |
| HERPUD1 | 1.41066479 | 0.001838478 |
| TSPAN14 | 1.405532522 | 0.00180366 |
| CCND3 | 1.369386878 | 5.61E-05 |

| | | |
|---|---|---|
| SH3BP5 | 1.358132128 | 0.001009764 |
| SLC1A4 | 1.353165908 | 0.008840062 |
| KLF2 | 1.349741132 | 0.003730295 |
| PIK3IP1 | 1.342086358 | 0.003220173 |
| KIF13B | 1.330435772 | 0.000319171 |
| CEBPB | 1.323560219 | 0.008895788 |
| AGO2 | 1.290440178 | 0.004582372 |
| ETS1 | 1.285697295 | 2.57616E-06 |
| HMGB2 | 1.27501757 | 0.000286778 |
| PXN | 1.270780613 | 0.001104541 |
| DDI2 | 1.25330982 | 0.001248711 |
| ZMYND8 | 1.243122734 | 0.0010387 |
| FOXO1 | 1.237885434 | 0.000782011 |
| NEAT1 | 1.233328536 | 0.007404282 |
| EMB | 1.231671028 | 0.000663748 |
| CYTIP | 1.23035984 | 1.01244E-05 |
| IRF2BPL | 1.229901384 | 0.006007675 |
| FAIM3 | 1.21977315 | 0.006546005 |
| GFOD1 | 1.211068089 | 0.001650821 |
| CNOT6L | 1.197856866 | 0.000343619 |
| XPO6 | 1.184795806 | 0.005605266 |
| SSH2 | 1.18038635 | 0.005768179 |
| CD7 | 1.180004985 | 5.58645E-05 |
| PARP8 | 1.175385705 | 0.000427296 |
| CELF2 | 1.145350859 | 3.29262E-05 |
| SRGN | 1.138981445 | 4.77006E-05 |
| IKZF1 | 1.119689424 | 0.001944128 |
| FGFBP2 | 1.113291549 | 0.001060493 |
| GALNT10 | 1.101496532 | 0.00215501 |
| MCTP2 | 1.098876563 | 0.006155812 |
| CXCR4 | 1.081581807 | 0.005321649 |
| PTGER2 | 1.069660577 | 0.001676092 |
| CHST11 | 1.04482673 | 0.002224091 |
| SYNE1 | 1.04004579 | 0.008691204 |
| ZFP36L2 | 1.035400828 | 0.001879153 |
| TSPYL2 | 1.034668006 | 0.005914345 |
| PPP2R5C | 1.028717733 | 0.003271968 |
| FOXN3 | 1.010675187 | 5.53071E-05 |
| MAPK1 | 1.005633042 | 0.008814597 |
| PIP4K2A | 0.956506703 | 0.00045056 |
| CEP78 | 0.946086124 | 0.009924402 |
| MYH9 | 0.913258503 | 0.001558758 |
| GLIPR1 | 0.909608942 | 0.006330655 |
| HECA | 0.819723272 | 0.008388118 |
| CD53 | 0.797214393 | 0.001126431 |
| CFLAR | 0.742624867 | 0.003985771 |
| CD164 | -0.764441713 | 0.009796686 |
| PLEK | -0.891867954 | 0.001381547 |
| LSP1 | -0.901097271 | 0.000940347 |
| IL2RB | -0.931285195 | 0.001763753 |
| MCL1 | -0.996124628 | 0.001961784 |
| ADAR | -1.03558784 | 0.003849879 |
| TAPBP | -1.051899971 | 0.003905663 |
| BCOR | -1.124108366 | 0.009827168 |
| PPP1CA | -1.141290856 | 0.001209068 |
| BZRAP1-AS1 | -1.147602335 | 0.003159585 |
| PLP2 | -1.155539018 | 0.001775028 |
| IL32 | -1.163965872 | 0.003626731 |
| PSMB9 | -1.166722805 | 0.002444652 |

| | | |
|---|---|---|
| BIRC3 | -1.18358659 | 0.00032543 |
| SP110 | -1.215377951 | 0.009605773 |
| MRPS6 | -1.221096397 | 0.005319444 |
| TOX | -1.261816611 | 0.001108557 |
| PHLDA1 | -1.276841013 | 0.00647335 |
| CD74 | -1.29458757 | 9.52801E-08 |
| PIM2 | -1.301235885 | 0.002497341 |
| SAMD9 | -1.316268308 | 2.88717E-05 |
| SBK1 | -1.328271869 | 0.005418545 |
| GBP5 | -1.333214033 | 0.003395589 |
| DENND3 | -1.355907918 | 0.009604126 |
| GBP4 | -1.371438773 | 0.004218374 |
| PARP14 | -1.381249545 | 0.000654058 |
| BTG2 | -1.389742079 | 0.006502528 |
| FAM105B | -1.396151802 | 0.001728706 |
| IFITM9P | -1.398869877 | 0.008850878 |
| IDH2 | -1.408819543 | 0.00069379 |
| C5orf56 | -1.409503447 | 0.004302722 |
| NAPA | -1.413055306 | 0.003035748 |
| HLA-DRA | -1.420828544 | 0.005525605 |
| NR3C1 | -1.421784961 | 2.11847E-06 |
| ORAI2 | -1.423451041 | 0.006762657 |
| NT5C3A | -1.430685032 | 0.001483797 |
| TRIM22 | -1.432369619 | 0.000372798 |
| RARRES3 | -1.448392546 | 0.000365351 |
| APOL1 | -1.500101941 | 0.005148456 |
| WWC3 | -1.535241989 | 0.000534554 |
| BCL2L1 | -1.550735626 | 0.000637087 |
| CFL1P2 | -1.564452934 | 0.00607379 |
| SLC25A1 | -1.57137887 | 0.00057295 |
| DRAP1 | -1.597391735 | 0.000628067 |
| TMSB4XP6 | -1.602714332 | 0.008364371 |
| SCARB2 | -1.615472663 | 0.008838042 |
| LAP3 | -1.618166504 | 0.005111353 |
| ZCCHC2 | -1.61902681 | 0.000370329 |
| TMEM123 | -1.623891871 | 2.77286E-05 |
| APOL3 | -1.653485831 | 0.000940508 |
| FASLG | -1.687910952 | 0.000150744 |
| PDIA3P | -1.705321357 | 0.009085529 |
| GBP1P1 | -1.711266824 | 0.005283224 |
| TIGIT | -1.713374648 | 5.59994E-05 |
| YIPF5 | -1.732327511 | 0.000988891 |
| PRMT7 | -1.760937691 | 0.003448236 |
| GBP1 | -1.773038796 | 0.000967265 |
| TRIM69 | -1.792082746 | 0.000552177 |
| SQLE | -1.808243274 | 0.001266225 |
| PARP9 | -1.823237402 | 0.000436809 |
| C2orf68 | -1.852354174 | 0.000827396 |
| IFIT5 | -1.858821678 | 0.000950583 |
| XAF1 | -1.863759579 | 1.62993E-06 |
| PARP12 | -1.864883553 | 0.0007554 |
| MYD88 | -1.866505762 | 0.001295288 |
| DDX60 | -1.868742034 | 1.5179E-07 |
| PTMS | -1.874918474 | 0.00233377 |
| SLAMF7 | -1.889713435 | 0.000103635 |
| BZRAP1 | -1.897862215 | 0.000768743 |
| TRIM21 | -1.920347541 | 0.001854435 |
| TRIM5 | -1.924044661 | 0.008265123 |
| EIF2AK2 | -1.929873808 | 1.53539E-06 |

| | | |
|---|---|---|
| EHD4 | -1.937759714 | 0.003098593 |
| BST2 | -1.941321365 | 0.000562132 |
| IRF9 | -1.943105679 | 0.004598973 |
| DDX58 | -1.953335628 | 1.00651E-05 |
| CD226 | -1.961179777 | 0.000604686 |
| OAS2 | -2.003509777 | 0.000218123 |
| IL15RA | -2.012705968 | 0.002199703 |
| DTX3L | -2.018565085 | 8.53993E-07 |
| C15orf39 | -2.018635193 | 0.001852639 |
| TRAF1 | -2.042651184 | 0.00026186 |
| TREX1 | -2.080236633 | 0.003458793 |
| GLIPR2 | -2.10359823 | 1.16464E-05 |
| SAMD9L | -2.106934065 | 4.96556E-06 |
| LAG3 | -2.111930806 | 0.001067625 |
| DDX60L | -2.135618 | 1.39941E-06 |
| LDLR | -2.143091102 | 8.01723E-06 |
| EPSTI1 | -2.148570445 | 2.37236E-06 |
| STAT1 | -2.161079929 | 6.95894E-07 |
| TRDC | -2.183193507 | 3.20996E-09 |
| TNFSF10 | -2.228946464 | 4.65682E-05 |
| RAB37 | -2.258466589 | 0.007607479 |
| NCS1 | -2.301353963 | 0.003080081 |
| RASGEF1B | -2.306283076 | 0.007534367 |
| OASL | -2.30871901 | 6.44055E-07 |
| PARP10 | -2.324360283 | 0.001547255 |
| ADAP1 | -2.345528575 | 0.003178393 |
| XCL2 | -2.373221382 | 0.006944423 |
| BCL3 | -2.39071305 | 6.61343E-06 |
| KLF10 | -2.399640035 | 0.006458489 |
| MT2A | -2.401331232 | 1.67214E-05 |
| HERC6 | -2.408574889 | 2.46146E-05 |
| HERC5 | -2.433649349 | 0.008330399 |
| ARHGAP31 | -2.440230856 | 0.007816143 |
| AC092580.4 | -2.455428631 | 0.001905575 |
| PLSCR1 | -2.491392531 | 3.17409E-07 |
| TCF7 | -2.568608781 | 0.000185699 |
| RTP4 | -2.609586086 | 8.05361E-05 |
| SPATS2L | -2.612854865 | 8.02755E-06 |
| RELB | -2.630771271 | 2.9636E-05 |
| RGS3 | -2.630881369 | 4.8194E-06 |
| DHX58 | -2.632976927 | 0.001309923 |
| TRAC | -2.704761607 | 0.000127384 |
| IFIH1 | -2.745083767 | 1.09483E-07 |
| IFI44 | -2.780335784 | 2.4789E-07 |
| IRF7 | -2.8070866 | 0.003173094 |
| CD82 | -2.817001559 | 2.30581E-08 |
| USP49 | -2.818025032 | 0.002303572 |
| USP18 | -2.830639001 | 0.004535263 |
| ITM2C | -2.859268143 | 0.00275393 |
| IFI6 | -2.960413609 | 0.000171466 |
| TNFSF14 | -2.97007046 | 2.12737E-06 |
| DNPH1 | -3.030539445 | 0.001750866 |
| LINC00299 | -3.229579982 | 0.000897108 |
| OAS1 | -3.380277713 | 0.004505519 |
| DENND5A | -3.422267882 | 0.005219281 |
| CTC-512J14.7 | -3.437789055 | 0.002728398 |
| OAS3 | -3.440471394 | 0.000942227 |
| MX1 | -3.470887319 | 0.000994382 |
| IFI44L | -3.50980718 | 0.001528504 |

| | | |
|---|---|---|
| C1orf61 | -3.548460774 | 0.005407629 |
| CMPK2 | -3.612874475 | 0.000209733 |
| RSAD2 | -3.627942927 | 5.8695E-05 |
| TACO1 | -3.751368843 | 0.000381598 |
| ETV7 | -3.903340752 | 0.005862492 |
| IFIT2 | -3.929901123 | 0.000105113 |
| IFIT3 | -4.102226081 | 0.004097288 |
| IL1RN | -4.159617746 | 0.009471023 |
| TTC26 | -4.390952147 | 0.003377369 |
| USP30-AS1 | -4.420994538 | 0.006471802 |
| IFIT1 | -4.814855238 | 0.000142683 |
| N4BP3 | -4.828458234 | 0.001181234 |
| TMEM78 | -4.860267527 | 0.005092175 |
| C1orf173 | -6.649298267 | 4.06286E-06 |
| SAMD7 | -7.678600858 | 0.009504592 |

| gene name | cytokine vs cytokine + Dex log2FC | cytokine vs cytokine + Dex p value |
|---|---|---|
| AREG | 5.043843254 | 3.36739E-12 |
| SPON2 | 4.848799958 | 7.26609E-24 |
| THAP8 | 4.813189072 | 0.006493324 |
| FFAR2 | 4.707601102 | 0.004903609 |
| CXCR4 | 4.608253294 | 3.55679E-30 |
| CH25H | 3.773403098 | 0.005386792 |
| TSC22D3 | 3.702040143 | 9.02038E-18 |
| DUSP1 | 3.575363024 | 1.57628E-05 |
| PIK3IP1 | 3.572319061 | 1.77702E-13 |
| FKBP5 | 3.538799548 | 2.2646E-33 |
| RASGRP2 | 2.967590491 | 1.47768E-05 |
| KLF9 | 2.789895271 | 5.48421E-12 |
| TLE1 | 2.747363259 | 0.000105104 |
| TRABD2A | 2.651203188 | 0.000511987 |
| FSD1 | 2.638247004 | 0.00663685 |
| KLF2 | 2.569273245 | 6.07337E-08 |
| ELOVL6 | 2.455060394 | 6.29295E-09 |
| VAV3 | 2.3097256 | 3.23814E-06 |
| AGPAT4 | 2.301752404 | 1.66535E-05 |
| YPEL1 | 2.230359712 | 0.000202741 |
| CCNG2 | 2.110140163 | 0.004818035 |
| HPGD | 2.106494426 | 7.00425E-05 |
| ENPP5 | 2.084805698 | 0.009358099 |
| FCRL6 | 2.013421563 | 0.000318776 |
| LPAR6 | 1.972106227 | 0.000128157 |
| BNC2 | 1.964331499 | 0.009778585 |
| TMEM173 | 1.91699234 | 0.000399592 |
| ZFP36 | 1.875760039 | 2.26459E-05 |
| ZFP36L2 | 1.834388855 | 2.99747E-08 |
| TXNIP | 1.830091369 | 8.77489E-07 |
| GLUL | 1.812156841 | 1.71338E-05 |
| SMAP2 | 1.796626511 | 7.31404E-09 |
| FAM115C | 1.794554063 | 0.000616337 |
| ABCB1 | 1.718080935 | 0.000125101 |
| FAM102A | 1.703086787 | 0.005059418 |
| PLEKHG3 | 1.684609654 | 0.004337878 |
| PARP8 | 1.651194758 | 5.09886E-07 |
| FAIM3 | 1.648022122 | 0.001682198 |
| RCBTB2 | 1.630075765 | 0.000781824 |
| TMCC3 | 1.615789034 | 0.000607999 |

| | | |
|---|---|---|
| ABLIM1 | 1.612482096 | 0.003416037 |
| MAPK1 | 1.583055597 | 1.95916E-05 |
| AIM1 | 1.575119118 | 0.00083424 |
| OGFRL1 | 1.566820022 | 0.008608822 |
| BIN1 | 1.564970922 | 0.000462625 |
| SLAMF6 | 1.535265091 | 0.000657431 |
| OSBPL5 | 1.473839868 | 0.000100317 |
| RGS1 | 1.45220068 | 0.002964234 |
| DYRK2 | 1.446385179 | 0.001285707 |
| CARD11 | 1.411138483 | 1.13551E-05 |
| TRERF1 | 1.393726502 | 0.00095667 |
| TSPAN5 | 1.386439518 | 0.006302498 |
| SYNE1 | 1.33864478 | 0.000637909 |
| PXN | 1.326836223 | 0.000178389 |
| SSH2 | 1.299691789 | 0.00164886 |
| GLCCI1 | 1.225573494 | 0.003559196 |
| TUBA4A | 1.207663141 | 0.003258579 |
| PARP4 | 1.197142061 | 0.000338119 |
| FAM65B | 1.189896168 | 0.00013488 |
| SELPLG | 1.16553563 | 0.003697083 |
| CD55 | 1.146242741 | 0.001153328 |
| BTG1 | 1.116915067 | 0.000804325 |
| CBFB | 1.104993401 | 8.20273E-05 |
| TMEM2 | 1.099635418 | 0.00079159 |
| NR1D2 | 1.050811873 | 0.002787404 |
| KIAA1551 | 1.042499258 | 0.000547079 |
| ST3GAL1 | 1.032815252 | 0.001970142 |
| SMARCA2 | 1.027942786 | 0.003310796 |
| FOXO1 | 1.020538508 | 0.003642932 |
| EVI2B | 1.008741026 | 0.003864572 |
| POLR3GL | 0.991723248 | 0.008760098 |
| CASP8 | 0.989761136 | 0.005283292 |
| CCDC69 | 0.986523867 | 0.006935976 |
| TMEM66 | 0.985719425 | 0.008923417 |
| DDIT4 | 0.984699198 | 0.00149051 |
| SSH1 | 0.905089486 | 0.004251249 |
| CD7 | 0.861770693 | 0.001666199 |
| SRGN | 0.780863699 | 0.004270824 |
| MSN | 0.694713635 | 0.001323824 |
| SLC38A1 | 0.59183351 | 0.009950435 |
| ID2 | -0.738090434 | 0.00532609 |
| RBPJ | -0.807975257 | 0.002049489 |
| CCL5 | -0.813858337 | 0.005261917 |
| VASP | -0.830544635 | 0.009460254 |
| SH2D2A | -0.856659669 | 0.002694664 |
| IL2RB | -0.859259968 | 0.00183395 |
| PRKX | -0.861891676 | 0.00355197 |
| CMTM6 | -0.908983075 | 0.00535513 |
| PLEK | -0.910009723 | 0.000548299 |
| CD44 | -0.945516299 | 0.006238405 |
| TMSB4XP4 | -0.956683124 | 0.000392379 |
| TGFBR3 | -0.990688091 | 0.003387914 |
| ZFP36L1 | -1.080145651 | 0.001873318 |
| GADD45B | -1.10083103 | 0.000650396 |
| TRGC1 | -1.156981249 | 0.000893749 |
| REL | -1.159243573 | 0.001370546 |
| TUBBP1 | -1.160918171 | 0.002663144 |
| PRR5L | -1.173545761 | 0.0022813 |
| STAT5A | -1.206841254 | 0.003312977 |

| | | |
|---|---|---|
| BHLHE40 | -1.215610187 | 2.53085E-05 |
| TRGC2 | -1.257616668 | 0.001043913 |
| MAP3K8 | -1.26717728 | 0.000341644 |
| PDE4B | -1.269866385 | 0.006678497 |
| BATF | -1.289809275 | 0.007509247 |
| IER3 | -1.308853624 | 0.009123558 |
| CREM | -1.310207068 | 0.001149936 |
| VIM | -1.337127623 | 7.5426E-05 |
| PTMS | -1.339149087 | 0.009609039 |
| TNFSF14 | -1.349395791 | 0.00942813 |
| IRF2BP2 | -1.354792299 | 4.58531E-05 |
| FRMD4B | -1.359049377 | 0.006682612 |
| TIGIT | -1.404863917 | 0.000498991 |
| BCL6 | -1.406273859 | 0.000549132 |
| PDE4A | -1.412096383 | 0.000460493 |
| TRDC | -1.412497115 | 5.74336E-05 |
| NFAT5 | -1.418190046 | 0.001214154 |
| TNFAIP8 | -1.422097892 | 0.000800025 |
| FOXP4 | -1.446555187 | 0.001337341 |
| ITGA1 | -1.462599114 | 0.005170582 |
| KDM6B | -1.529372143 | 0.001415732 |
| SOCS3 | -1.628302447 | 0.006005922 |
| JUNB | -1.633450476 | 6.03898E-07 |
| LTB | -1.656557958 | 0.001985663 |
| LGALS1 | -1.656679886 | 0.001015806 |
| TNFSF10 | -1.664170271 | 0.000755773 |
| NCS1 | -1.751781511 | 0.00144436 |
| TRAF1 | -1.760168533 | 0.000261394 |
| TRAC | -1.760771818 | 0.007718554 |
| TNFRSF18 | -1.818785405 | 0.000464477 |
| SATB1 | -1.861145419 | 1.58267E-07 |
| TPM3P8 | -1.943909229 | 0.006881615 |
| FEZ1 | -2.045040708 | 0.000145393 |
| RGS16 | -2.065659343 | 9.02523E-10 |
| TNF | -2.156979893 | 2.27488E-06 |
| CD83 | -2.217384016 | 2.06317E-08 |
| CD274 | -2.226145361 | 0.000120979 |
| MFSD2A | -2.311035708 | 0.004687912 |
| CCR7 | -2.324021166 | 0.001689433 |
| BCL2A1 | -2.341071025 | 0.00048591 |
| CD82 | -2.381449736 | 5.2578E-08 |
| TMEM217 | -2.440233784 | 0.009525414 |
| EGR2 | -2.48394869 | 0.000126826 |
| RGCC | -2.488755814 | 0.002640976 |
| CSF1 | -2.493895707 | 2.73335E-05 |
| ADAM19 | -2.59074428 | 1.51552E-08 |
| DUSP4 | -2.66846923 | 1.51835E-09 |
| GRAMD1B | -2.681751436 | 0.000506191 |
| TNFRSF9 | -2.975052357 | 1.29189E-06 |
| BATF3 | -3.009516493 | 0.000441672 |
| HBEGF | -3.141112394 | 0.000234841 |
| LTA | -3.275528828 | 3.00389E-16 |
| TNFRSF4 | -3.328205338 | 5.19627E-07 |
| XCL1 | -3.431044241 | 1.89323E-05 |
| XCL2 | -3.586179936 | 1.32689E-05 |
| IL1RN | -3.895222537 | 0.000501944 |
| GPR183 | -5.519477275 | 8.88639E-08 |
